# Discovery and validation of candidate smoltification gene expression biomarkers across multiple species and ecotypes of Pacific salmonids

**DOI:** 10.1101/474692

**Authors:** Aimee Lee S. Houde, Oliver P. Günther, Jeffrey Strohm, Tobi J. Ming, Shaorong Li, Karia H. Kaukinen, David A. Patterson, Anthony P. Farrell, Scott G. Hinch, Kristina M. Miller

## Abstract

Early marine survival of juvenile salmon is intimately associated with their physiological condition during ocean entry and especially smoltification. Smoltification is a developmental parr–smolt transformation allowing salmon to acquire the trait of seawater tolerance in preparation for marine living. Traditionally, this developmental process has been monitored using gill Na^+^/K^+^ATPase (NKA) activity or plasma hormones, but gill gene expression can be reliably used. Here, we describe the discovery of candidate genes from gill tissue for staging smoltification using comparisons of microarray studies with particular focus on the commonalities between anadromous Rainbow trout and Sockeye salmon datasets, as well as literature comparison encompassing more species. A subset of 37 candidate genes mainly from the microarray analyses was used for Taq-Man qPCR assay design and their monthly expression patterns were validated using gill samples from four groups, representing three species and two ecotypes: Coho salmon, Sockeye salmon, stream-type Chinook salmon, and ocean-type Chinook salmon. The best smoltification biomarkers, as measured by consistent changes across these four groups, were genes involved in ion regulation, oxygen transport, and immunity. Smoltification gene expression patterns (using the top 10 biomarkers) were confirmed by significant correlations with NKA activity and were associated with changes in body brightness, caudal fin darkness, and caudal peduncle length. We incorporate gene expression patterns of pre-smolt, smolt, and de-smolt trials from acute seawater transfers using a companion study to develop a preliminary seawater tolerance classification model for ocean-type Chinook salmon. This work demonstrates the potential of gene expression biomarkers to stage smoltification and classify juveniles as pre-smolt, smolt, or de-smolt.

## 1. Introduction

Beyond their cultural importance, salmonids can provide over a billion dollars annually to the economies of countries with recreational and commercial fisheries (e.g. Canada, Pinfold, 2011). Yet, populations of several salmonid species are declining on the Pacific and Atlantic coasts, and lower early marine survival of juveniles is associated with these declines (Beamish et al., 2010; Friedland et al., 2003; Mills et al., 2013). To increase salmonid populations and augment fisheries, hatchery breeding programs are used (Fraser, 2008). As well, aquaculture is used to alleviate some of the fishing pressure on wild populations (Naylor et al., 2000) and provide additional economic opportunities (Bostock et al., 2010). However, the success of both hatcheries and aquaculture is known to be limited by the physiological condition of the smolt life stage during the transition from freshwater to seawater (e.g. Chittenden et al., 2008; Stien et al., 2013). Consequently tools to measure the physiological condition of smolts are routinely used and improvements in them sought to inform culture and decisions for optimizing smolt performance.

All salmonid species begin their lives in freshwater as eggs, alevins, and fry, and then the juvenile anadromous forms become smolts to successfully outmigrate to seawater, where rapid bodily growth and increased reproductive success are greatly improved over freshwater residence. A trade-off may be survival because of increased predation, variable prey availability, and other risks in the marine environment (Quinn, 2005). The developmental process preparing salmonids for the transition from freshwater to marine habitats is termed smoltification or parr–smolt transformation, which is characterized by changes in behaviour, skin pigmentation, body morphology, and physiology (reviewed by McCormick et al., 1998; 2013; Björnsson et al., 2011). Changes in behaviour include increased negative rheotaxis (i.e. downstream movement) and schooling (i.e. the loss of territorial behaviour). The schooling behaviour may lower the risks of predation in river and the early marine environment. Changes in skin pigmentation include acquiring silver skin pigmentation and dark caudal fin tips. Changes in body morphology include a more streamlined body shape, elongation of the caudal peduncle, and associated lower body condition and increased buoyancy. These changes in pigmentation and morphology may be adaptations to marine habitats as camouflage from predators and increased swimming performance in open water, respectively.

The physiological changes during smoltification are equally numerous, such as increased hemoglobin, metabolism, and seawater tolerance. Higher hemoglobin may increase the oxygen-capacity of the blood because seawater is typically lower in dissolved oxygen than freshwater(Seear et al., 2010). Higher metabolism may be to meet the increased energetic demands during smoltification and migration (Robertson and McCormick, 2012). Of the physiological changes, acquired seawater tolerance may be the most important (McCormick et al., 1998; 2013; Björnsson et al., 2011; ?). Indeed, juvenile salmonids that are unprepared for increased salinity, i.e. pre-smolts that have not completed the parr–smolt transformation or de-smolts that have remained in freshwater too long and have reverted to a physiology more suited to freshwater, have greatly reduced survival and slower growth because of internal ionic and osmotic disturbances from the excess ions in seawater relative to freshwater. As a result, smoltification is regarded as a ‘physiological window’ for seawater tolerance, one that can narrow because of higher water temperature, which may have implications with global climate change (e.g. Bassett et al., 2018). Moreover, the physiological window may be altered in the culture relative to natural environment, with hatchery juveniles generally having lower seawater tolerance than wild juveniles (e.g. Shrimpton et al., 1994; Chittenden et al., 2008). Knowing the smolt status of juveniles in particular is critical for hatchery and aquaculture operations to optimize the timing the smolt release directly into seawater. Federal hatcheries guidelines in British Columbia, Canada suggest that the time of release should coincide with that of the wild migration (MacKinlay et al., 2004), but certain hatcheries may have a specific range of dates used every year. Altogether, hatcheries and aquaculture can benefit from tools that reliably measure the smolt status of salmonids for planning releases and modifying the culture environment, if necessary.

In general existing tools take advantage of known changes associated with smotification. For example, salmonids generally need to reach a critical body size prior to smoltification. Photoperiod and, to a lesser extent, temperature also drives smoltification (McCormick et al., 1998; 2013; Björnsson et al., 2011). Since an increase in day length activates the light-brain-pituitary axis to release a cascade of hormones including growth hormone, insulin-like growth factor I, cortisol, and thyroid, these hormones can be monitored in plasma samples. Growth hormone and cortisol stimulate the development of gill ionocytes and their associated Na^+^/K^+^-ATPase (McCormick, 1993; Evans et al., 2005), the activity of which can be monitored in gill samples. Thyroid hormones may be involved in the changes in behaviour and skin pigmentation, which are useful visual indicators of smoltification. All the same, smoltification research has mainly focussed on species and ecotypes that migrate to seawater after one or more years in freshwater, e.g. Coho salmon (*Oncorhynchus kisutch*), stream-type Chinook salmon (*O. tshawytscha*, see Bourret et al., 2016), Sockeye salmon (*O. nerka*), anadromous Rainbow trout (*O. mykiss*), Atlantic salmon (*Salmo salar)*, and Brook trout (*Salvelinus fontinalis*). However, species and ecotypes that migrate to the ocean after less than a year in freshwater: e.g. ocean-type Chinook salmon (*O. tshawytscha*, see Bourret et al., 2016), Pink salmon (*O. gorbuscha*), and Chum salmon (*O. keta*), enter seawater at a smaller body size and may remain longer in estuaries than the other groups. In these species and ecotypes, smoltification may not depend on photoperiod and may be body size based (Clarke et al., 1992; 1994; Gallagher et al., 2013, but see Hoffnagle and Fivizzani, 1998). Thus, tools to define smolt status have focussed on gill Na^+^/K^+^-ATPase activity and plasma hormone concentrations.

Recently, techniques for monitoring smoltification have shifted to candidate gill gene expression using quantitative PCR (qPCR) for hormones and their receptors (e.g. Kiilerich et al., 2007; Hecht et al., 2014), as well as the precursors to Na^+^/K^+^-ATPase (e.g. Nilsen et al., 2007; Piironen et al., 2013). In particular, the gill expression of Na^+^/K^+^-ATPase α-1 isoforms for ‘a’ freshwater and ‘b’ seawater ion regulation (c.f. Richards et al., 2003; Shrimpton et al., 2005), which typically change reciprocally during smoltification, are compared. More recently, smoltification has been examined at the genomic level using microarrays (e.g. Seear et al., 2010; Robertson and McCormick, 2012; Sutherland et al., 2014) which have identified gill expression patterns for the upregulation of biological functions such as ion regulation, metabolism, oxygen transport, growth, structural integrity (e.g. collagen), calcium uptake (i.e. nutrient limitation for growth), and immunity, as well as downregulation of immunity and a few ion regulation and hormones. The upregulation of innate immunity is suggested as a preparation for exposure to new pathogens in marine environments (Boulet et al., 2012), while the downregulation of anti-viral immunity (Sutherland et al., 2014) is suggested to be due to the suppression by cortisol (Lemmetyinen et al., 2013). Despite these recent advances, it is not known if expression patterns of specific genes for smoltification can be reliably applied across salmonid species and different ecotypes.

Therefore, our objective was to discover candidate genes for smoltification and validate a subset of these genes using new samples from multiple species with different ecologies. To this end, we used mapping approaches to discover candidate smoltification genes by a meta-analysis of microarray gene expression patterns across studies. In particular, we focused on a comparison between anadromous Rainbow trout (Sutherland et al., 2014) and in-house Sockeye salmon datasets, as well as mining the literature for a wider collection of salmonid studies based on gene names. We then selected a subset of candidate genes for validation. These genes were developed into TaqMan qPCR assays and tested for expected gene expression patterns using various hatchery and wild sources of gill samples from Coho salmon, Sockeye salmon, stream-type Chinook salmon, and ocean-type Chinook salmon. We used the Fluidigm BioMarkM HD platform for measuring gene expression, a high throughput microfluidics-based technology that can individually quantify 96 assays across 96 samples at once. We focused on these four groups because of their population declines in Southern British Columbia (BC), Canada and subsequent hatchery supplementation (DFO 2013; Beamish et al., 2009; Noakes et al., 2000). In particular, the Sockeye salmon were from the endangered population of Cultus Lake, BC (COSEWIC, 2003).

We hypothesize that a suite of biomarkers will be consistently associated with the smoltification process across species and ecotypes, and the specific level of activation of this smoltification biomarker panel alone could predict smolt status. As such, the present study would mark the first step in a process by identifying biomarkers that change with monthly smolt development. Our companion study examines the gene expression associated with seawater survival using pre-smolt, smolt, and de-smolt juveniles (e.g. oceantype Chinook salmon, Houde et al., 2018). Using the smolt status for the trials of the companion study, here we explore a preliminary seawater tolerance classification model for ocean-type Chinook salmon. We examined how the seawater tolerance changed during monthly development for ocean-type Chinook salmon in the present study.

## 2. Materials and Methods

### 2.1. Candidate smoltification gene discovery

Smoltification candidate genes for gill tissue were identified using two approaches: (1) comparisons between a Sockeye salmon (*Oncorhynchus nerka*) cGRASP 44K internal microarray dataset of the Molecular Genetics Lab, Pacific Biological Station, Nanaimo, BC and the signatures of four external cGRASP microarray studies, i.e. 44K: Sutherland et al. (2014) and 16K: Boulet et al. (2012); Robertson and McCormick (2012); Lemmetyinen et al. (2013), and (2) a literature mining of significant gene names across published studies. Statistical analyses were conducted in R 3.1.2 (R Core Team). Methods for the Sockeye salmon microarray studies are described by Miller et al. 2009; 2011). The Sockeye salmon dataset is composed of 7 parr and 8 smolt samples for 27,104 features. This dataset was filtered with a 50% threshold for missing values and imputation of missing values was performed with the mean value over available samples. The Rainbow trout (*Oncorhynchus mykiss*) dataset (Sutherland et al., 2014) was downloaded from NCBI’s Gene Expression Omnibus (GEO) public repository using the *GEOquery* R package (Sean and Meltzer, 2007) and the processing steps of the authors were honoured.

For the direct comparisons between the internal Sockeye salmon and external microarray datasets, first significant features that separated parr and smolt for the Sockeye salmon dataset were identified using the robust empirical Bayes method of the *limma* R package (Ritchie et al., 2015). Features with a false discovery rate (FDR) < 0.05 were considered significant. Next, to identify the top 100 features that separated parr and smolt for both species, significant features of the Rainbow trout and the Sockeye salmon datasets (both 44K platforms) were combined and analyzed collectively using a sparse independent principal component analysis (sIPCA) with the *mixOmics* R package (Rohart et al., 2017). These 100 features were examined for overlap with the identified significant features from the Sockeye salmon robust limma analysis described above. For the remaining three datasets using the 16K platform, both the 16K and 44K features were mapped to Atlantic salmon (*Salmo salar)* gene IDs from NCBI (see below for details), enabling comparisons across platforms. Similarly, the 16K features were examined for overlap with the identified significant features for both Sockeye salmon and Rainbow trout datasets.

Mining published literature involved discovering the overlap of significant gene names across five microarray studies that used the gill tissue of salmonid fishes, i.e. the four external microarray studies and Seear et al. (2010) that used a TRAITS/SGP microarray. Given the small number of studies and that different gene subunits invariably contribute to a protein, the study tables were visually examined for overlap using generalized gene names. Names that significantly separated parr and smolt in at least two microarray studies were recorded and were organized by smoltification biological function. Additional candidate gene studies (*n* = 5) examining the expression of specific ion regulation, hormone, and hormone receptor genes for gill tissue were also considered.

### 2.2. qPCR assay design

A Microsoft Access relational database containing mRNA sequences of salmonids was produced for qPCR TaqMan assay design, with the objective of developing assays that were gene specific and worked across several species. The database is available from the authors. Until recently, there were limited mRNA sequence data for the genus *Oncorhynchus*, so sequences were generated for Coho (*O. kisutch*), Sockeye (*O. nerk*a), and Chinook (*O. tshawytscha*) salmon from pools of six to eight individuals per species. Samples were enriched for GRASP microarray features using SureSelectXT (Agilent) and then sequenced using IonTorrent (Thermo-Fisher) following the manufacturer kits. Akbarzadeh et al. (2018) provides greater methodological details.

Sequences were mapped to the Atlantic salmon genome using the methods described by Houde et al. (2019). The database also contained maps of microarray features to the Atlantic salmon genome, specifically GRASP (16K, 32K, and 44K), TRAITS (version 1, 2.1, and 2.2) and SIQ. Altogether, database retrieval of mRNA sequences for the three species used microarray features IDs, gene IDs, gene names, or official gene symbols. Available sequences of several salmonid species, i.e. from the database and additional searches of the NCBI repository, were aligned using MEGA7.0.14 (Kumar et al., 2016). Sequences of closely related genes, e.g. duplicate genes, were also included in the alignment. Sequence regions that differed among closely related genes and were conserved among species were used as template in Primer Express 3.0.1 (Thermo Fisher) using the default setting optimized for the BioMark platform. Primer and TaqMan probe combinations that mismatched in at least one base pair at the 3’ end of closely related genes were preferentially selected. Another check of potential gene specificity and workability across salmonids used NCBI Primer-Blast. One or two assays were designed per candidate gene.

### 2.3. Assay efficiencies

Assay design efficiencies were measured using pools of cDNA samples for six to nine salmonid species. Species-specific pools contained a mix of five tissues (gill, liver, heart, kidney, and brain tissue) from several individuals, with the exception of Rainbow trout and Atlantic salmon (only gill tissue). For each species, cDNA was diluted using a five-fold serial dilutions 1 to 1/625. Following Fluidigm BioMark^ℒ^ prescribed methods, target cDNA sequences were enriched using a specific target amplification (STA) method that included small concentrations of the assay primers as well as three housekeeping genes: Coil-P84, 78d16.1, and MrpL40 (Miller et al., 2017). Specifically, for each reaction, 3.76 µL 1X TaqMan PreAmp master mix (Applied Biosystems), 0.2 µM of each of the primers, and 1.24 µL of cDNA. Samples were run on a 14 cycle PCR program, with excess primers removed with EXO-SAP-IT (Affymetrix), and diluted 1 in 5 with DNA suspension buffer.

The sample dilutions were run in duplicate and most assays in singleton following the Fluidigm platform instructions. Specifically, for sample reactions, 3.0 µL 2X TaqMan mastermix (Life Technologies), 0.3 µL 20X GE sample loading reagent, and 2.7 µL STA product. For assay reactions, 3.3 µL 2X assay loading reagent, 0.7 µL DNA suspension buffer, 1.08 µL forward and reverse primers (50 µM), and 1.2 µL probe (10 µM). The PCR was 50 °C for 2 min, 95 °C for 10 min, followed by 40 cycles of 95 °C for 15 s, and then 60 °C for 1 min. Data were extracted using the Real-Time PCR Analysis Software (Fluidigm) using Ct thresholds set manually for each assay. Linear models of the Ct values by the log of the dilutions for each group-assay set were produced using the plyr R package (Wickham, 2011). Efficiency values were calculated using 10^−1/slope – 1^, and assays with values between 0.9 and 1.1 were ideal.

### 2.4. Validation samples

Juveniles representing the four groups (three species and two ecotypes) were collected monthly between November 2015 to May 2016 which spanned the smoltification period at four Salmon Enhancement Program (SEP) hatchery facilities. Nitinat Hatchery and Quinsam Hatchery on Vancouver Island, BC for Coho salmon (*Oncorhynchus kisutch*) and Chinook salmon (ocean-type, *O. tshawytscha*) (Table 1). Inch Creek Hatchery and Chehalis Hatchery on mainland BC, respectively, for Sockeye salmon (*O. nerka*) and Chinook salmon (stream-type, *O. tshawytscha*). In addition, wild (i.e. natural-born) juvenile counterparts of Coho salmon and Sockeye salmon were collected from the hatchery-supplemented source rivers and lakes using baited traps, dip nets, seines, or downstream fences. We targeted 20–30 individuals monthly for each set and the last collection date was as close as possible to the hatchery release date.

**Table 1:** Summary of samples sizes for the four groups collected from four hatcheries and their wild source counterpart. Presented is the number of juveniles of Coho salmon (*Oncorhynchus kisutch*), Sockeye salmon (*O. nerka*), Chinook salmon (stream-type, *O. tshawytscha*), and Chinook salmon (ocean-type, *O. tshawytscha*). Juveniles were collected from freshwater unless denoted by the symbol ‘E’ which denotes juveniles from an estuary where they were exposed to seawater for about two weeks. Digital photograph were collected for the Nitinat and Quinsam juveniles in March, April, and May. Nitinat wild Coho salmon were collected from Campass Creek, a neighbouring tributary of Nitinat River, which was smaller and thus more feasible for catching juveniles with traps than Nitinat River.

|  | November | December | January | February | March | Early April | Late April | May |
| --- | --- | --- | --- | --- | --- | --- | --- | --- |
| **Nitinat Hatchery** |  |  |  |  |  |  |  |  |
| Coho salmon (Nitinat River), age 1+ |  |  |  |  |  |  |  |  |
| Hatchery | 20 | - | 20 | 20 | 20 | 20 | - | 20 |
| Wild | 30 | - | 30 | 30 | 30 | 30 | - | 20 |
| Chinook salmon (Nitinat River), age 0+ |  |  |  |  |  |  |  |  |
| Hatchery | - | - | 20 | 20 | 20 | 20 | 20, 20 E | 20 E |
| Chinook salmon (Sarita River), age 0+ |  |  |  |  |  |  |  |  |
| Hatchery | - | - | 20 | 20 | 20 | 20 | - | 20 |
| **Quinsam Hatchery** |  |  |  |  |  |  |  |  |
| Coho salmon (Quinsam River), age 1+ |  |  |  |  |  |  |  |  |
| Hatchery | 30 | - | 30 | 30 | 30 | 30 | - | 30 |
| Wild | 30 | - | 30 | 35 | 30 | 29 | - | 30 |
| Chinook salmon (Quinsam River), age 0+ |  |  |  |  |  |  |  |  |
| Hatchery | - | - | - | 30 | 30 | 30 | - | 30 |
| **Inch Creek Hatchery** |  |  |  |  |  |  |  |  |
| Sockeye salmon (Cultus Lake), age 1+ |  |  |  |  |  |  |  |  |
| Hatchery | 20 | 20 | 20 | 20 | 20 | 20 | - | - |
| Wild | 20 | - | - | 20 | 17 | 10 | - | - |
| **Chehalis Hatchery** |  |  |  |  |  |  |  |  |
| Chinook salmon (Chilko River), age 1+ |  |  |  |  |  |  |  |  |
| Hatchery | - | 20 | 20 | 19 | 20 | 30 | - | - |
| Chinook salmon (Upper Fraser Summer Red), age 1+ |  |  |  |  |  |  |  |  |
| Hatchery | - | 20 | 20 | 20 | 20 | 20 | - | - |

Fish were euthanized using buffered MS-222 (300 mg L^−1^) then measured for length (± 0.1 cm) and mass (± 0.01 g). Body condition was calculated as 100 x mass ÷ length^3^ (Fulton, 1904). For the months of March, April, and May, Nitinat and Quinsam hatchery and wild juveniles were also digitally photographed (Nikon Coolpix AW110) using a camera stand with a light grey background and a length scale. Photographs were examined for skin pigmentation and body morphology (detailed by Houde et al., 2015) to generate LAB colour space values for anterior, posterior, and caudal fin regions, which were subjected to a principal component analysis, as well as morphology values using 21 landmarks which were subjected to a relative warp analysis using tpsRelw32 software (Rohlf, 2017). Gill tissue from the right side was then placed into a cryovial and immediately frozen with liquid nitrogen or dry ice for Na^+^/K^+^-ATPase activity. Gill tissue from the left side (used for gene expression) was placed into RNAlater (Ambion) for 24 h before freezing or the whole fish was placed into a Whirl-pack bag and then immediately frozen between slabs of dry ice for later gill dissection. Tissues were stored at −80 °C until used for measurements.

### 2.5. Gene expression

We targeted a minimum subset of eight individual fish each month for gill gene expression and measured Na^+^/K^+^-ATPase activity (McCormick, 1993) in around half of these samples. For gene expression, gill tissue was homogenized in TRIzol (Ambion) and BCP reagent using stainless steel beads on a MM301 mixer mill (Retsch Inc.). RNA was extracted from the homogenate using the ‘No-Spin Procedure’ of MagMAX-96 Total RNA Isolation kits (Ambion) and a Biomek FXP automation workstation (Beckman-Coulter). RNA yield was quantified using the A260 value and extracts were normalized to 62.5 ng mL^−1^. Normalized RNA was reverse transcribed to cDNA using SuperScript VILO synthesis kits (Invitrogen). Normalized RNA and cDNA were stored at −80 °C between steps.

Gene expression was quantified using the assays and samples in singleton with STA enriched cDNA and the Fluidigm platform as described above. We included additional assays for candidate genes of thermal and hypoxia stress to assess cross-reactivity with candidate smoltification genes (data available from authors). Each gene expression chip contained three housekeeping genes (i.e. Coil-P84, 78d16.1, and MrpL40, Miller et al., 2017), dilutions of a group-specific cDNA pool, and a group-specific calibrator sample. For determining the optimal normalization gene(s) from the three housekeeping (HK) candidates, gene expression of each HK was first linearly transformed (efficiency minimum textsuperscriptCt – sample Ct). Values were then used in the NormFinder R function (Andersen et al., 2004) with groupings for constituents (e.g. hatchery location) by month to identify the gene or gene pair with the lowest stability (standard deviation). Sample gene expression was normalized with the ΔΔ.Ct method (Livak and Schmittgen, 2001) using the mean (for single gene) or geometric mean (for pair of genes) and the group-specific calibrator sample. Gene expression was then log transformed: log_2_(2^Δ Δct^).

### 2.6. Statistical analysis for validating genes

Candidate smoltification genes were validated using a correlation analysis based on principal components analyses (PCA) across groups and within groups. Analyses were performed using R 3.4.4 at a significance level of α = 0.05. Across the four groups, the expression values of all freshwater monthly gill samples were placed into a single PCA. Loadings and scores were visualized using the *fviz pca function* of the *factoextra* R package (Kassambara and Mundt, 2017). The PC axis best separating earlier and later months was identified. Candidate genes were ranked as biomarkers based on the significance of Pearson correlations between each gene assay and this PC axis. A second PCA and visualization was performed using the top 10 biomarkers with *p* < 0.05. Additional Pearson correlations examined the relationships between gene expression patterns (PC1 and PC2 of the second PCA) and Na^+^/K^+^ATPase activity, as well as body length, mass, condition, morphology, and skin pigmentation. The same approach was used to examine each of the four groups separately. Student’s t-tests also examined gene expression differences for all 37 gene assays between freshwater and seawater samples collected at the same time in late April for Nitinat ocean-type Chinook salmon.

### 2.7. Seawater tolerance classification model

We conducted a companion study using ocean-type Chinook salmon, for which juveniles were exposed to salinity treatments (freshwater, brackish, and seawater) during four trials covering the smoltification period (Houde et al., 2018). We categorized each trial as either pre-smolt, smolt, or de-smolt based on fish survival over several days after acute seawater transfer for a subset of individuals. The PCA pattern for the ocean-type Chinook salmon in the present study was applied to the freshwater juveniles of the companion study. Next, the gene expression PC axis thresholds that best separated three smolt statuses were identified by the maximum of Youden’s J statistic (sensitivity + specificity 1, Youden, 1950) from receiver operating characteristic (ROC) analysis using the *pROC* R package. The resulting thresholds were used to classify fish as seawater tolerant (smolt) or intolerant (pre-smolt and de-smolt). This seawater tolerance classification model was then applied to the constituents of ocean-type Chinook salmon in the present study, to examine how seawater tolerance progressed with development on a monthly basis.

## 3. Results

### 3.1. Candidate smoltification genes using microarray comparisons

Limma analysis of the Sockeye salmon dataset identified 1,296 significant features that separated parr and smolt. By comparison, the Rainbow trout dataset had a published 400 feature parr–smolt signature. Combining the Sockeye salmon and Rainbow trout signatures using sIPCA to identify the top 100 features, there were 53 upregulated and 23 downregulated features that were also significant for the Sockeye salmon limma analysis (Table S1). Upregulated features were represented by six biological functions of metabolism (*n* = 21), immunity (8), oxygen transport (7), growth (7), structural integrity (5), ion regulation (4), and calcium uptake (1); the ion regulation and oxygen transport functions were predominantly at the top end of fold changes. Downregulated features were represented mainly by the function of immunity (17); other functions included ion regulation (1), growth (1), repressor of circadian rhythm (1), and unknown (3). Comparisons to three additional studies using the 16K platforms identified 7 upregulated and 14 downregulated significant features for parr to smolt for both species (Table S2). Upregulated features contained the functions of immunity (4), metabolism (2), and ion regulation (1). Downregulated features all contained the function of immunity.

### 3.2. Candidate smoltification genes using literature mining

Across the five published microarray studies, there were 15 upregulated and 6 downregulated genes for parr-to-smolt that were significant in at least two studies (Table S3). Including the studies that examined specific genes, several of these studies found that the ion regulation genes Na^+^/K^+^-ATPase α-1 were significant, specifically for parr-to-smolt there was an upregulation of isoform ‘b’ and downregulation of isoform ‘a’ (e.g. Nilsen et al., 2007; Piironen et al., 2013; Stefansson et al., 2007). There was also support for parr-to-smolt upregulation of other ion regulation genes, i.e. cystic fibrosis transmembrane conductance regulator I (e.g. Nilsen et al., 2007) and Na^+^/K^+^/2Clcotransporter (e.g. Nilsen et al., 2007; Stefansson et al., 2007). Although many plasma hormones change appreciably during smoltification (e.g. McCormick et al., 2013), the majority of the associated genes were unchanged for parrto-smolt in the five microarray studies. However, directed qPCR parr-to-smolt revealed significant upregulation of glucocorticoid (cortisol) receptor (Kiilerich et al., 2007, but see Hecht et al., 2014), growth hormone (Hecht et al., 2014) and receptor (Hecht et al., 2014; Kiilerich et al., 2007; Stefansson et al., 2007), insulin-like growth factor and receptor (Stefansson et al., 2007), and thyroid receptor beta (Hecht et al., 2014). Also, there may be a significant downregulation for parr-to-smolt of prolactin receptor (Kiilerich et al., 2007, but see Hecht et al., 2014). Hence, we included assays to these genes in our test panel.

### 3.3. qPCR assays of select candidate genes

A total of 45 candidate smoltification genes were selected for TaqMan qPCR design: 25 upregulated and 20 downregulated for parr to smolt (Table 2). The majority of the candidate genes (*n* = 34) were from the microarray analyses using both Sockeye salmon and Rainbow trout; 13 of these genes were also present in the literature review. Of the 34 genes, 28 were from the 44K analysis and mainly represented the extremes of the fold changes (Table S1), and six were from the 16K analysis and represented most of the available genes for this analysis (Table S2). Another two genes (S100A4 and FKBP5) were identified as highly differentially expressed by Sutherland et al. (2014) for Rainbow trout and were added by visual inspection of Sockeye salmon boxplots. The last nine genes were from the literature mining to fill eight biological functions, i.e. ion regulation, oxygen transport, metabolism, growth, calcium uptake, structural integrity, immunity, and hormones, so there would be at least two representative genes.

**Table 2:**
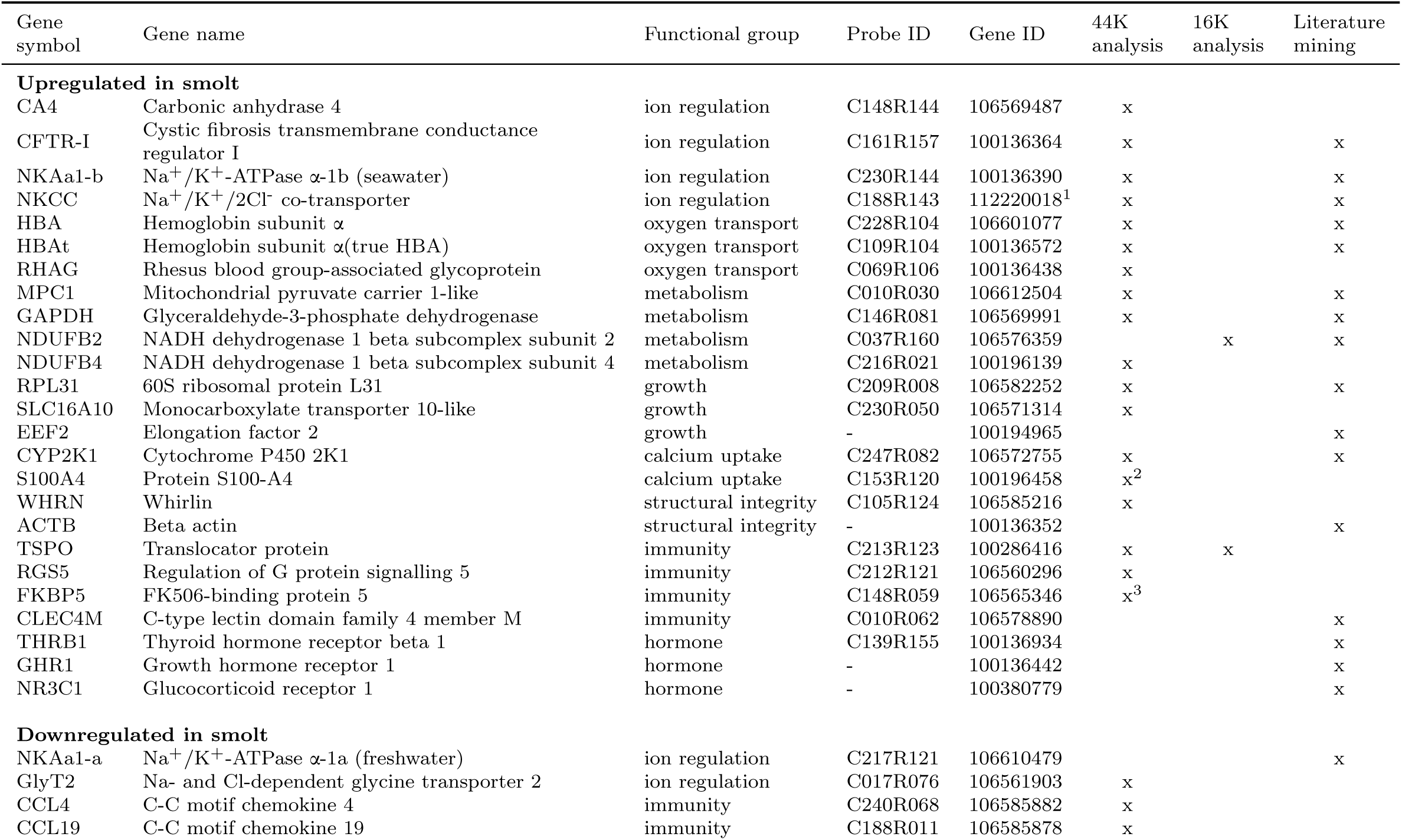
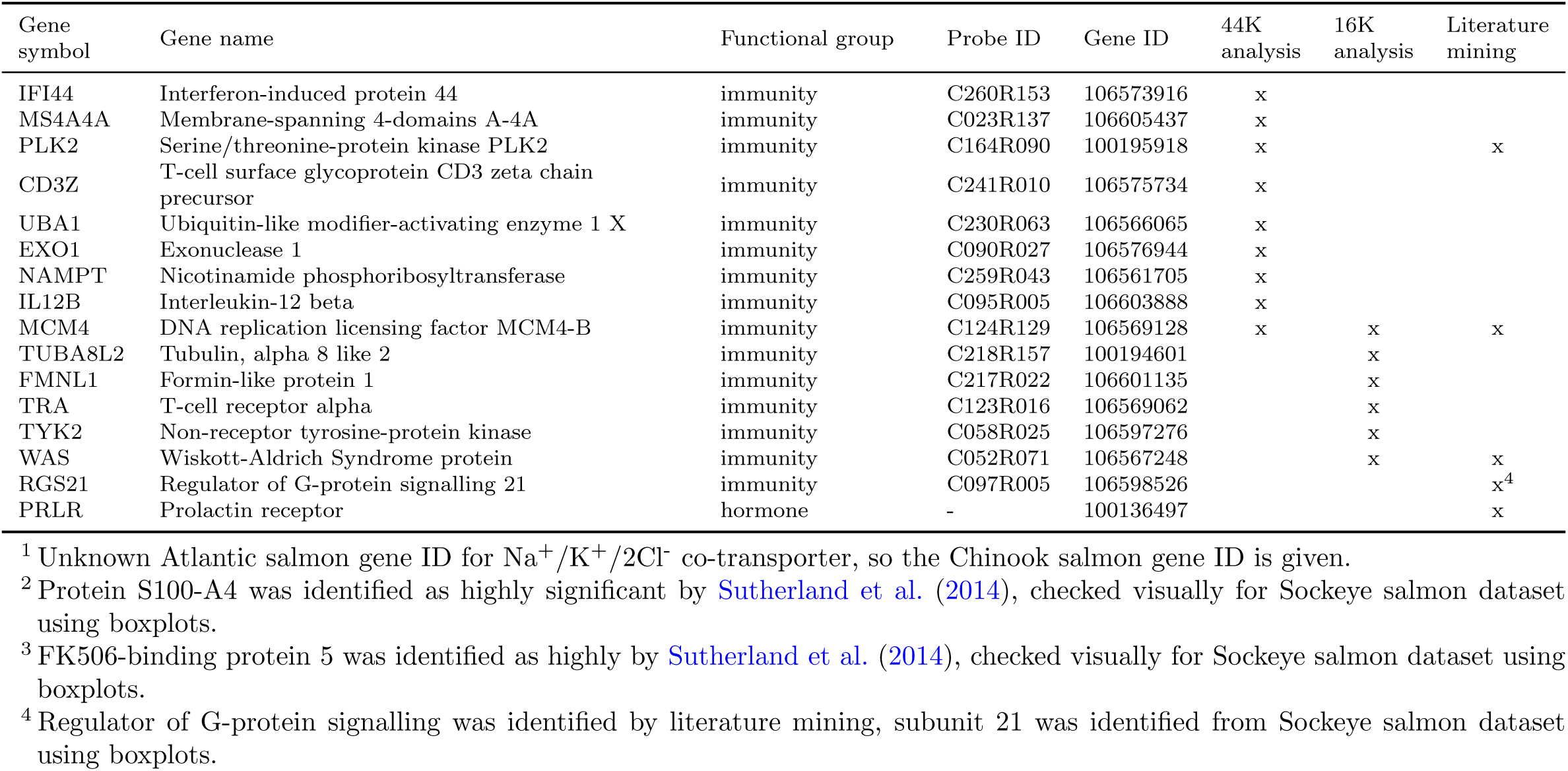
Summary of the candidate smoltification biomarkers for qPCR assay design using gill tissue. Presented for each gene is the smoltification functional group, feature (probe) ID for the 44K cGRASP microarray, and Atlantic salmon (*Salmo salar)* gene ID. The symbol x indicates that the gene was significant for parr to smolt for a specified analysis. Stars represent genes that were added after visualization of Rainbow trout *(Oncorhynchus mykiss*) and Sockeye salmon (*O. nerka*) boxplots.

For efficiency testing, two assays were designed for the top 12 upregulated and 10 downregulated genes (set 1), remaining genes had one assay design (set 2) (Table S4). Eight out of 45 assays did not pass the efficiency criteria (i.e. CD3Z, GAPDH, GlyT2, NKCC, RGS5, TYK2, S100A4, and WHRN) across species, thus leaving 20 upregulated gene assays and 17 downregulated genes (Table S5). For the genes with two assays (set 1), the assay with the highest efficiency across species was selected for further analysis with validation juvenile samples.

### 3.4. Validation of smoltification genes

Across all four groups, a PCA of gill expression of 37 candidate genes identified that PC2 separated earlier and later months (Figure S1). PC1 was associated with group differences. The expression of 32 genes was significantly correlated (*p* < 0.05) with PC2 (summary in Table 3; statistics in Table S6–S10; data in Appendix 1–4). The top 10 genes based on correlation significance were represented by five upregulated biomarkers and five downregulated biomarkers in smolts (Figure 1).

**Table 3:**
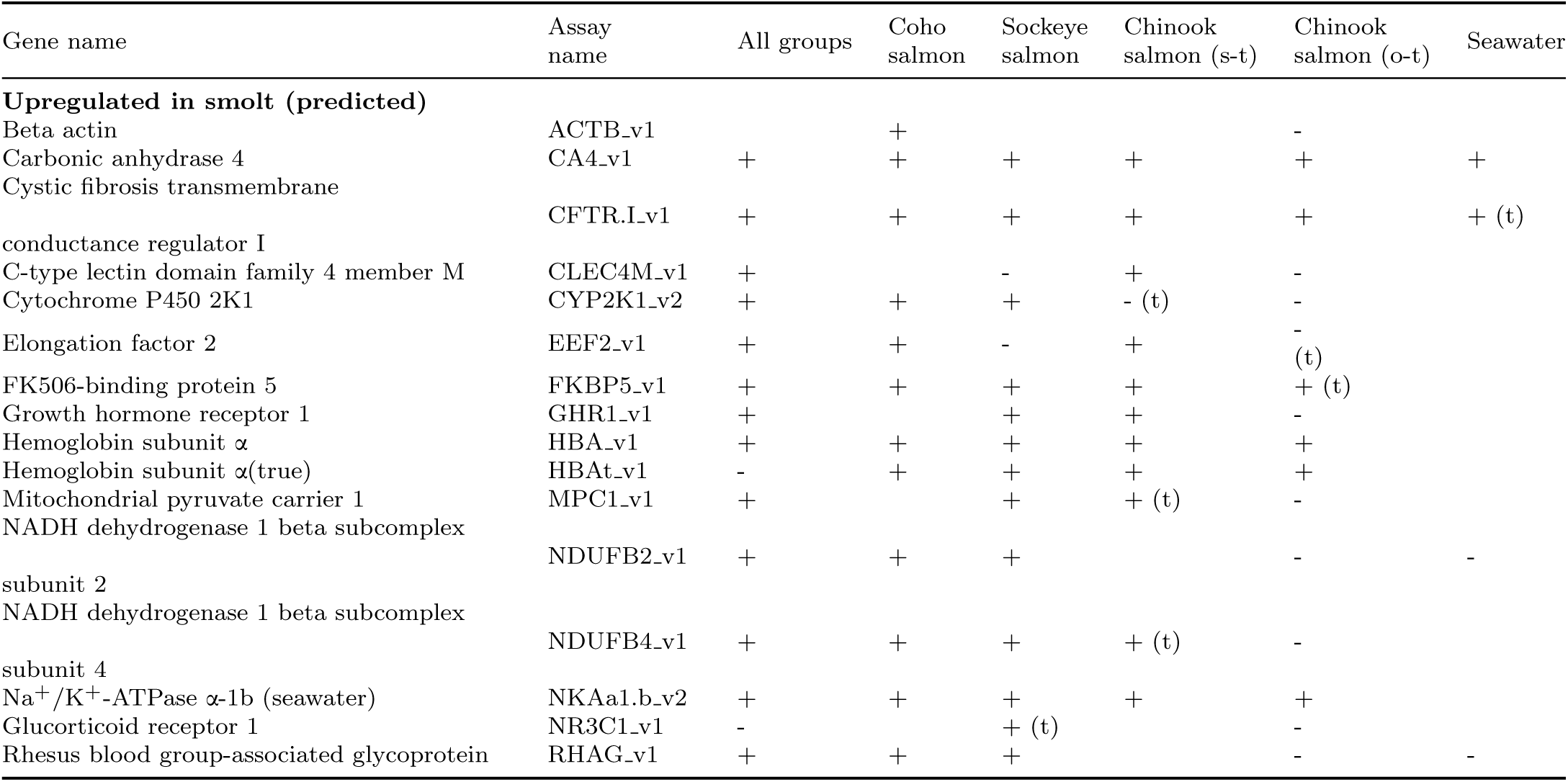
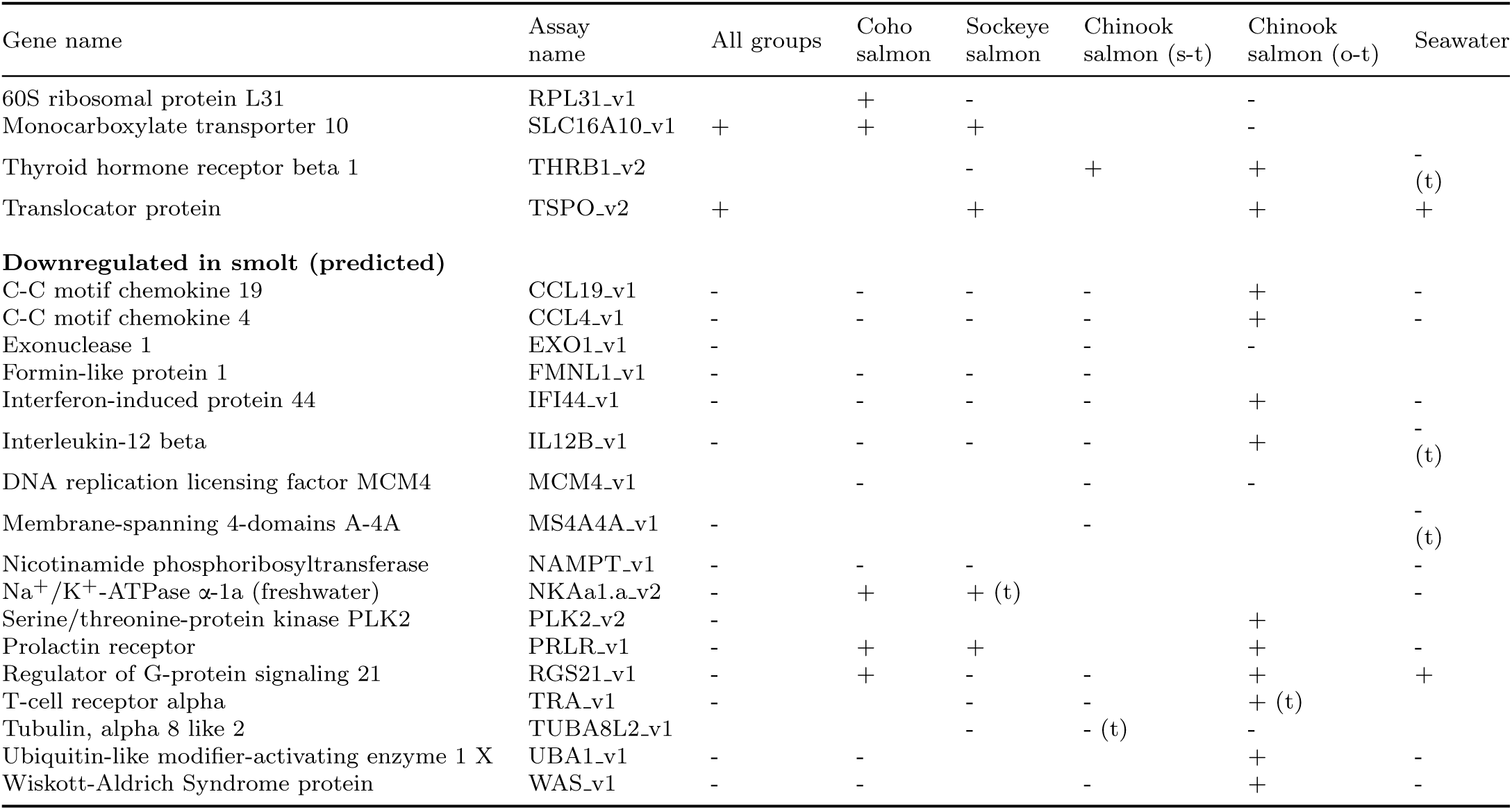
Summary of the gill smoltification gene expression patterns for the four groups. The expression values of the. 37 candidate genes were subjected to principal component analysis (PCA) for all four groups and each group separately: Coho salmon, Sockeye salmon, stream-type Chinook salmon, and ocean-type Chinook salmon. Gene expression relationships with the main PC axis separating earlier and later months was examined. Student t-tests examined expression differences between freshwater and seawater Nitinat ocean-type Chinook salmon sampled at the same time in late April; estuary juveniles were exposed to seawater for about two weeks. Presented are the significant (*p <* 0.05) expression patterns: + for positive correlation with smoltification or higher in seawater and for negative correlation with smoltification or lower in seawater. Trends (*p <* 0.1) are presented with a t in brackets. Pearson correlations and *p*-values are displayed in Table S6–S10; beanplots of the data are displayed in Appendix 1–4. Freshwater and seawater mean differences and statistics are displayed in Table S11; beanplots of the data are displayed in Appendix 4.

**Figure 1:**
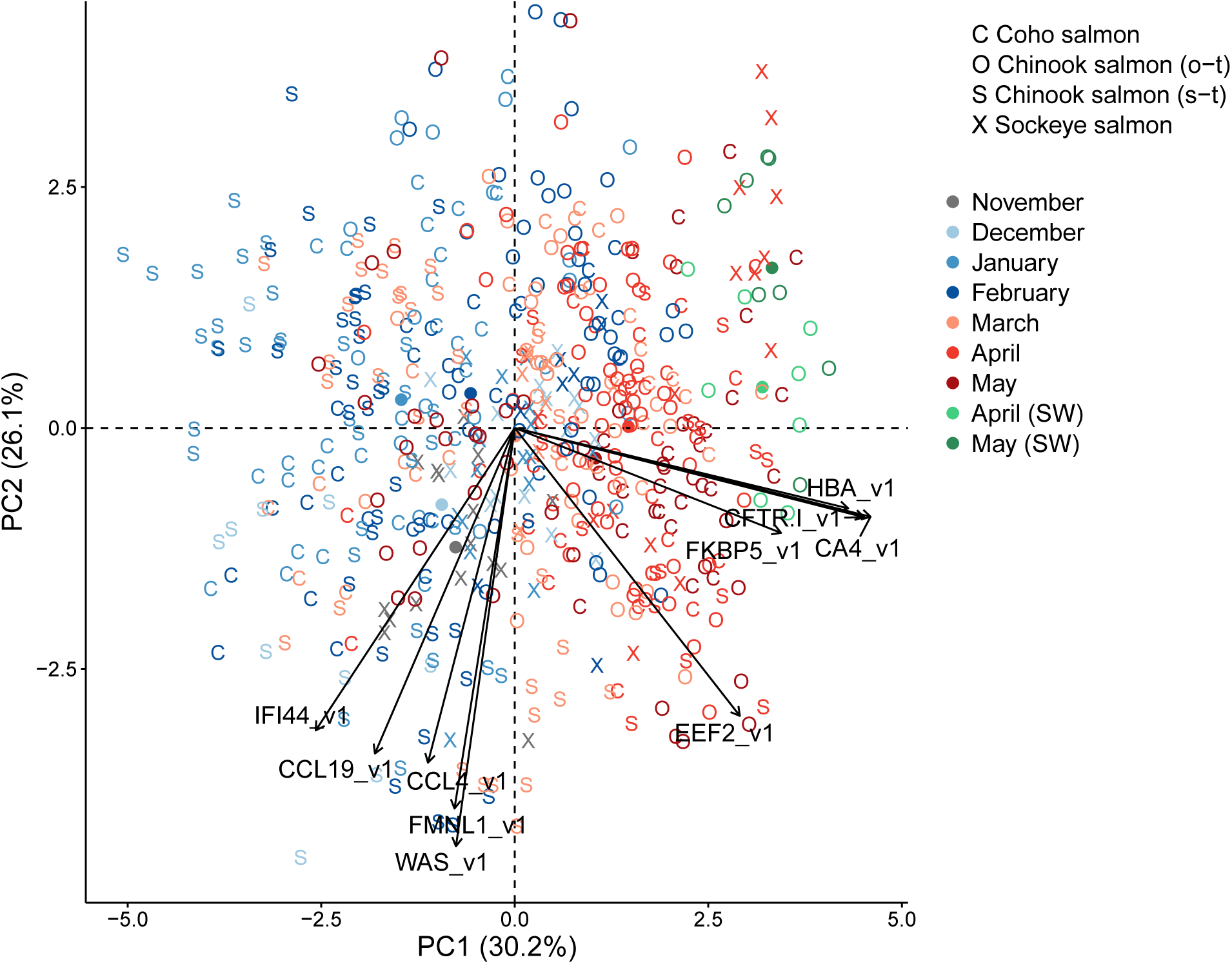
Canonical plots of the first two principal components of the top 10 biomarkers for smoltification using all four groups. Groups are Coho salmon (*Oncorhynchus kisutch*), Sockeye salmon (*O. nerka*), Chinook salmon (stream-type, *O. tshawytscha*), and Chinook salmon (ocean-type, *O. tshawytscha*). Percentage in brackets is the variation explained by the component. Monthly sample centroids are represented by the circle of the same colour. Black arrows represent loading vectors of the biomarkers. Legend symbol SW is for seawater and these individuals were not used in the PCA.

Within each of the four groups, PCAs of gill expression of 37 candidate genes identified that PC2 separated earlier and later months (Figure S2). PC1 was associated with different sets, i.e. hatchery or wild and source population. Coho salmon had 26 genes, Sockeye salmon had 28 genes, stream-type Chinook salmon 21 genes, and ocean-type Chinook salmon had 30 genes with expression values significantly correlated with PC2 (Table 3). Notably, ocean-type Chinook salmon had metabolic and growth genes downregulated, and eight immunity genes upregulated during smoltification, opposite the prediction. Five biomarkers, i.e. CA4, CFTR-I, HBA, HBAt, NKAa1b, were consistently upregulated across all groups (Figure 2). An additional four biomarkers, i.e. CCL19, CCL4, IFI44, IL12B, were consistently downregulated for Coho salmon, Sockeye salmon, and stream-type Chinook salmon, but upregulated for ocean-type Chinook salmon.

**Figure 2:**
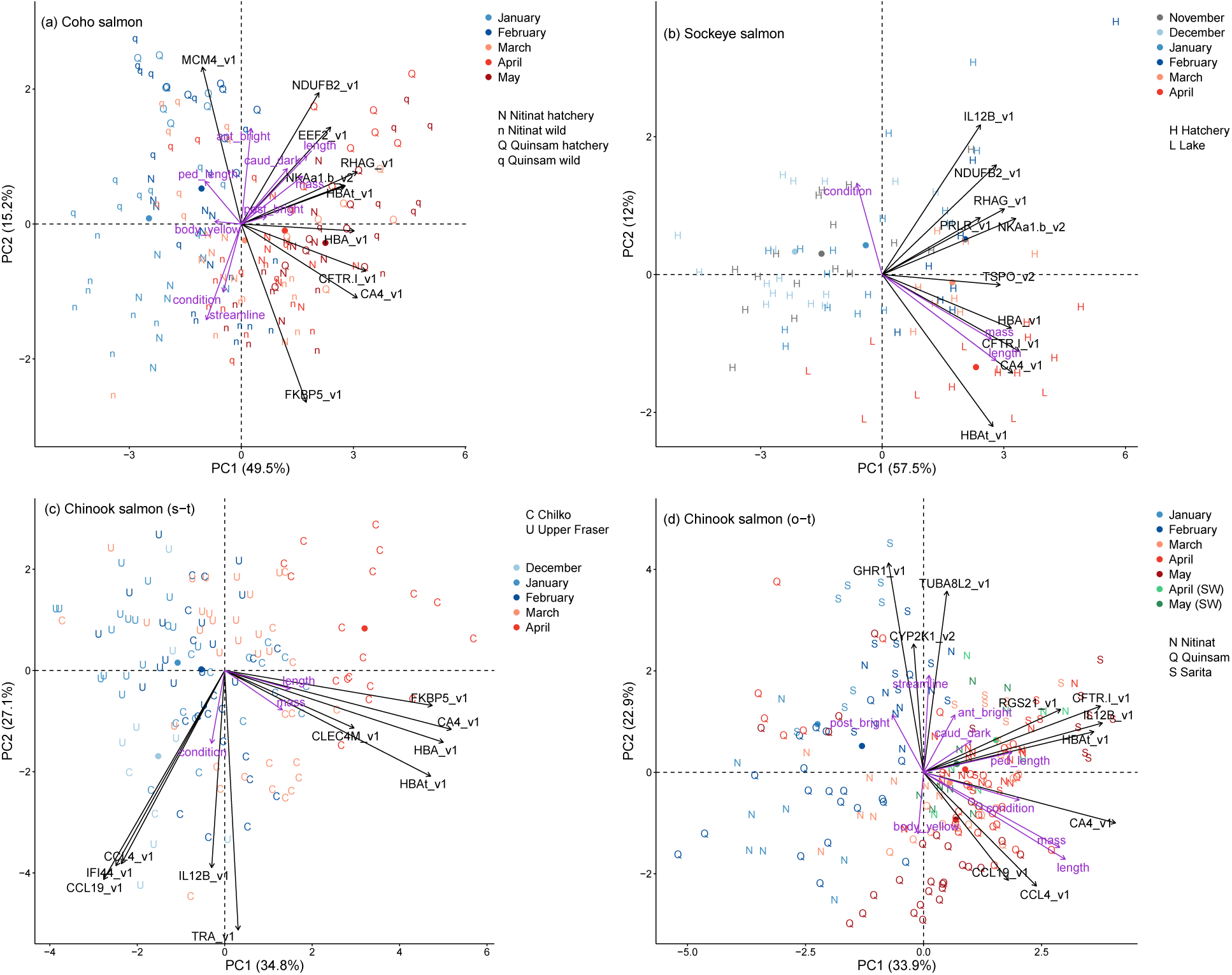
Canonical plots of the first two principal components of the top 10 biomarkers for smoltification using each of the four groups. (a) Coho salmon, (b) Sockeye salmon, (c) stream-type Chinook salmon, and (d) ocean-type Chinook salmon. Purple arrows represent loading vectors of the body variables. See Figure 1 legend.

Comparing Nitinat ocean-type Chinook salmon collected at the same time in late April from freshwater and seawater (about two weeks exposure to an estuary), 13 of the 30 genes were differently expressed between environments (Table 3; Table S11; Appendix 4). Interestingly, the genes predicted to be downregulated during smoltification were first upregulated in freshwater and only downregulated in seawater.

### 3.5. Relationship to gill NKA activity and body variables

Smoltification biomarker panels for each of the four groups, i.e. PC1 and PC2 using the top 10 genes (Figure 2), were significantly correlated with gill Na^+^/K^+^-ATPase activity (Figure 3). Body length and mass were positively correlated with PC1 for each of the four groups, as expected for juveniles growing during smoltification (Figure 2, statistics in Table S12). Body condition was also correlated with PC1 for ocean-type Chinook salmon, whereas it was correlated with PC2 for Coho salmon, Sockeye salmon, and stream-type Chinook salmon. Monthly body length, mass, and condition for all groups are presented in Figure S3.

**Figure 3:**
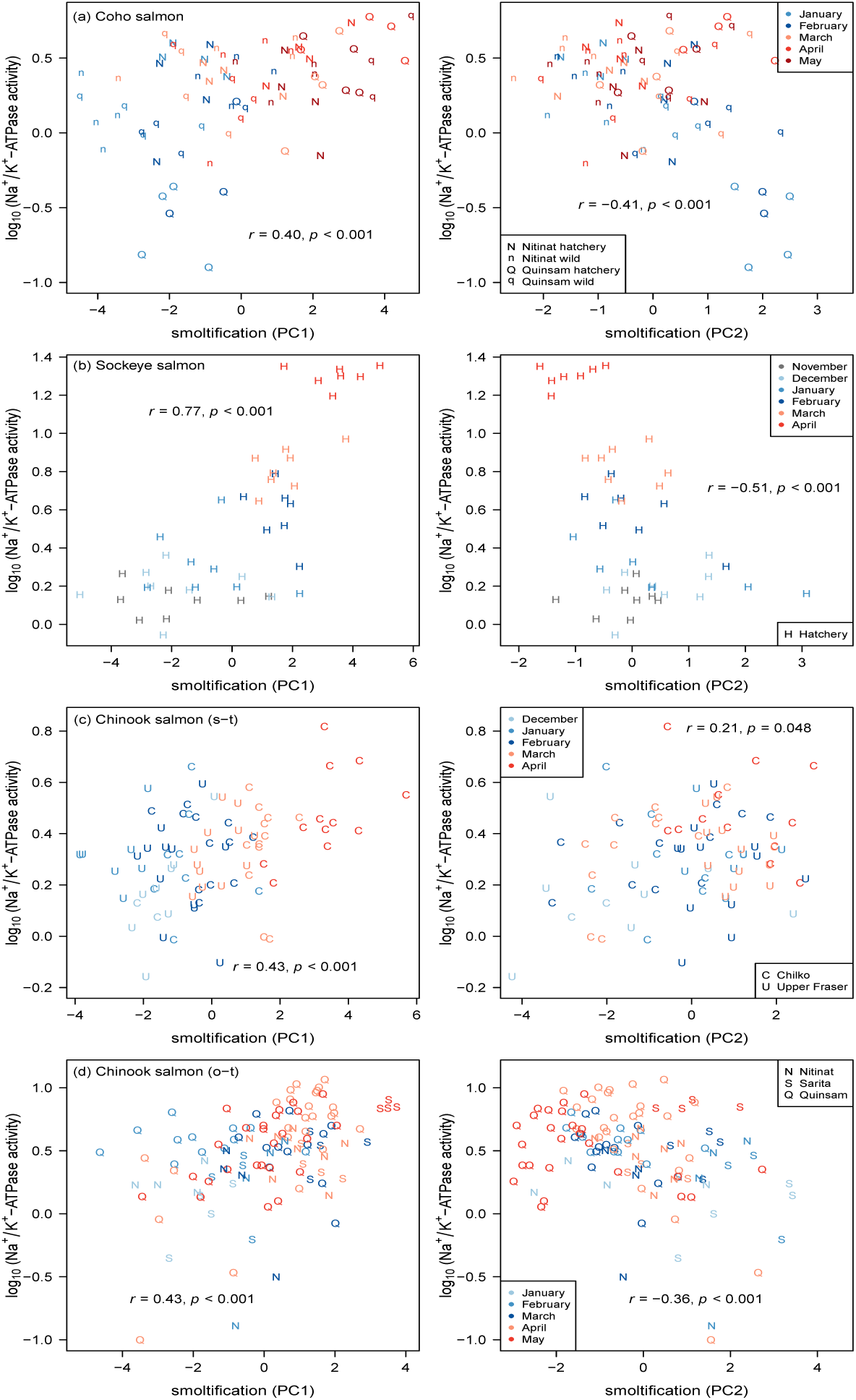
Relationships between smoltification gene expression patterns and Na^+^/K^+^ATPase activity for the four groups. By row: (a) Coho salmon, (b) Sockeye salmon, (c) stream-type Chinook salmon, and (d) ocean-type Chinook salmon. Gene expression patterns used the top 10 biomarkers. Gill Na^+^/K^+^-ATPase activity units are µmol ADP (mg protein)^−1^ h^−1^, which are presented as log_10_. There were no samples for Sockeye salmon from Cultus Lake in April. Legend symbol SW is for seawater.

Photographs to examine for correlations with skin pigmentation and body morphology were available only for the Coho salmon and ocean-type Chinook salmon. We considered the first four principal component axes (PCs) for skin pigmentation and two relative warps axes (RWs) for body morphology. For skin pigmentation, Coho salmon and Chinook salmon PC1 (53.8 and 37.3%) and PC2 (23.1 and 31.2%) were primarily associated with the posterior and anterior region brightness, respectively. Coho salmon PC3 (10.1%) was associated with body (posterior, anterior, and caudal fin) region yellowness and PC4 (5.7%) with caudal fin darkness; these traits were PC4 (6.5%) and PC3 (19.3%) for Chinook salmon, respectively. For body morphology, we considered the RWs for truncated to streamlined body shape, i.e. Coho salmon RW2 (12.6%) and Chinook salmon RW5 (ΔΔ%), and caudal peduncle length, i.e. RW7 (4.5 and 3.9%), because of their relationship with smoltification (Björnsson et al., 2011; McCormick et al., 1998; 2013).

The smoltification biomarker PC1s for both groups were positively correlated with caudal fin darkness (Figure 2). Coho salmon PC1 also had a positive trend for posterior brightness, as well as negative correlations with streamlined to truncated shape and caudal peduncle length and there was a trend for body yellowness. PC2 was correlated with anterior brightness. Chinook salmon PC1 was also positively correlated with caudal peduncle length. PC2 was correlated with posterior brightness, anterior brightness, body yellowness, and streamlined to truncated shape. Monthly skin pigment and body morphology are presented in Figure S4 and Figure S5, respectively.

### 3.6. Seawater tolerance classification model

The initial PCA of the gill expression using 37 candidate genes for ocean-type Chinook salmon in the present study indicated a pre-smolt to smolt pattern for PC2, and suggested a smolt to de-smolt pattern for PC3 (Figure 4a). Specifically, Quinsam May juveniles separated from earlier months along PC3. De-smoltification was also suspected for Quinsam May juveniles because of a decrease in gill Na^+^/K^+^-ATPase activity (mean ± SE, April 5.7 ± 0.7 and May 3.9 ± 0.4 µmol ADP (mg protein)^−1^ h^−1^, Student’s t-test *p* = 0.028). PC3 was significantly correlated with the expression of 25 genes (Table S10).

A new PCA using the top 20 biomarkers (*p <* 1×10^−5^ for both PC2 and PC3) maintained patterns as expected (Figure 4b), and the freshwater individuals of a companion study (Houde et al., 2018) were projected into this PCA. These freshwater individuals were assigned a smolt status at the trial-level based on the survival (over several days) of other individuals from the same trial during acute seawater transfer. By maximum of Youden’s J statistic and ROC analysis, the best PC2 threshold separating pre-smolt and smolt trials was 0.01, and the best PC3 threshold separating smolt and de-smolt trials was −1.40 (Figure 5). Individuals were classified as seawater tolerant (smolt) or intolerant (pre-smolt and de-smolt) using the areas defined by the thresholds.

**Figure 4:**
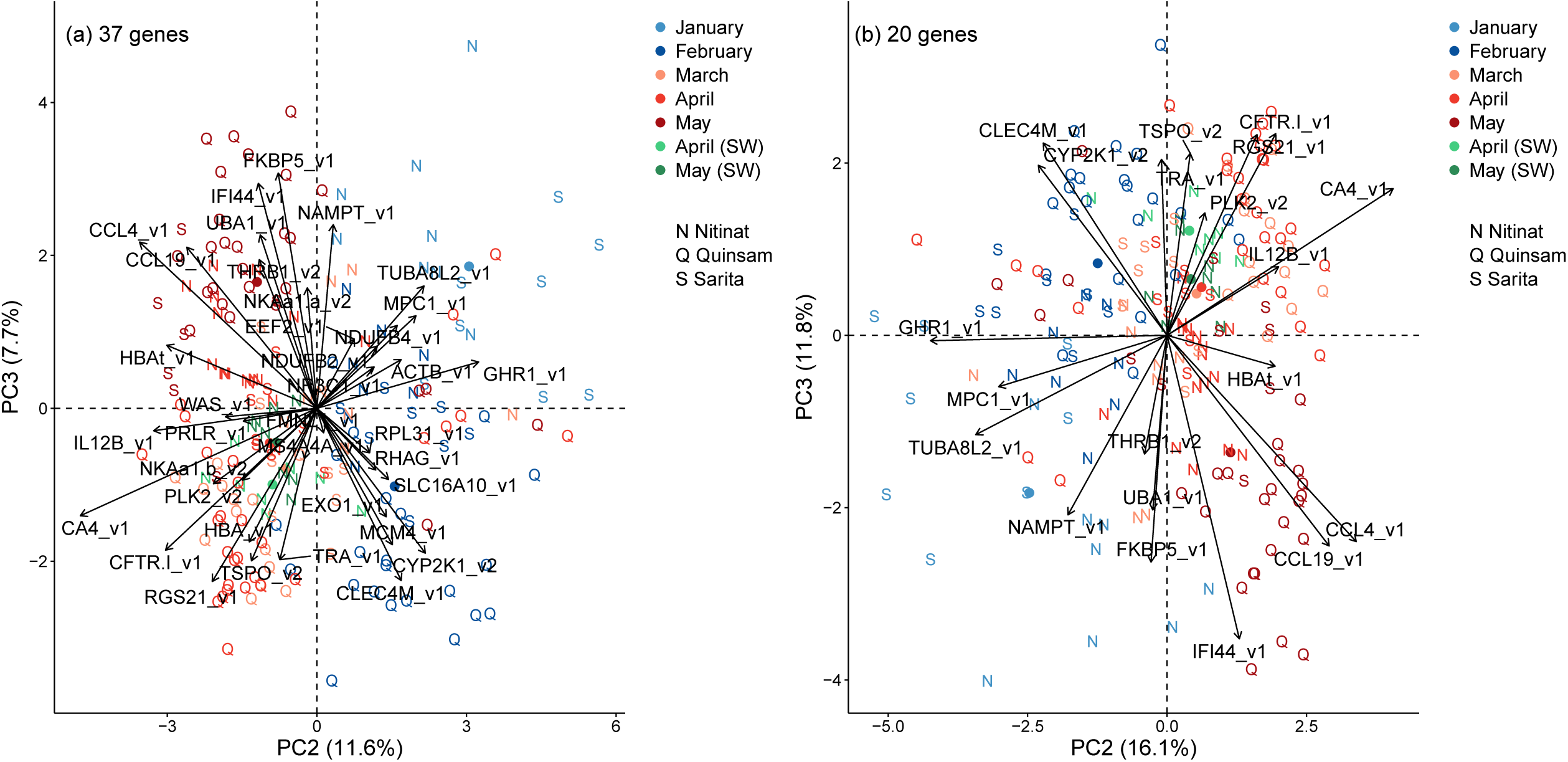
Canonical plots of the second and third principal components for the candidate genes using ocean-type Chinook salmon. Displayed are (a) all 37 genes and (b) the top 20 genes. See Figure 1 legend.

**Figure 5:**
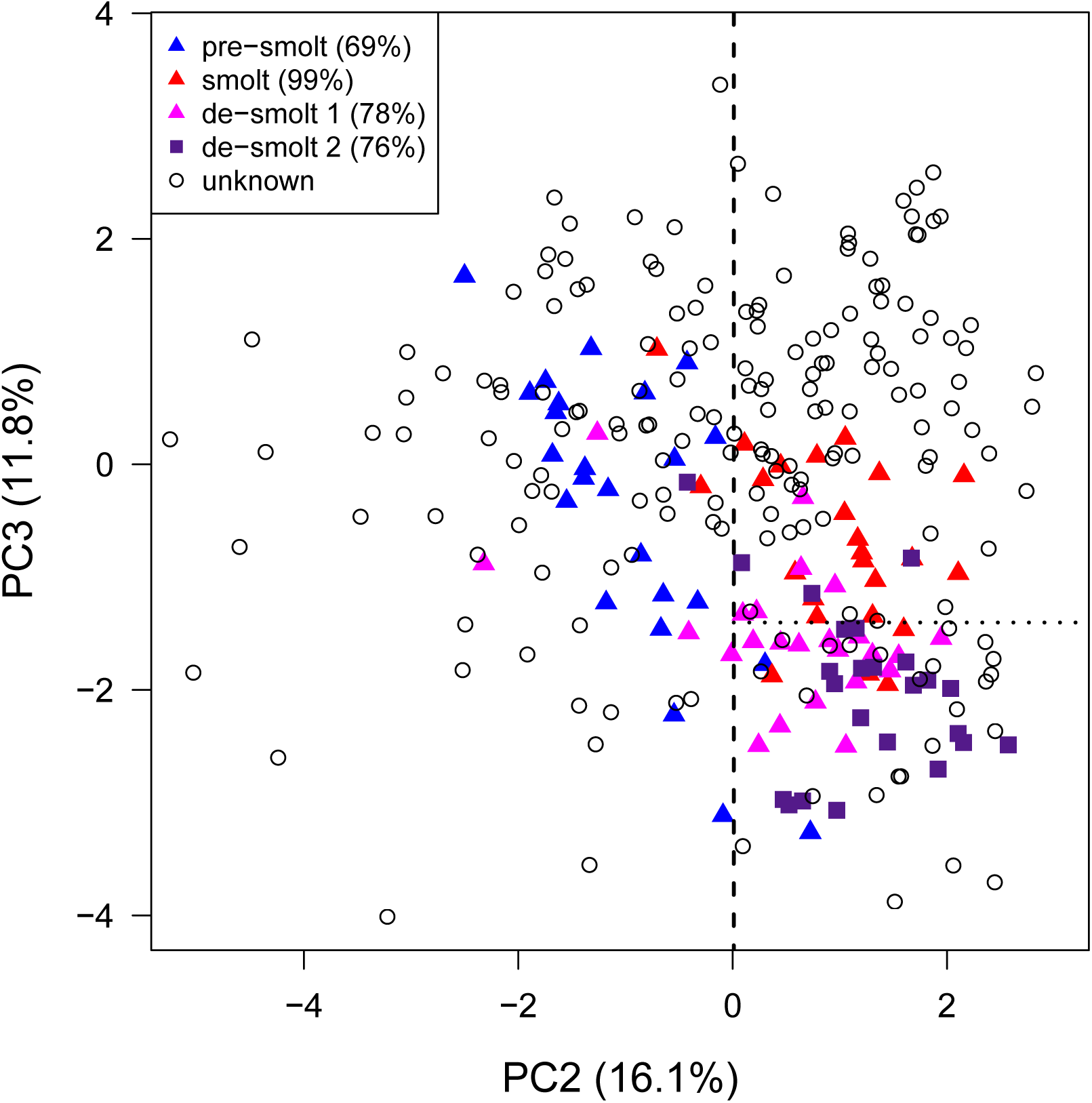
Seawater tolerance classification model using gene expression patterns of ocean-type Chinook salmon. Freshwater individuals with a smolt status are from the four trials of the companion study of Houde et al. (2018). Percentages for smolt statuses represent the trial seawater survival. The plot is based on the PCA using the top 20 biomarkers displayed in Figure 4b, and individuals of the companion study were projected into PC2 and PC3. Dashed lines represent the PC axis thresholds that separate (1) pre-smolt and smolt and (2) smolt and de-smolt. Thresholds were determined using Youden’s J statistic and ROC analysis. Juveniles within the ‘smolt’ area were classified as seawater tolerant and juveniles within the ‘pre-smolt’ and ‘de-smolt’ areas were classified as not seawater tolerant.

The classification model was applied to the unknown smolt status ocean-type Chinook salmon of the present study. Nitinat and Sarita juveniles were largely classed as seawater intolerant pre-smolt from January to March and seawater tolerant smolt in April and May (Table 4). On the other hand, Quinsam juveniles were classed as seawater intolerant pre-smolt in February, seawater tolerant smolt in March and April, and largely seawater intolerant de-smolt in May.

**Table 4:** Modelled seawater tolerance by monthly development for ocean-type Chinook salmon. The classification model used gene expression pattern thresholds for delineating seawater tolerant (smolt) and intolerant (pre-smolt and de-smolt). Month symbols are in chronological order of development and are the first three letters.

| Month | Seawater tolerance |  |  |
| --- | --- | --- | --- |
| Nitinat |  |  |  |
|  | pre-smolt | smolt | de-smolt |
| Jan | 6 | 0 | 2 |
| Feb | 8 | 0 | 0 |
| Mar | 7 | 1 | 0 |
| Apr | 1 | 14 | 1 |
| Sarita |  |  |  |
|  | pre-smolt | smolt | de-smolt |
| Jan | 8 | 0 | 0 |
| Feb | 8 | 0 | 0 |
| Mar | 4 | 4 | 0 |
| Apr | 2 | 6 | 0 |
| May | 2 | 5 | 1 |
| Quinsam |  |  |  |
|  | pre-smolt | smolt | de-smolt |
| Feb | 18 | 4 | 0 |
| Mar | 0 | 14 | 0 |
| Apr | 6 | 22 | 0 |
| May | 4 | 2 | 20 |

## 4. Discussion

Comparing gill gene expression for anadromous Rainbow trout (Sutherland et al., 2014) with our internal Sockeye salmon dataset, we discovered numerous common candidate smoltification genes. A subset of 25 upregulated and 20 downregulated genes, mainly representing the fold change extremes of the 44K analysis, were selected for TaqMan qPCR assay design. Of these 45, 20 upregulated and 17 downregulated genes passed our assay efficiency criteria and were then tested on monthly gill samples of Coho salmon, Sockeye salmon, stream-type Chinook salmon, and ocean-type Chinook salmon. We identified 32 common smoltification biomarkers; however, within each group variation of the smoltification biomarkers ranged from 21 to 30 genes. Nevertheless, in all cases, the smoltification biomarkers could be reduced to the top 10 genes and still retain good separation among the monthly samples along the smoltification axis. Indeed, smoltification gene expression patterns (i.e. PC1 and PC2 of the biomarker panels using the top 10 genes for each group) were confirmed by correlations with gill Na^+^/K^+^-ATPase activity. We recommend the smoltification biomarkers panels using the top 10 genes for the four groups (Figure 2). For species and ecotypes not examined in the present study, we recommend the smoltification biomarker panel using the top 10 genes for the groups combined (Figure 1).

### 4.1. Common gill smoltification genes among groups

Across the four groups, ion regulation (carbonic anhydrase 4, CA4 and cystic fibrosis transmembrane conductance regulator I, CFTR-I, and Na^+^/K^+^ATPase α-1b, NKAa1-b) and oxygen transport (hemoglobin alpha, HBAt and HBA) genes were upregulated during smoltification. Another oxygen transport gene (Rhesus blood group-associated glycoprotein, RHAG) was also upregulated for Coho salmon and Sockeye salmon. CFTR-I and NKA1a-b are important ion regulators for gill ionocytes that help remove excess chloride and sodium ions, respectively, from the body of fish in seawater (Evans et al., 2005).

Furthermore, four immunity genes (C-C motif chemokine 19, CCL19; C-C motif chemokine 4, CCL4; interferon-induced protein 44, IFI44; and interleukin-12 beta, IL12B) were downregulated during smoltification for Coho salmon, Sockeye salmon, and stream-type Chinook salmon, but upregulated for ocean-type Chinook salmon (elaborated below). Yet, these four genes had lower expression in seawater than freshwater for ocean-type Chinook salmon. The majority of immunity genes (300 out of 360), such as chemokines, can be downregulated during seawater acclimation, possibly because of a trade-off between the energetic costs of osmoregulation and pathogen resistance in seawater (Johansson et al., 2016). These eight genes were predominantly at the top end of upregulated and downregulated genes (based on fold change) in the 44K analysis, but were not detected in the 16K analysis. The upregulated genes and chemokines were also identified by literature mining. Four uncharacterized features showed downregulation in the 44K analysis, but limited sequence template precluded assay design. They may be worth pursuing should more sequence data become available.

The consistency of these ion regulator genes across groups suggests that the Na^+^/K^+^/2Cl^−^ cotransporter (NKCC), also within gill ionocytes, may also be a good species-wide smoltification biomarker (see Nilsen et al., 2007; Stefansson et al., 2007). Unfortunately, our single assay for NKCC only worked for Rainbow trout, possibly because at the time we had limited sequence information; thus, we were not able to examine this gene for our target salmonids. Relative to the other ion regulators, carbonic anhydrase has received lesser research attention. Yet recently, carbonic anhydrase genes were under rapid genetic selection for osmoregulation of Rainbow trout introduced from high to low salinities (Willoughby et al., 2018). Carbonic anhydrase can be important for both acid-base and ion regulation because of the productions of H^+^ and HCO3needed for Na^+^ and Clexchange in gill tissue (Gilmour, 2012; Havird et al., 2013). CA4 was the second most powerful single predictor of smoltification after CFTR-I using all groups.

Hemoglobin isoforms change from juvenile to adult types during smoltification of Coho salmon and Sockeye salmon (Vanstone et al., 1964). The adult type may have a higher oxygen affinity and weaker Bohr effect than the juvenile type, suggesting an adaptation to the lower oxygen tension of seawater than freshwater. Yet, Fyhn et al. (1991) found that the isoforms shifted after smoltification for stream- and ocean-type Chinook salmon, suggesting that they may be more body size dependent. However, our results, which show a contribution of hemoglobin genes to smoltification in stream-type and ocean-type Chinook salmon, suggest that hemoglobin may still have a smoltification role for Chinook salmon, although it may not be related to the isoform switching.

Our confidence in the smoltification biomarkers is strengthened by the gene expression similarities in response to higher salinity. In the companion study, we used these same candidate gene assays on juvenile ocean-type Chinook salmon exposed to freshwater (0 PSU), brackish (20 PSU), and seawater (28 or 29 PSU) for six days (Houde et al., 2018). Ion regulation genes (i.e. CA4, CFTR-I, and NKAa1-b) and an oxygen transport gene (i.e. HBA) that were upregulated during smoltification in the present study also had higher expression in brackish and seawater than freshwater in the companion study. Similarly, the four immunity genes (i.e. CCL19, CCL4, IFI44, and IL12B) that were downregulated during smoltification had lower expression in brackish and seawater than freshwater. Overall, the strongest gill smoltification biomarkers, consistent for all the Pacific salmonid groups examined, were likely in preparation for higher salinity and decreased dissolved oxygen in marine relative to freshwater environments.

### 4.2. Different gill smoltification genes among groups

Beyond ion regulation and oxygen transport, gene expression patterns for the remaining six upregulated biological functions were dependent on the group or did not fit the prediction based microarray or literature information. In particular, three metabolic genes (NADH dehydrogenase 1 beta subcomplex subunit 2 and 4, NDUFB2 and NDUFB4, and mitochondrial pyruvate carrier 1, MPC1) were generally upregulated for Coho salmon, Sockeye salmon, and stream-type Chinook salmon, but downregulated for ocean-type Chinook salmon. Expression of metabolic genes can be related to body growth (Salem et al., 2007), and importantly photoperiod in known to influence growth of stream-type Chinook salmon but not ocean-type Chinook salmon (Clarke et al., 1992; 1994). Conceivably, metabolic gene expression may differ as a result of photoperiod dependence, but a mechanistic link would need to be found.

Three growth genes (monocarboxylate transporter 10, SLC16A10; elongation factor 2, EEF2; and 60S ribosomal protein L31, RPL31) were also generally upregulated for Coho salmon or stream-type Chinook salmon, but downregulated for Sockeye salmon or ocean-type Chinook salmon even though these two groups also continued to grow. Thus, elongation factors and ribosomal genes may not be consistently upregulated during smoltification, e.g. downregulation of elongation factor 1B and upregulation of ribosomal proteins (Lemmetyinen et al., 2013), downregulation of ribosomal proteins (Seear et al., 2010), and mixture of up and downregulation of ribosomal proteins (Boulet et al., 2012; Robertson and McCormick, 2012).

The structural integrity gene (beta actin, ACTB) did not change with smoltification for Sockeye salmon and stream-type Chinook salmon. Hecht et al. (2014) also found no change with ACTB for Rainbow trout. The calcium uptake gene (cytochrome P450 2K1, CYP2K1) was upregulated for Coho salmon and Sockeye salmon, but downregulated for stream- and ocean-type Chinook salmon. Another calcium uptake gene, protein S100A4 (S100A4) had the largest parr-to-smolt difference in expression for the Rainbow trout microarray study (Sutherland et al., 2014); unfortunately, our assay for S100A4 did not work for Chinook salmon and Sockeye salmon, so this gene was not examined further. One gene each represented the structural integrity and calcium uptake biological functions. Future work should examine other structural integrity genes such as collagen, SPARC, or tropomyosin (e.g. Seear et al., 2010; Lemmetyinen et al., 2013) and develop an assay for S100A4 which works on a broader range of species to examine the consistency of regulation across species and ecotypes.

No across group support was found for any of the hormone genes and for just one of three immunity genes predicted to be upregulated during smoltification. The immunity gene FK506–binding protein 5 (FKPBP5) was upregulated for Coho salmon, Sockeye salmon, stream-type Chinook salmon, with a similar trend for ocean-type Chinook salmon. On the other hand, translocator protein (TSPO) was upregulated for Sockeye salmon and ocean-type Chinook salmon only, and c-type lectin domain family 4 member M (CLEC4M) was upregulated for stream-type Chinook salmon only. In contrast to the Sockeye salmon and ocean-type Chinook salmon examined in the present study, Atlantic salmon, Rainbow Trout, and Brook trout in previous studies (Seear et al., 2010; Boulet et al., 2012; Lemmetyinen et al., 2013; Sutherland et al., 2014) showed upregulation of c-type lectins 2 or 4M. Growth hormone receptor 1 (GHR1) was upregulated for Sockeye salmon and stream-type Chinook salmon but downregulated for ocean-type Chinook salmon. Glucocorticoid (cortisol) receptor 1 (NR3C1) was upregulated for Sockeye salmon (trend) but downregulated for stream-type Chinook salmon. Thyroid hormone receptor beta 1 (THRB1) was upregulated for both types of Chinook salmon only. Although plasma values of these hormones are well associated with smoltification across species (e.g. McCormick et al., 2013), our results can be added to other studies suggesting that the gene expression patterns of these hormones or their receptors are not necessarily in line with plasma patterns (e.g. Kiilerich et al., 2007; Stefansson et al., 2007; Hecht et al., 2014). Overall, the immunity and hormone gene expression patterns suggest that there are species and ecotype differences during smoltification, or that these genes are functioning outside of the smoltification process. Further studies should examine the reproducibility of these patterns across species and ecotypes.

Beyond the four immunity genes that were generally downregulated during smoltification described above, predicted downregulation of remaining gill genes depended on the group. In particular, ocean-type Chinook salmon generally displayed an upregulation of certain immunity genes that where then generally downregulated in seawater. Immunity genes appear to be downregulated during smoltification for certain species and ecotypes, e.g. Sockeye salmon, while other species and ecotypes may not have a downregulation of these genes until reaching higher salinity, e.g. ocean-type Chinook salmon. Furthermore, ion regulation Na^+^/K^+^-ATPase α-1a (NKAa1-a) and prolactin receptor (PRLR) were lower in seawater than freshwater for ocean-type Chi-nook salmon, similar to the results of Houde et al. (2018). Higher expression of both genes was associated with mortality in seawater (Houde et al., 2018), indicating that the expression should decrease for seawater acclimation (also see Flores and Shrimpton, 2012).

### 4.3. Relationship to gill NKA activity

Elevated gill Na^+^/K^+^-ATPase activity has been associated with seawater survival of Atlantic salmon (e.g. Stich et al., 2015; 2016) and ocean-type Chinook salmon (Houde et al., 2018), as well as with the risk of predation for Rainbow trout (Kennedy et al., 2007). Similar correlations existed between log10 Na^+^/K^+^-ATPase (NKA) activity and the primary smoltification gene expression pattern (PC1) for Coho salmon (0.40), stream-type Chinook (0.43) salmon, and ocean-type Chinook salmon (0.43). The correlation was even stronger for Sockeye salmon (0.77), perhaps because the 44K candidate gene discovery analysis used this species. Although only moderate correlations are common between gene expression and protein activity (Schwänhausser et al., 2011; Kanerva et al., 2014), possibly because of post-transcriptional and post-translational modifications (Maier et al., 2009), changes in gene expression may be one of the first indicators of a physiological change or response (Feder and Walser, 2005; Miller et al., 2017). Furthermore, high NKA activity prior to seawater entry may not be necessary if a juvenile can rapidly increase NKA activity once in seawater (Madsen and Naamansen, 1989; Bassett et al., 2018). In support of this idea, we observed that ocean-type Chinook salmon smolts achieved a higher median NKA activity in seawater than in either pre-smolts or de-smolts (i.e.10.2 vs. < 7.5 µmol ADP (mg protein)^−1^ h^−1^, Houde et al., 2018).

### 4.4. Relationship to body appearance

Gill gene expression patterns were associated with skin pigmentation and body morphology changes during smoltification. Lower body condition, more streamlined body shape, elongation of caudal peduncle, increased body silvering, and darkening of caudal fin margins are commonly used indices of smoltification (McCormick et al., 1998; 2013; Björnsson et al., 2011). Correspondingly, the primary smoltification gene expression pattern (PC1) for Coho salmon and ocean-type Chinook salmon was associated with caudal fin darkness, as well as a positive trend with body brightness for Coho salmon. Although the PC1 pattern for Chinook salmon was also positively associated with caudal peduncle length, it was negative for Coho salmon. As far as we are aware, this is the first study to find relationships between gene expression patterns and body appearance during smoltification. Conceivably, caudal fin darkness may be a proxy of smoltification across other species and ecotypes but we did not have photographs of stream-type Chinook salmon and Sockeye salmon to test this possibility. Further research should examine whether these patterns occur in additional species and ecotypes.

### 4.5. Seawater tolerance model

Our preliminary seawater tolerance classification model for ocean-type Chinook salmon incorporated the gene expression patterns of freshwater presmolt, smolt, and de-smolt trials from a companion study using acute seawater transfers (Houde et al., 2018). Similar to Di Cicco et al. (2018) for classifying viral disease states, we statistically identified the gene expression (PC2 and PC3) thresholds that best separated pre-smolt, smolt, and de-smolt trials to classify individuals as seawater tolerant (smolt) or intolerant (pre-smolt and de-smolt). Our preliminary model appears to detect the gain as well as the loss of seawater tolerance using smolt status. Nitinat and Sarita juveniles were seawater tolerant in April and/or May around the hatchery release times, while Quinsam juveniles that were seawater tolerant in March and April were seawater intolerant (de-smolt) around the release times in May. The de-smoltification of May Quinsam juveniles was also confirmed by lower Na^+^/K^+^-ATPase activity. Even so, our discovery process for the candidate genes focussed on smoltification, i.e. pre-smolt to smolt. Other genes (e.g. FKBP5, IFI44, NAMPT, and UBA1) and a longer sampling period into the summer may improve resolution between smolts and de-smolts.

Similar seawater tolerance classification models may be produced for Coho salmon, stream-type Chinook salmon, and Sockeye salmon. Our preliminary model for ocean-type Chinook salmon used the freshwater smolt status at the level of the trial, with other individuals acutely transferred to seawater for measures of survival (Houde et al., 2018). A more direct approach of linking freshwater gene expression to seawater survival at the level of the individual would have been more powerful. For example, a small gill biopsy a few days before seawater transfer followed by a survival measure covering a few days after transfer, e.g. six days (Houde et al., 2018). Additional data are needed between individual gene expression and subsequent seawater tolerance to improve the model.

### 4.6. Conclusion

Ion regulation, oxygen transport, and certain immunity genes were the best gill smoltification biomarkers, with consistent changes across multiple populations samples in the four groups examined. These genes were mainly selected from the 44K microarray discovery analysis and represented the top end of upregulated or downregulated genes based on fold changes. The directional shifts in expression were also similar between exposure to freshwater and brackish or seawater (Houde et al., 2018), implying an important role for higher salinity acclimation. Smoltification gene expression patterns had significant relationships with gill Na^+^/K^+^-ATPase activity, as well as caudal fin darkness for both Coho salmon and ocean-type Chinook salmon. Metabolic genes were upregulated and immunity genes were downregulated for photoperiod dependent species and ecotypes, i.e. stream-type Chinook salmon, Coho salmon, and Sockeye salmon, but the opposite occurred for photoperiod independent species and ecotypes, i.e. ocean-type Chinook salmon. We have provided a preliminary seawater tolerance classification model of presmolt, smolt, and de-smolt for ocean-type Chinook salmon. To expand this model for classification to other species and ecotypes, additional individual-level data linking freshwater gene expression and its association to seawater survival are required. Beyond the gill smoltification biomarkers, we are also developing biomarkers predictive of other divergent stressors, e.g. general stress and imminent mortality (Evans et al., 2011; Miller et al., 2011; Jeffries et al., 2012, 2014); viral disease development (Miller et al., 2017); and salinity, thermal, and hypoxia stress (Houde et al., 2018), to support the development of a ‘Salmon Fit-Chip’ tool to rapidly and inexpensively assess the physiological condition of 100s to 1000s of fish.

## Acknowledgements

This research was supported by the Natural Sciences and Engineering Research Council of Canada through a Postdoctoral Fellowship and by Mitacs/ Pacific Salmon Foundation through an Accelerate Internship to ASH. Funding was for the research was provided by Genome British Columbia, the Pacific Salmon Commission, and Fisheries and Oceans Canada Genomic Research and Development Fund. APF is supported by a Canada Research Chair. We are thankful for the smoltification advice of S. McCormick, M. Shrimpton, and S. Sharron. We thank the managers and staff at Nitinat Hatchery, Quinsam Hatchery, Chehalis Hatchery, and Inch Creek Hatchery, as well as A. Schulze, E. Di Cicco, C. Rycroft, O. Dyck, T. Smith, D. Callander, K Robinson, L. Pon, and J. Hills for help with sample collection and processing.

## Supplementary Tables

**Table S1:** Summary of the results from the 44K analysis using the combined Sockeye salmon and Rainbow trout datasets for gill tissue. The Rainbow trout dataset was from Sutherland et al. (2014). Sockeye salmon and Rainbow trout datasets were combined and analyzed collectively using a sparse independent principal component analysis to identify the top 100 features (i.e. Probe IDs) that separated parr and smolt. Presented are the 76 features that overlapped with the identified significant features of the separate Sockeye salmon robust limma analysis, which are ordered by fold change. Mean values are for parr and smolt of the Sockeye salmon dataset. Bold italics are gene names with Probe ID used for qPCR assay development; normal italics are gene names and Probe IDs matching to same gene ID.

| Probe ID | Gene symbol | Gene name | Functional group | Mean for parr | Mean for smolt | log<sub>2</sub> fold change | Fold change | Adjusted *p*-value |
| --- | --- | --- | --- | --- | --- | --- | --- | --- |
| Upregulated in smolt |  |  |  |  |  |  |  |  |
| ***C148R144*** | ***CA4*** | ***Carbonic anhydrase 4*** | ion regulation | -2.40 | 0.82 | 3.22 | 9.31 | 0.00 |
| ***C209R008*** | ***RPL31*** | ***60S ribosomal protein L31*** | growth | -1.04 | 1.08 | 2.12 | 4.34 | 0.01 |
| ***C228R104*** | ***HBA*** | ***Hemoglobin subunit alpha*** | oxygen transport | -1.66 | 0.32 | 1.98 | 3.96 | 0.00 |
| ***C105R124*** | ***WHRN*** | ***Whirlin*** | structural integrity | -0.84 | 0.97 | 1.81 | 3.51 | 0.00 |
| ***C069R106*** | ***RHAG*** | ***Rhesus blood group-associated glycoprotein*** | oxygen transport | -0.68 | 0.92 | 1.60 | 3.04 | 0.00 |
| ***C109R104*** | ***HBAt*** | ***Hemoglobin subunit alpha (true HBA)*** | oxygen transport | -1.25 | 0.30 | 1.55 | 2.93 | 0.03 |
| ***C011R079*** | ***RHAG*** | ***Rhesus blood group-associated glycoprotein*** | oxygen transport | -0.48 | 1.07 | 1.55 | 2.93 | 0.00 |
| ***C212R121*** | ***RGS5*** | ***Regulator of G-protein signalling 5*** | immunity | -0.93 | 0.48 | 1.41 | 2.65 | 0.03 |
| ***C161R157*** | ***CFTR-I*** | ***Cystic fibrosis transmembrane conductance regulator I*** | ion regulation | -1.65 | -0.39 | 1.26 | 2.40 | 0.01 |
| ***C230R144*** | ***NKAa1-b*** | ***Na<sup>+</sup>/K<sup>+</sup>-ATPase α-1b (seawater)*** | ion regulation | -1.22 | 0.04 | 1.26 | 2.39 | 0.01 |
| ***C213R123*** | ***TSPO*** | ***Translocator protein*** | immunity | -0.99 | 0.10 | 1.09 | 2.13 | 0.00 |
| C158R093 | RGS5 | Regulator of G-protein signaling 5 (gene ID 106584144) | immunity | -0.22 | 0.86 | 1.08 | 2.12 | 0.00 |
| ***C230R050*** | ***SLC16A10*** | ***Monocarboxylate transporter 10-like*** | growth | -0.17 | 0.91 | 1.08 | 2.11 | 0.00 |
| C146R037 | CDA | Cytidine deaminase | metabolism | -1.12 | -0.11 | 1.01 | 2.01 | 0.00 |
| ***C010R030*** | ***MPC1*** | ***Mitochondrial pyruvate carrier 1-like*** | metabolism | -0.44 | 0.55 | 0.98 | 1.98 | 0.04 |
| ***C247R082*** | ***CYP2K1*** | ***Cytochrome P450 2K1*** | calcium uptake | -0.40 | 0.51 | 0.91 | 1.88 | 0.04 |
| ***C188R143*** | ***NKCC*** | ***Na<sup>+</sup>/K<sup>+</sup>/2Cl<sup>-</sup> co-transporter*** | ion regulation | -0.85 | 0.01 | 0.86 | 1.82 | 0.04 |
| C111R149 | SEC61G | Protein transport protein Sec61 subunit gamma | growth | -0.08 | 0.77 | 0.85 | 1.80 | 0.01 |
| C109R066 | TIMM17A | Mitochondrial import inner membrane translocase subunit Tim17-A | metabolism | -0.48 | 0.35 | 0.84 | 1.78 | 0.01 |
| ***C146R081*** | ***GAPDH*** | ***Glyceraldehyde-3-phosphate dehydrogenase*** | metabolism | -1.07 | -0.25 | 0.82 | 1.77 | 0.03 |

| Probe ID | Gene symbol | Gene name | Functional group | Mean<br>for parr | Mean<br>for smolt | log<sub>2</sub> fold<br>change | Fold<br>change | Adjusted<br>*p*-value |
| --- | --- | --- | --- | --- | --- | --- | --- | --- |
| C183R158 | BCKDK | 3-methyl-2-oxobutanoate dehydrogenase<br>[lipoamide] kinase | metabolism | -0.61 | 0.19 | 0.80 | 1.74 | 0.01 |
| C005R080 | GLRX5 | Glutaredoxin-related protein 5 | oxygen transport | -0.52 | 0.24 | 0.77 | 1.70 | 0.02 |
| C069R043 | HIGD1A | HIG1 domain family member 1A | metabolism | 0.05 | 0.81 | 0.76 | 1.69 | 0.02 |
| C165R074 | RNF222 | RING finger protein 222 | structural integrity | -0.62 | 0.13 | 0.75 | 1.68 | 0.03 |
| C202R155 | At5g50100 | Uncharacterized protein At5g50100 | growth | -0.16 | 0.58 | 0.74 | 1.67 | 0.00 |
| C085R077 | CTNS | Cystinosin | immunity | -0.09 | 0.64 | 0.72 | 1.65 | 0.02 |
| C196R101 | PSPH | Phosphoserine phosphatase | metabolism | -0.46 | 0.24 | 0.70 | 1.63 | 0.04 |
| C221R144 | COX7A2 | Cytochrome c oxidase polypeptide VIIa-liver/heart | metabolism | -0.53 | 0.17 | 0.70 | 1.62 | 0.03 |
| C213R046 | PSPH | Phosphoserine phosphatase | metabolism | -0.42 | 0.27 | 0.69 | 1.62 | 0.01 |
| *C030R160* | *TSPO* | *Translocator protein* | immunity | -0.71 | -0.02 | 0.69 | 1.61 | 0.02 |
| C010R060 | MPC2 | Mitochondrial pyruvate carrier 2 | metabolism | -0.27 | 0.42 | 0.68 | 1.61 | 0.05 |
| C117R042 | MPC2 | Mitochondrial pyruvate carrier 2 | metabolism | -0.42 | 0.26 | 0.68 | 1.60 | 0.02 |
| C156R115 | MPC2 | Mitochondrial pyruvate carrier 2 | metabolism | -0.40 | 0.27 | 0.67 | 1.59 | 0.02 |
| C057R083 | PPT2 | lysosomal thioesterase PPT2-A-like | metabolism | 0.02 | 0.68 | 0.66 | 1.58 | 0.00 |
| C111R151 | HMGB3 | High mobility group protein B3 | immunity | 0.30 | 0.93 | 0.63 | 1.55 | 0.02 |
| C010R151 | CYCS | Cytochrome c, somatic | metabolism | -0.48 | 0.15 | 0.63 | 1.55 | 0.03 |
| C208R060 | CHCHD7 | Coiled-coil-helix-coiled-coil-helix<br>domain-containing protein 7 | metabolism | 0.06 | 0.69 | 0.63 | 1.54 | 0.04 |
| C142R087 | ALAD | Delta-aminolevulinic acid dehydratase | oxygen transport | -0.60 | 0.02 | 0.62 | 1.54 | 0.03 |
| C066R146 | MRPS17 | 28S ribosomal protein S17 | growth | -0.31 | 0.31 | 0.62 | 1.54 | 0.02 |
| C167R048 | BCKDK | 3-methyl-2-oxobutanoate dehydrogenase [lipoamide]<br>kinase | metabolism | 0.08 | 0.70 | 0.62 | 1.54 | 0.00 |
| C204R009 | UBALD1 | UBA-like domaing containing 1 | immunity | -0.46 | 0.16 | 0.62 | 1.54 | 0.00 |
| C175R095 | CCDC56 | Coiled-coil domain-containing protein 56 | metabolism | 0.04 | 0.64 | 0.61 | 1.52 | 0.02 |
| C172R080 | TOMM6 | Translocase of outer mitochondrial membrane 6 | immunity | -0.28 | 0.32 | 0.60 | 1.52 | 0.02 |
| C224R050 | PSMD9 | 26S proteasome non-ATPase regulatory subunit 9 | growth | -0.21 | 0.38 | 0.60 | 1.51 | 0.03 |
| C251R097 | TMEM254 | Transmembrane protein 254 | structural integrity | -0.14 | 0.45 | 0.59 | 1.51 | 0.04 |
| ***C216R021*** | ***NDUFB4*** | ***NADH dehydrogenase 1 beta subunit 4*** | metabolism | -0.22 | 0.37 | 0.59 | 1.51 | 0.03 |
| C211R097 | COX17 | Cytochrome c oxidase copper chaperone | metabolism | -0.22 | 0.36 | 0.58 | 1.50 | 0.03 |
| C238R060 | MRPL37 | 39S ribosomal protein L37 | growth | -0.18 | 0.35 | 0.53 | 1.44 | 0.04 |
| C045R135 | COX7A2L | Cytochrome c oxidase subunit VIIa-related protein | metabolism | -0.01 | 0.51 | 0.52 | 1.44 | 0.04 |
| C101R159 | TMED2 | Transmembrane emp24 domain-containing protein 2<br>precursor | structural integrity | -0.10 | 0.41 | 0.51 | 1.42 | 0.02 |
| C096R011 | ALAD | Delta-aminolevulinic acid dehydratase | oxygen transport | -0.55 | -0.07 | 0.48 | 1.40 | 0.01 |
| C245R002 | UQCRC2 | Cytochrome b-c1 complex subunit 2 | metabolism | 0.03 | 0.47 | 0.44 | 1.36 | 0.05 |
| C184R070 | ARPC5L | Actin-related protein 2/3 complex subunit 5-like<br>protein | structural integrity | -0.03 | 0.39 | 0.41 | 1.33 | 0.02 |
| Downregulated in smolt |  |  |  |  |  |  |  |  |
| *C240R068* | *CCL4* | *C-C motif chemokine 4* | immunity | 1.19 | -0.55 | -1.74 | -3.33 | 0.00 |
| *C017R076* | *GlyT2* | *NA- and Cl-dependent glycine transporter 2* | ion regulation | 0.84 | -0.80 | -1.64 | -3.13 | 0.02 |
| *C188R011* | *CCL19* | *C-C motif chemokine 19* | immunity | 0.92 | -0.59 | -1.51 | -2.86 | 0.00 |
| *C260R153* | *IFI44* | *Interferon-induced protein 44* | immunity | 0.40 | -0.78 | -1.18 | -2.27 | 0.00 |
| C124R109 | At5g5010 | Uncharacterized protein At5g50100 | growth | 0.41 | -0.67 | -1.09 | -2.12 | 0.02 |
| *C023R137* | *MS4A4A* | *Membrane-spanning 4-domains A-4A* | immunity | 0.40 | -0.58 | -0.99 | -1.98 | 0.02 |
| *C164R090* | *PLK2* | *Serine/threonine-protein kinase PLK2* | immunity | 0.31 | -0.64 | -0.95 | -1.93 | 0.01 |
| *C241R010* | *CD3Z* | *T-cell surface glycoprotein CD3 zeta chain precursor* | immunity | 0.19 | -0.74 | -0.93 | -1.90 | 0.01 |
| *C230R063* | *UBA1* | Ubiquitin-like modifier-activating enzyme 1 X | immunity | 0.49 | -0.43 | -0.92 | -1.89 | 0.04 |
| C244R027 |  | UNKNOWN |  | 0.29 | -0.62 | -0.91 | -1.88 | 0.02 |
| *C090R027* | *EXO1* | *Exonuclease 1* | immunity | 0.11 | -0.78 | -0.89 | -1.85 | 0.00 |
| C245R123 |  | UNKNOWN |  | 0.25 | -0.61 | -0.86 | -1.81 | 0.02 |
| *C259R043* | *NAMPT* | *Nicotinamide phosphoribosyltransferase* | immunity | 0.14 | -0.62 | -0.76 | -1.70 | 0.04 |
| *C095R005* | *IL12B* | *Interleukin-12 beta* | immunity | 0.31 | -0.43 | -0.74 | -1.67 | 0.00 |
| C223R096 | KDM5B | Lysine demethylase 5B | immunity | 0.34 | -0.36 | -0.70 | -1.62 | 0.04 |
| C164R017 | BHLHE40 | Basic helix-loop-helix family member E40 | repressor of circadian rhythm | -0.26 | -0.91 | -0.65 | -1.57 | 0.03 |
| C081R077 | NFX1 | Nuclear transcription factor, X-box binding 1 | immunity | 0.19 | -0.44 | -0.62 | -1.54 | 0.03 |
| C146R113 | DTX3 | Deltex E3 ubiquitin ligase 3 | immunity | 0.15 | -0.46 | -0.61 | -1.52 | 0.02 |
| C229R071 | ASAP2 | ArfGAP with SH3 domain, ankyrin repeat and PH domain 2 | immunity | 0.25 | -0.36 | -0.61 | -1.52 | 0.01 |
| C124R132 | RIPK4 | Receptor interacting serine-threonine kinase 4 | immunity | 0.31 | -0.27 | -0.58 | -1.50 | 0.01 |
| C057R087 |  | Transposable element |  | 0.05 | -0.50 | -0.54 | -1.46 | 0.03 |
| *C124R129* | *MCM4* | *DNA replication licensing factor MCM4-B* | immunity | -0.06 | -0.55 | -0.49 | -1.41 | 0.04 |
| C177R004 | BCL10 | B-cell lymphoma/leukemia 10 | immunity | 0.08 | -0.40 | -0.48 | -1.40 | 0.04 |

**Table S2:** Summary of the results from the 16K signature analysis using the Sockeye salmon and Rainbow trout datasets for gill tissue. The Rainbow trout dataset was from Sutherland et al. (2014). Presented are the mapped 44K features that were significant for both Sockeye salmon and Rainbow trout, which are ordered by fold change. Bold italics are gene names with Probe ID used for qPCR assay development; normal italics are gene names and Probe IDs matching to same gene ID.

| Probe ID | Gene symbol | Gene name | Functional group | 16K study | Sockeye salmon |  | Rainbow trout |  |
| --- | --- | --- | --- | --- | --- | --- | --- | --- |
|  |  |  |  |  | Log<sub>2</sub> fold change | p-value | Log<sub>2</sub> fold change | p-value |
| Upregulated in smolt |  |  |  |  |  |  |  |  |
| **C213R123** | **TSPO** | **Translocator protein** | immunity | Robertson and McCormick (2012) | 1.04 | <0.01 | 0.50 | <0.01 |
| C071R152 | TSPO | Translocator protein | immunity | Robertson and McCormick (2012) | 0.70 | <0.01 | 0.44 | <0.01 |
| C030R160 | TSPO | Translocator protein | immunity | Robertson and McCormick (2012) | 0.64 | <0.01 | 0.54 | <0.01 |
| C004R095 | TSPO | Translocator protein | immunity | Robertson and McCormick (2012) | 0.61 | <0.01 | 0.45 | <0.01 |
| C183R077 | SGK3 | Serine/threonine kinase 3 | ion regulation | Robertson and McCormick (2012) | 0.48 | <0.01 | 0.66 | 0.04 |
| C057R039 | CYCS | Cytochrome C, somatic | metabolism | Boulet et al. (2012) | 0.48 | 0.04 | 0.98 | <0.01 |
| **C037R160** | **NDUFB2** | **NADH dehydrogenase 1 beta subunit 2** | metabolism | Robertson and McCormick (2012) | 0.29 | 0.02 | 0.26 | 0.02 |
| Downregulated in smolt |  |  |  |  |  |  |  |  |
| **C218R157** | **TUBA8L2** | **Tubulin, alpha 8 like 2** | immunity | Lemmetyinen et al. (2013) | -0.73 | <0.01 | -0.28 | 0.04 |
| **C217R022** | **FMNL1** | **Formin-like protein 1** | immunity | Lemmetyinen et al. (2013) | -0.62 | <0.01 | -0.30 | 0.01 |
| C118R080 | PBRM1 | Polybromo-1 | immunity | Robertson and McCormick (2012) | -0.60 | 0.02 | -0.29 | 0.01 |
| C217R123 | TRA | T-cell receptor alpha | immunity | Robertson and McCormick (2012) | -0.58 | 0.05 | -0.71 | 0.01 |
| **C123R016** | **TRA** | **T-cell receptor alpha** | immunity | Robertson and McCormick (2012) | -0.56 | 0.03 | -0.81 | <0.01 |
| C189R144 | TRA | T-cell receptor alpha | immunity | Robertson and McCormick (2012) | -0.54 | 0.03 | -0.74 | 0.01 |
| C185R169 | TRA | T-cell receptor alpha | immunity | Robertson and McCormick (2012) | -0.52 | <0.01 | -0.63 | 0.01 |
| C062R132 | FMNL1 | Formin-like protein 1 | immunity | Lemmetyinen et al. (2013) | -0.51 | 0.03 | -0.39 | 0.03 |
| C135R158 | FMNL1 | Formin-like protein 1 | immunity | Lemmetyinen et al. (2013) | -0.50 | 0.04 | -0.35 | 0.01 |
| **C124R129** | **MCM4** | **DNA replication licensing factor MCM4-B** | immunity | Robertson and McCormick (2012) | -0.49 | <0.01 | -0.57 | <0.01 |
| **C058R025** | **TYK2** | **Non-receptor tyrosine-protein kinase TYK2** | immunity | Robertson and McCormick (2012) | -0.46 | 0.01 | -0.34 | <0.01 |
| **C052R071** | **WAS** | **Wiskott-Aldrich syndrome protein** | immunity | Robertson and McCormick (2012) | -0.42 | 0.01 | -0.77 | <0.01 |
| C236R165 | TYK2 | Non-receptor tyrosine-protein kinase TYK2 | immunity | Robertson and McCormick (2012) | -0.36 | 0.03 | -0.42 | <0.01 |
| C131R157 | TYK2 | Non-receptor tyrosine-protein kinase TYK2 | immunity | Robertson and McCormick (2012) | -0.36 | 0.02 | -0.40 | <0.01 |

**Table S3:**
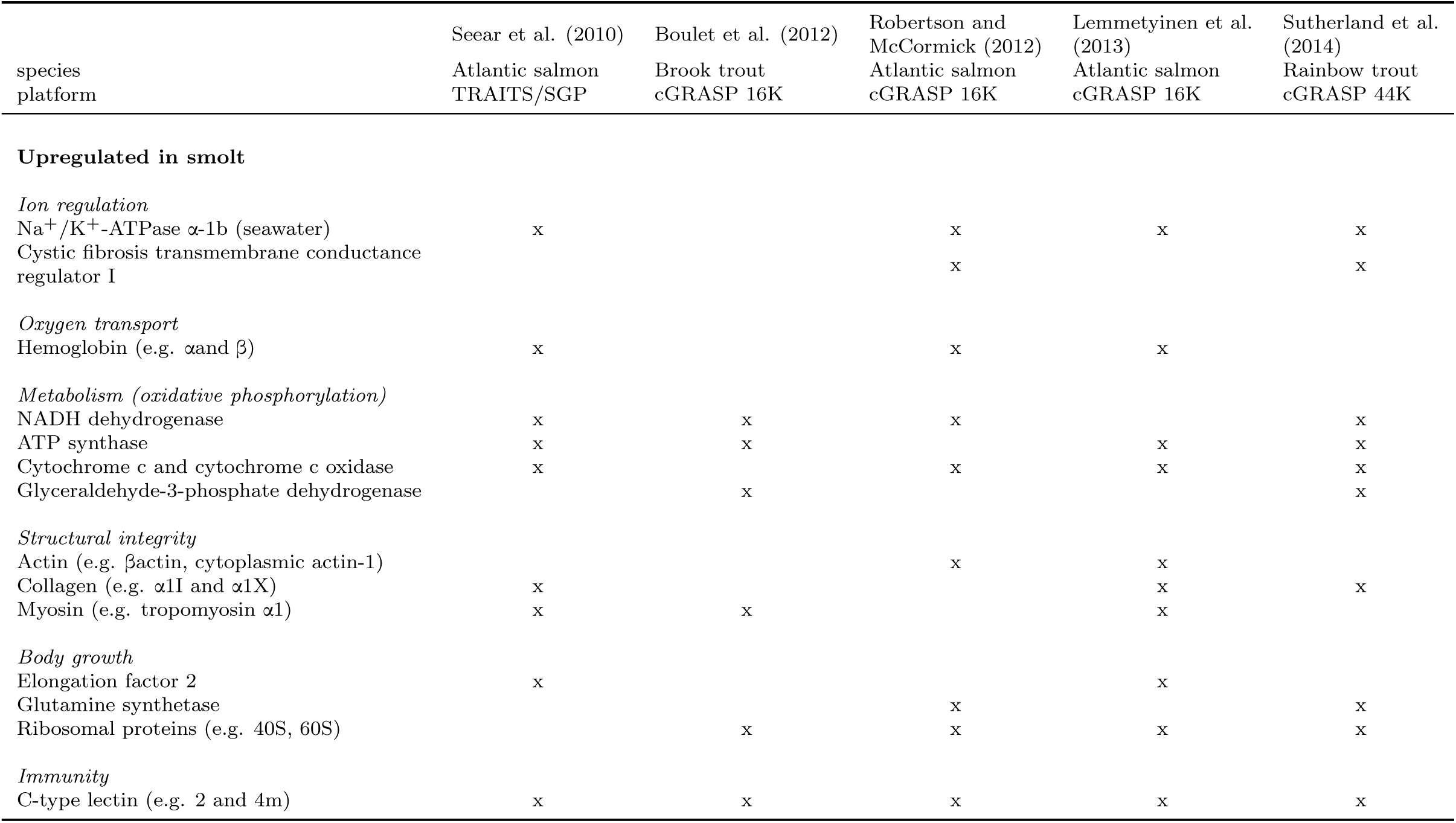
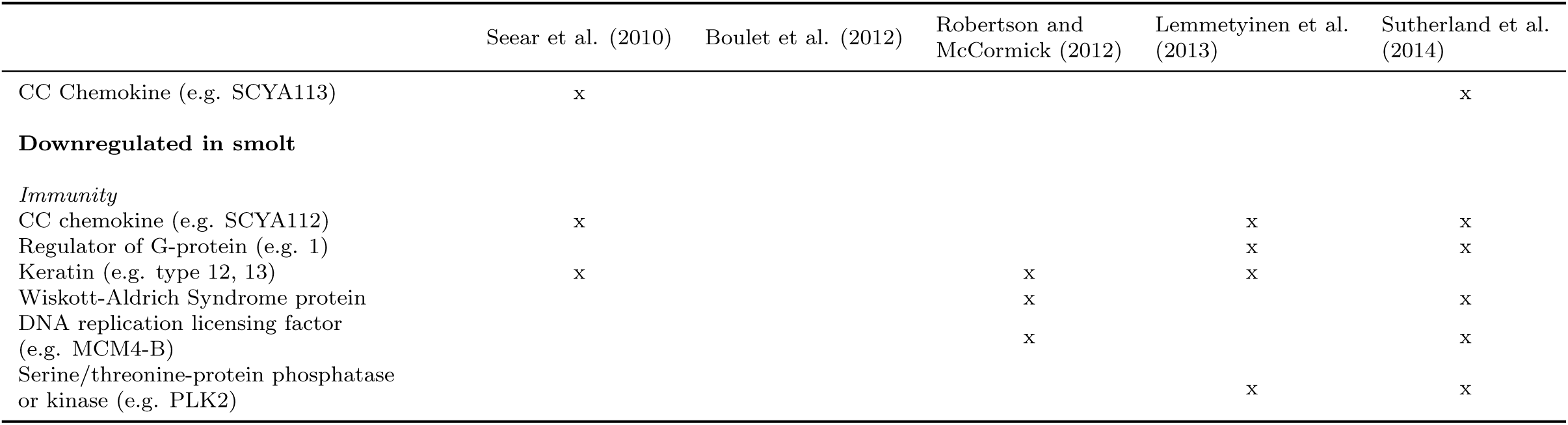
Summary of the gene names associated with smoltification across published microarray studies using gill tissue. Presented are generalized gene names organized by smoltification functional group for microarray studies including Atlantic salmon (*Salmo salar)*, Brook trout (*Salvelinus fontinalis*), and Rainbow trout (*Oncorhynchus mykiss*). The symbol x indicates that a gene name was significant for separating parr and smolt.

**Table S4:**
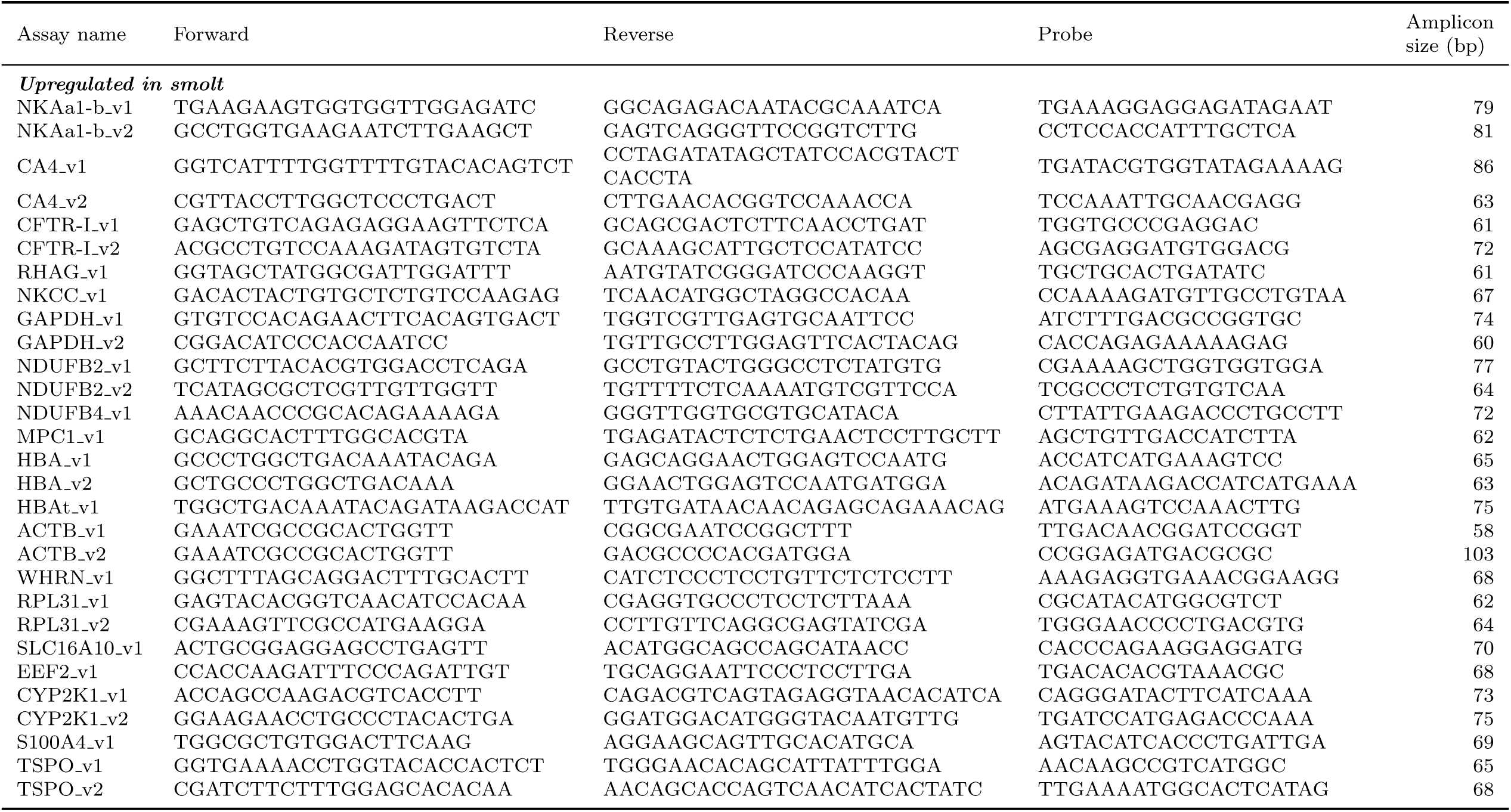
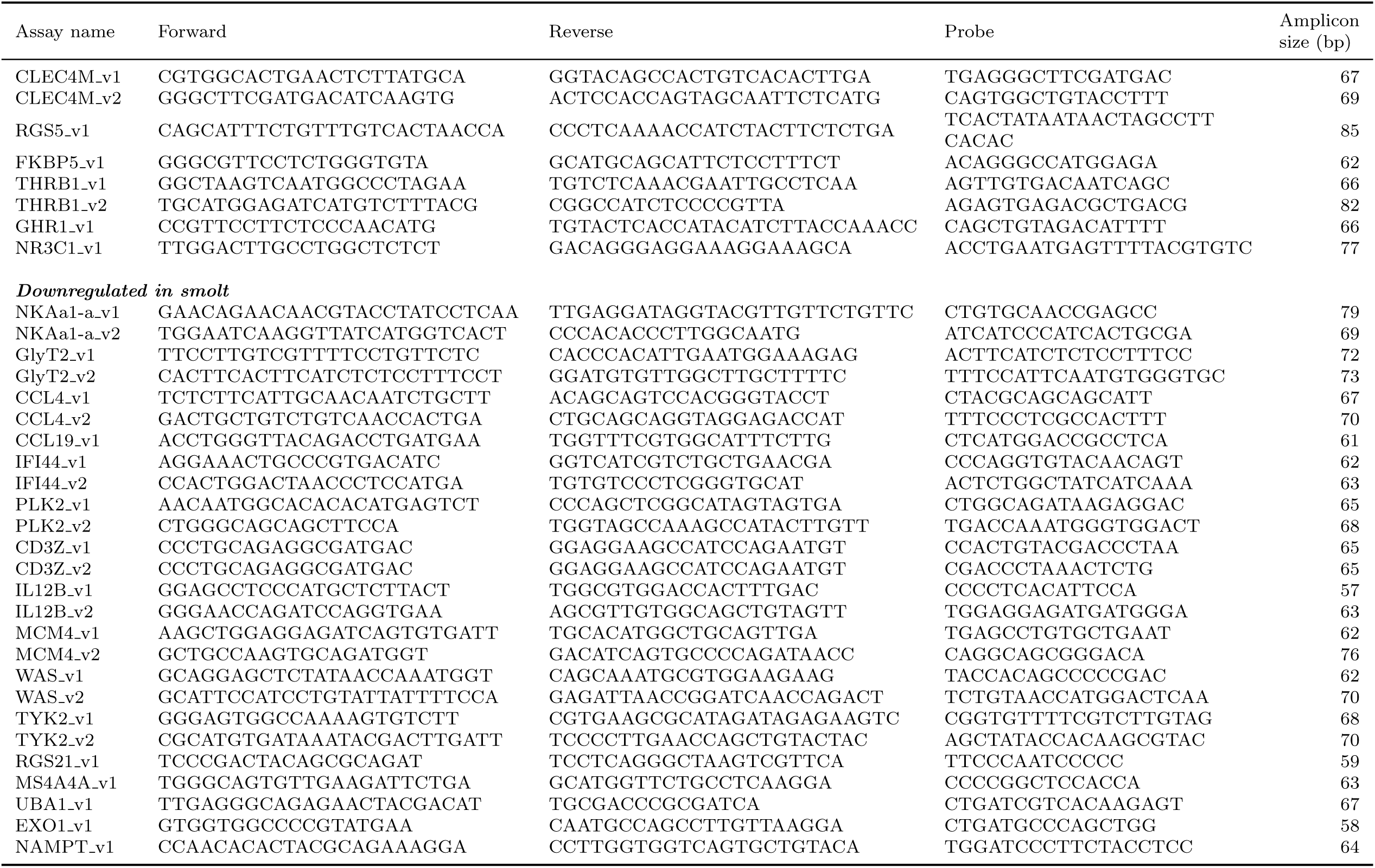
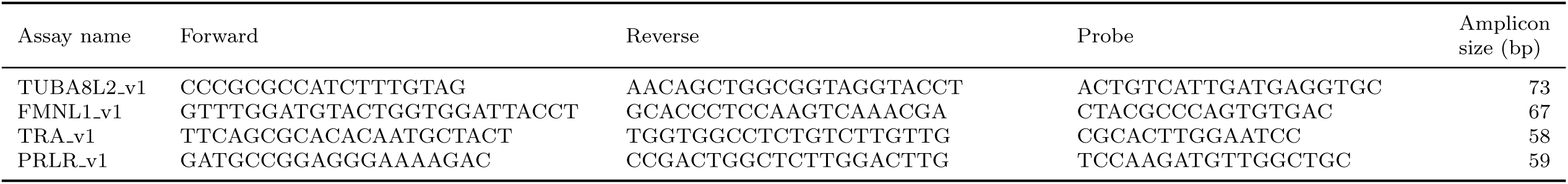
Summary of qPCR TaqMan assay designs for candidate smoltification genes. Presented are the forward, reverse, and TaqMan probe sequences, as well as the amplicon size. Assay names use the symbols described in Table 2 in main text. Two assays were designed for the top 12 upregulated and 10 downregulated genes (set 1), remaining genes had one assay design (set 2): v1 is version 1 and v2 is version 2.

**Table S5:**
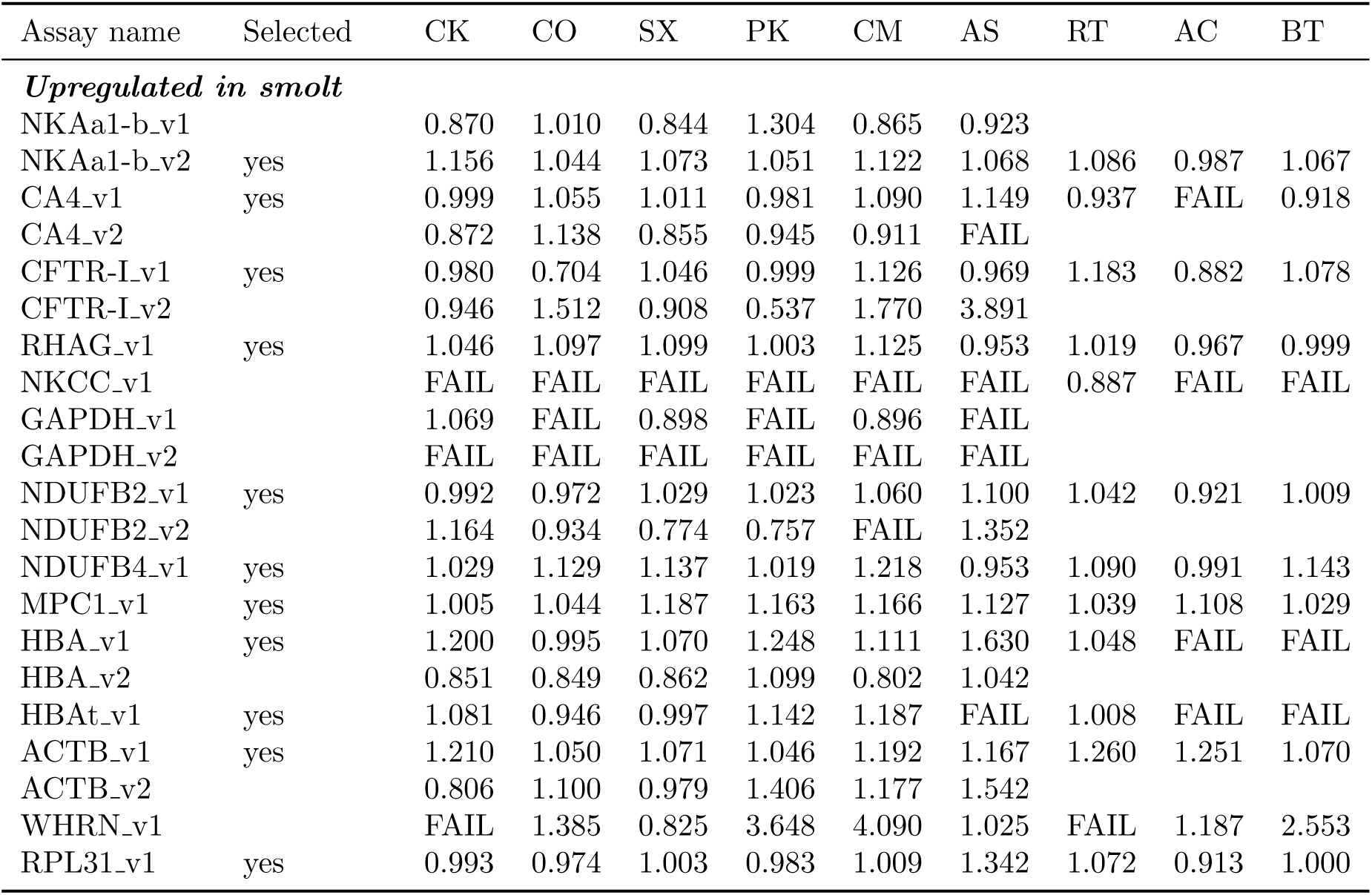
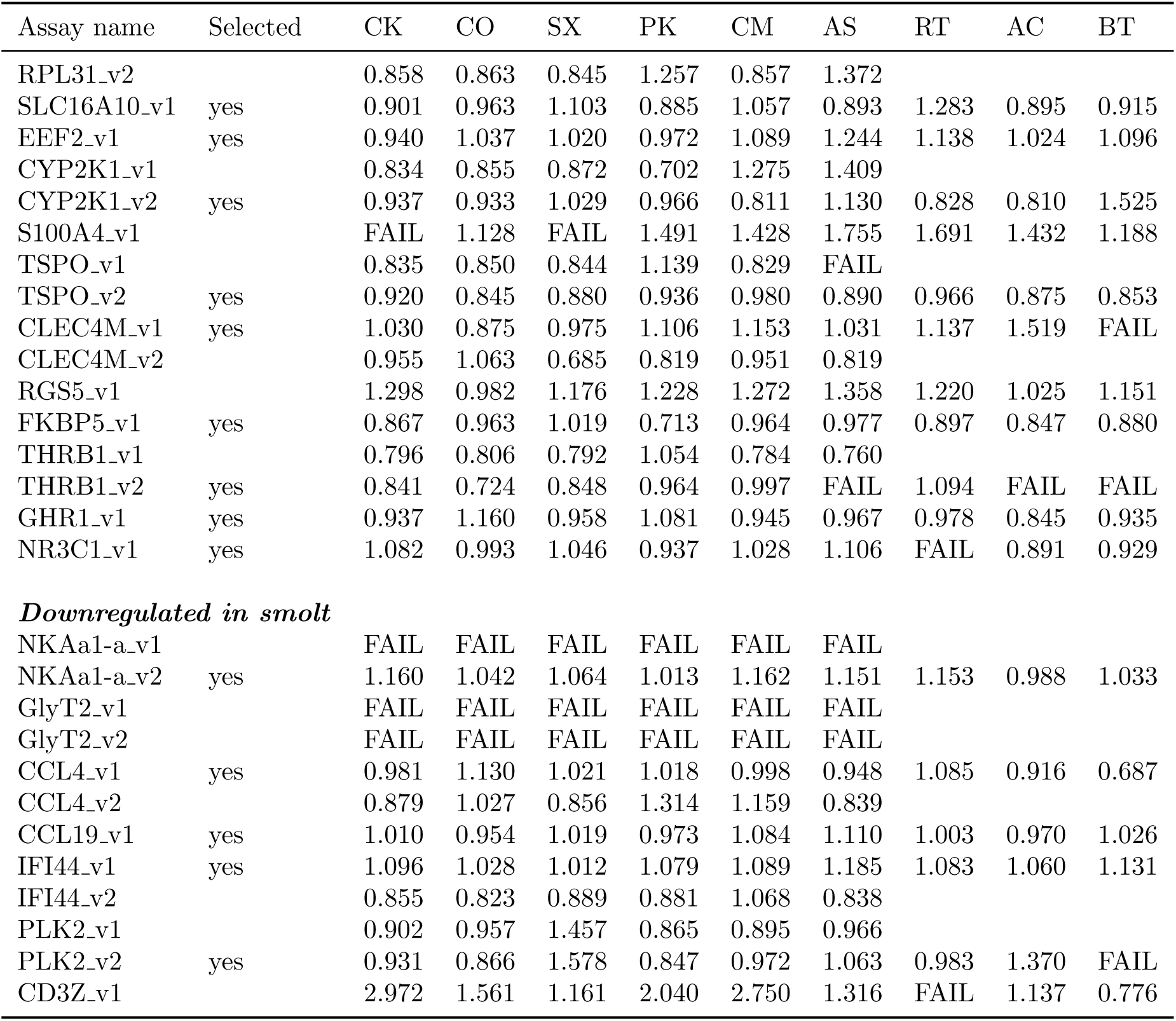
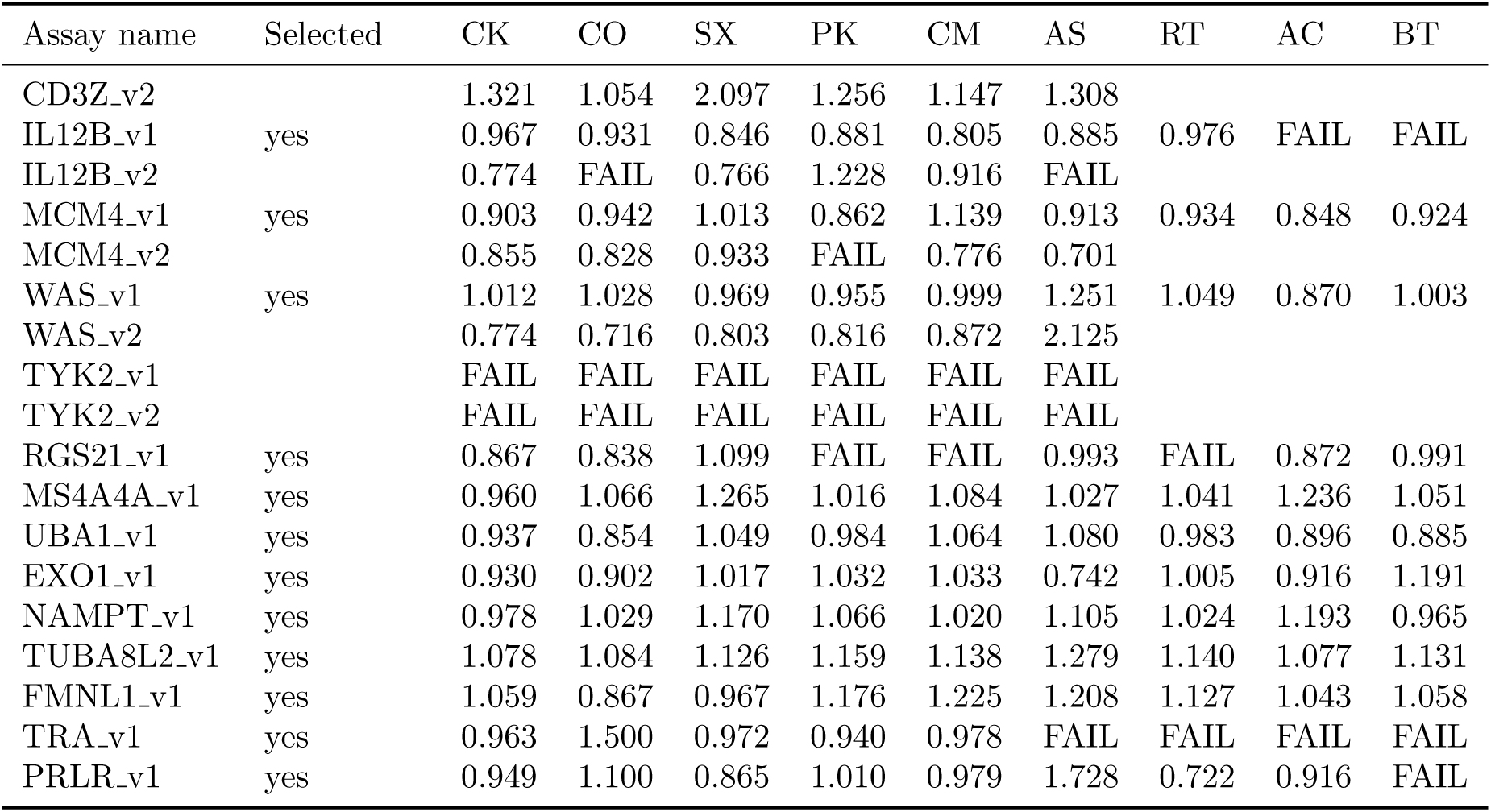
Summary of efficiency values for the qPCR TaqMan assay designs using six to nine salmonid species. Species abbreviations: CK= Chinook salmon (*Oncorhynchus tshawytscha)*, CO= Coho salmon (*O. kisutch*), SX= Sockeye salmon (*O. nerka*), PK= Pink salmon (*O. gorbuscha*), CM= Chum salmon (*O. keta*), AS= Atlantic salmon (*Salmo salar)*, RT= Rainbow trout (*O. mykiss*), AC= Arctic charr (*Salvelinus alpinus)*, and BT= Bull trout (*Salvelinus confluentus)*. Both assay designs of set 1 were tested for efficiency using six species. The best assay design of set 1 and all of set 2 single assay designs were then tested using nine species. Efficiency values between 0.8 and 1.2 were considered good. FAIL indicates no detectable amplification. See Table 2 legend in main text for additional details.

**Table S6:** Summary of the Pearson correlations between PC2 and the expression of 37 candidate smoltification genes using all four groups.

| Assay name | $r$ | $p$ |
| --- | --- | --- |
| ACTB_v1 | -0.03628 | 0.402779 |
| CA4_v1 | 0.650688 | 0 |
| CCL19_v1 | -0.45388 | 0 |
| CCL4_v1 | -0.37056 | 0 |
| CFTR.L_v1 | 0.723941 | 0 |
| CLEC4M_v1 | 0.343777 | 4.44E-16 |
| CYP2K1_v2 | 0.226706 | 1.19E-07 |
| EEF2_v1 | 0.414287 | 0 |
| EXO1_v1 | -0.16926 | 8.48E-05 |
| FKBP5_v1 | 0.518222 | 0 |
| FMNL1_v1 | -0.3929 | 0 |
| GHR1_v1 | 0.199101 | 3.54E-06 |
| HBA_v1 | 0.624489 | 0 |
| HBA_t_v1 | -0.13112 | 0.002398 |
| IFI44_v1 | -0.56695 | 0 |
| IL12B_v1 | -0.32228 | 2.26E-14 |
| MCM4_v1 | -0.00672 | 0.876895 |
| MPC1_v1 | 0.147246 | 0.000642 |
| MS4A4A_v1 | -0.23536 | 3.72E-08 |
| NAMPT_v1 | -0.16426 | 0.000137 |
| NDUFB2_v1 | 0.302671 | 8.92E-13 |
| NDUFB4_v1 | 0.232095 | 5.79E-08 |
| NKAa1.a_v2 | -0.09342 | 0.030888 |
| NKAa1.b_v2 | 0.339919 | 6.66E-16 |
| NR3C1_v1 | -0.14933 | 0.000536 |
| PLK2_v2 | -0.15146 | 0.000445 |
| PRLR_v1 | -0.17944 | 3.04E-05 |
| RGS21_v1 | -0.10182 | 0.018594 |
| RHAG_v1 | 0.244954 | 9.76E-09 |
| RPL31_v1 | 0.06362 | 0.14205 |
| SLC16A10_v1 | 0.103012 | 0.017256 |
| THRB1_v2 | 0.068314 | 0.114846 |
| TRA_v1 | -0.22363 | 1.77E-07 |
| TSPO_v2 | 0.133854 | 0.001936 |
| TUBA8L2_v1 | -0.04732 | 0.275042 |
| UBA1_v1 | -0.15486 | 0.000328 |
| WAS_v1 | -0.4355 | 0 |

**Table S7:** Summary of the Pearson correlations between PC2 and the expression of 37 candidate smoltification genes for Coho salmon.

| Assay name | $r$ | $p$ |
| --- | --- | --- |
| ACTB_v1 | -0.1777 | 0.026462 |
| CA4_v1 | -0.86386 | 0 |
| CCL19_v1 | 0.243586 | 0.002182 |
| CCL4_v1 | 0.178953 | 0.025403 |
| CFTR.L_v1 | -0.88924 | 0 |
| CLEC4M_v1 | -0.0952 | 0.23714 |
| CYP2K1_v2 | -0.22418 | 0.004903 |
| EEF2_v1 | -0.50957 | 1.09E-11 |
| EXO1_v1 | 0.118789 | 0.139682 |
| FKBP5_v1 | -0.58557 | 8.88E-16 |
| FMNL1_v1 | 0.399286 | 2.43E-07 |
| GHR1_v1 | -0.1223 | 0.128273 |
| HBA_v1 | -0.7775 | 0 |
| HBA_t_v1 | -0.66312 | 0 |
| IFI44_v1 | 0.377681 | 1.17E-06 |
| IL12B_v1 | 0.232621 | 0.003475 |
| MCM4_v1 | 0.435372 | 1.35E-08 |
| MPC1_v1 | -0.03564 | 0.658675 |
| MS4A4A_v1 | 0.107597 | 0.181235 |
| NAMPT_v1 | 0.266893 | 0.000757 |
| NDUFB2_v1 | -0.42869 | 2.36E-08 |
| NDUFB4_v1 | -0.38738 | 5.86E-07 |
| NKAa1.a_v2 | -0.25199 | 0.001507 |
| NKAa1.b_v2 | -0.68527 | 0 |
| NR3C1_v1 | 0.053141 | 0.509986 |
| PLK2_v2 | 0.000665 | 0.993427 |
| PRLR_v1 | -0.22419 | 0.0049 |
| RGS21_v1 | -0.23058 | 0.003781 |
| RHAG_v1 | -0.73892 | 0 |
| RPL31_v1 | -0.40186 | 2.00E-07 |
| SLC16A10_v1 | -0.39925 | 2.43E-07 |
| THRB1_v2 | -0.03892 | 0.629536 |
| TRA_v1 | 0.107788 | 0.180456 |
| TSPO_v2 | -0.05689 | 0.480554 |
| TUBA8L2_v1 | 0.054845 | 0.49649 |
| UBA1_v1 | 0.351538 | 6.80E-06 |
| WAS_v1 | 0.378703 | 1.09E-06 |

**Table S8:** Summary of the Pearson correlations between PC2 and the expression of 37 candidate smoltification genes for Sockeye salmon.

| Assay name | $r$ | $p$ |
| --- | --- | --- |
| ACTB_v1 | 0.082718 | 0.468612 |
| CA4_v1 | 0.793261 | 0 |
| CCL19_v1 | -0.22605 | 0.045165 |
| CCL4_v1 | -0.35073 | 0.00153 |
| CFTR.L_v1 | 0.832734 | 0 |
| CLEC4M_v1 | -0.38243 | 0.000505 |
| CYP2K1_v2 | 0.327642 | 0.003203 |
| EEF2_v1 | -0.51509 | 1.19E-06 |
| EXO1_v1 | -0.02941 | 0.796958 |
| FKBP5_v1 | 0.232226 | 0.039454 |
| FMNL1_v1 | -0.24315 | 0.030835 |
| GHR1_v1 | 0.490326 | 4.50E-06 |
| HBA_v1 | 0.813886 | 0 |
| HBA_t_v1 | 0.682975 | 4.10E-12 |
| IFI44_v1 | -0.46057 | 1.95E-05 |
| IL12B_v1 | 0.592312 | 8.93E-09 |
| MCM4_v1 | -0.16452 | 0.147384 |
| MPC1_v1 | 0.454375 | 2.60E-05 |
| MS4A4A_v1 | 0.154063 | 0.175218 |
| NAMPT_v1 | 0.244134 | 0.03014 |
| NDUFB2_v1 | 0.621709 | 9.67E-10 |
| NDUFB4_v1 | 0.515412 | 1.17E-06 |
| NKAa1.a_v2 | 0.211878 | 0.060856 |
| NKAa1.b_v2 | 0.75616 | 8.88E-16 |
| NR3C1_v1 | 0.203574 | 0.071946 |
| PLK2_v2 | 0.09461 | 0.406893 |
| PRLR_v1 | 0.597026 | 6.35E-09 |
| RGS21_v1 | -0.41606 | 0.000137 |
| RHAG_v1 | 0.80588 | 0 |
| RPL31_v1 | -0.25337 | 0.024257 |
| SLC16A10_v1 | 0.398443 | 0.000276 |
| THRB1_v2 | 0.245944 | 0.028901 |
| TRA_v1 | -0.44742 | 3.56E-05 |
| TSPO_v2 | 0.733132 | 1.55E-14 |
| TUBA8L2_v1 | 0.265784 | 0.017912 |
| UBA1_v1 | -0.0791 | 0.488367 |
| WAS_v1 | -0.11068 | 0.331505 |

**Table S9:** Summary of the Pearson correlations between PC2 and the expression of 37 candidate smoltification genes for stream-type Chinook salmon.

| Assay name | $r$ | $p$ |
| --- | --- | --- |
| ACTB_v1 | 0.063431 | 0.47514 |
| CA4_v1 | -0.58541 | 3.19E-13 |
| CCL19_v1 | 0.709524 | 0 |
| CCL4_v1 | 0.621435 | 4.00E-15 |
| CFTR.L_v1 | -0.24342 | 0.00544 |
| CLEC4M_v1 | -0.33614 | 9.84E-05 |
| CYP2K1_v2 | 0.157624 | 0.074423 |
| EEF2_v1 | -0.31222 | 0.000316 |
| EXO1_v1 | 0.321477 | 0.000203 |
| FKBP5_v1 | -0.60248 | 4.24E-14 |
| FMNL1_v1 | 0.275064 | 0.001606 |
| GHR1_v1 | -0.22987 | 0.008777 |
| HBA_v1 | -0.56708 | 2.45E-12 |
| HBA_t_v1 | -0.44971 | 8.97E-08 |
| IFI44_v1 | 0.746626 | 0 |
| IL12B_v1 | 0.385703 | 6.36E-06 |
| MCM4_v1 | 0.308279 | 0.000379 |
| MPC1_v1 | -0.16822 | 0.056697 |
| MS4A4A_v1 | 0.196692 | 0.025475 |
| NAMPT_v1 | 0.132866 | 0.133346 |
| NDUFB2_v1 | -0.13715 | 0.121179 |
| NDUFB4_v1 | -0.15982 | 0.070426 |
| NKAa1.a_v2 | -0.09251 | 0.297078 |
| NKAa1.b_v2 | -0.1793 | 0.042038 |
| NR3C1_v1 | 0.071568 | 0.420253 |
| PLK2_v2 | 0.101785 | 0.251057 |
| PRLR_v1 | -0.01846 | 0.835538 |
| RGS21_v1 | 0.224464 | 0.010548 |
| RHAG_v1 | -0.12244 | 0.166865 |
| RPL31_v1 | 0.051096 | 0.56525 |
| SLC16A10_v1 | 0.033999 | 0.702085 |
| THRB1_v2 | -0.2421 | 0.005707 |
| TRA_v1 | 0.462402 | 3.45E-08 |
| TSPO_v2 | -0.01029 | 0.90786 |
| TUBA8L2_v1 | 0.157099 | 0.075406 |
| UBA1_v1 | 0.10494 | 0.236586 |
| WAS_v1 | 0.308611 | 0.000373 |

**Table S10:** Summary of the Pearson correlations between PC2 or PC3 and the expression of 37 candidate smoltification genes for ocean-type Chinook salmon.

| Assay name | $r$ (PC2) | $p$ (PC2) | $r$ (PC3) | $p$ (PC3) |
| --- | --- | --- | --- | --- |
| ACTB_v1 | 0.295468 | 9.16E-05 | 0.111455 | 0.147902 |
| CA4_v1 | -0.82921 | 0 | -0.24564 | 0.001244 |
| CCL19_v1 | -0.45581 | 4.21E-10 | 0.368436 | 7.68E-07 |
| CCL4_v1 | -0.62148 | 0 | 0.380255 | 3.14E-07 |
| CFTR.L_v1 | -0.53029 | 1.03E-13 | -0.3243 | 1.60E-05 |
| CLEC4M_v1 | 0.295834 | 8.97E-05 | -0.3938 | 1.08E-07 |
| CYP2K1_v2 | 0.380222 | 3.15E-07 | -0.33119 | 1.03E-05 |
| EEF2_v1 | 0.131025 | 0.08855 | 0.158466 | 0.039022 |
| EXO1_v1 | 0.24312 | 0.0014 | -0.24796 | 0.001113 |
| FKBP5_v1 | -0.13548 | 0.078149 | 0.53799 | 3.86E-14 |
| FMNL1_v1 | 0.02361 | 0.759903 | -0.05587 | 0.469299 |
| GHR1_v1 | 0.567149 | 8.88E-16 | 0.106027 | 0.168788 |
| HBA_v1 | -0.23681 | 0.001876 | -0.30541 | 5.13E-05 |
| HBA_t_v1 | -0.52606 | 1.75E-13 | 0.144307 | 0.060452 |
| IFI44_v1 | -0.20517 | 0.007275 | 0.51451 | 7.12E-13 |
| IL12B_v1 | -0.57207 | 4.44E-16 | -0.05122 | 0.507131 |
| MCM4_v1 | 0.264691 | 0.000486 | -0.31182 | 3.48E-05 |
| MPC1_v1 | 0.347345 | 3.47E-06 | 0.211596 | 0.005607 |
| MS4A4A_v1 | -0.04324 | 0.575596 | -0.11498 | 0.135411 |
| NAMPT_v1 | 0.057584 | 0.455739 | 0.419977 | 1.19E-08 |
| NDUFB2_v1 | 0.212028 | 0.005508 | 0.142497 | 0.063782 |
| NDUFB4_v1 | 0.281835 | 0.000197 | 0.189455 | 0.013345 |
| NKAa1.a_v2 | -0.0336 | 0.663534 | 0.27528 | 0.00028 |
| NKAa1.b_v2 | -0.26095 | 0.000588 | -0.1643 | 0.032274 |
| NR3C1_v1 | 0.200761 | 0.008663 | 0.094512 | 0.22022 |
| PLK2_v2 | -0.36256 | 1.18E-06 | -0.17241 | 0.024564 |
| PRLR_v1 | -0.25758 | 0.000696 | -0.02923 | 0.70518 |
| RGS21_v1 | -0.36651 | 8.85E-07 | -0.39598 | 9.04E-08 |
| RHAG_v1 | 0.206415 | 0.006921 | -0.14098 | 0.066685 |
| RPL31_v1 | 0.1849 | 0.015784 | -0.10159 | 0.187427 |
| SLC16A10_v1 | 0.251176 | 0.000953 | -0.16253 | 0.034209 |
| THRB1_v2 | -0.20179 | 0.008319 | 0.340011 | 5.72E-06 |
| TRA_v1 | -0.12969 | 0.09187 | -0.34609 | 3.78E-06 |
| TSPO_v2 | -0.22918 | 0.002645 | -0.3487 | 3.16E-06 |
| TUBA8L2_v1 | 0.375566 | 4.50E-07 | 0.279671 | 0.000221 |
| UBA1_v1 | -0.1985 | 0.009462 | 0.396463 | 8.69E-08 |
| WAS_v1 | -0.32037 | 2.05E-05 | -0.01946 | 0.801146 |

**Table S11:** Summary of the Student’s t-tests examining the difference in expression of 37 candidate smoltification genes between freshwater and seawater for ocean-type Chinook salmon. Nitinat juveniles were sampled at the same time in late April in both environments. Seawater juveniles were contained within a netpen in an estuary for about two weeks prior to sampling. Presented are the mean values in freshwater and seawater; *t* and *p*-values for the mean difference; and significance coding for the difference, i.e. *** *p <* 0.001, ** 0.001 *< p <* 0.01, * 0.01 *< p <* 0.05,. 0.05 *< p <* 0.1

| Assay name | Freshwater | Seawater | $t$-value | $p$-value | Signif. |
| --- | --- | --- | --- | --- | --- |
| ACTB_v1 | 0.781499 | 0.614146 | 0.454184 | 0.656659 |  |
| CA4_v1 | -0.16458 | 1.665436 | -5.67612 | 7.47E-05 | *** |
| CCL19_v1 | -1.1124 | -2.10943 | 2.783734 | 0.014903 | * |
| CCL4_v1 | 0.324899 | -2.61555 | 7.859551 | 1.13E-05 | *** |
| CFTR.L_v1 | 0.367682 | 1.632086 | -1.77163 | 0.107445 |  |
| CLEC4M_v1 | -0.72391 | -0.34098 | -0.60563 | 0.5545 |  |
| CYP2K1_v2 | 0.229148 | 0.581438 | -0.42066 | 0.680431 |  |
| EEF2_v1 | 1.00145 | 1.11542 | -0.5718 | 0.576693 |  |
| EXO1_v1 | 0.365374 | 0.453229 | -0.49978 | 0.625311 |  |
| FKBP5_v1 | -0.03752 | 0.537554 | -1.37563 | 0.196721 |  |
| FMNL1_v1 | 0.401456 | 0.076089 | 0.927794 | 0.369291 |  |
| GHR1_v1 | 0.5083 | 0.260142 | 0.467005 | 0.647695 |  |
| HBA_v1 | 1.379431 | 1.06952 | 0.756477 | 0.463339 |  |
| HBA_t_v1 | -0.64149 | -1.82298 | 1.600008 | 0.132927 |  |
| IFI44_v1 | 0.128963 | -0.9276 | 3.608317 | 0.002892 | ** |
| IL12B_v1 | 1.002072 | 0.131237 | 1.740927 | 0.103722 |  |
| MCM4_v1 | 0.276983 | 0.247993 | 0.131698 | 0.897428 |  |
| MPC1_v1 | 0.889465 | 0.567753 | 0.939307 | 0.363611 |  |
| MS4A4A_v1 | 0.973487 | 0.1996 | 2.029614 | 0.061889 | . |
| NAMPT_v1 | 0.92058 | -0.05864 | 2.503539 | 0.025465 | * |
| NDUFB2_v1 | 1.034136 | 0.233148 | 2.567294 | 0.022735 | * |
| NDUFB4_v1 | 0.708509 | 0.322051 | 1.252024 | 0.231117 |  |
| NKAa1.a_v2 | 1.05694 | -1.17179 | 7.392296 | 1.28E-05 | *** |
| NKAa1.b_v2 | 0.312624 | 0.763602 | -1.44866 | 0.169462 |  |
| NR3C1_v1 | 0.876391 | 0.443096 | 1.640321 | 0.123257 |  |
| PLK2_v2 | 1.101163 | 0.811895 | 0.877102 | 0.395861 |  |
| PRLR_v1 | 1.421913 | -0.68146 | 6.365303 | 3.24E-05 | *** |
| RGS21_v1 | -0.32839 | 1.33284 | -6.60176 | 1.19E-05 | *** |
| RHAG_v1 | 1.739602 | 0.695542 | 3.181861 | 0.006896 | ** |
| RPL31_v1 | 0.646804 | 0.496804 | 0.554798 | 0.589087 |  |
| SLC16A10_v1 | 1.488335 | 1.127572 | 0.762769 | 0.460651 |  |
| THRB1_v2 | 1.371714 | 0.648913 | 1.865934 | 0.083299 | . |
| TRA_v1 | 0.125936 | 0.032976 | 0.305942 | 0.764628 |  |
| TSPO_v2 | 1.123814 | 0.046151 | 2.58888 | 0.021507 | * |
| TUBA8L2_v1 | 0.132761 | -0.18029 | 1.625875 | 0.126282 |  |
| UBA1_v1 | 0.745251 | -0.2679 | 3.094738 | 0.008873 | ** |
| WAS_v1 | 0.272223 | -0.28009 | 2.243392 | 0.041572 | * |

**Table S12:**
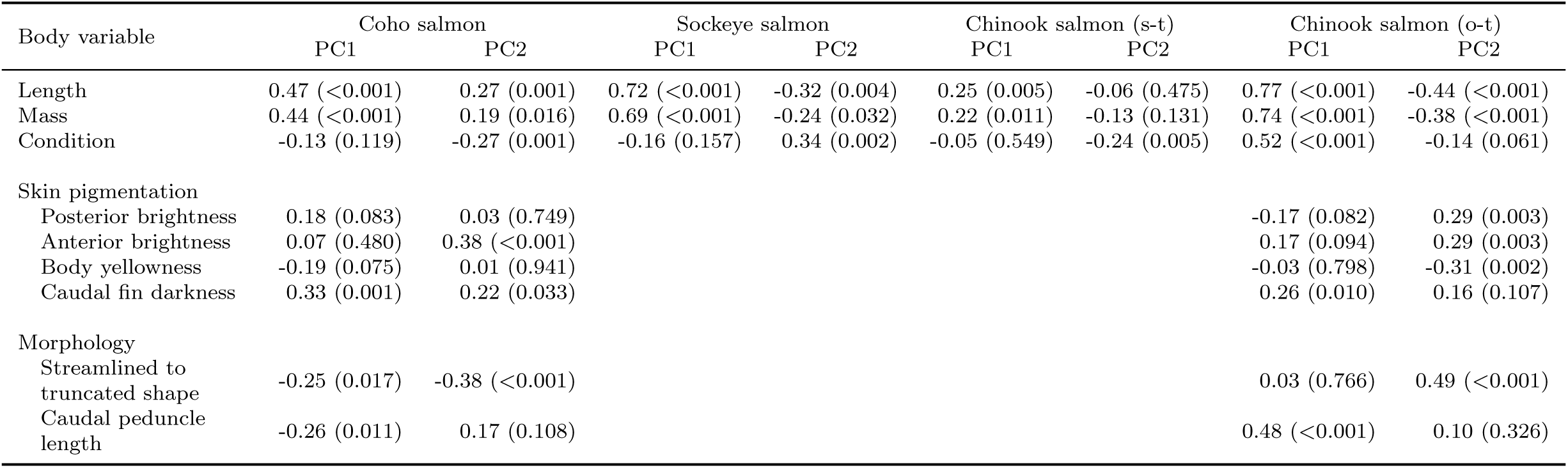
Summary of body variable correlations with smoltification gene expression patterns for the four groups. Presented are the Pearson correlations (*r)* with *p*-values in brackets for body variables and smoltification gene expression patterns (PC1 and PC2) using Coho salmon, Sockeye salmon, stream-type Chinook salmon, and ocean-type Chinook salmon. Smoltification gene expression patterns (PC1 and PC2), using the top 10 biomarkers, for each group are displayed in Figure 2.

**Figure S1:**
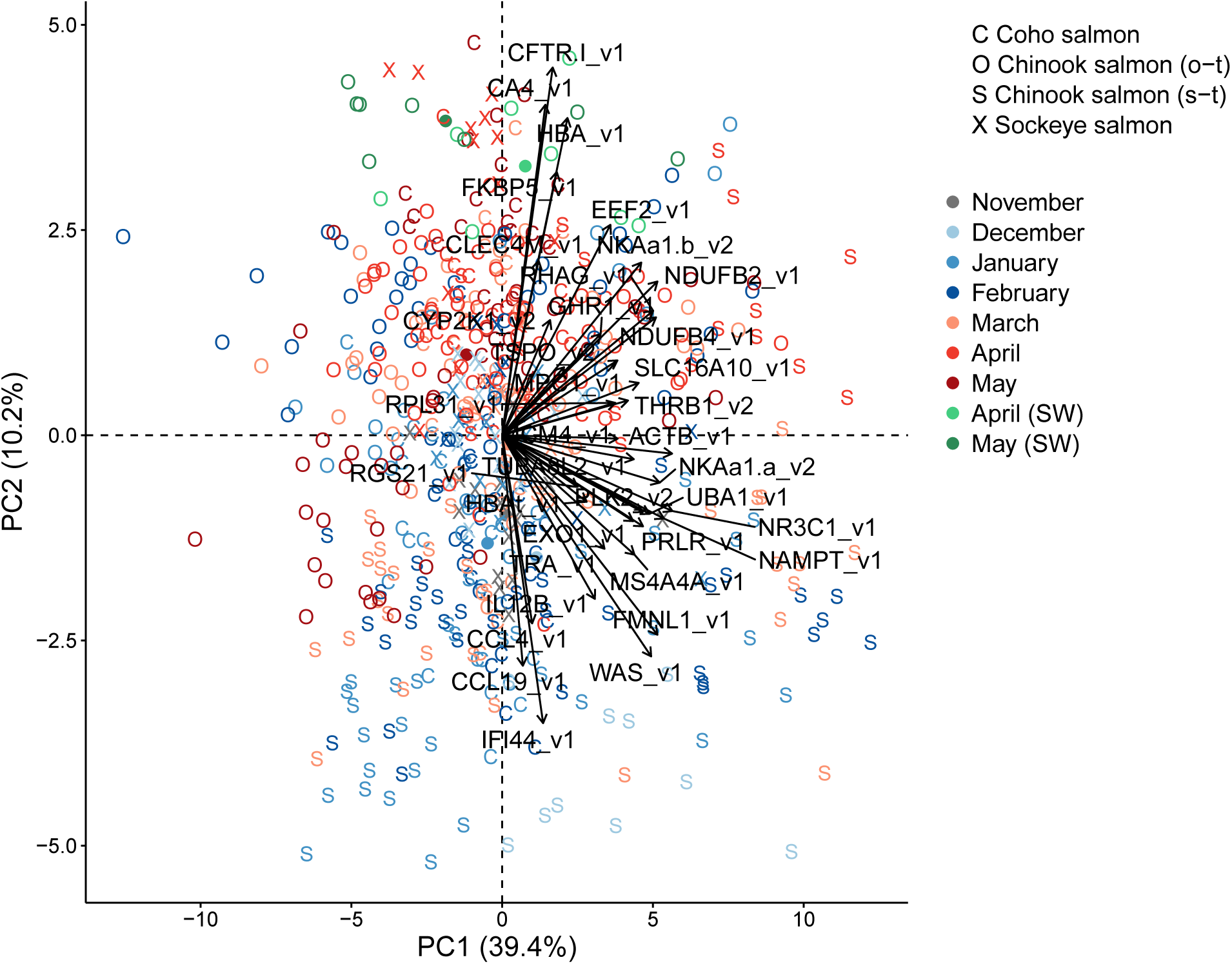
Canonical plots of the first two principal components of all 37 candidate genes for smoltification using all four groups. Groups are Coho salmon (*Oncorhynchus kisutch*), Sockeye salmon, Chinook salmon (stream-type, *O. tshawytscha*), and Chinook salmon (ocean-type, *O. tshawytscha*). See Figure 1 legend.

**Figure S2:**
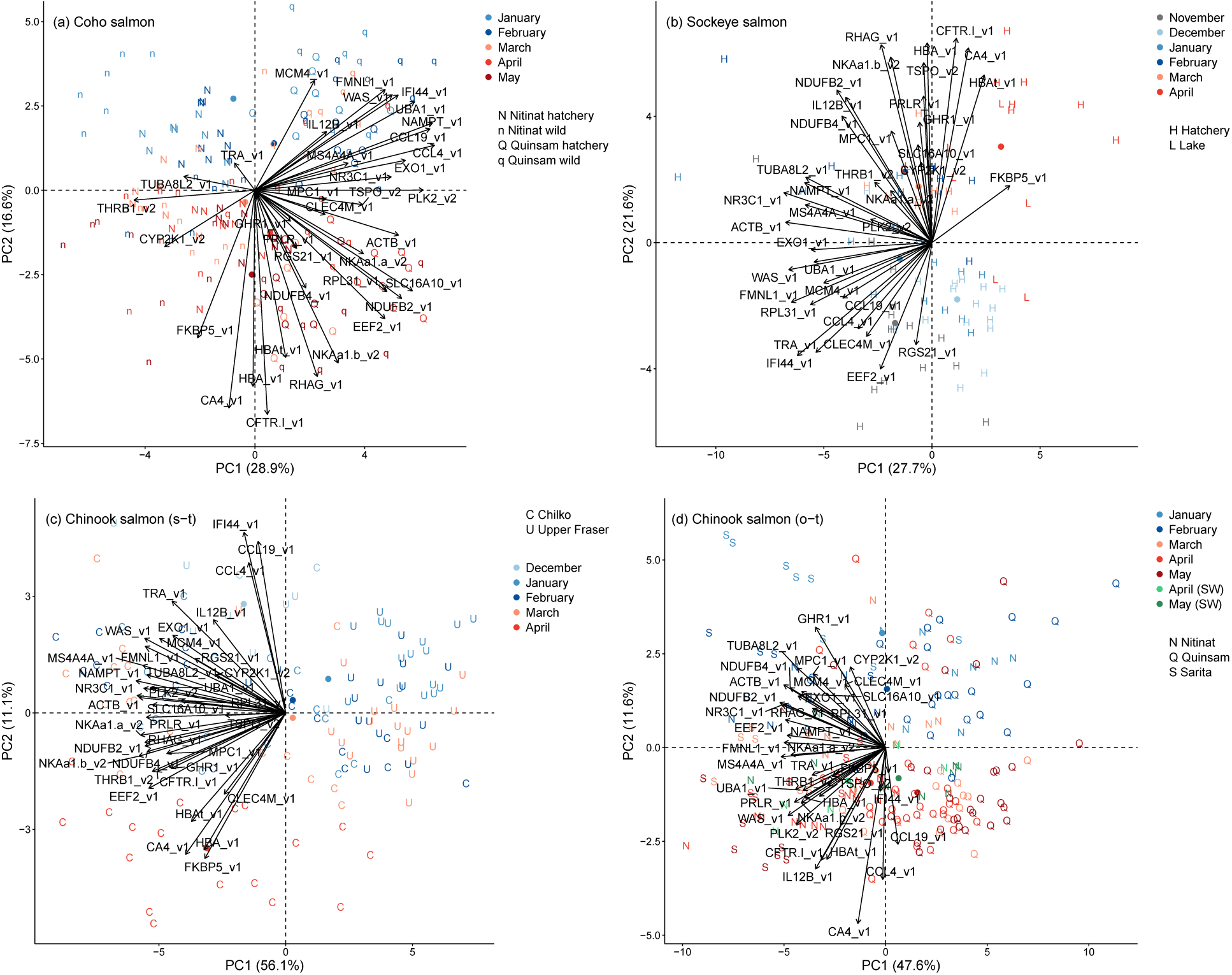
Canonical plots of the first two principal components of all 37 candidate genes for smoltification using each of the four groups. (a) Coho salmon, (b) Sockeye salmon, (c) stream-type Chinook salmon, and (d) ocean-type Chinook salmon. See Figure 1 legend.

**Figure S3:**
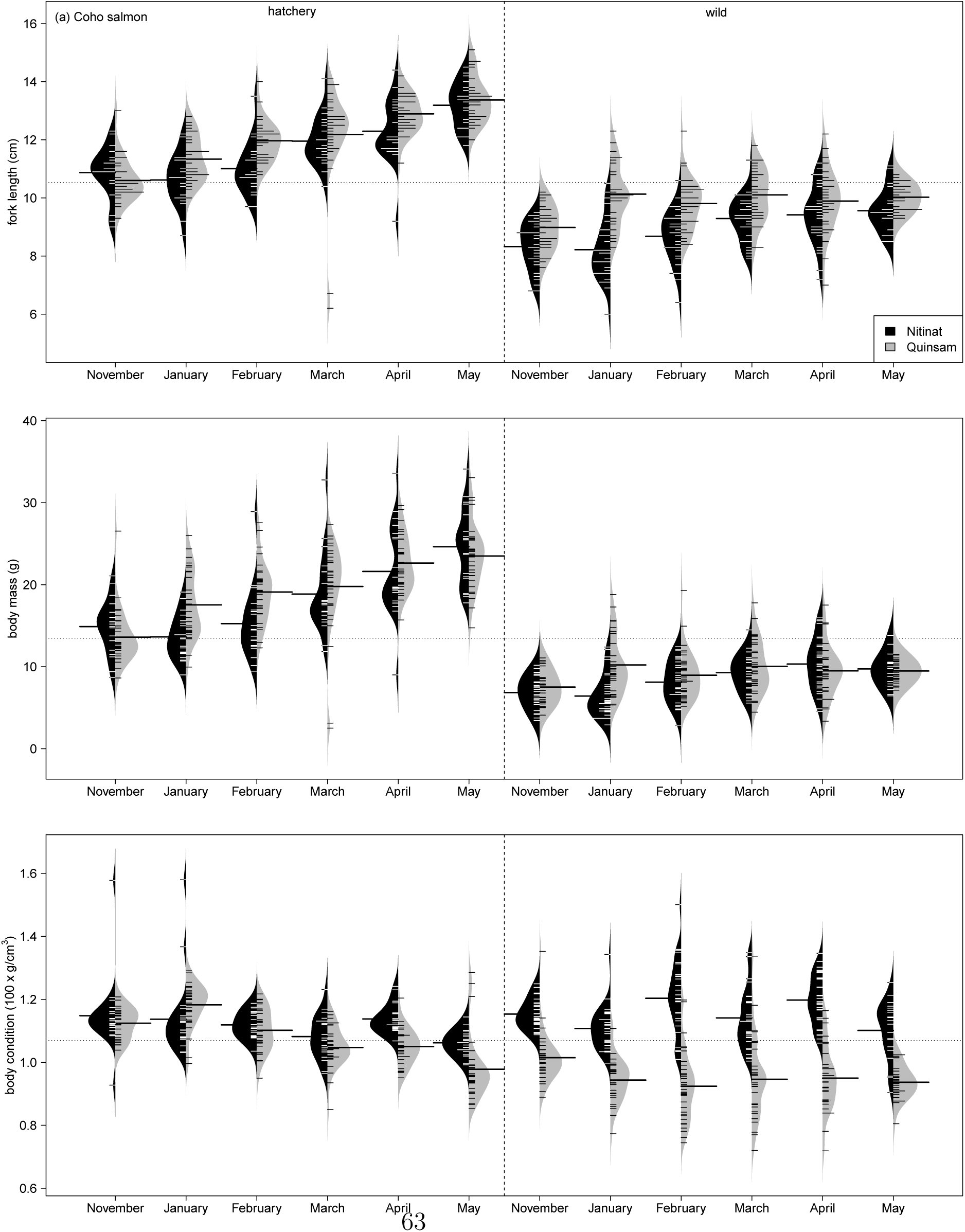
Beanplots of body length, mass, and condition by monthly development for each of the four groups. (a) Coho salmon, (b) Sockeye salmon, (c) stream-type Chinook salmon, and (d) ocean-type Chinook salmon. Solid horizontal line is the mean for each bean; dashed horizontal line is the overall mean across beans. Month symbol E is early and L is late.

**Figure S4:**
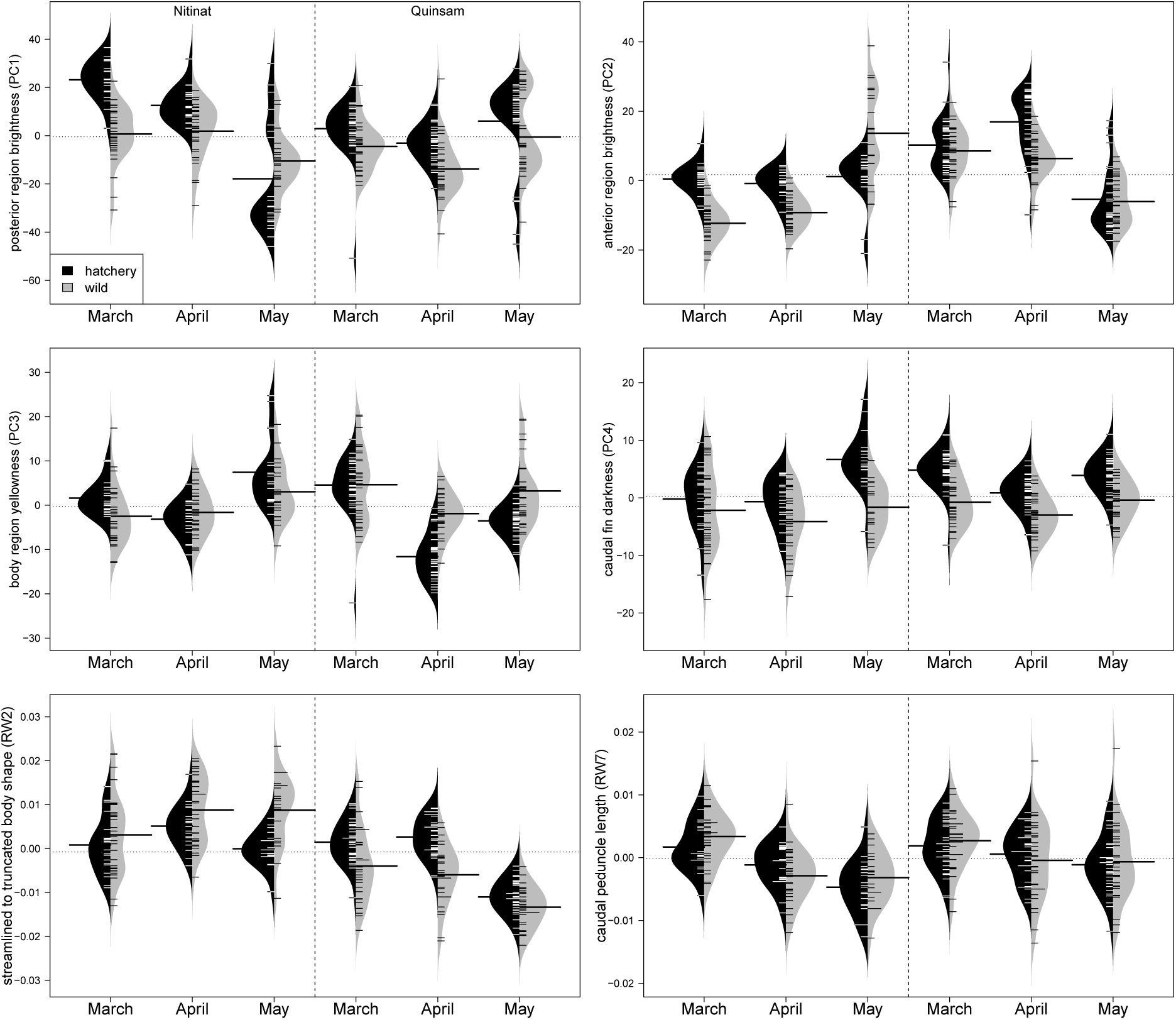
Beanplots of skin pigmentation and body morphology by monthly development for Coho salmon. Solid horizontal line is the mean for each bean; dashed horizontal line is the overall mean across beans.

**Figure S5:**
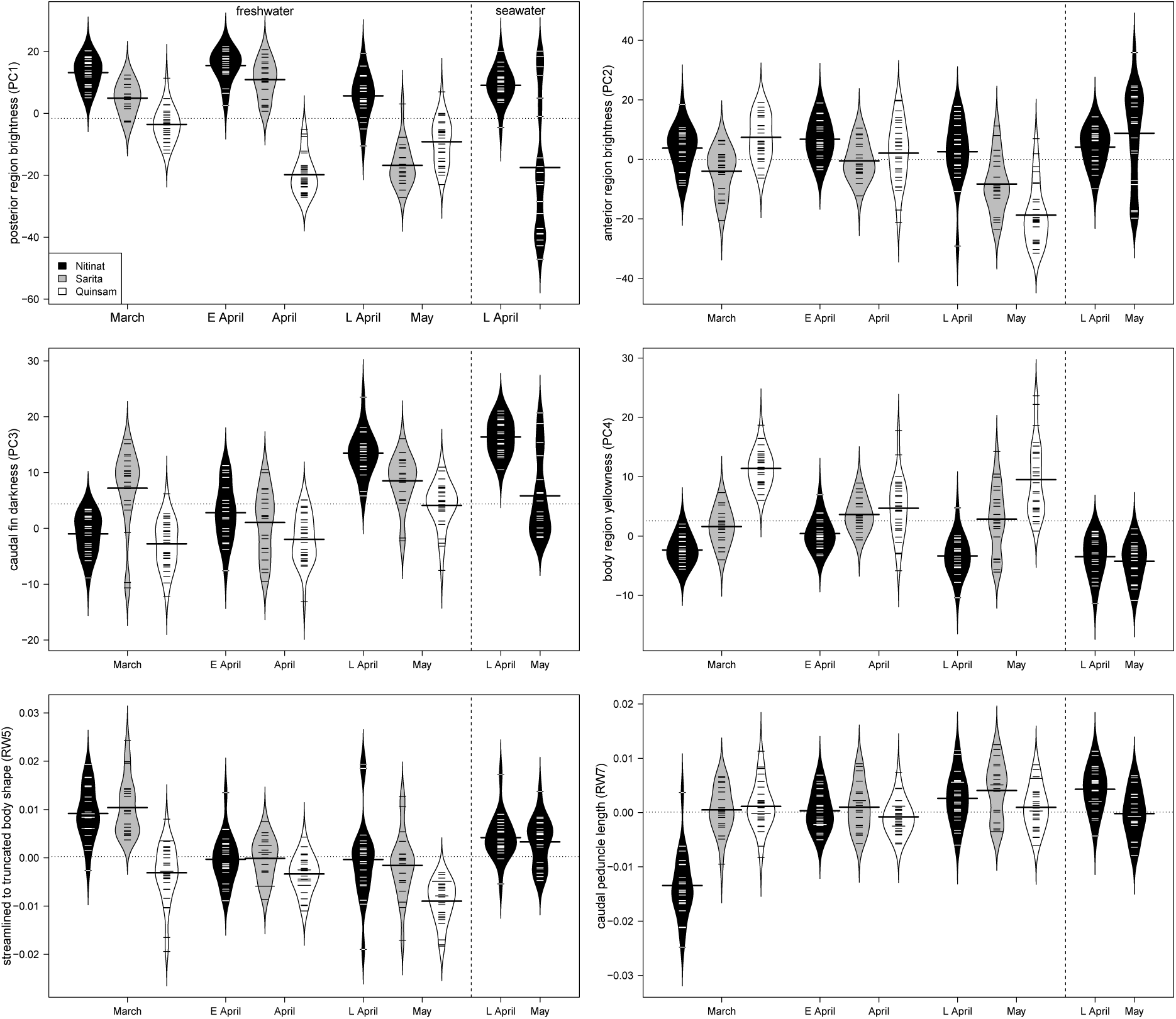
Beanplots of skin pigmentation and body morphology by monthly development for ocean-type Chinook salmon. Solid horizontal line is the mean for each bean; dashed horizontal line is the overall mean across beans. Month symbol E is for early and L is for late.

## Supplementary Appendices

*Available online at BioRxiv: https://doi.org/10.1101/474692

Appendix 1. Beanplots of the expression of 37 candidate smotification genes by monthly development for Coho salmon. Solid horizontal line is the mean for each bean; dashed horizontal line is the overall mean across beans.

Appendix 2. Beanplots of the expression of 37 candidate smotification genes by monthly development for Sockeye salmon. Solid horizontal line is the mean for each bean; dashed horizontal line is the overall mean across beans.

Appendix 3. Beanplots of the expression of 37 candidate smotification genes by monthly development for stream-type Chinook salmon. Solid horizontal line is the mean for each bean; dashed horizontal line is the overall mean across beans.

Appendix 4. Beanplots of the expression of 37 candidate smotification genes by monthly development for ocean-type Chinook salmon. Solid horizontal line is the mean for each bean; dashed horizontal line is the overall mean across beans. Month symbol E is early and L is late.

