## appendix 1 for "Discovery and validation of candidate smoltification gene expression biomarkers across multiple species and ecotypes of Pacific salmonids"

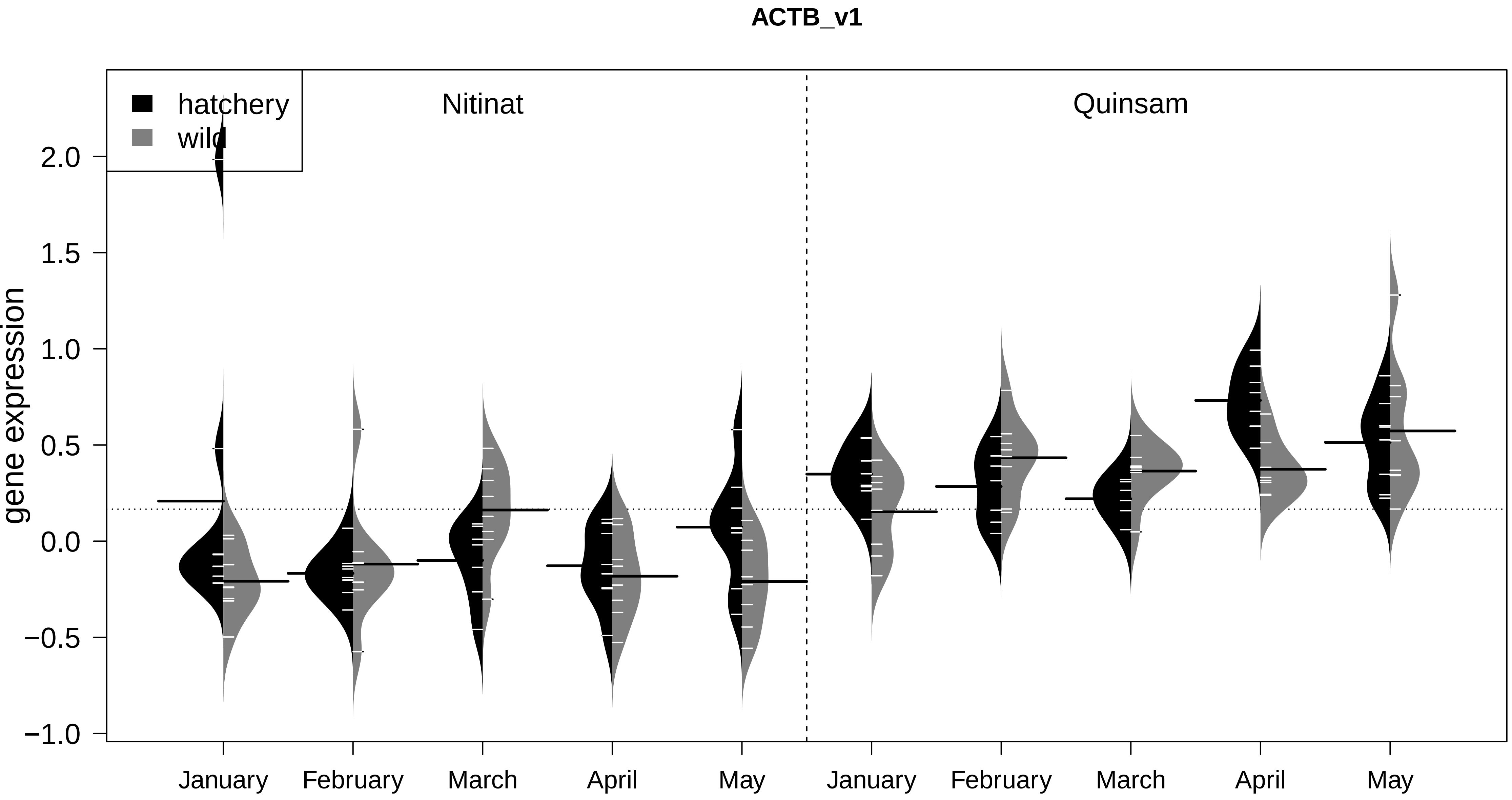
ACTB_v1
hatchery
wild
Nitinat
Quinsam
gene expression
2.0
1.5
1.0
0.5
0.0
−0.5
−1.0
January
February
March
April
May
January
February
March
April
May

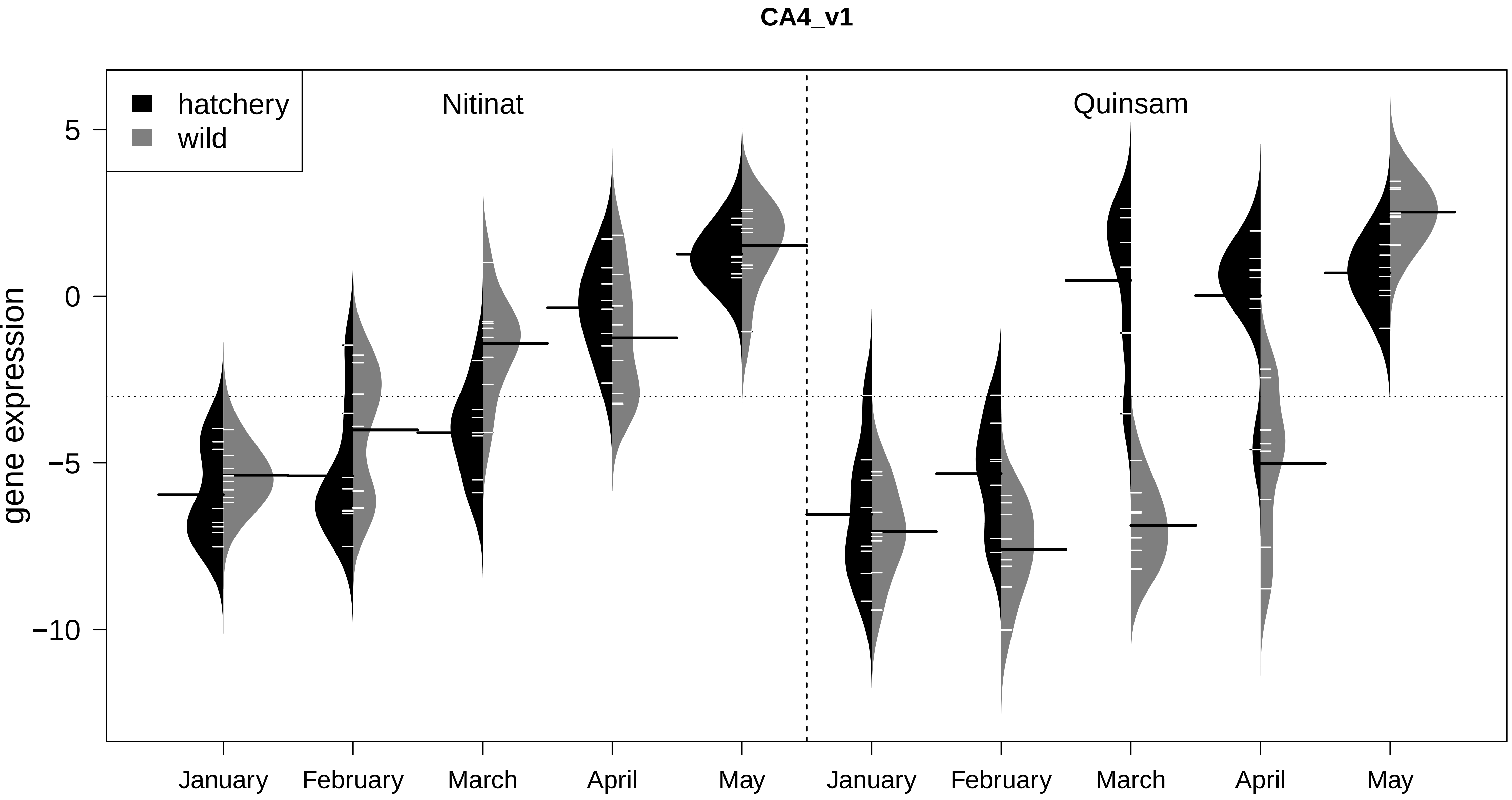

CA4_v1
hatchery
wild
Nitinat
Quinsam
5
0
−5
−10
gene expression
January
February
March
April
May
January
February
March
April
May

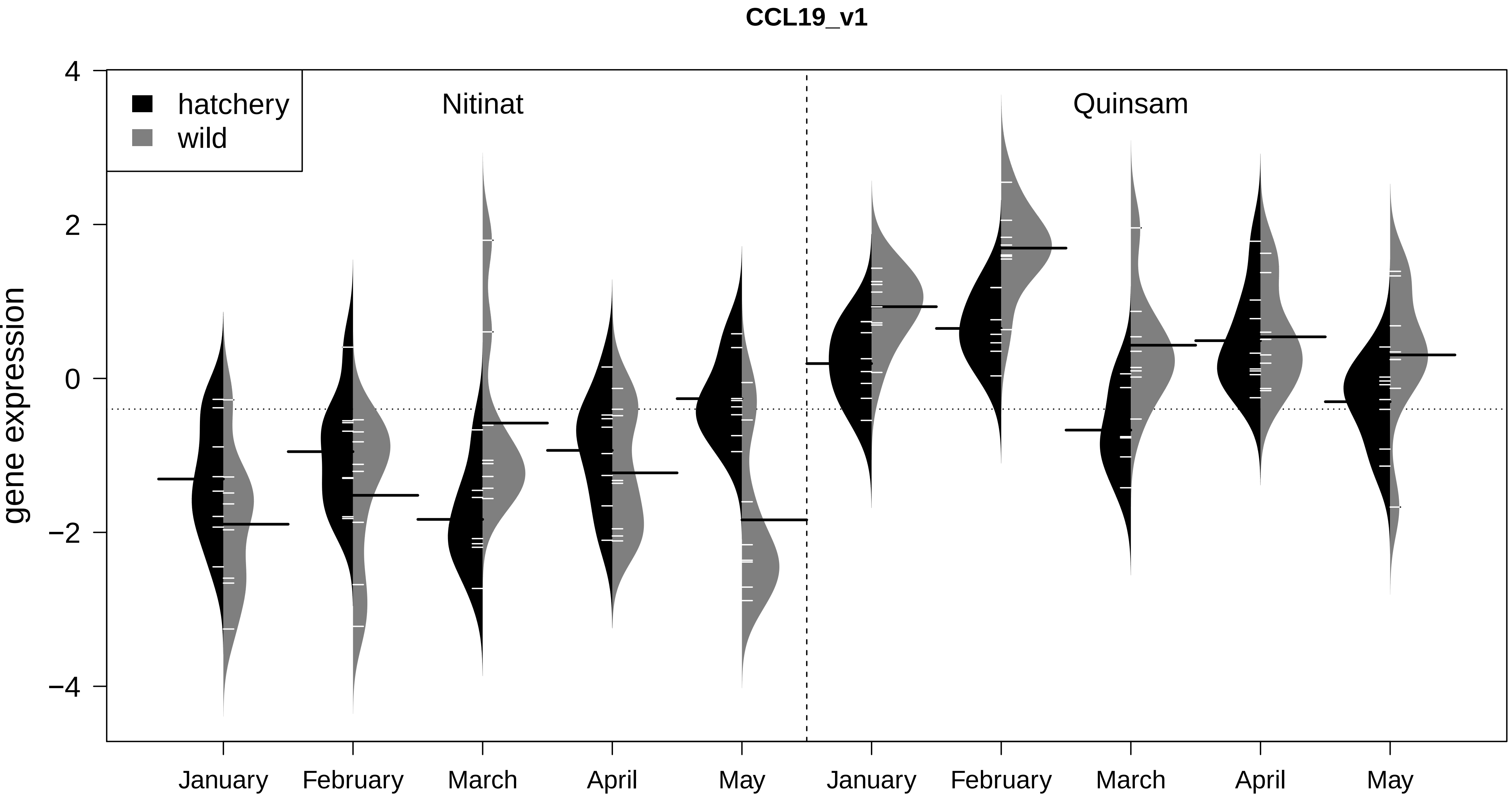

CCL19_v1
hatchery
wild
Nitinat
Quinsam
gene expression
4
2
0
−2
−4
January
February
March
April
May
January
February
March
April
May

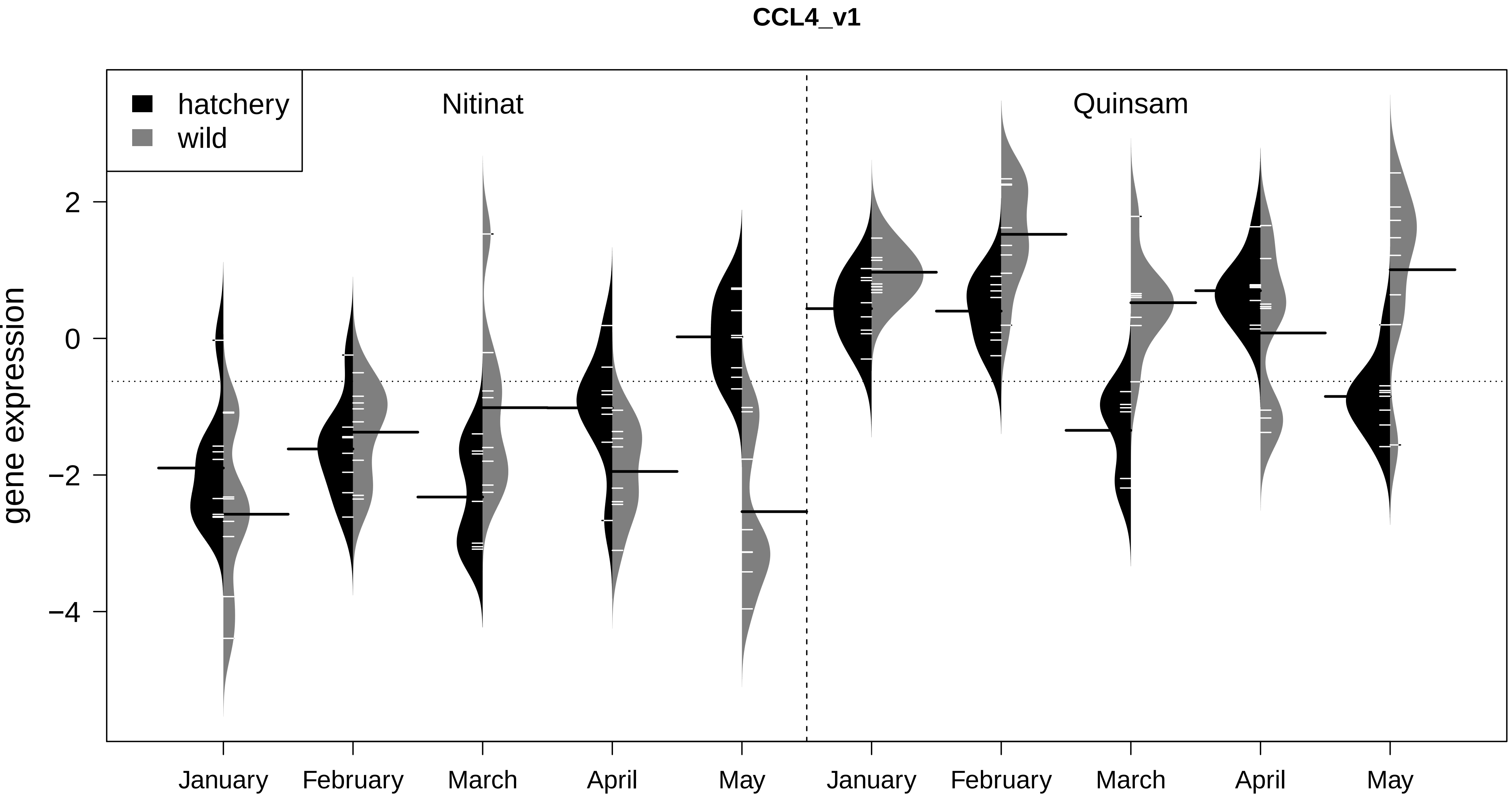

CCL4_v1
hatchery
wild
Nitinat
Quinsam
gene expression
2
0
−2
−4
January
February
March
April
May
January
February
March
April
May

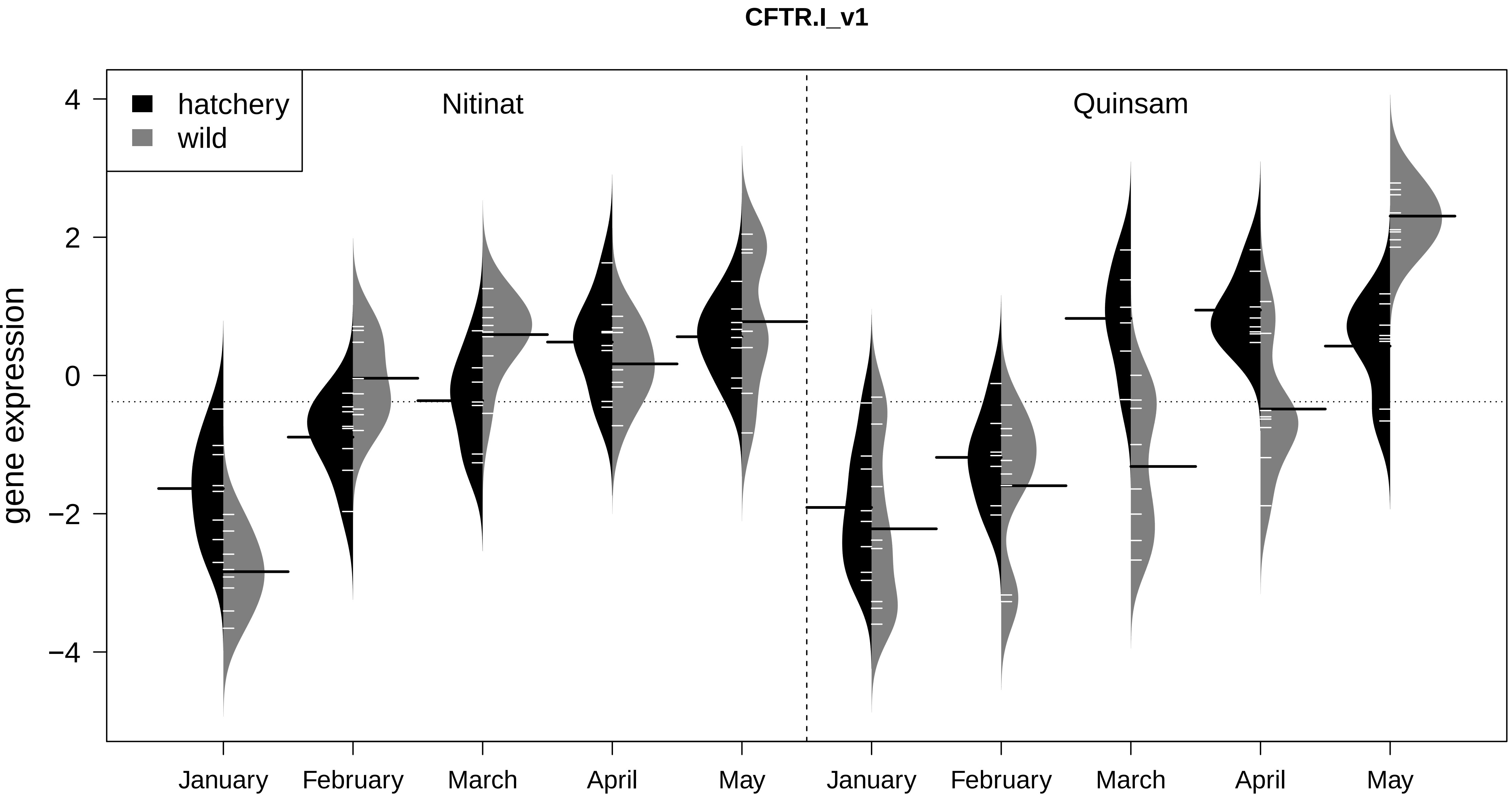

CFTR.I_v1
hatchery
wild
Nitinat
Quinsam
gene expression
4
2
0
−2
−4
January
February
March
April
May
January
February
March
April
May

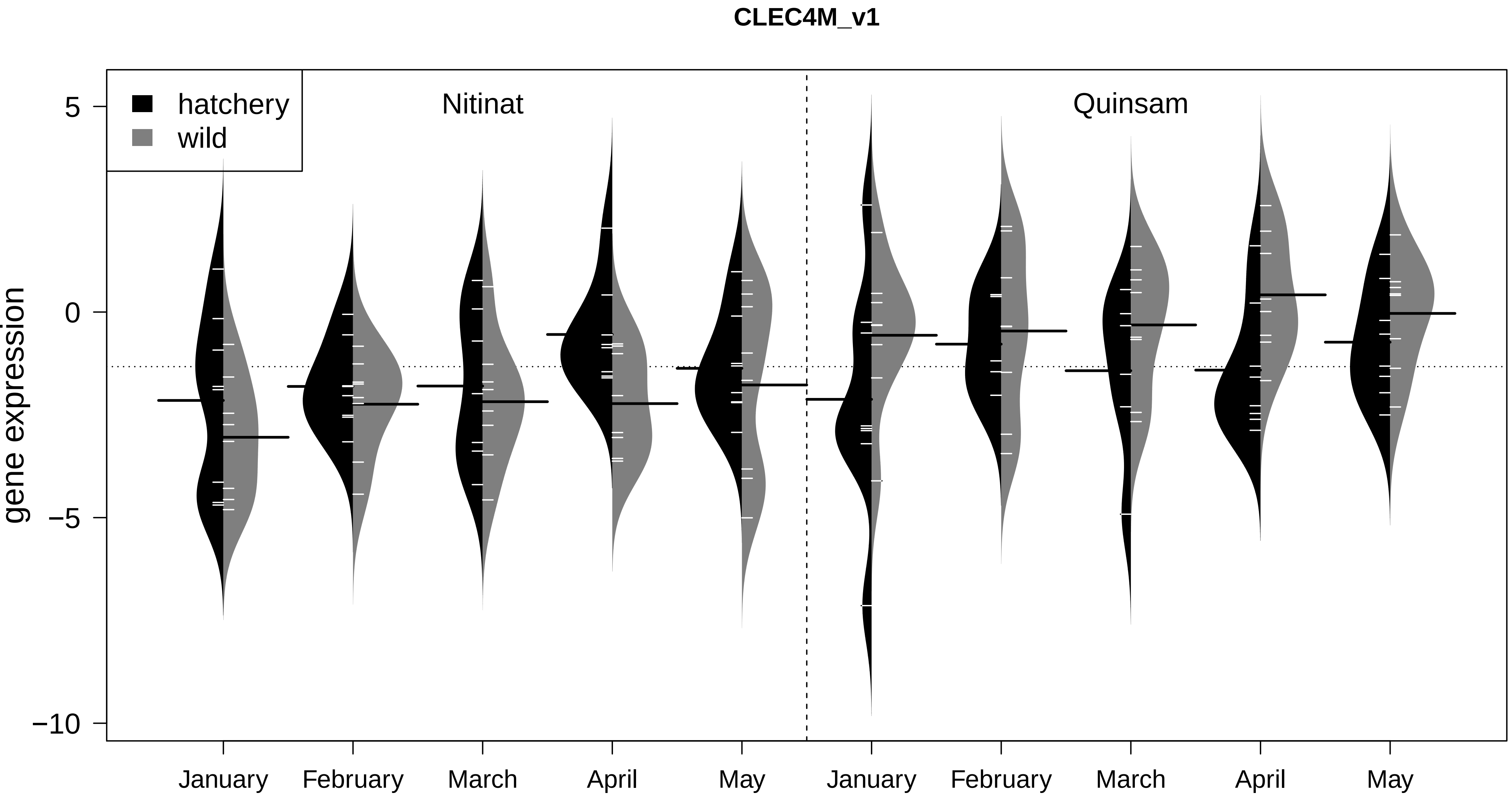
CLEC4M_v1
hatchery
wild
Nitinat
Quinsam
gene expression
5
0
−5
−10
January
February
March
April
May
January
February
March
April
May

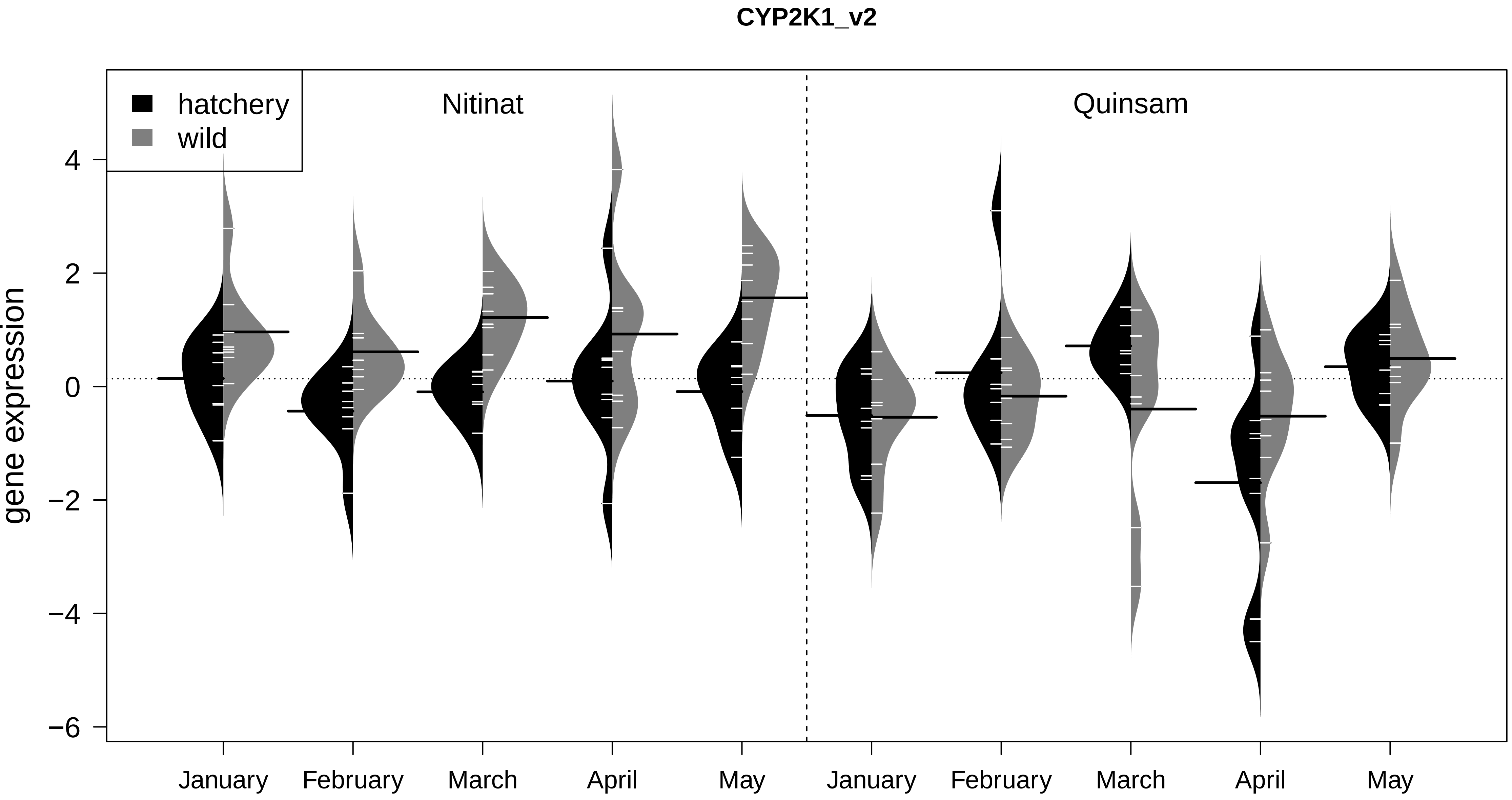
CYP2K1_v2
hatchery
wild
Nitinat
Quinsam
gene expression
4
2
0
−2
−4
−6
January
February
March
April
May
January
February
March
April
May

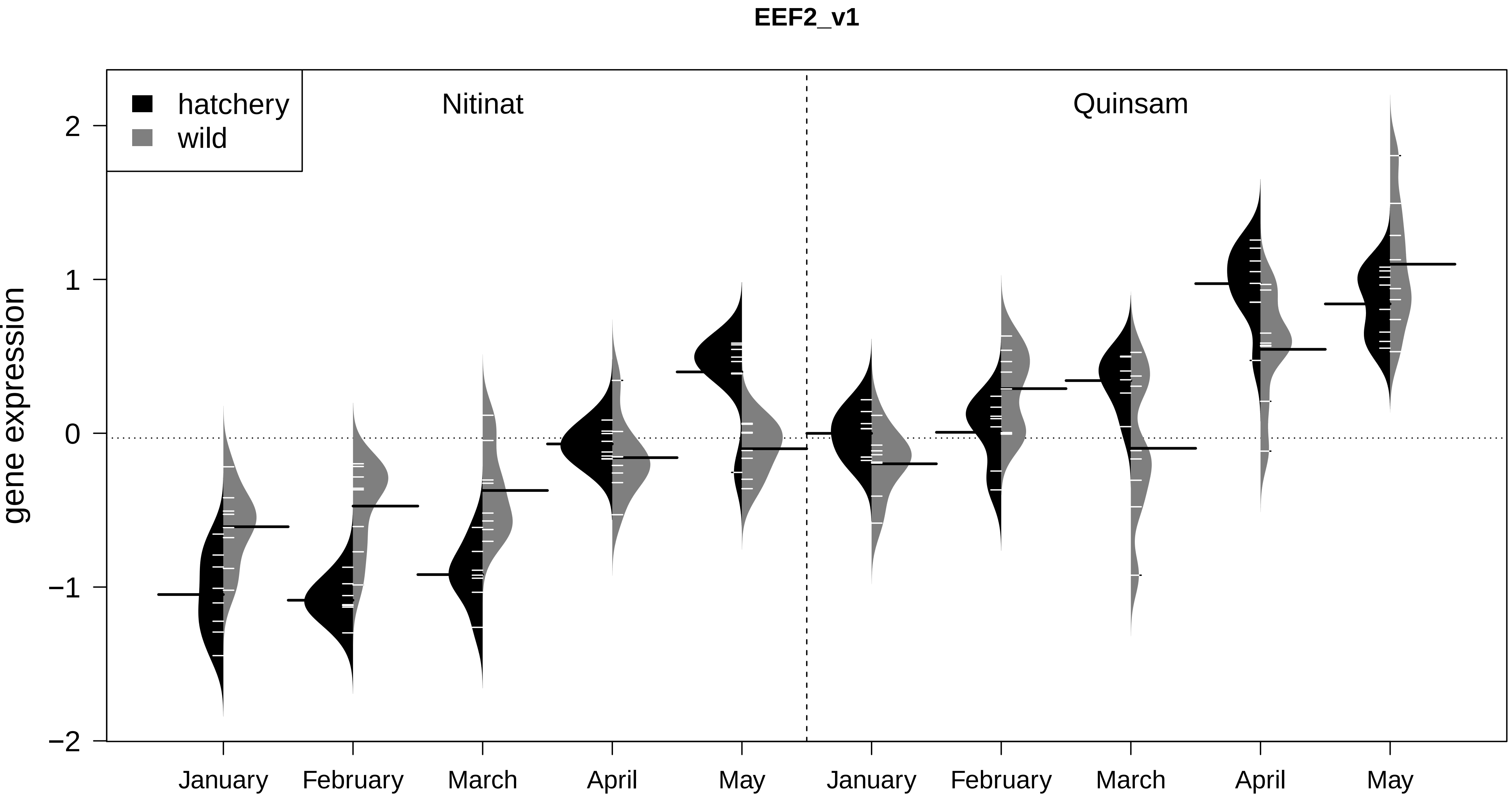

EEF2_v1
hatchery
wild
Nitinat
Quinsam
gene expression
2
1
0
−1
−2
January
February
March
April
May
January
February
March
April
May

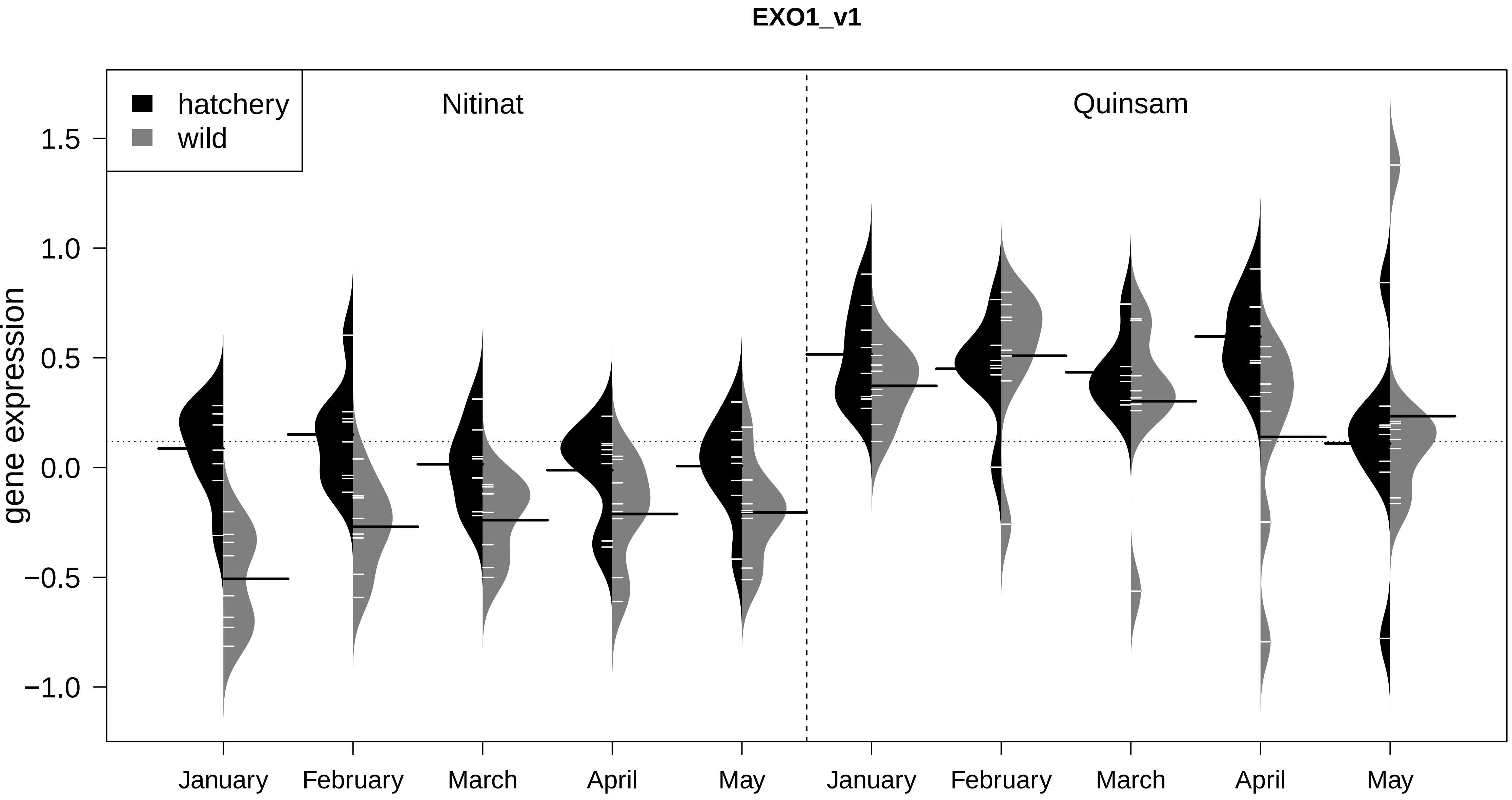

EXO1_v1
hatchery
wild
Nitinat
Quinsam
gene expression
1.5
1.0
0.5
0.0
−0.5
−1.0
January
February
March
April
May
January
February
March
April
May

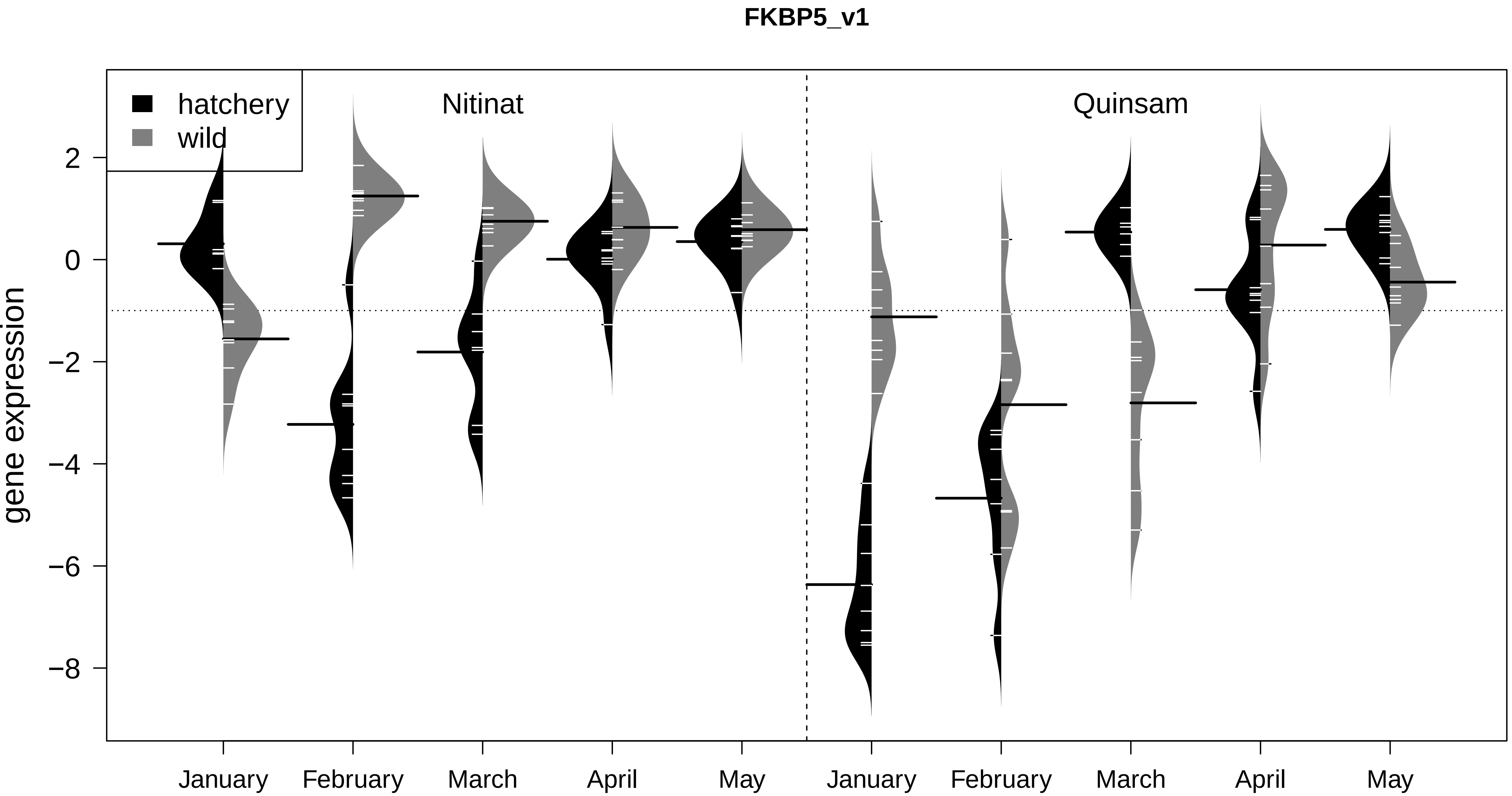

FKBP5_v1
hatchery
wild
Nitinat
Quinsam
gene expression
2
0
−2
−4
−6
−8
January
February
March
April
May
January
February
March
April
May

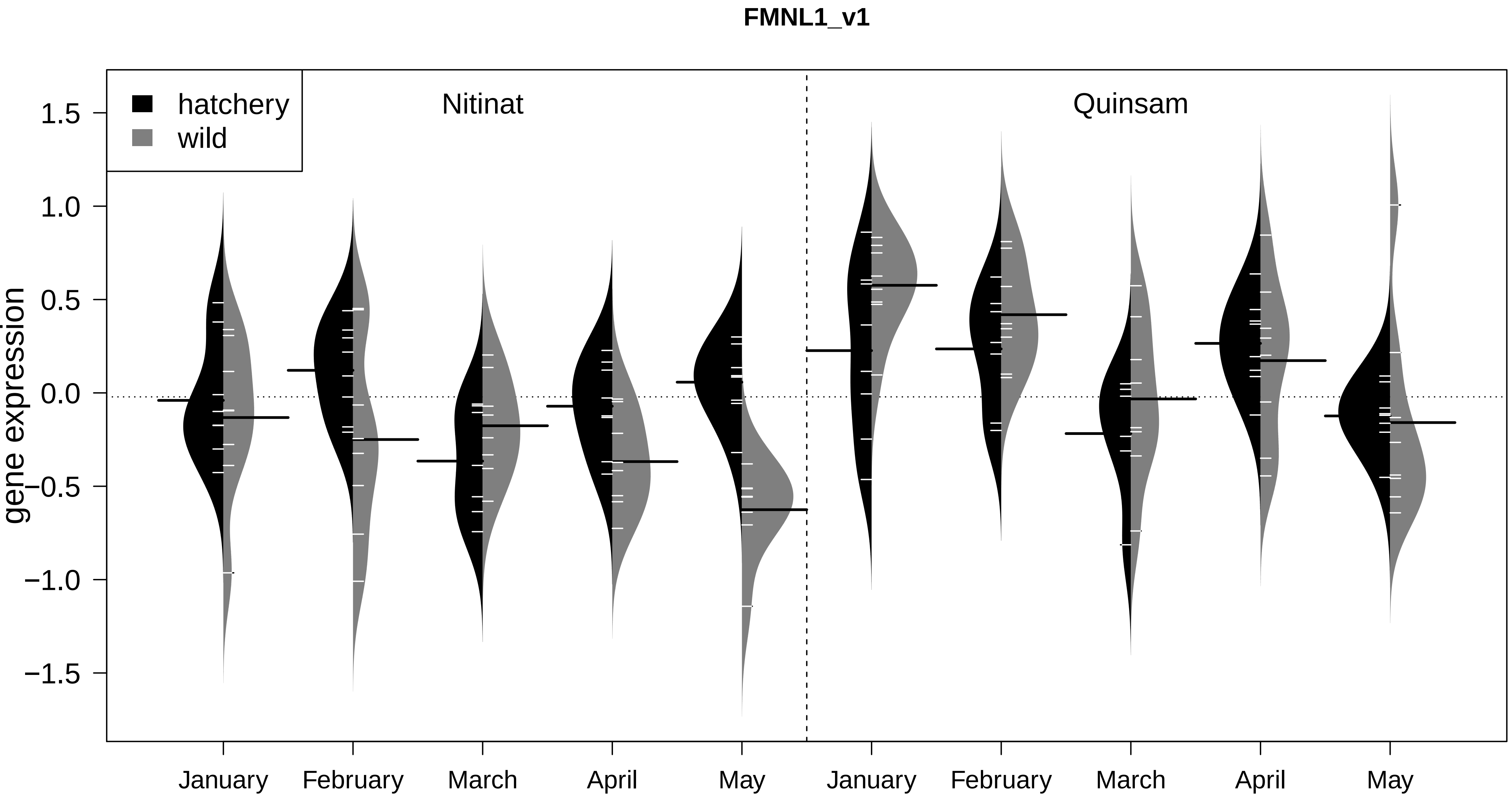

FMNL1_v1
hatchery
wild
Nitinat
Quinsam
gene expression
1.5
1.0
0.5
0.0
−0.5
−1.0
−1.5
January
February
March
April
May
January
February
March
April
May

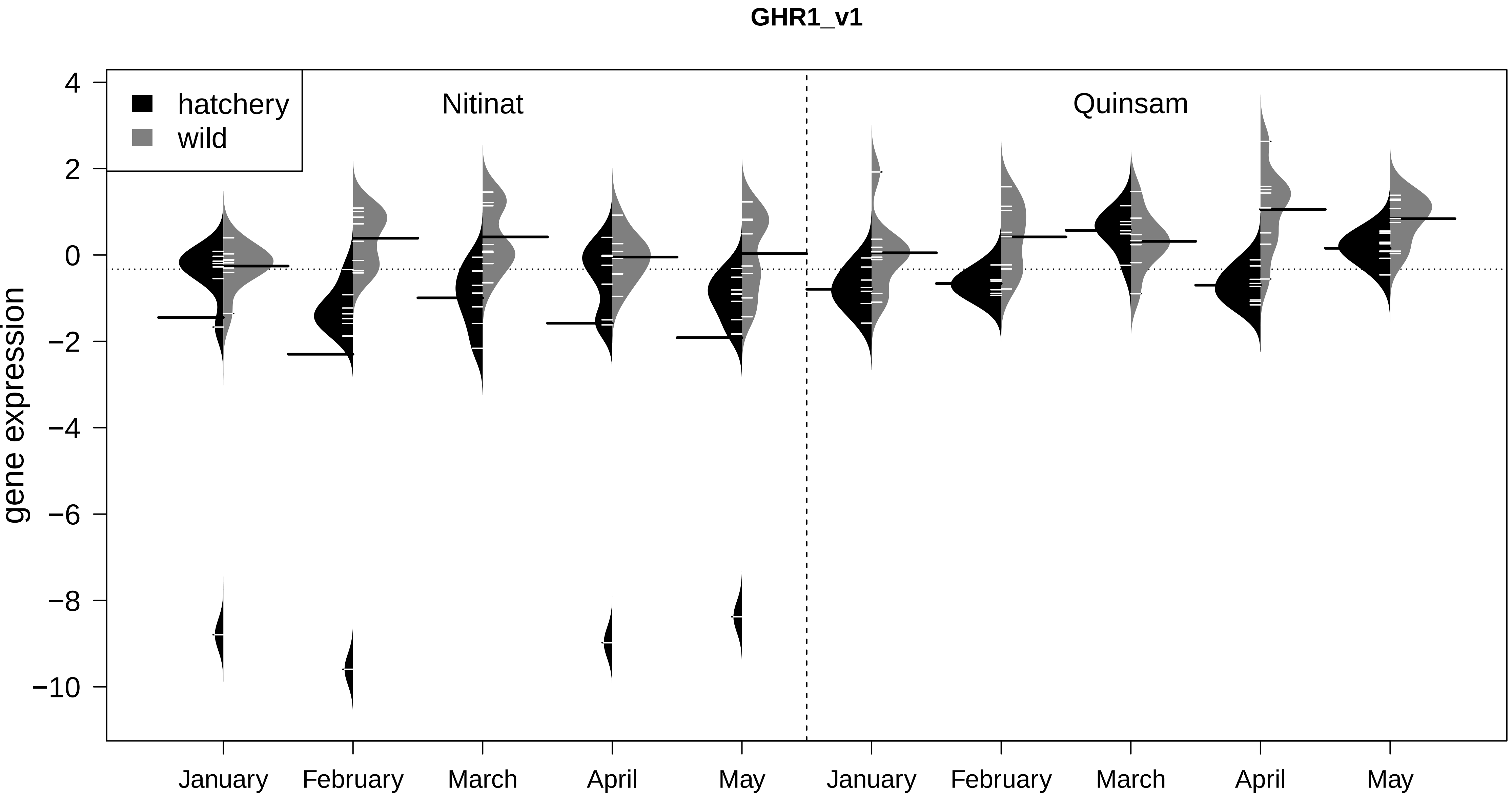

GHR1_v1
hatchery
wild
Nitinat
Quinsam
gene expression
4
2
0
−2
−4
−6
−8
−10
January
February
March
April
May
January
February
March
April
May

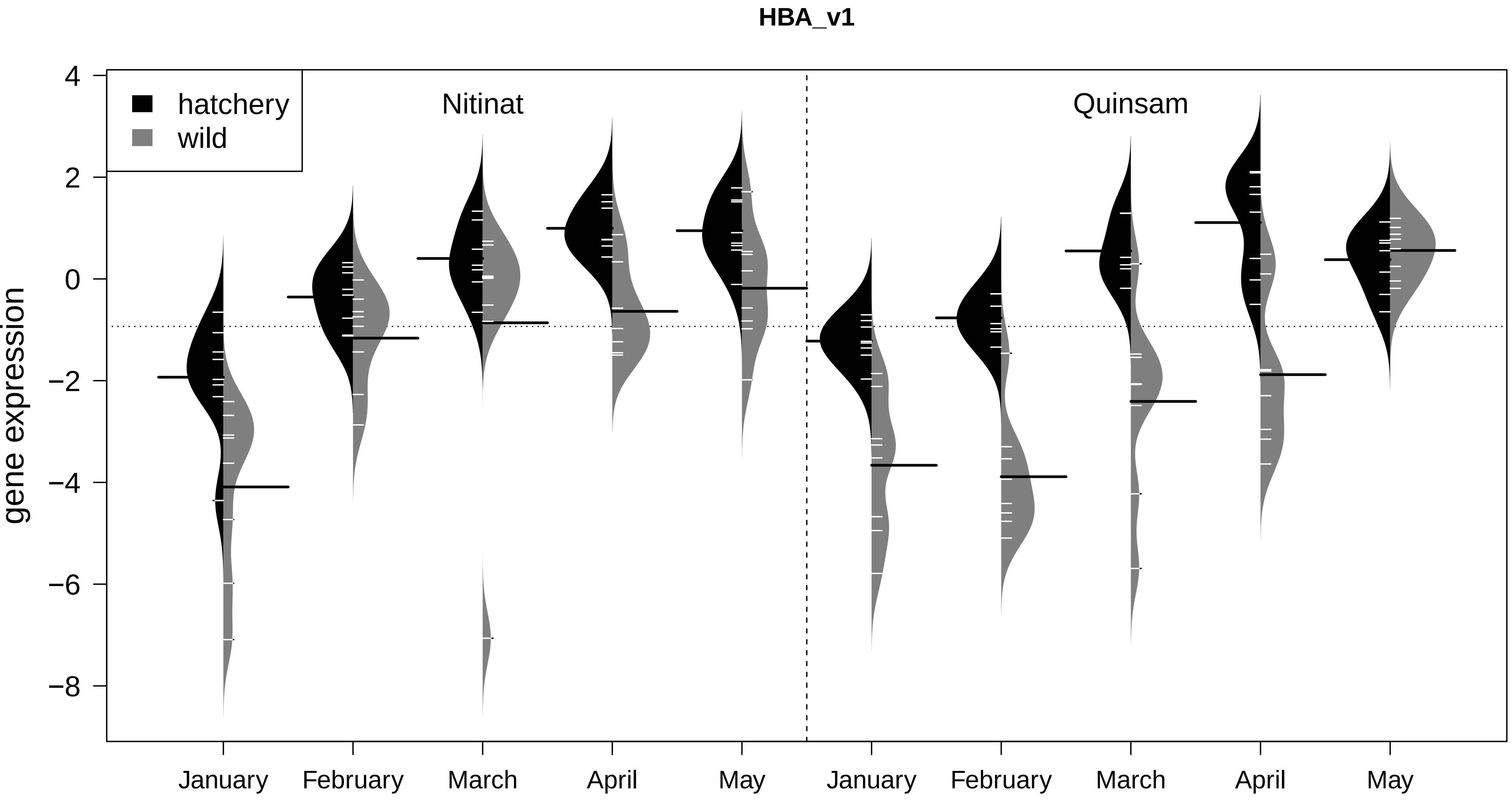
HBA_v1
hatchery
wild
Nitinat
Quinsam
gene expression
4
2
0
−2
−4
−6
−8
January
February
March
April
May
January
February
March
April
May

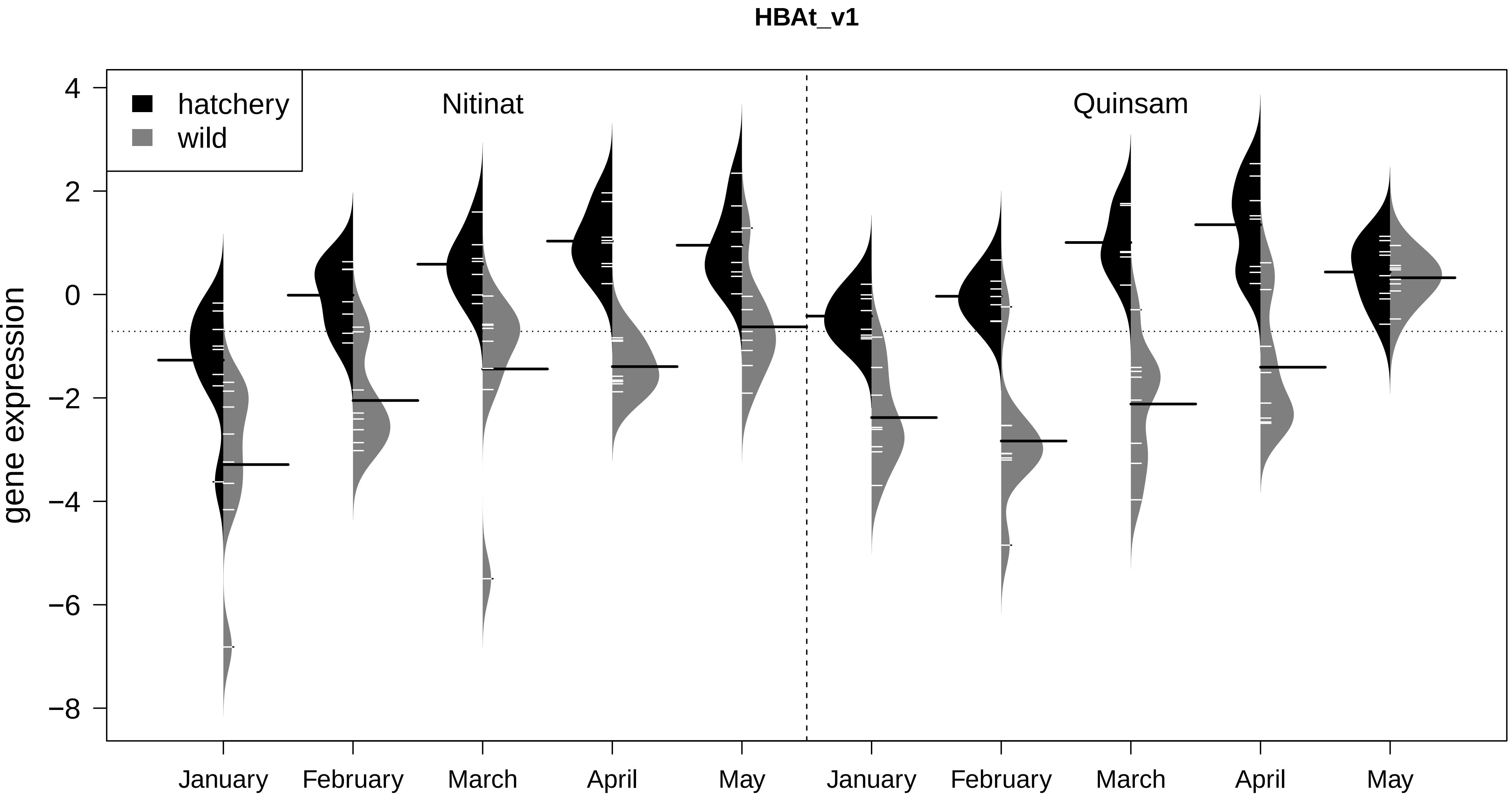
HBAt_v1
hatchery
wild
Nitinat
Quinsam
gene expression
4
2
0
−2
−4
−6
−8
January
February
March
April
May
January
February
March
April
May

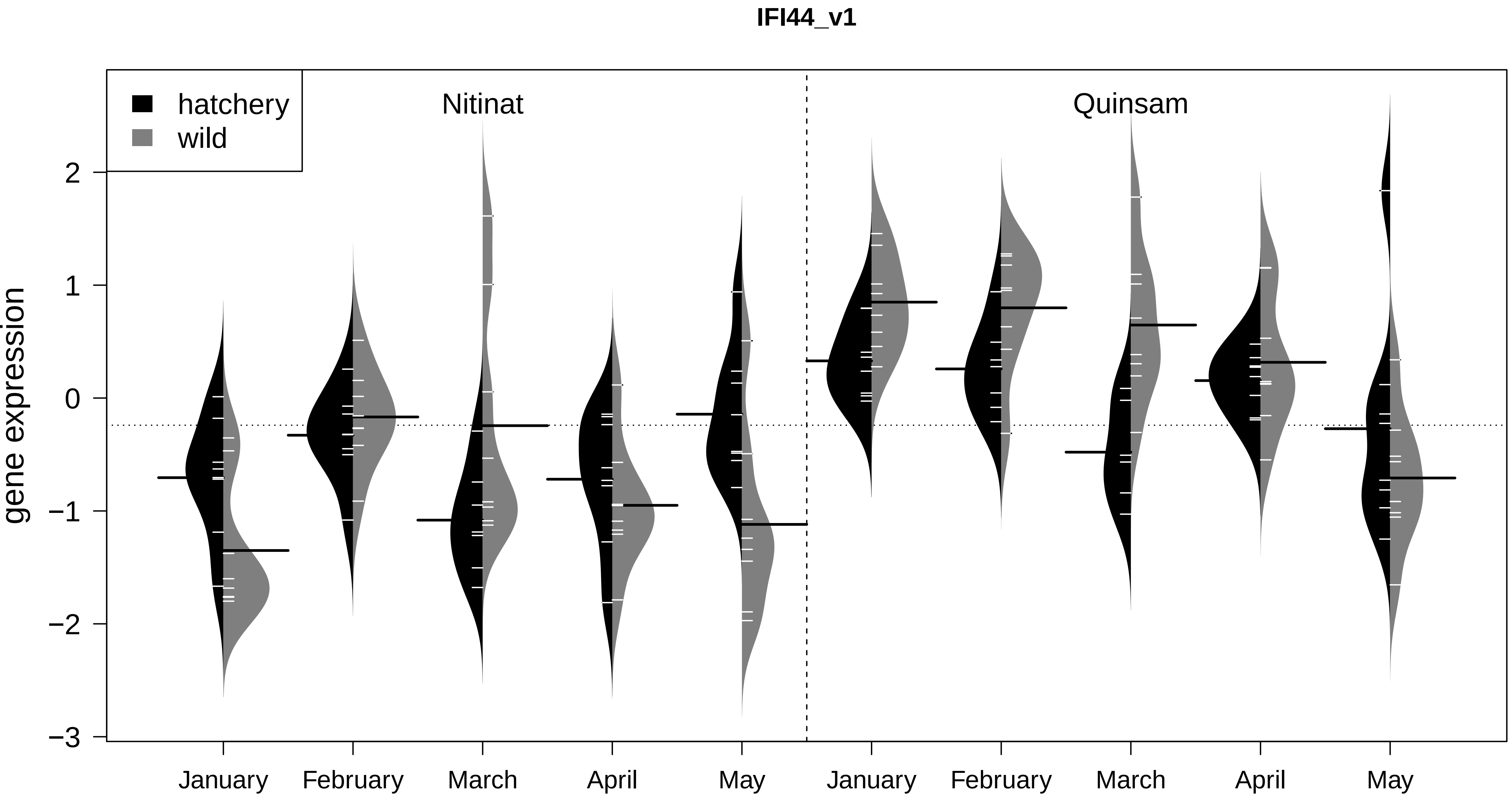

IFI44_v1
hatchery
wild
Nitinat
Quinsam
gene expression
2
1
0
−1
−2
−3
January
February
March
April
May
January
February
March
April
May

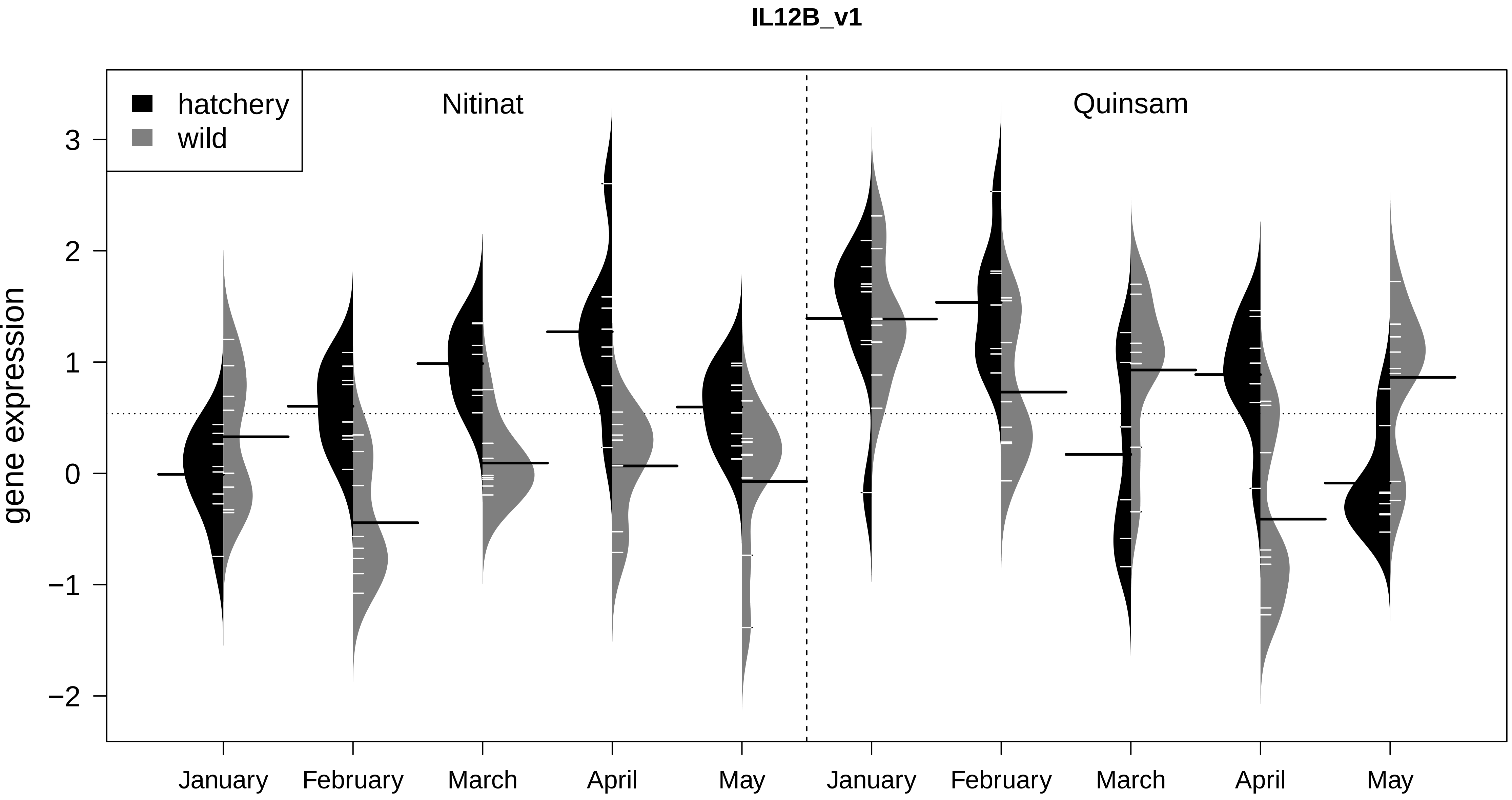
IL12B_v1
hatchery
wild
Nitinat
Quinsam
gene expression
3
2
1
0
−1
−2
January
February
March
April
May
January
February
March
April
May

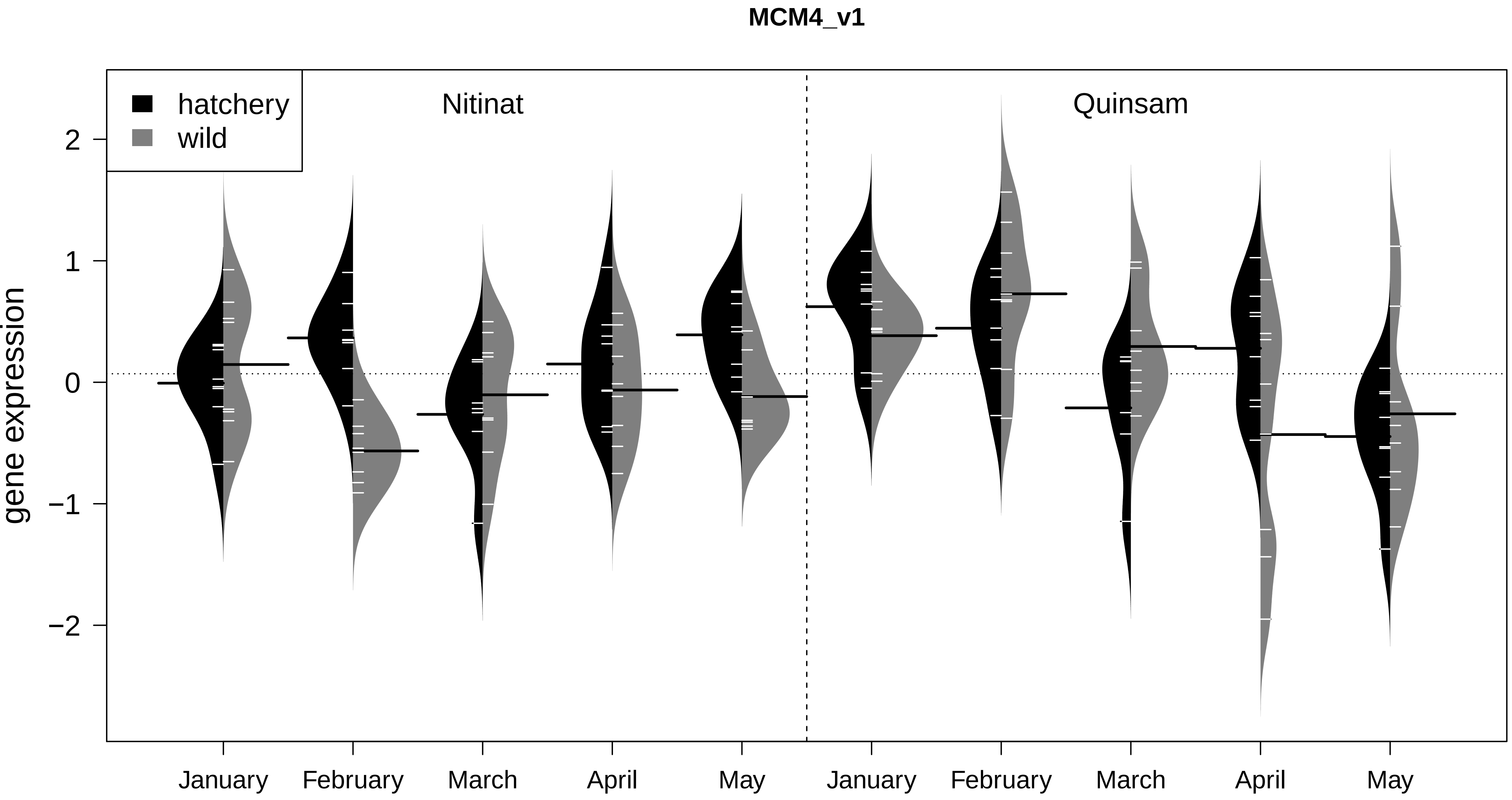

MCM4_v1
hatchery
wild
Nitinat
Quinsam
gene expression
2
1
0
−1
−2
January
February
March
April
May
January
February
March
April
May

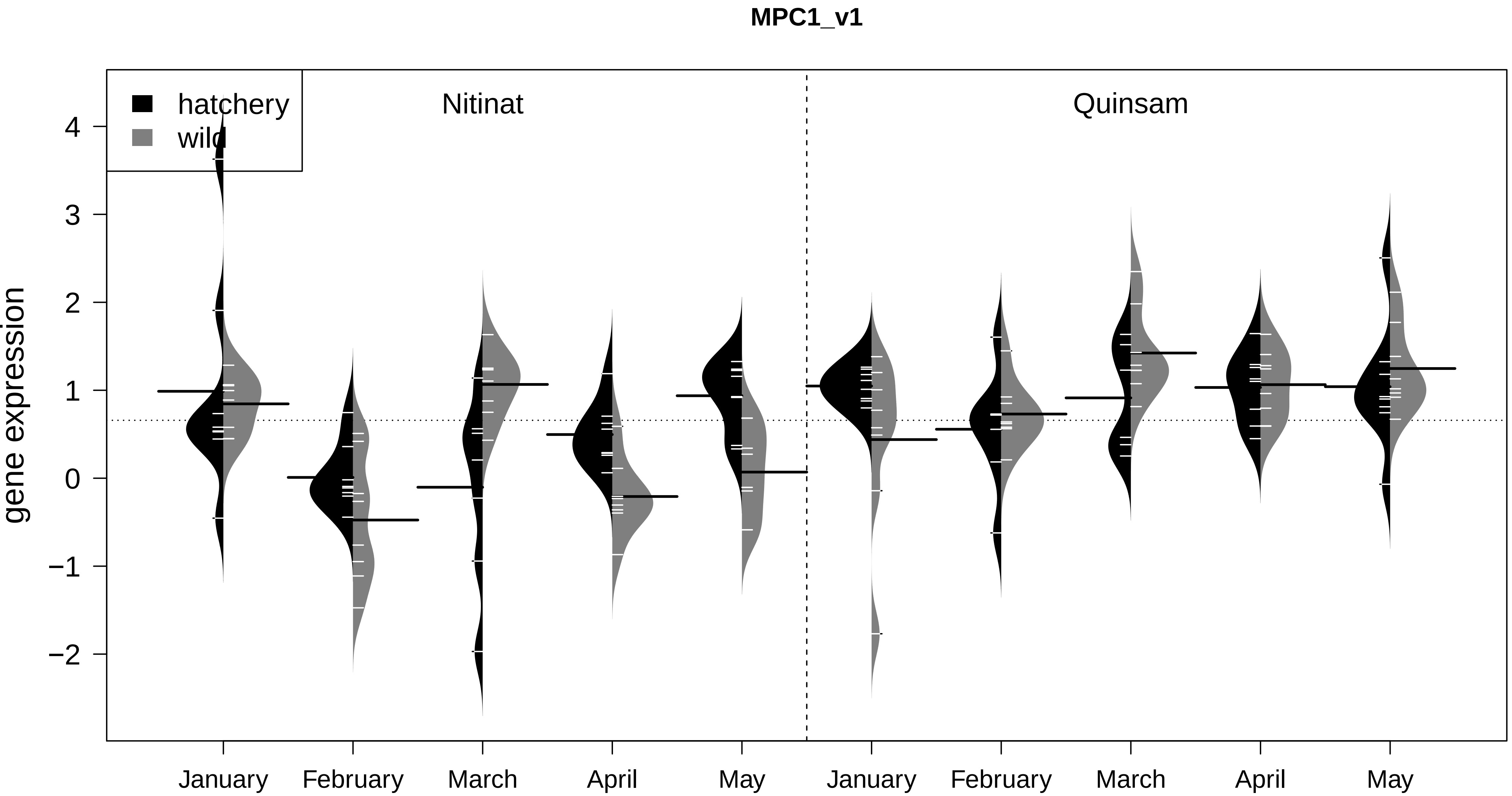

MPC1_v1
hatchery
wild
Nitinat
Quinsam
gene expression
4
3
2
1
0
−1
−2
January
February
March
April
May
January
February
March
April
May

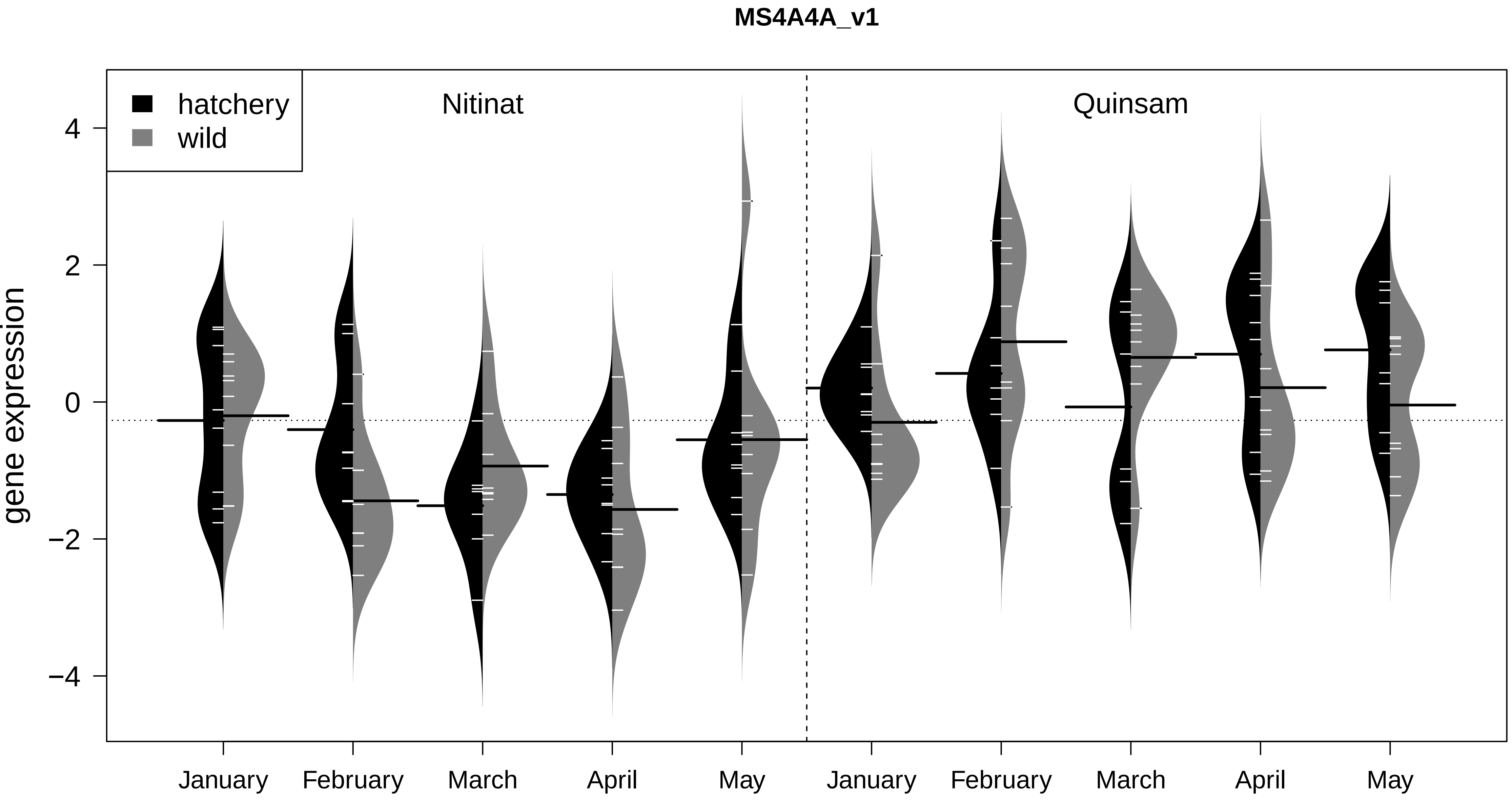

MS4A4A_v1
hatchery
wild
Nitinat
Quinsam
gene expression
4
2
0
−2
−4
January
February
March
April
May
January
February
March
April
May

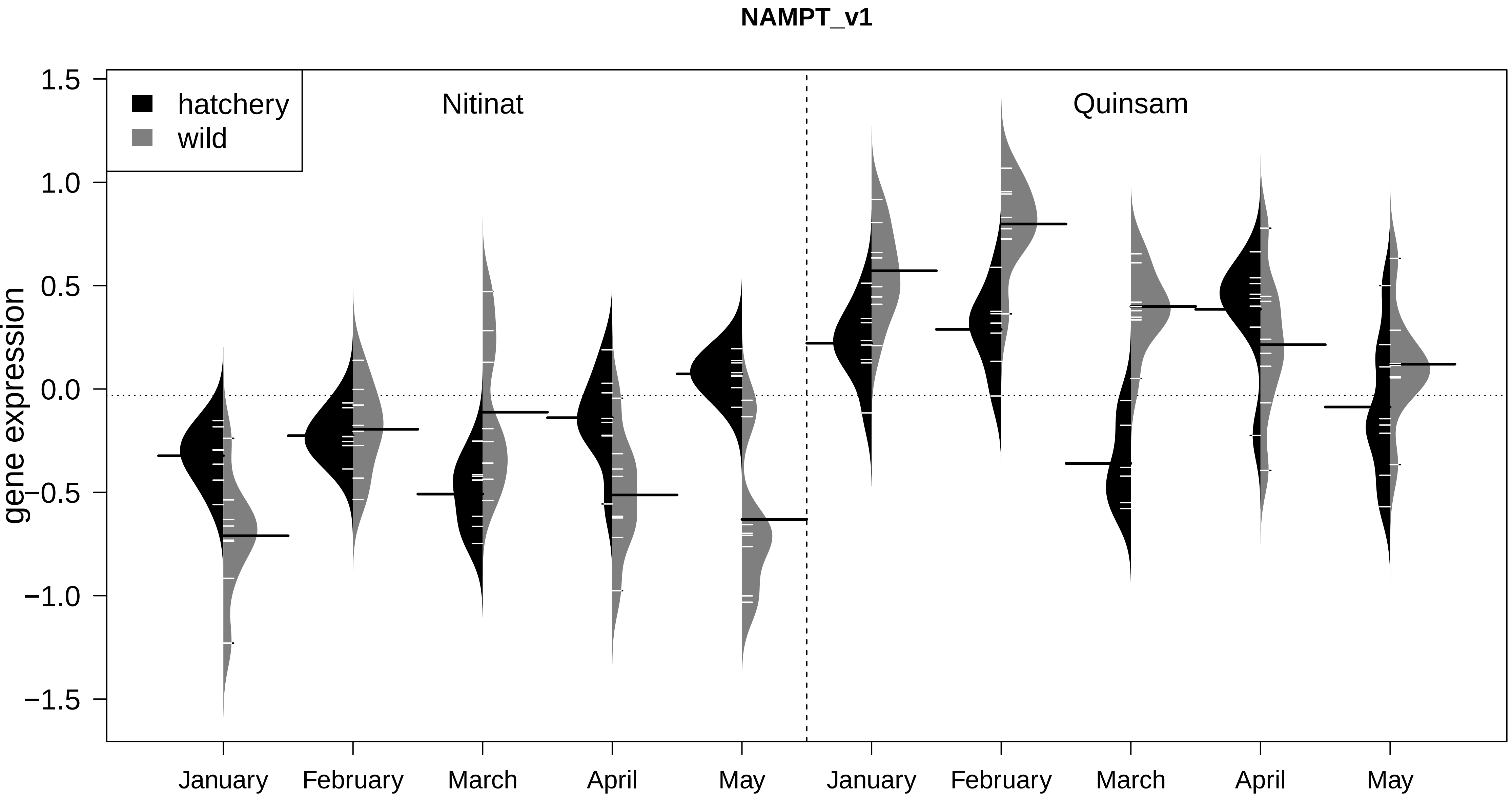

NAMPT_v1
hatchery
wild
Nitinat
Quinsam
gene expression
1.5
1.0
0.5
0.0
−0.5
−1.0
−1.5
January
February
March
April
May
January
February
March
April
May

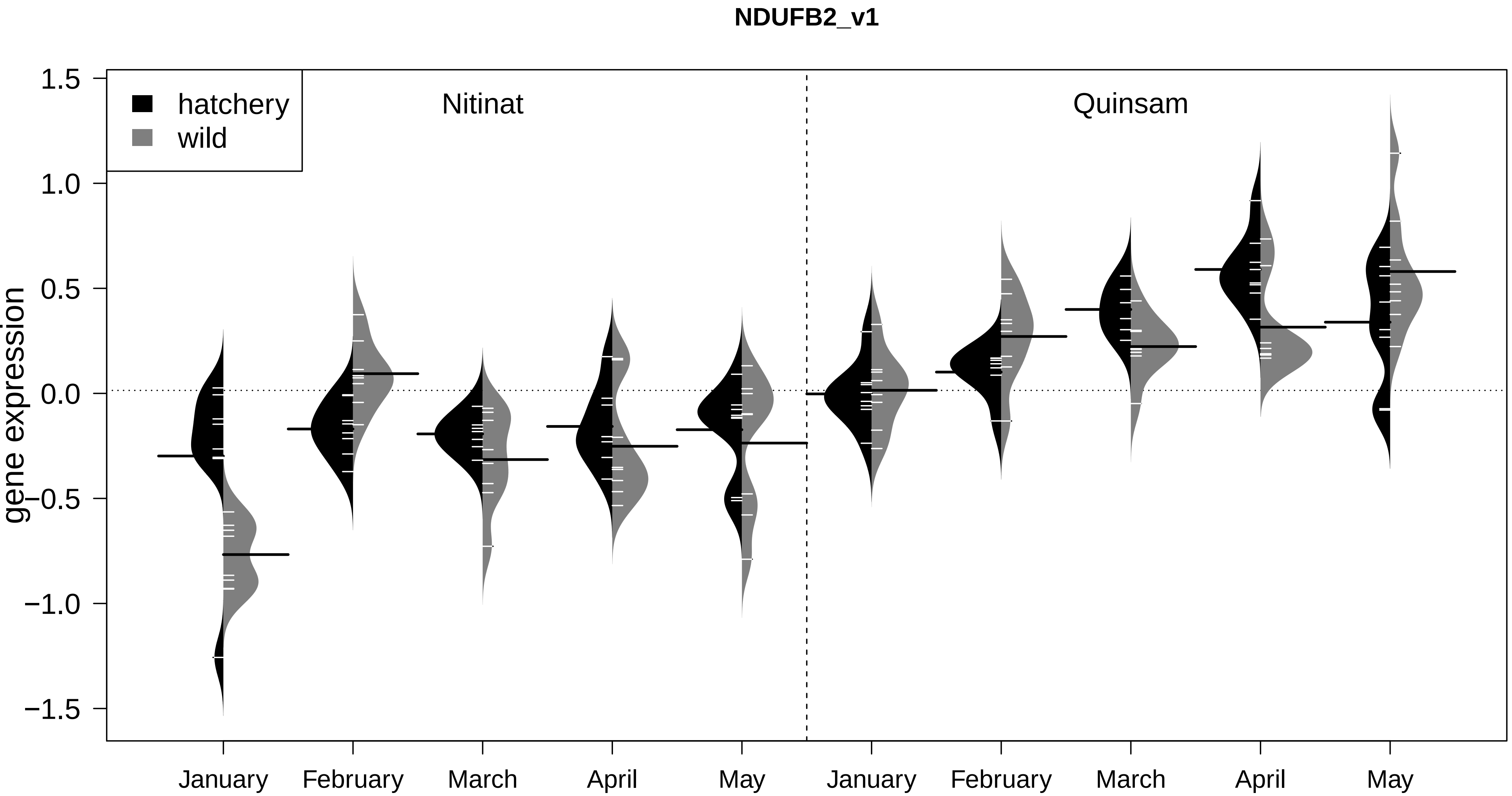
NDUFB2_v1
hatchery
wild
Nitinat
Quinsam
gene expression
1.5
1.0
0.5
0.0
−0.5
−1.0
−1.5
January
February
March
April
May
January
February
March
April
May

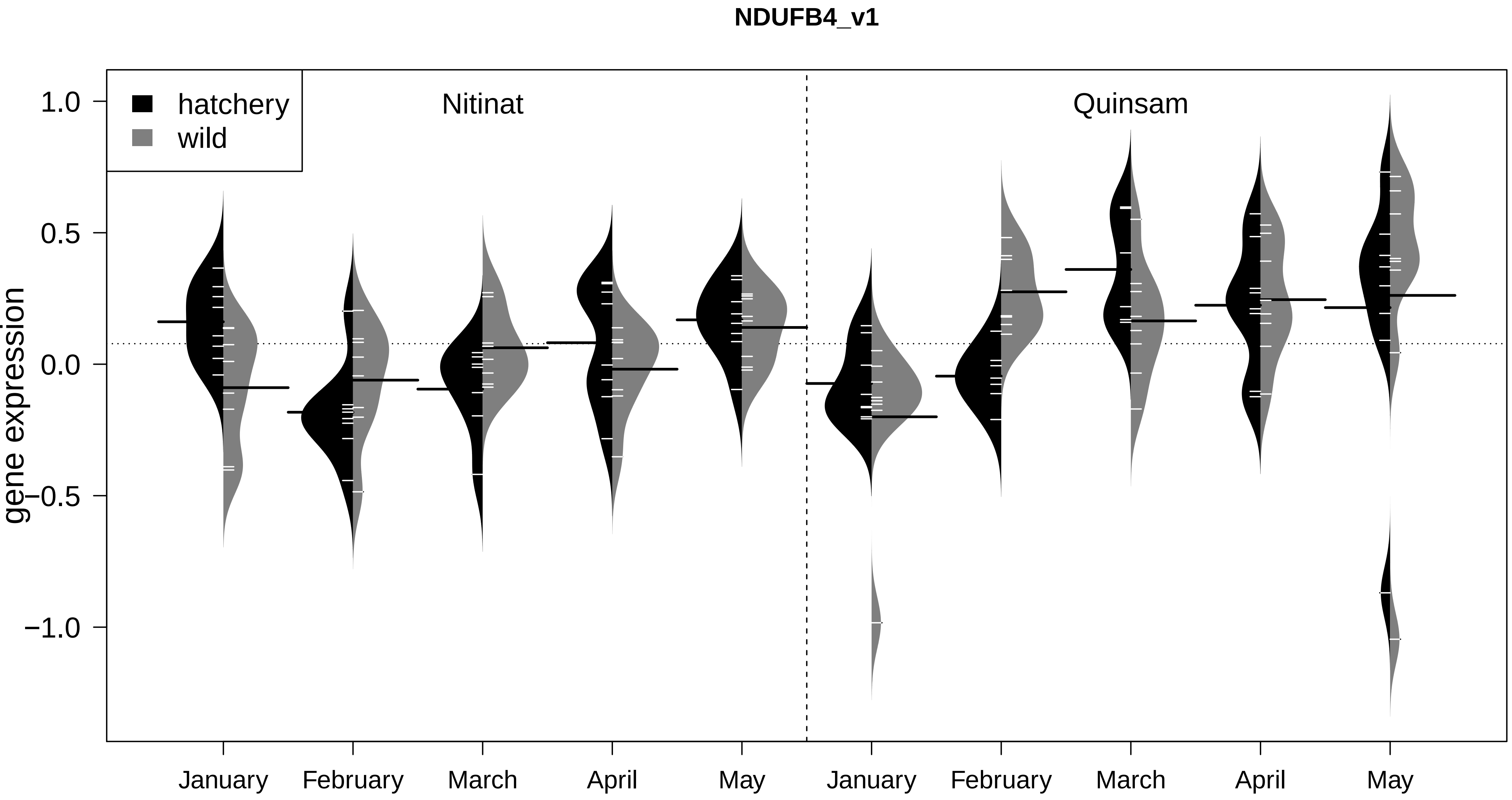
NDUFB4_v1
hatchery
wild
Nitinat
Quinsam
gene expression
1.0
0.5
0.0
−0.5
−1.0
January
February
March
April
May
January
February
March
April
May

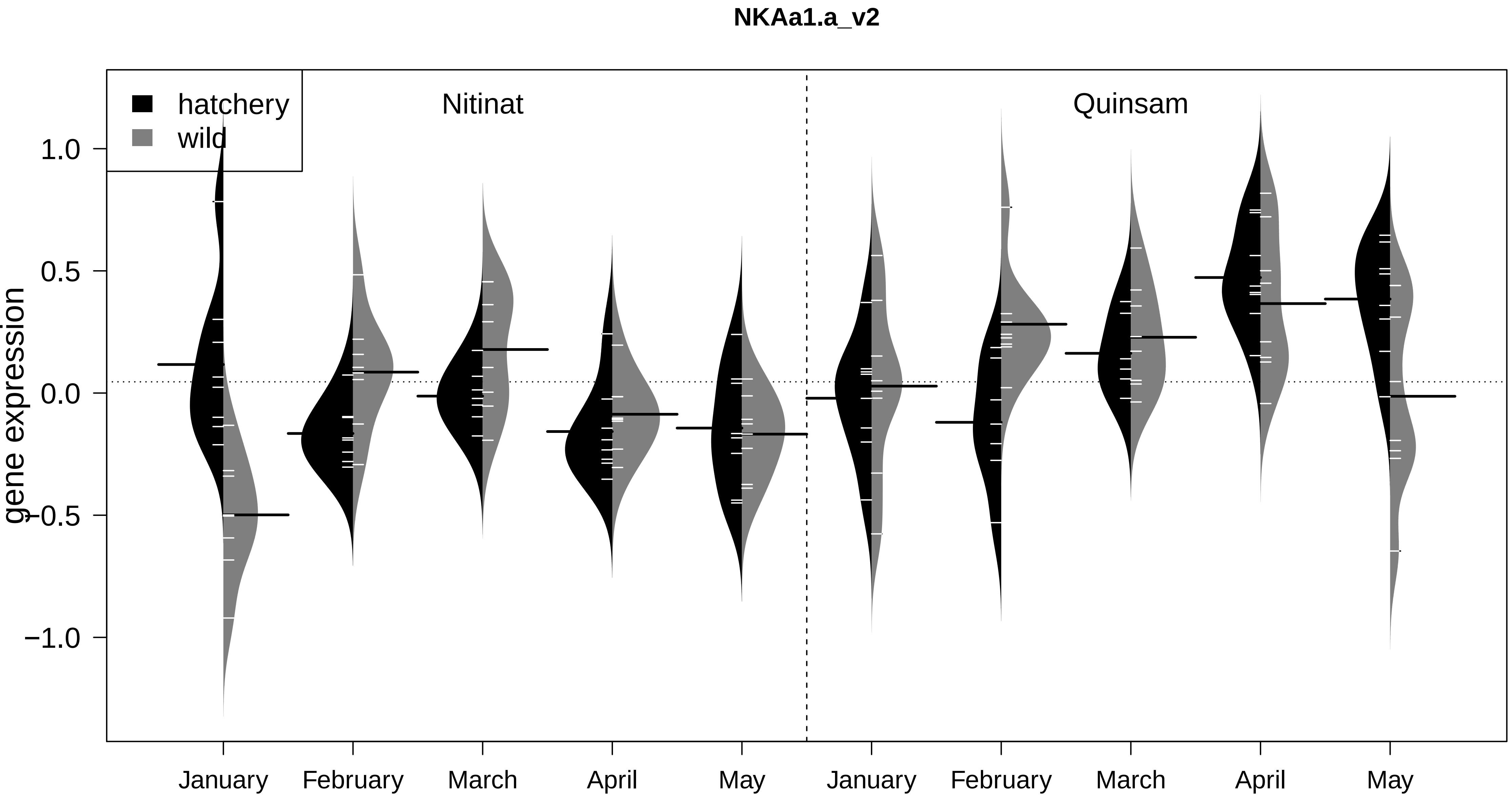
NKAa1.a_v2
hatchery
wild
Nitinat
Quinsam
gene expression
1.0
0.5
0.0
−0.5
−1.0
January
February
March
April
May
January
February
March
April
May

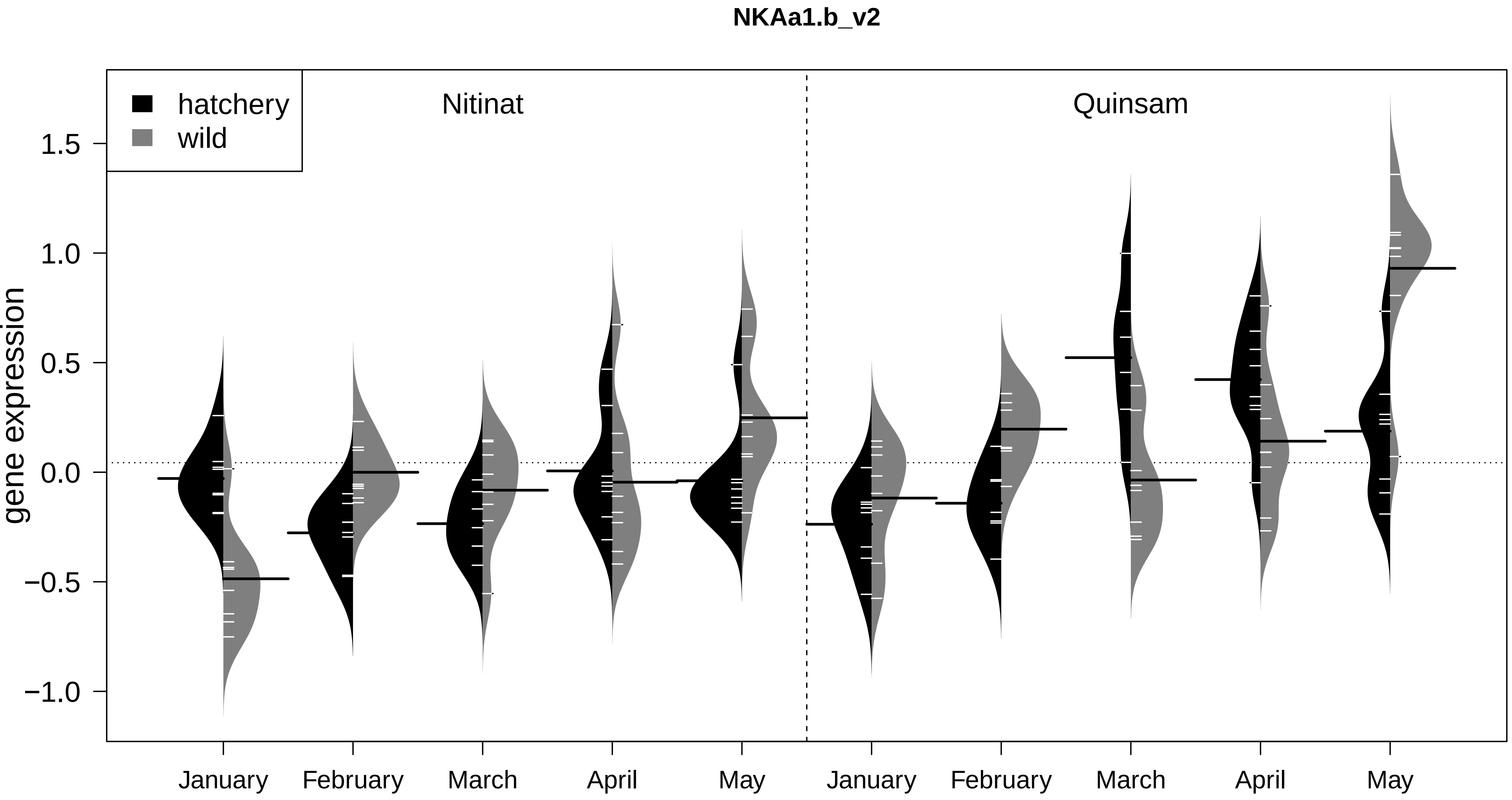

NKAa1.b_v2
hatchery
wild
Nitinat
Quinsam
gene expression
1.5
1.0
0.5
0.0
−0.5
−1.0
January
February
March
April
May
January
February
March
April
May

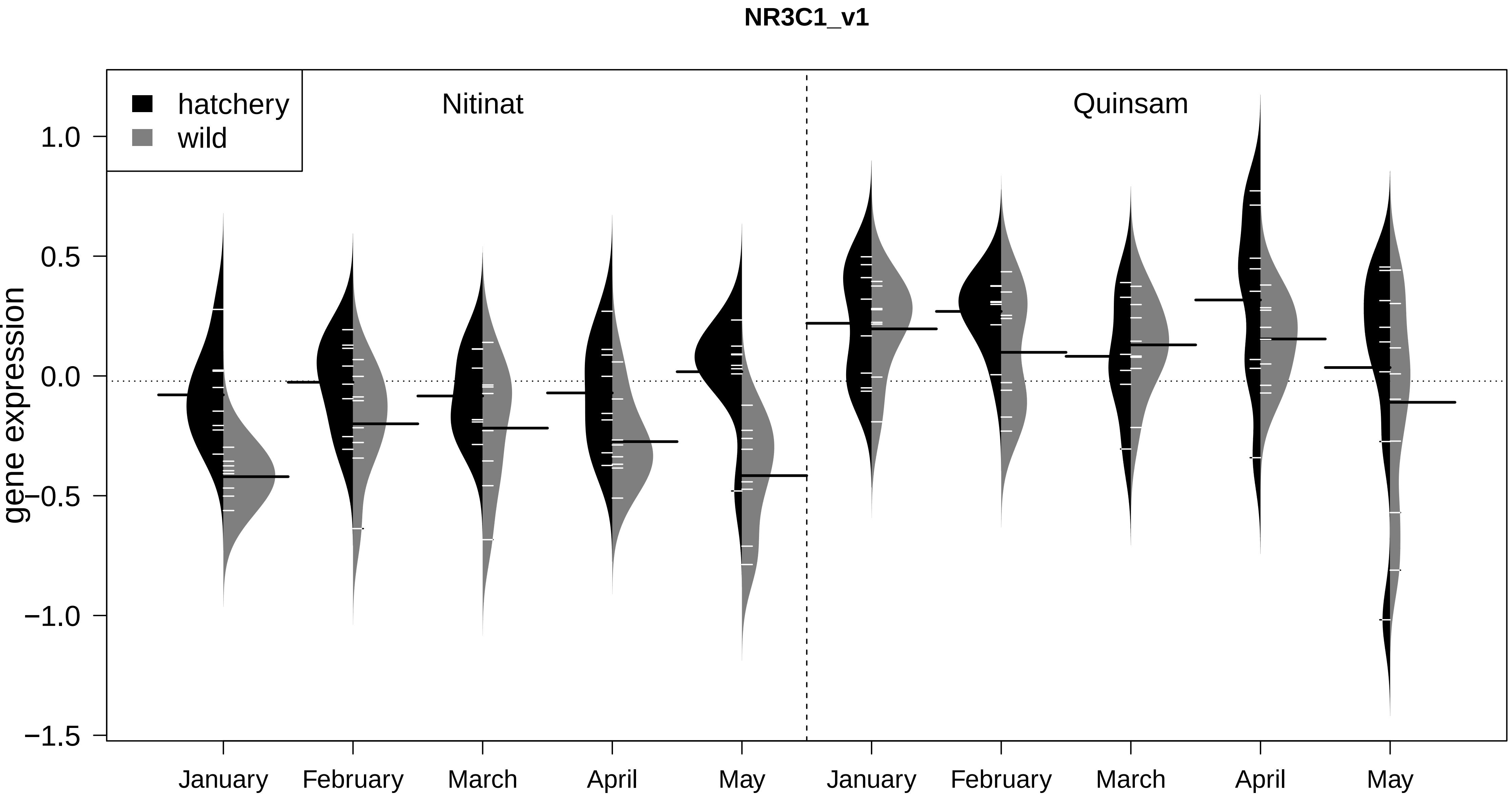

NR3C1_v1
hatchery
wild
Nitinat
Quinsam
gene expression
1.0
0.5
0.0
−0.5
−1.0
−1.5
January
February
March
April
May
January
February
March
April
May

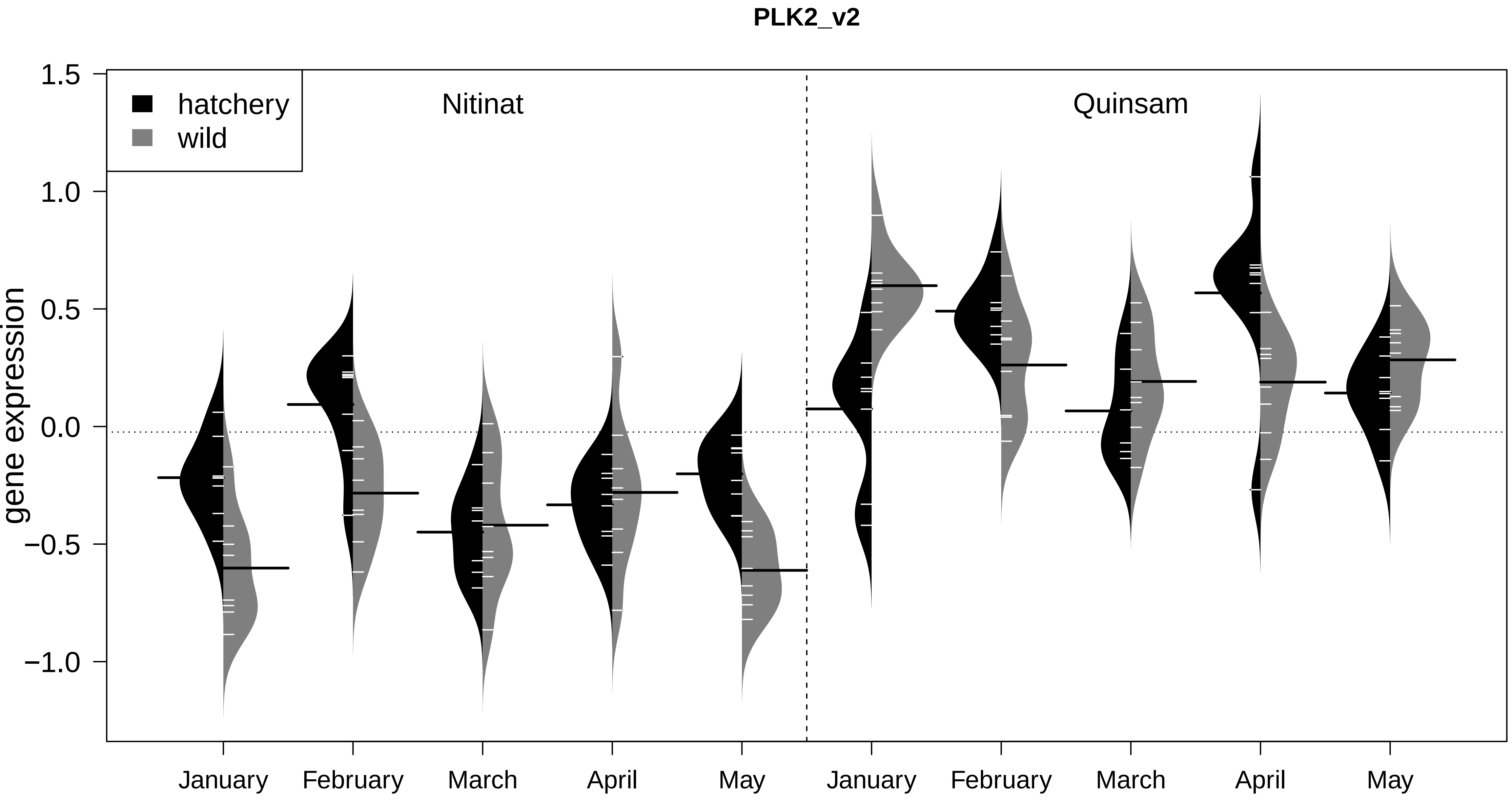
PLK2_v2
hatchery
wild
Nitinat
Quinsam
gene expression
1.5
1.0
0.5
0.0
−0.5
−1.0
January
February
March
April
May
January
February
March
April
May

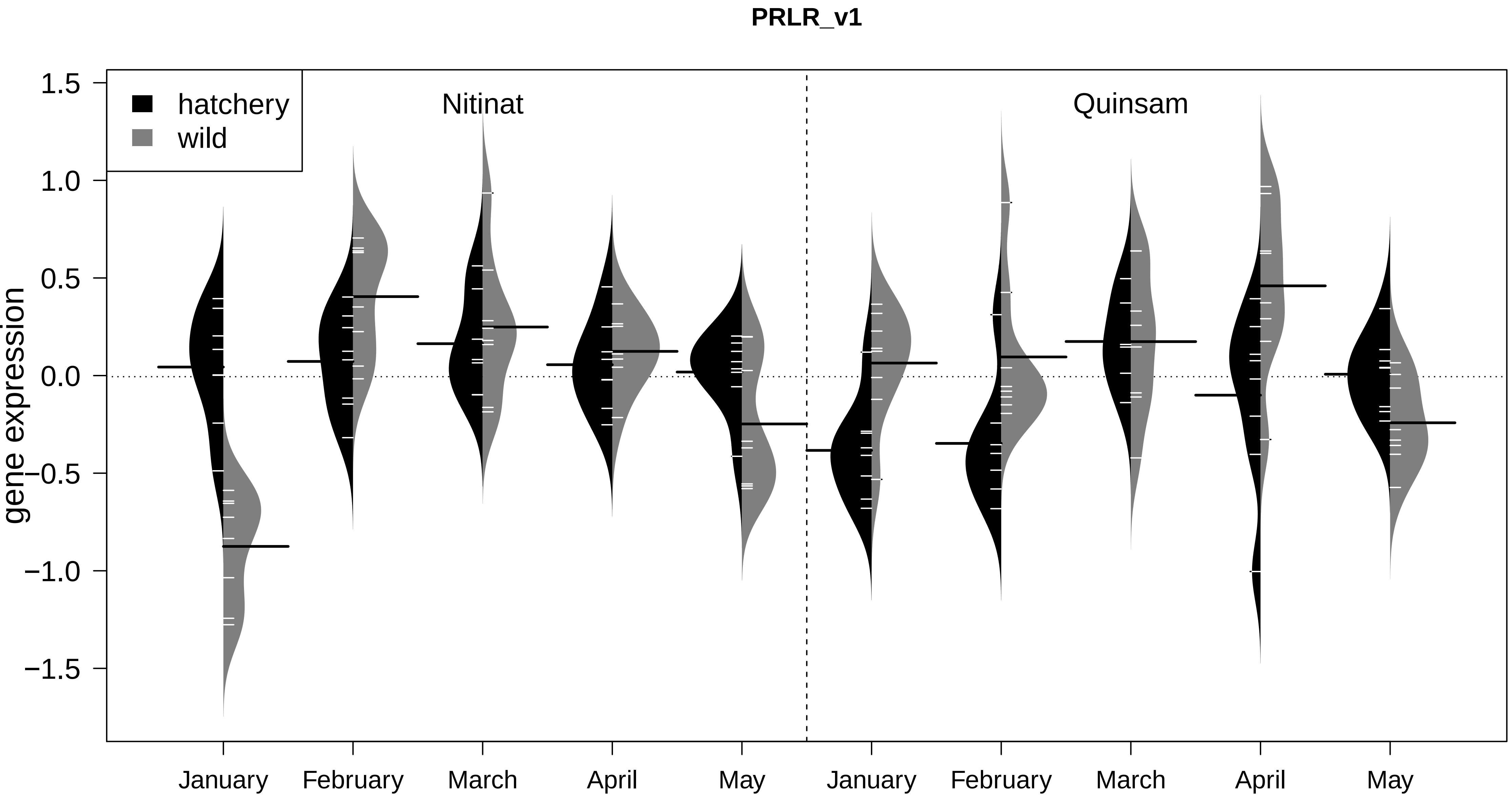
PRLR_v1
hatchery
wild
Nitinat
Quinsam
gene expression
1.5
1.0
0.5
0.0
−0.5
−1.0
−1.5
January
February
March
April
May
January
February
March
April
May

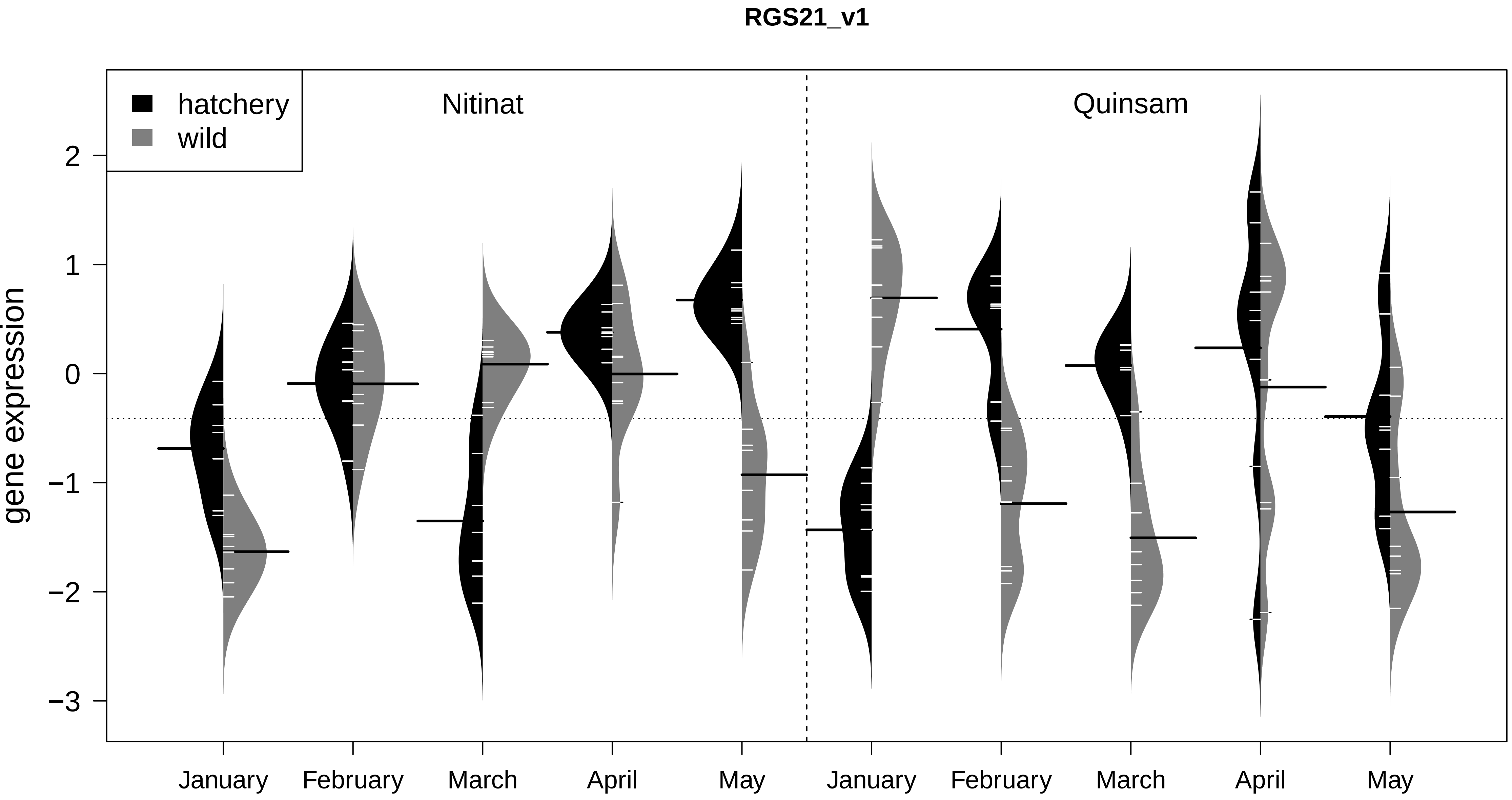
RGS21_v1
hatchery
wild
Nitinat
Quinsam
gene expression
2
1
0
−1
−2
−3
January
February
March
April
May
January
February
March
April
May

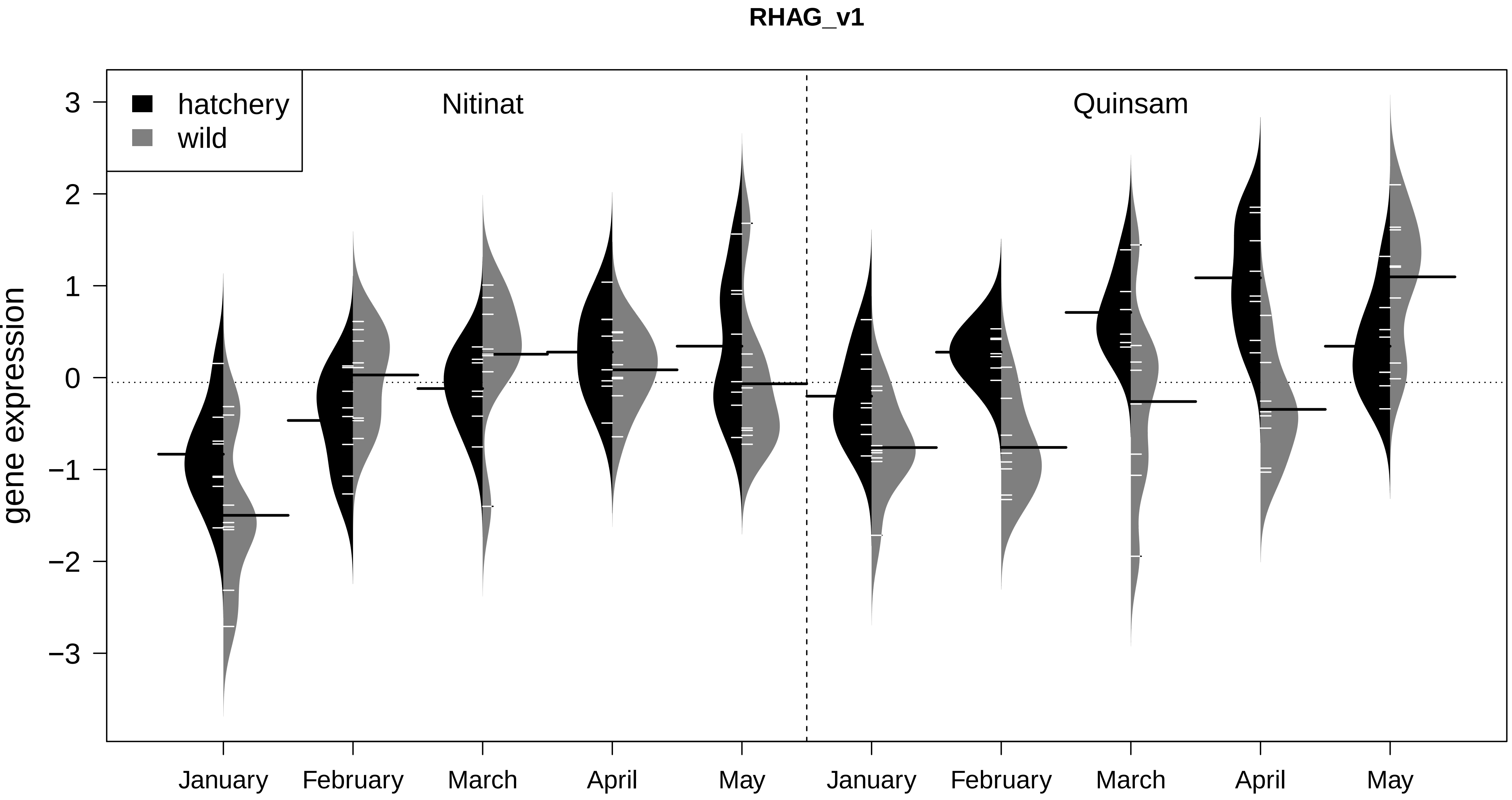

RHAG_v1
hatchery
wild
Nitinat
Quinsam
gene expression
3
2
1
0
−1
−2
−3
January
February
March
April
May
January
February
March
April
May

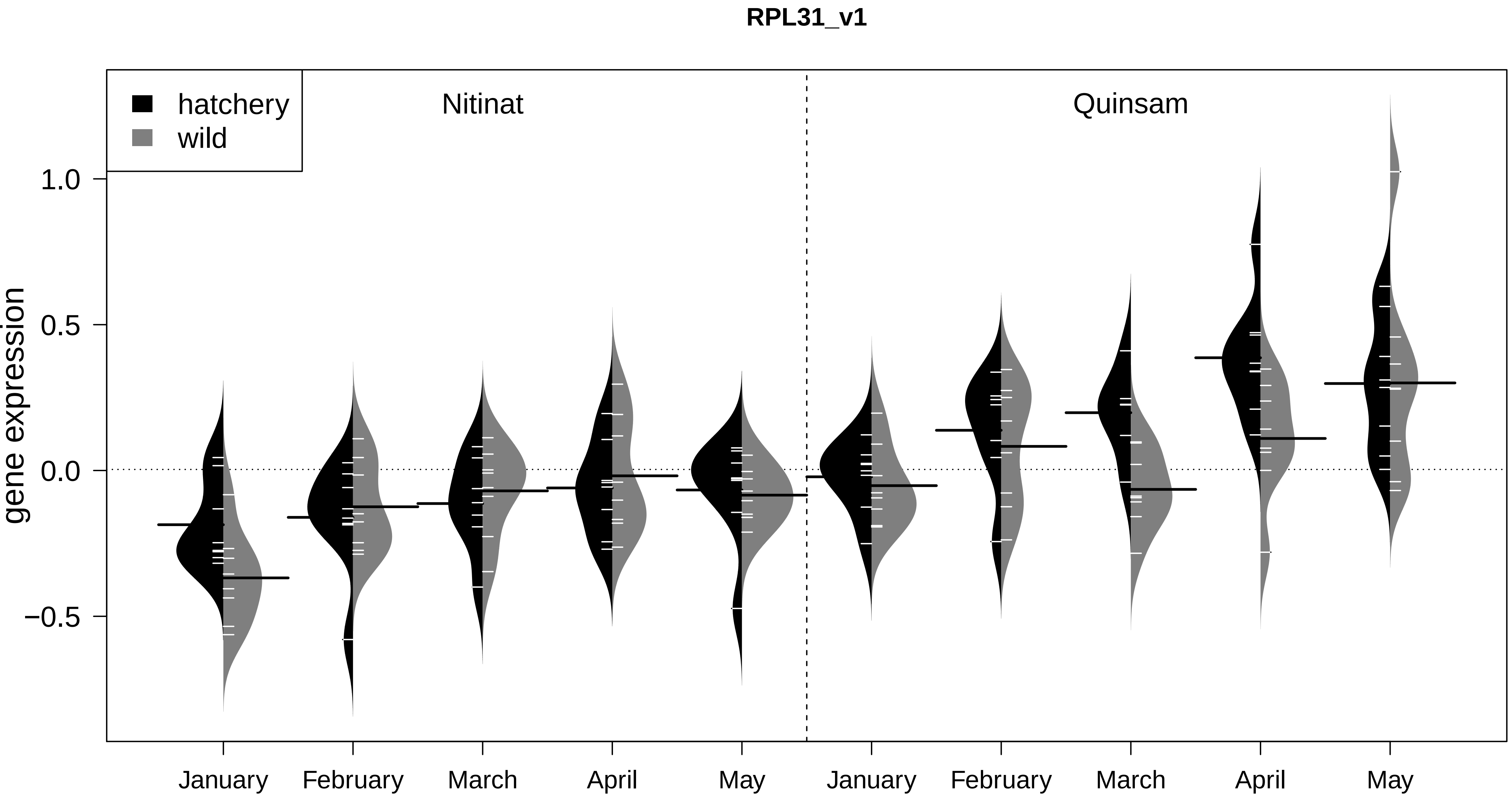

RPL31_v1
hatchery
wild
Nitinat
Quinsam
gene expression
1.0
0.5
0.0
−0.5
January
February
March
April
May
January
February
March
April
May

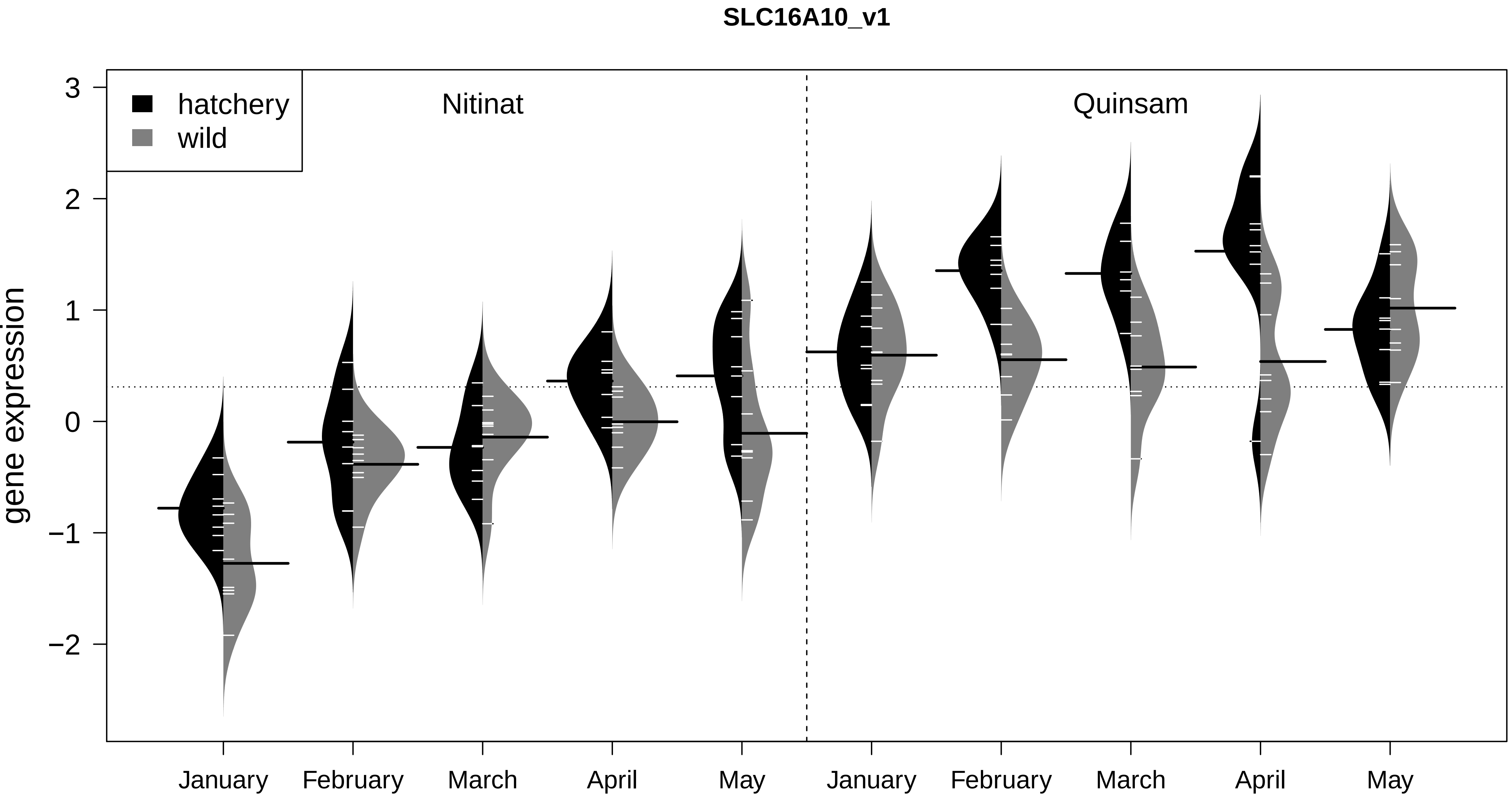

SLC16A10_v1
hatchery
wild
Nitinat
Quinsam
gene expression
3
2
1
0
−1
−2
January
February
March
April
May
January
February
March
April
May

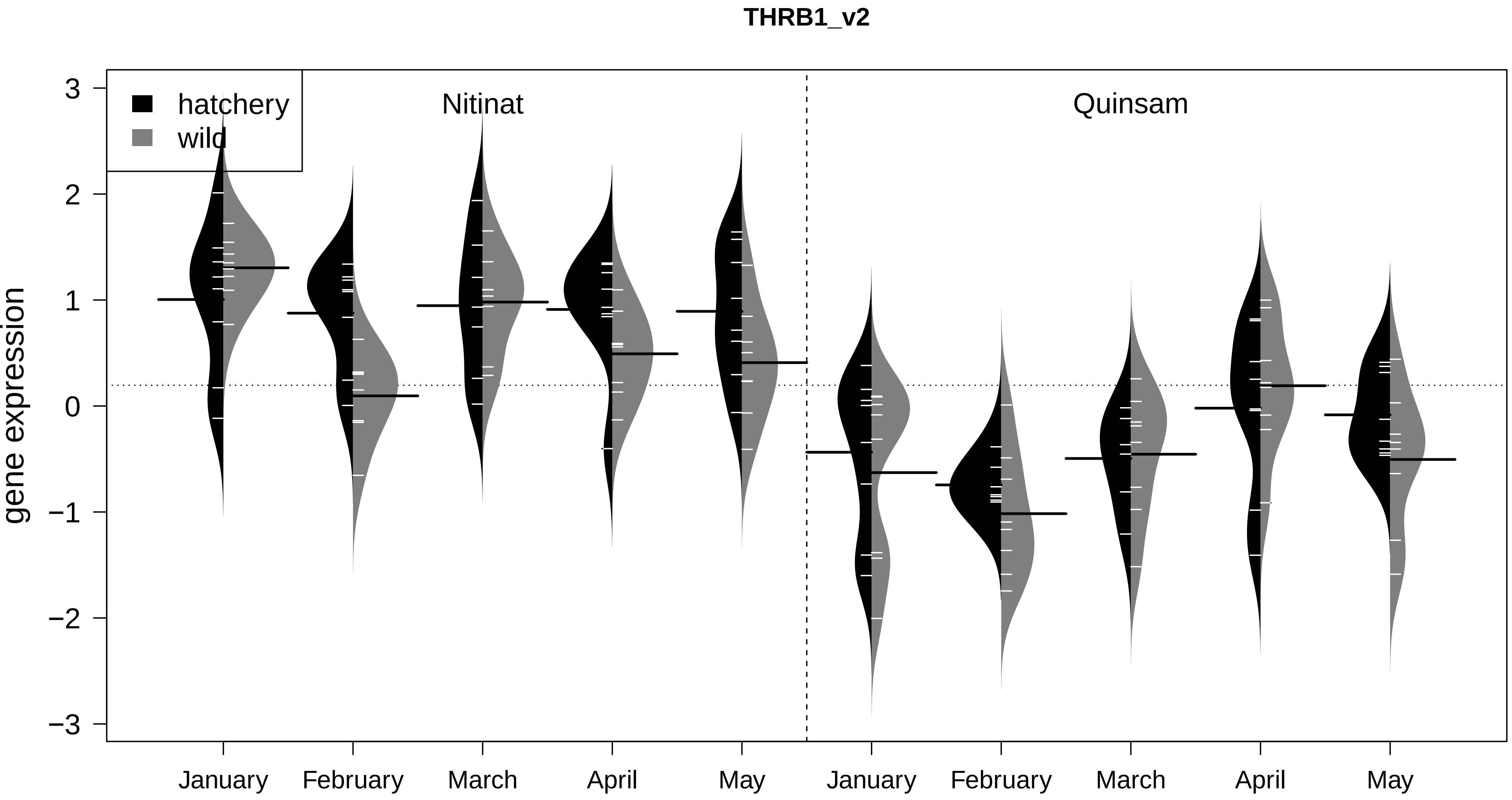
THRB1_v2
hatchery
wild
Nitinat
Quinsam
gene expression
3
2
1
0
−1
−2
−3
January
February
March
April
May
January
February
March
April
May

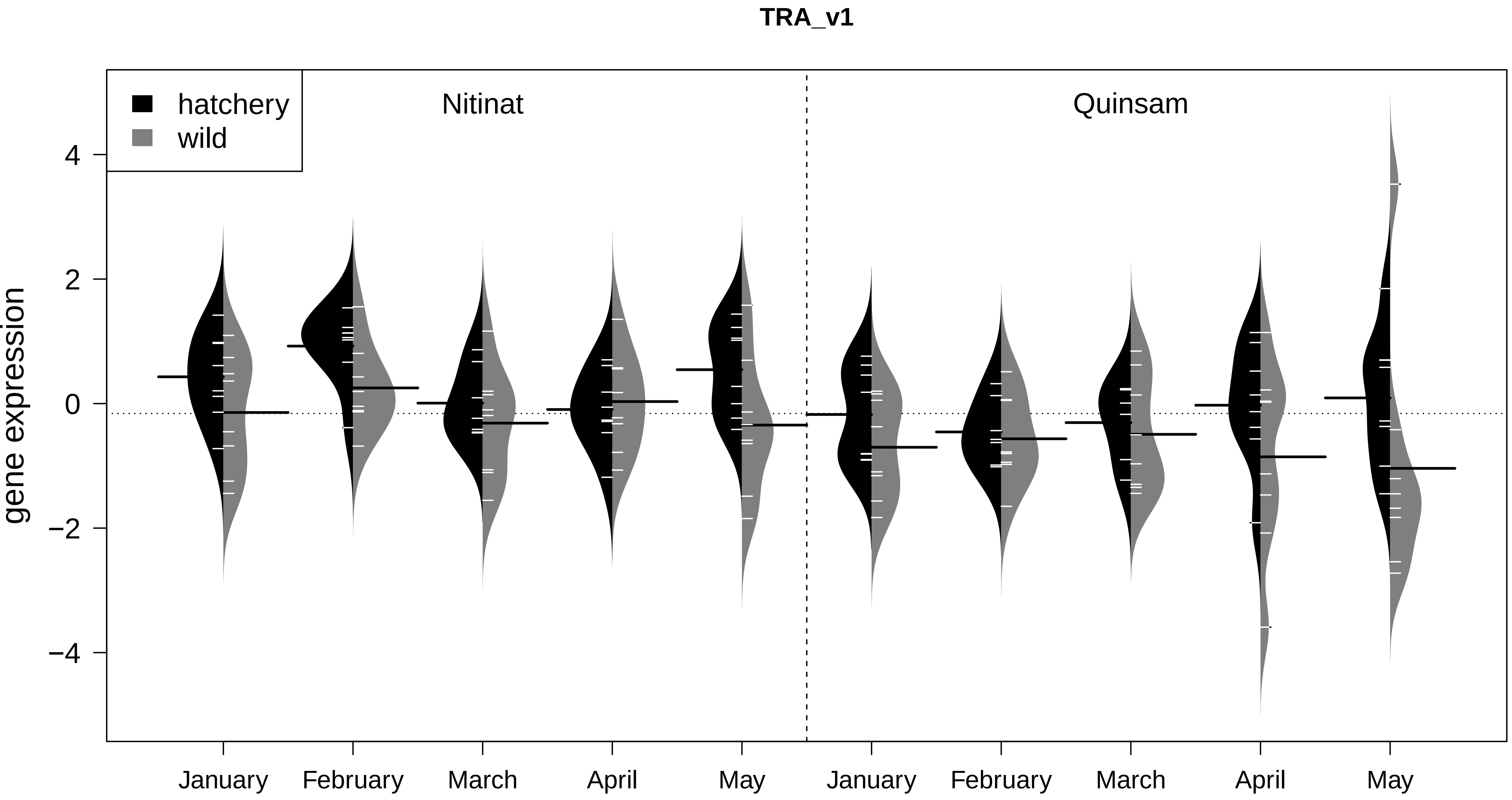
TRA_v1
hatchery
wild
Nitinat
Quinsam
gene expression
4
2
0
−2
−4
January
February
March
April
May
January
February
March
April
May

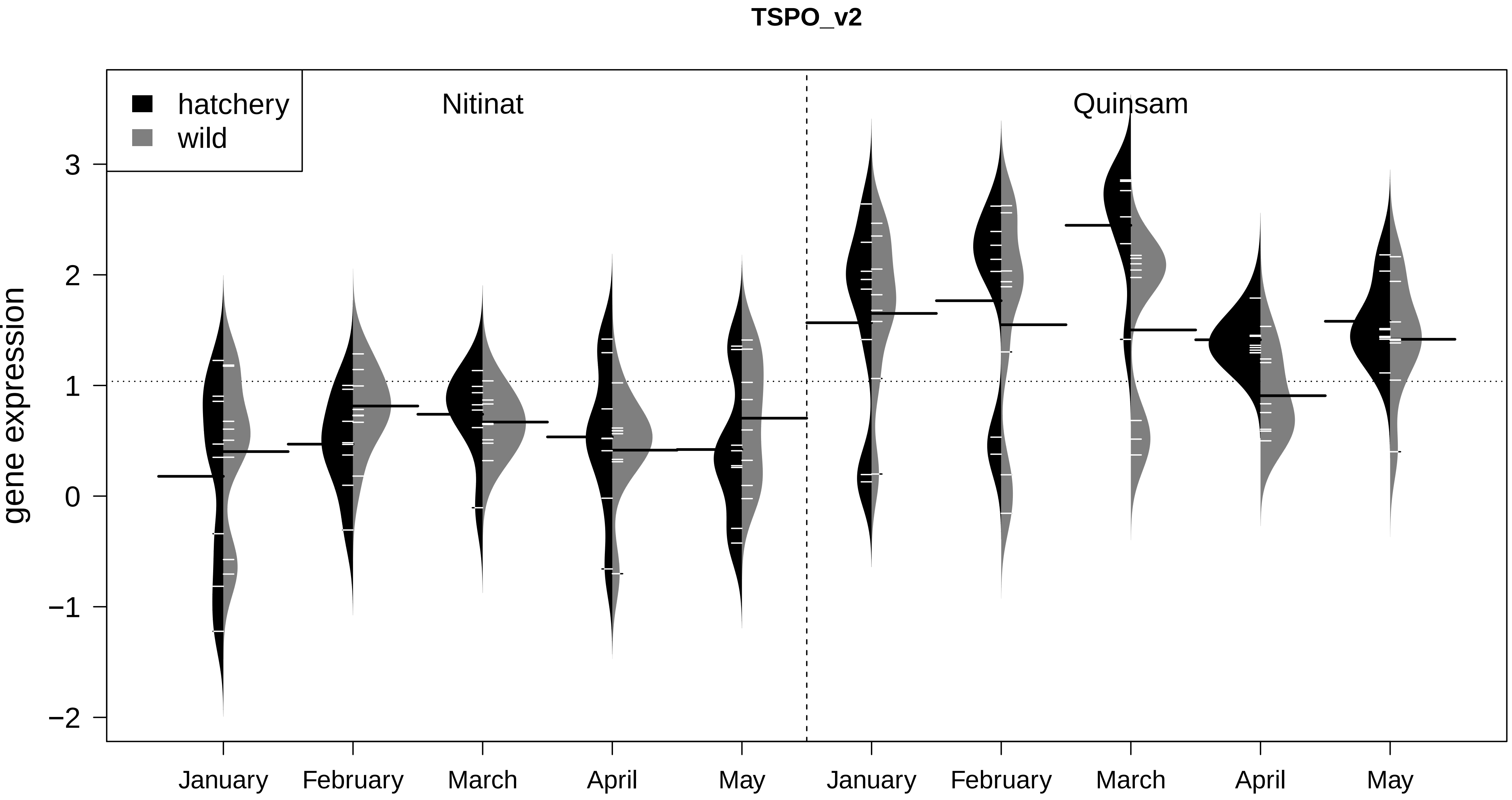
TSPO_v2
hatchery
wild
Nitinat
Quinsam
gene expression
3
2
1
0
−1
−2
January
February
March
April
May
January
February
March
April
May

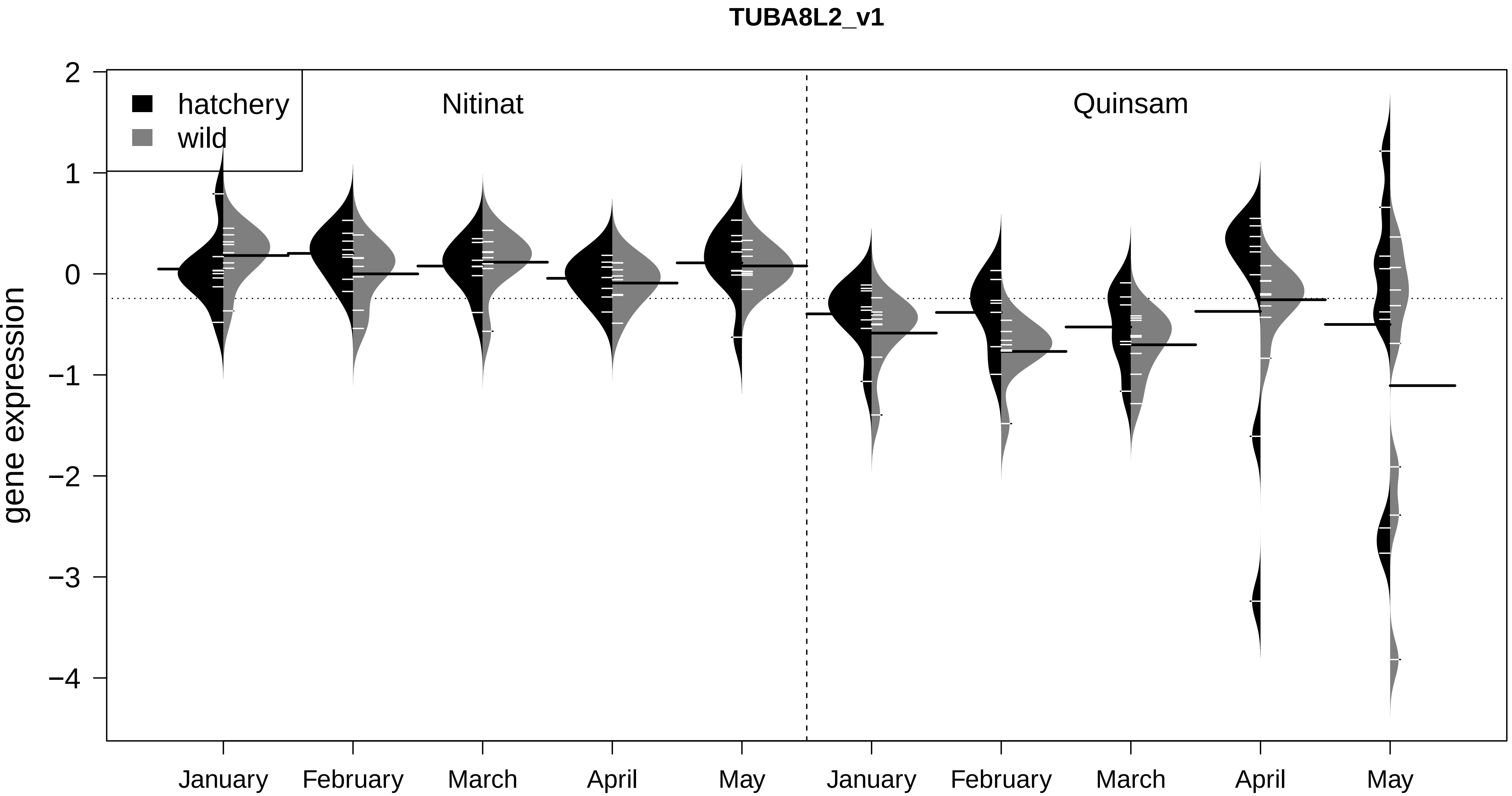

TUBA8L2_v1
hatchery
wild
Nitinat
Quinsam
gene expression
2
1
0
−1
−2
−3
−4
January
February
March
April
May
January
February
March
April
May

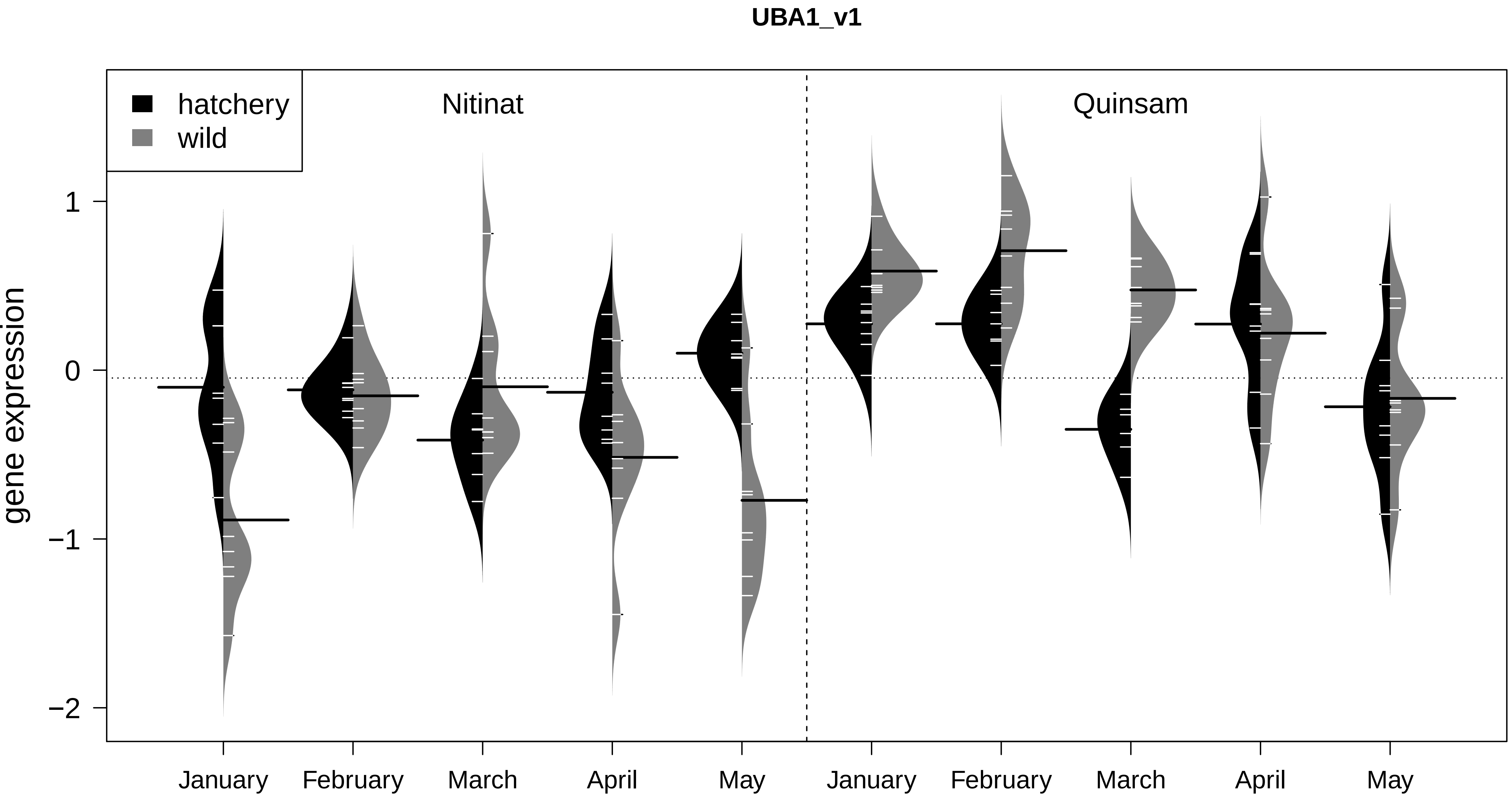
UBA1_v1
hatchery
wild
Nitinat
Quinsam
gene expression
1
0
−1
−2
January
February
March
April
May
January
February
March
April
May

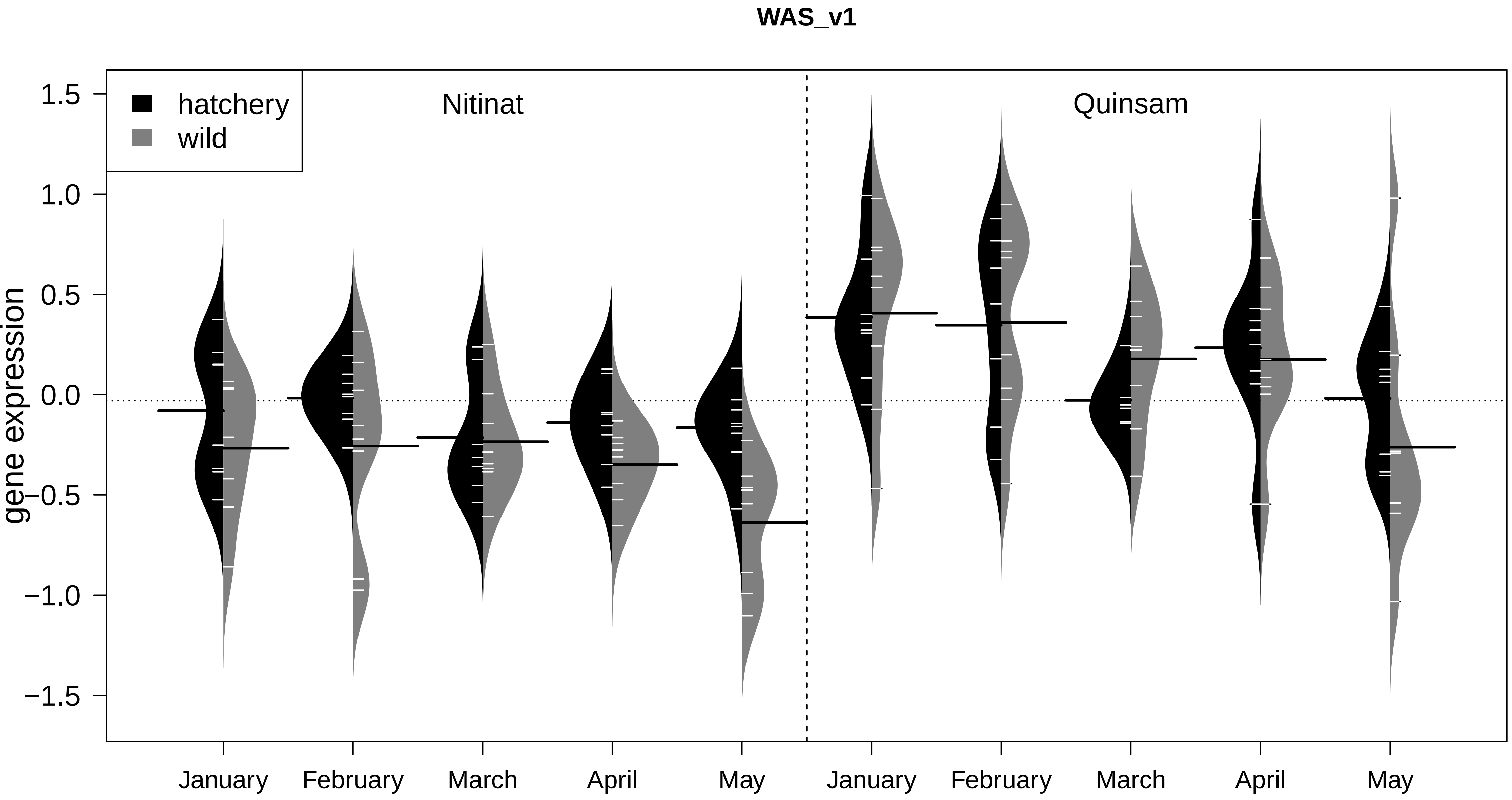

WAS_v1
hatchery
wild
Nitinat
Quinsam
gene expression
1.5
1.0
0.5
0.0
−0.5
−1.0
−1.5
January
February
March
April
May
January
February
March
April
May
