## appendix 2 for "Discovery and validation of candidate smoltification gene expression biomarkers across multiple species and ecotypes of Pacific salmonids"

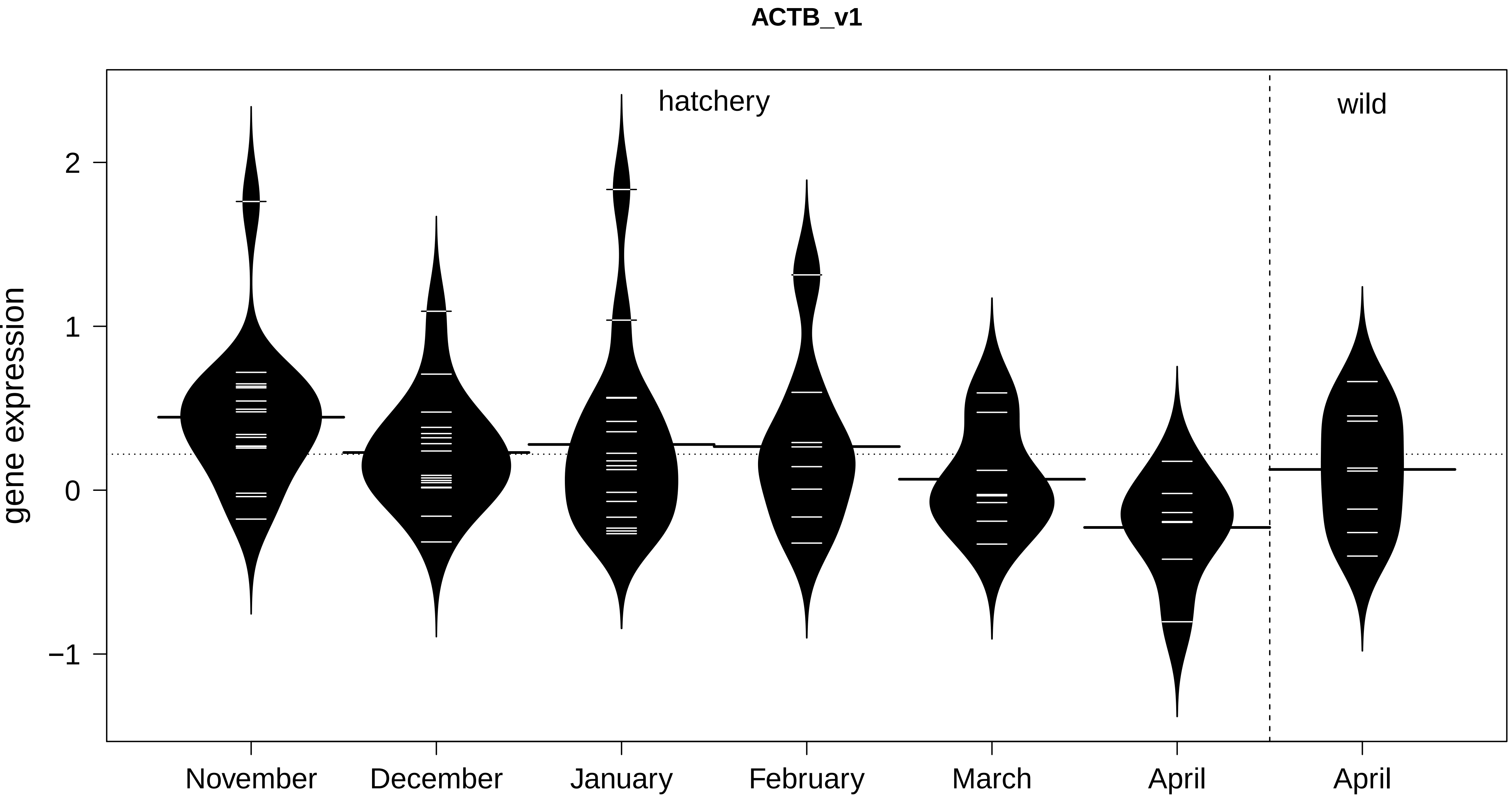
ACTB_v1
hatchery
wild
gene expression
2
1
0
−1
November
December
January
February
March
April
April

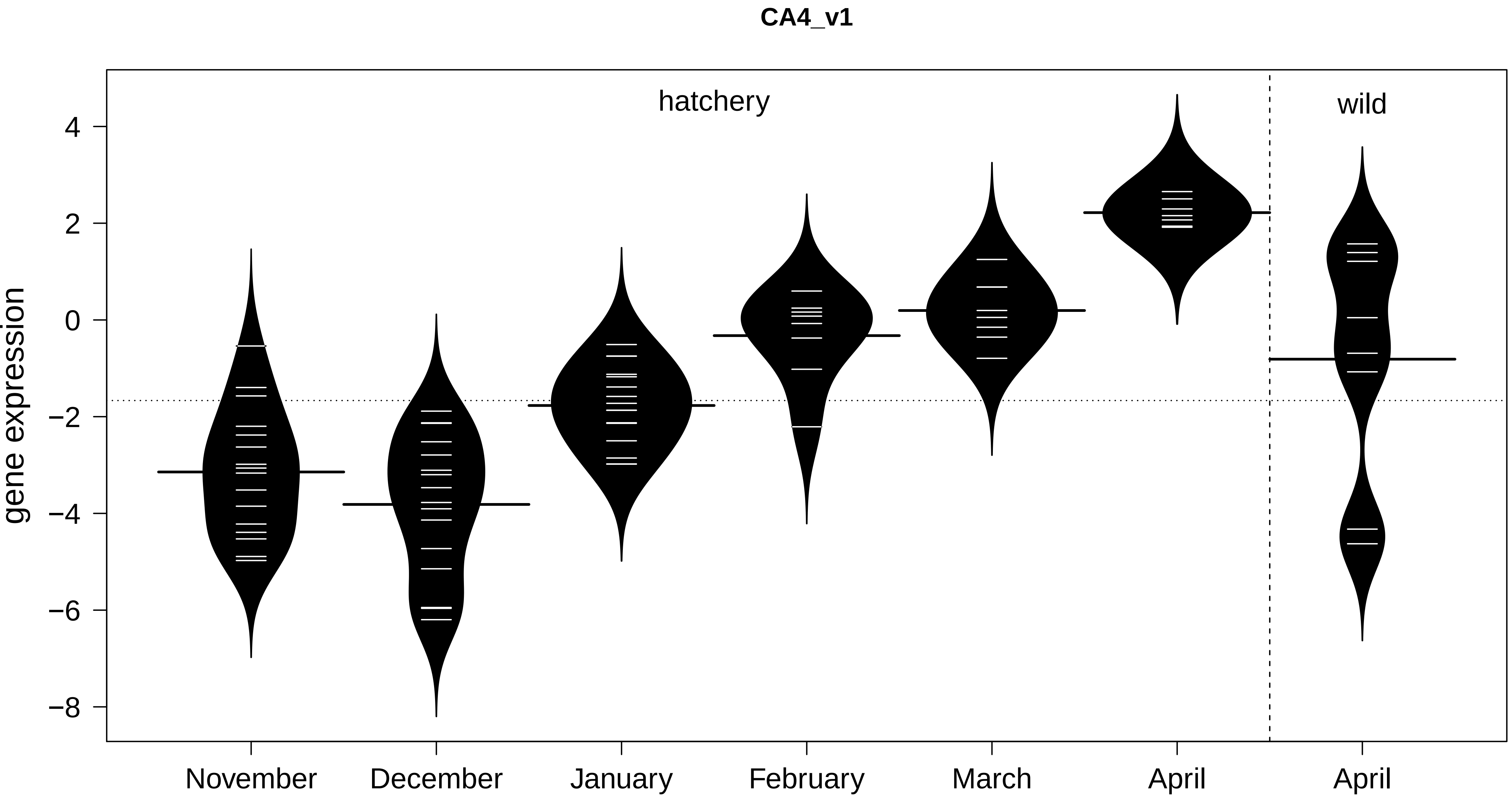
CA4_v1
hatchery
wild
gene expression
4
2
0
−2
−4
−6
−8
November
December
January
February
March
April
April

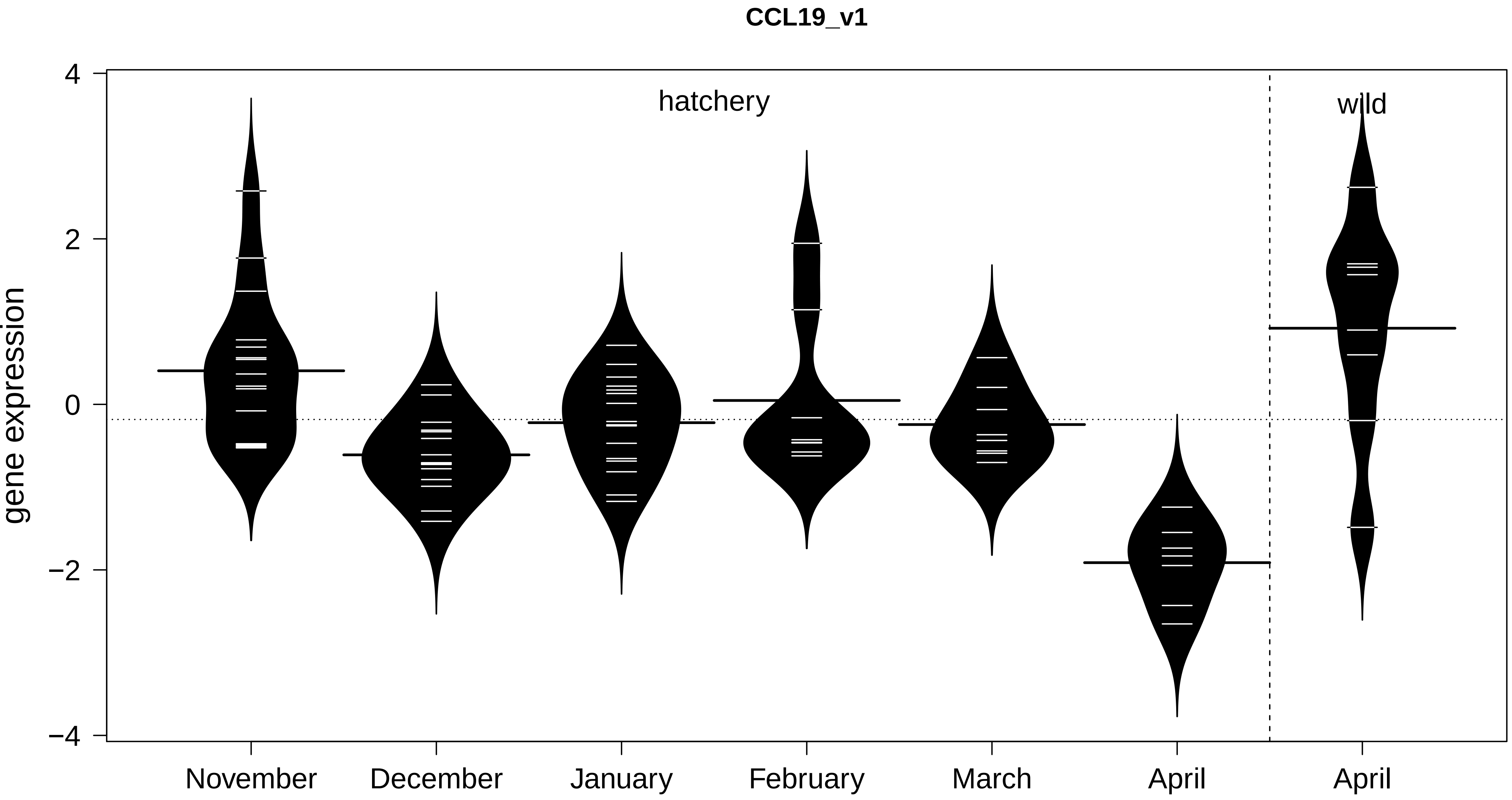

CCL19_v1
gene expression
4
2
0
−2
−4
hatchery
wild
November
December
January
February
March
April
April

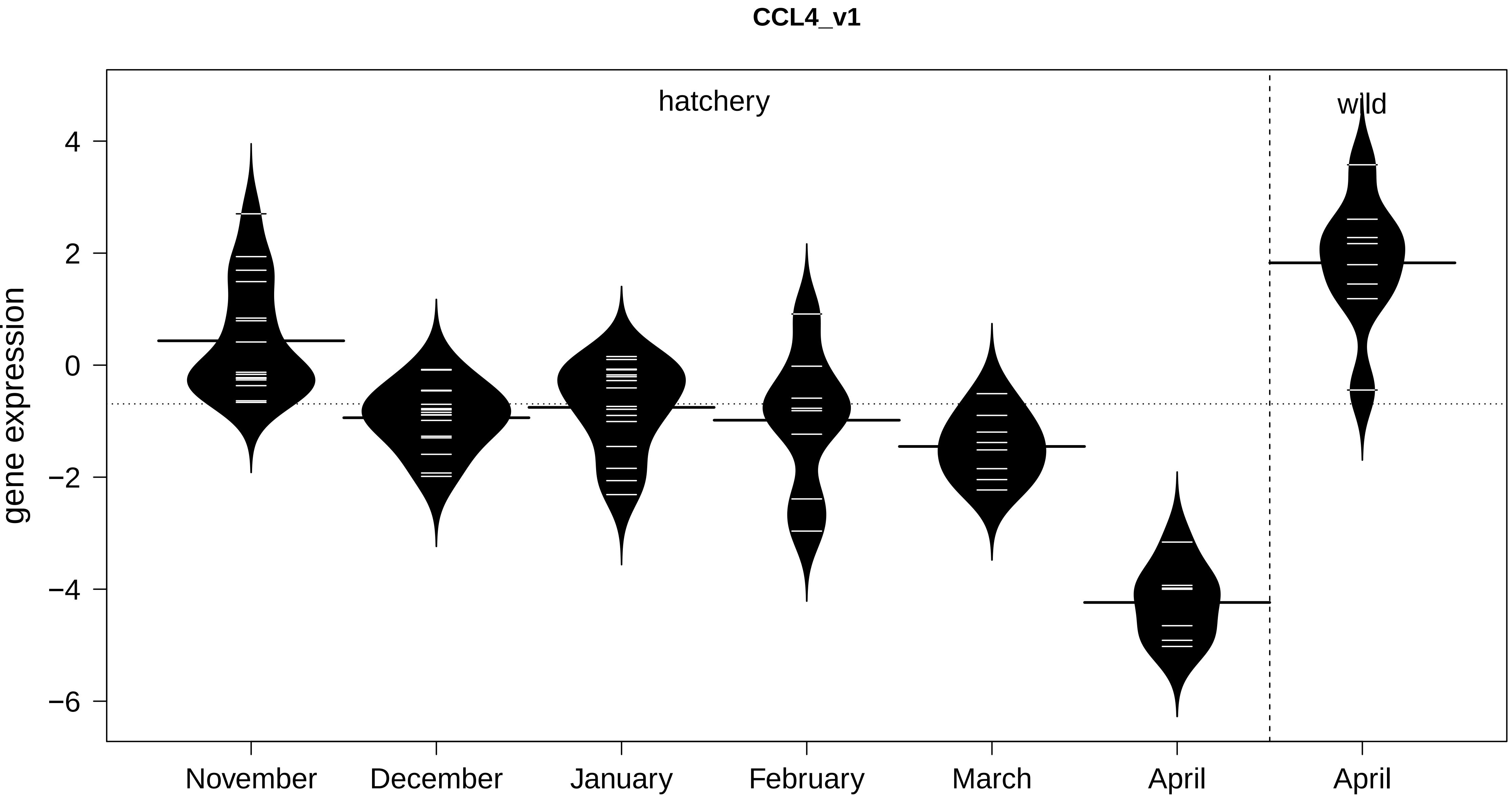

CCL4_v1
hatchery
wild
gene expression
4
2
0
−2
−4
−6
November
December
January
February
March
April
April

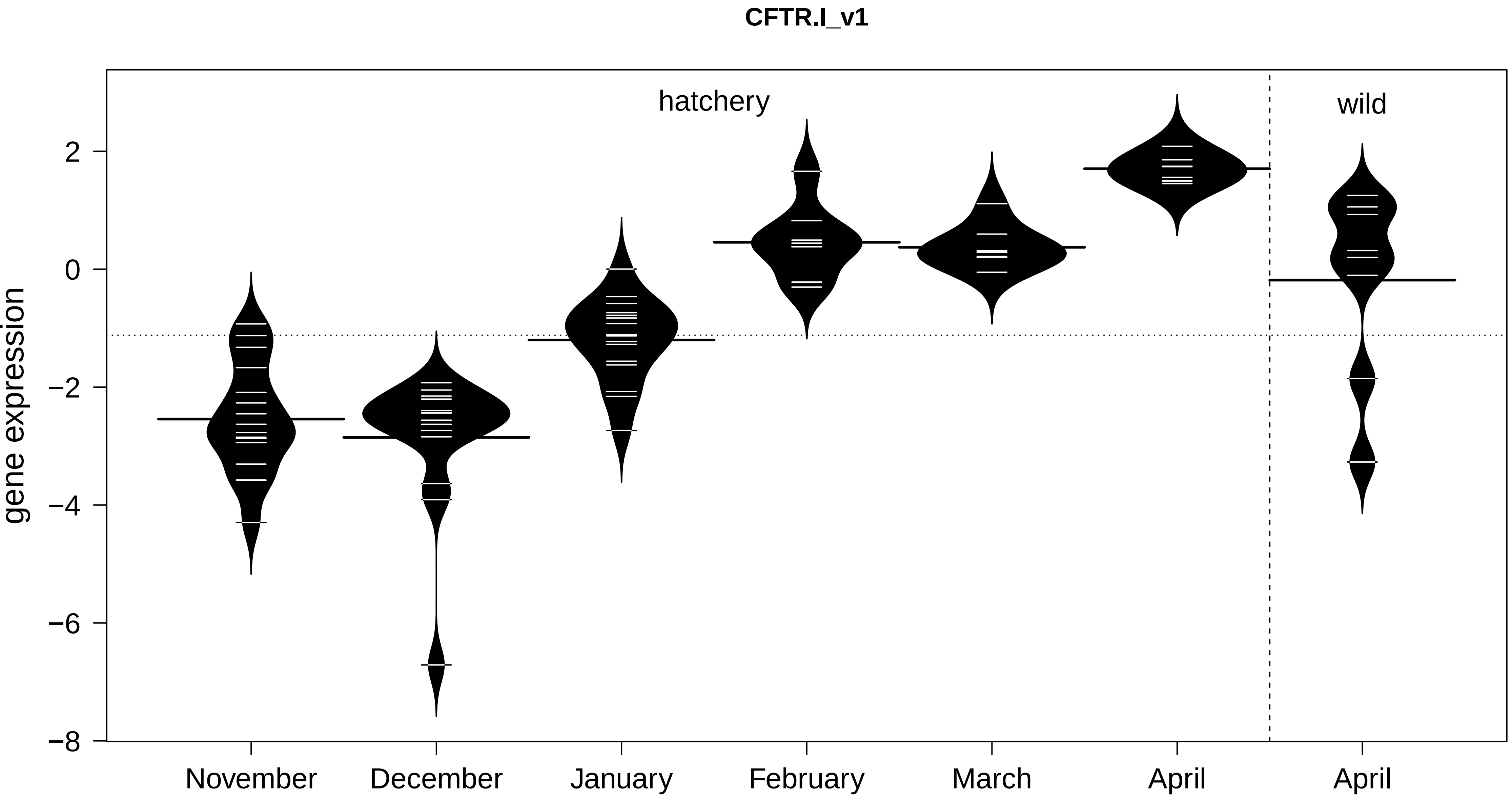
CFTR.I_v1
hatchery
wild
gene expression
2
0
−2
−4
−6
−8
November
December
January
February
March
April
April

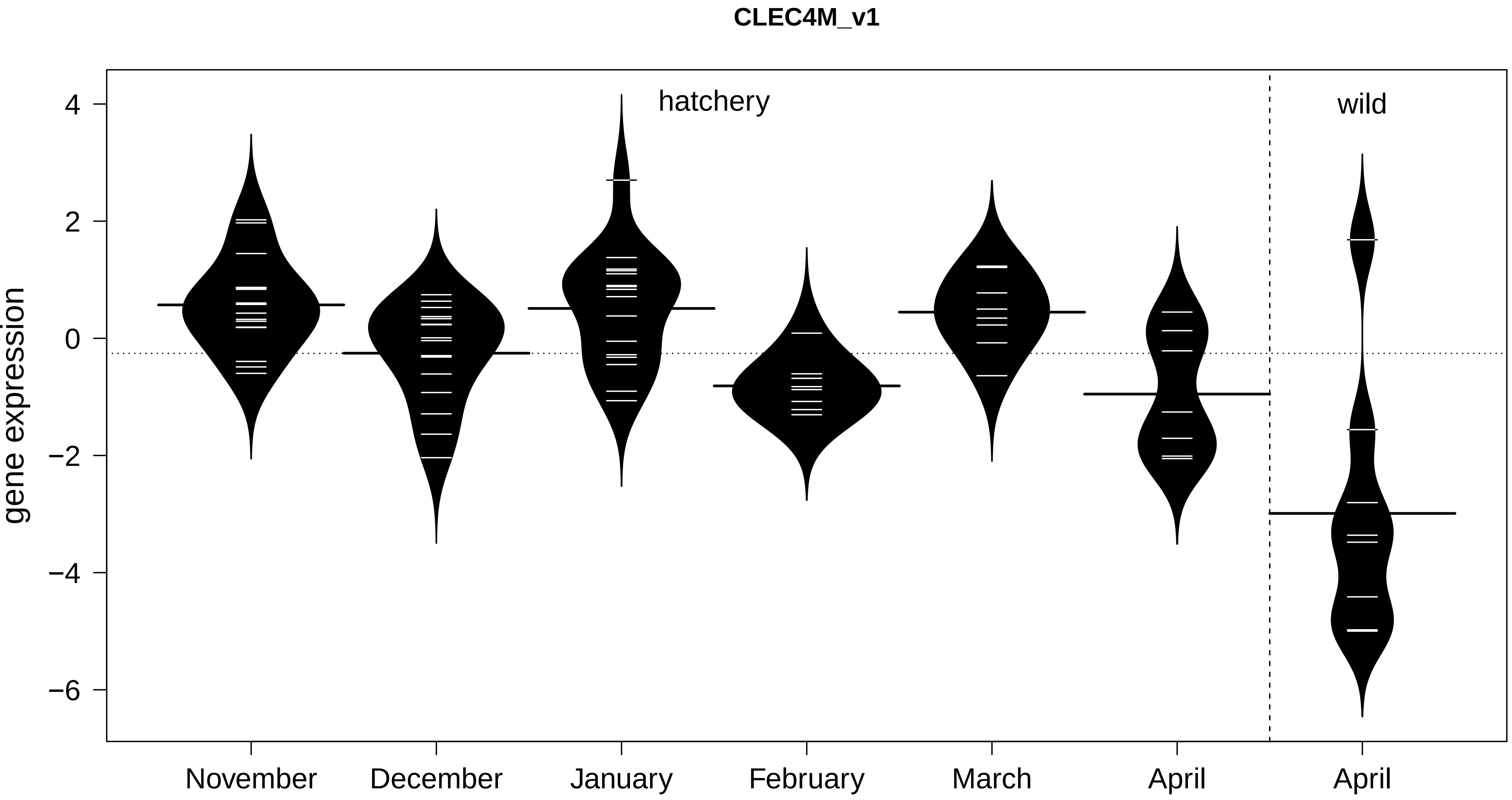
CLEC4M_v1
hatchery
wild
gene expression
4
2
0
−2
−4
−6
November
December
January
February
March
April
April

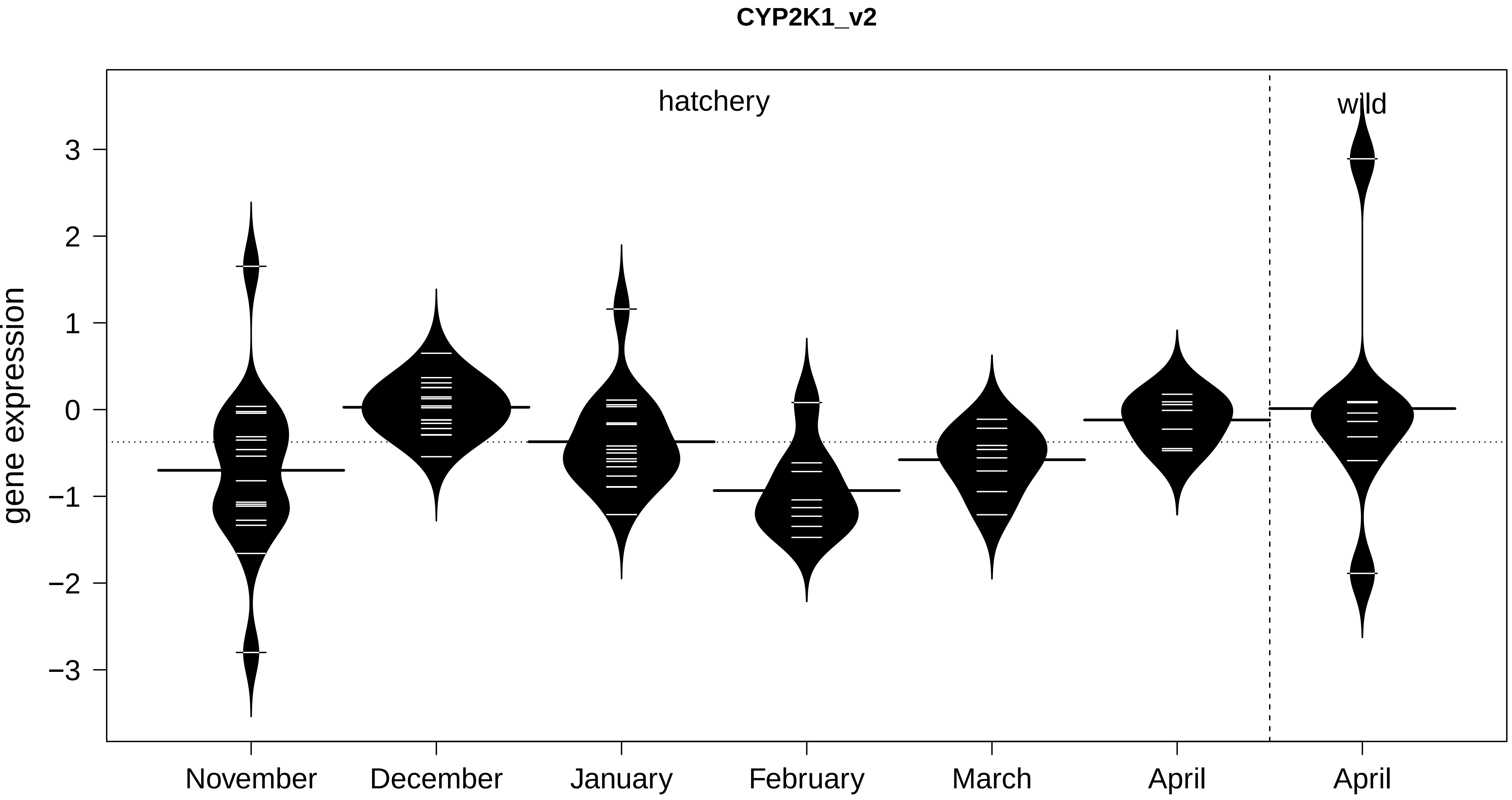

CYP2K1_v2
hatchery
wild
gene expression
3
2
1
0
−1
−2
−3
November
December
January
February
March
April
April

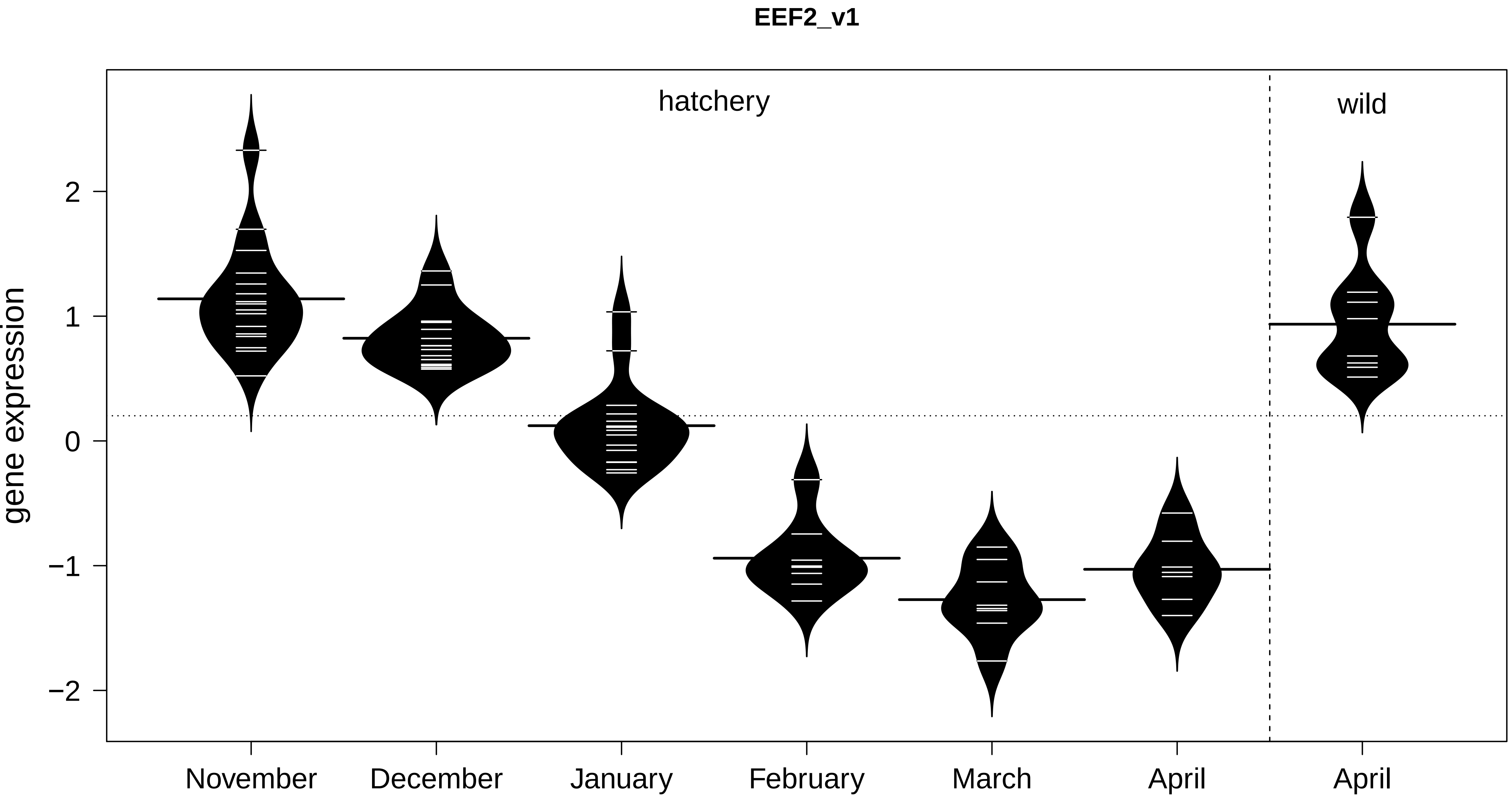
EEF2_v1
hatchery
wild
gene expression
2
1
0
−1
−2
November
December
January
February
March
April
April

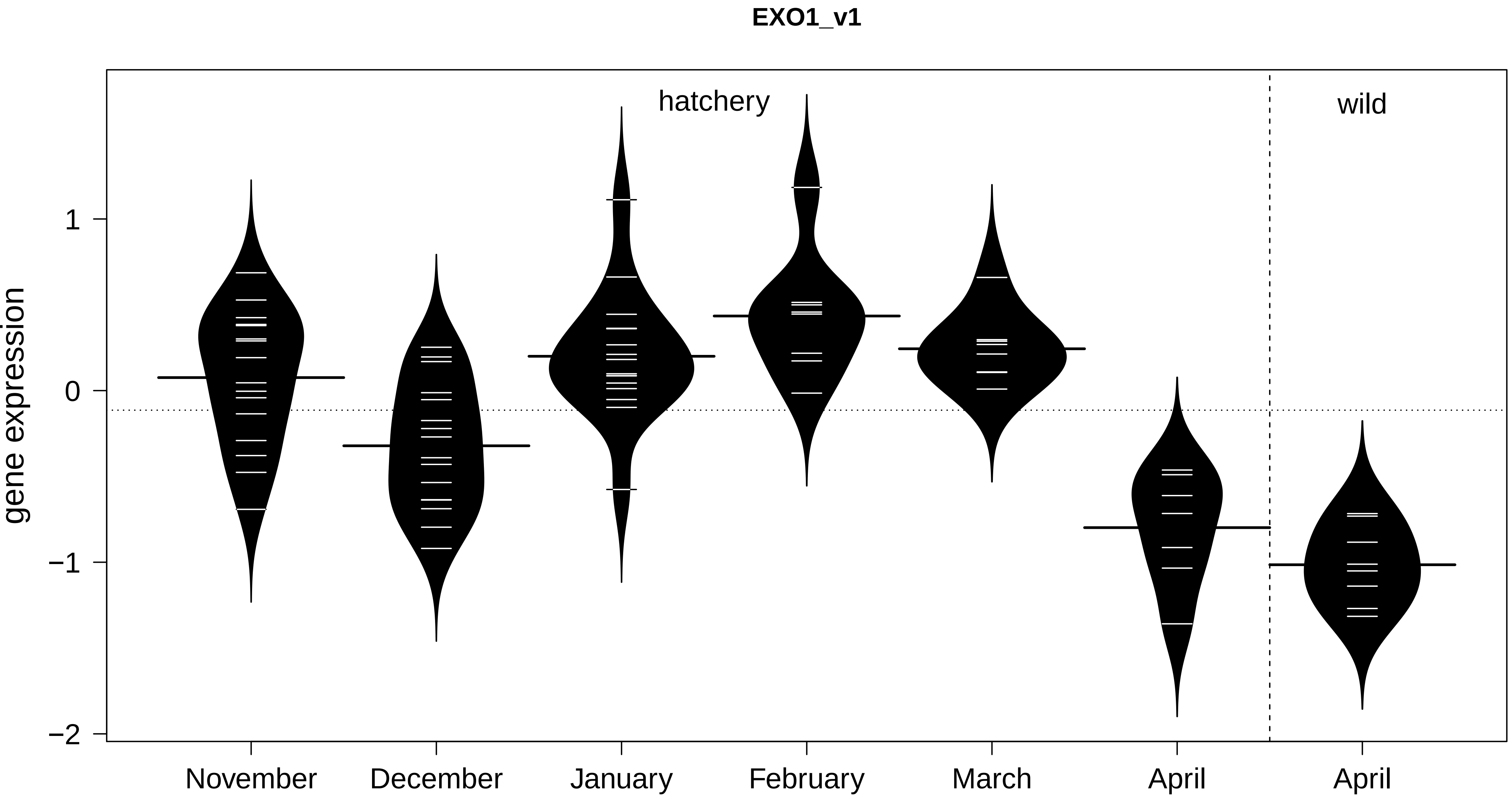
EXO1_v1
hatchery
wild
gene expression
1
0
−1
−2
November
December
January
February
March
April
April

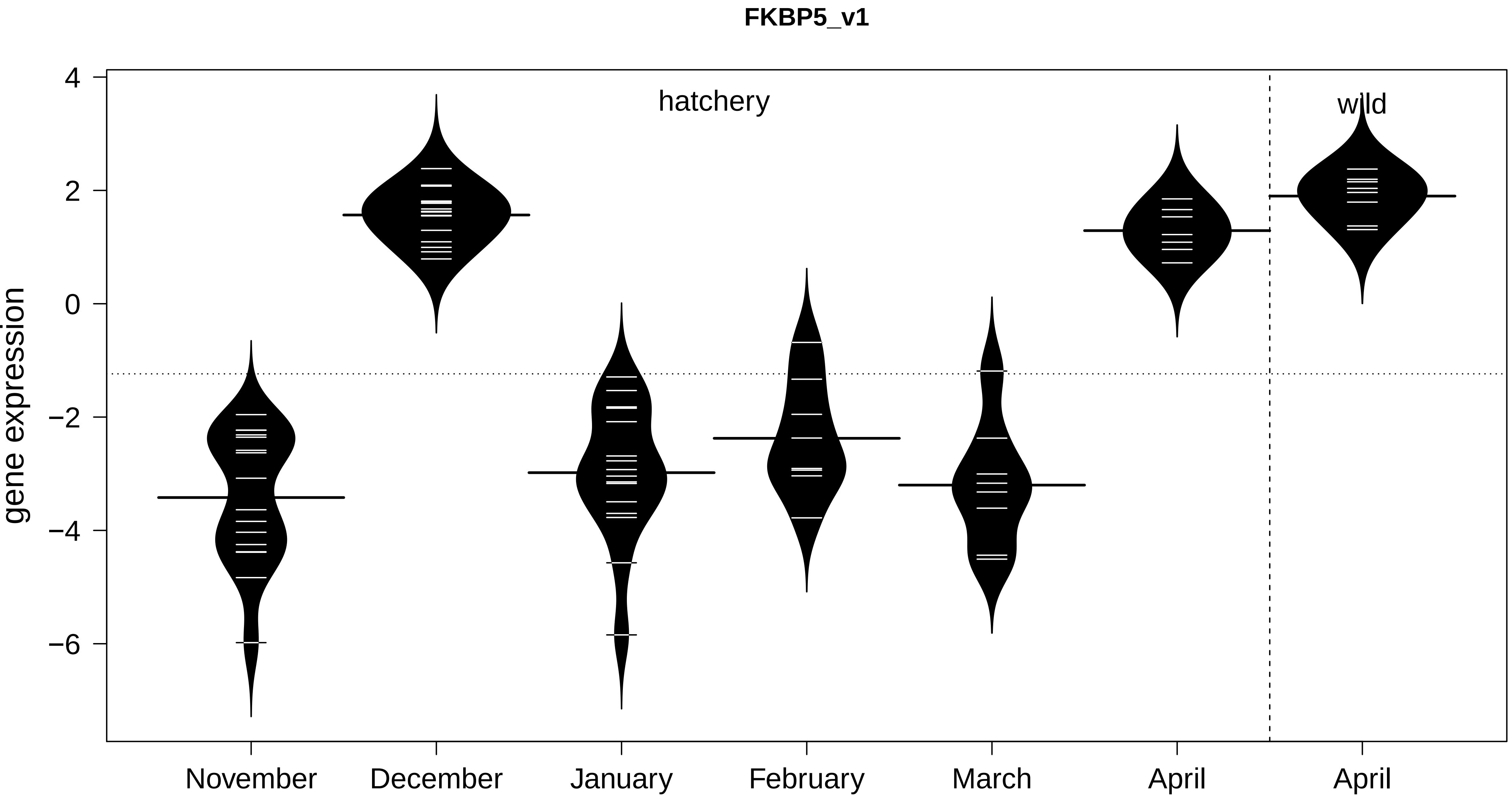

FKBP5_v1
hatchery
wild
gene expression
4
2
0
−2
−4
−6
November
December
January
February
March
April
April

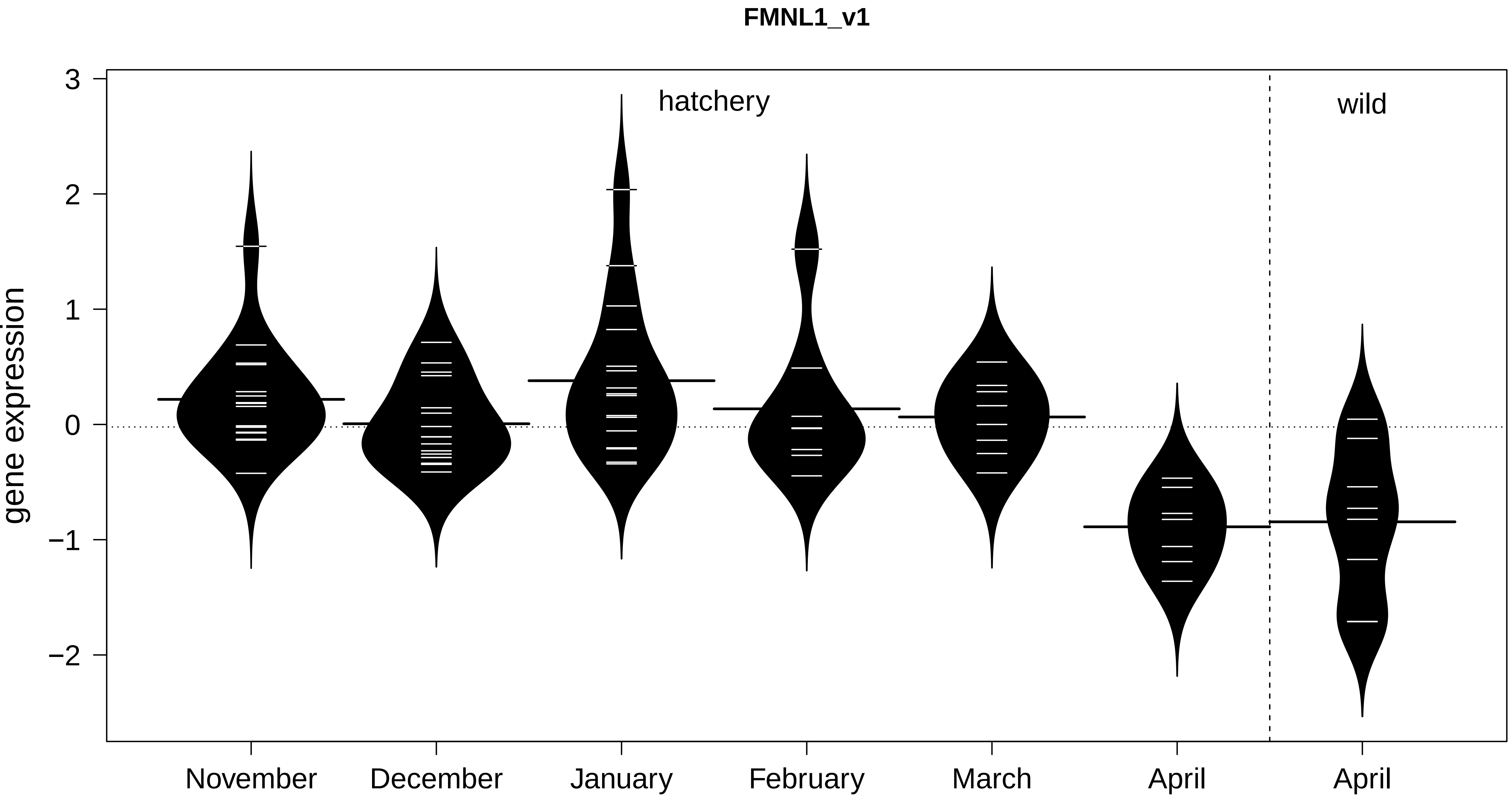

FMNL1_v1
gene expression
3
2
1
0
−1
−2
hatchery
wild
November
December
January
February
March
April
April

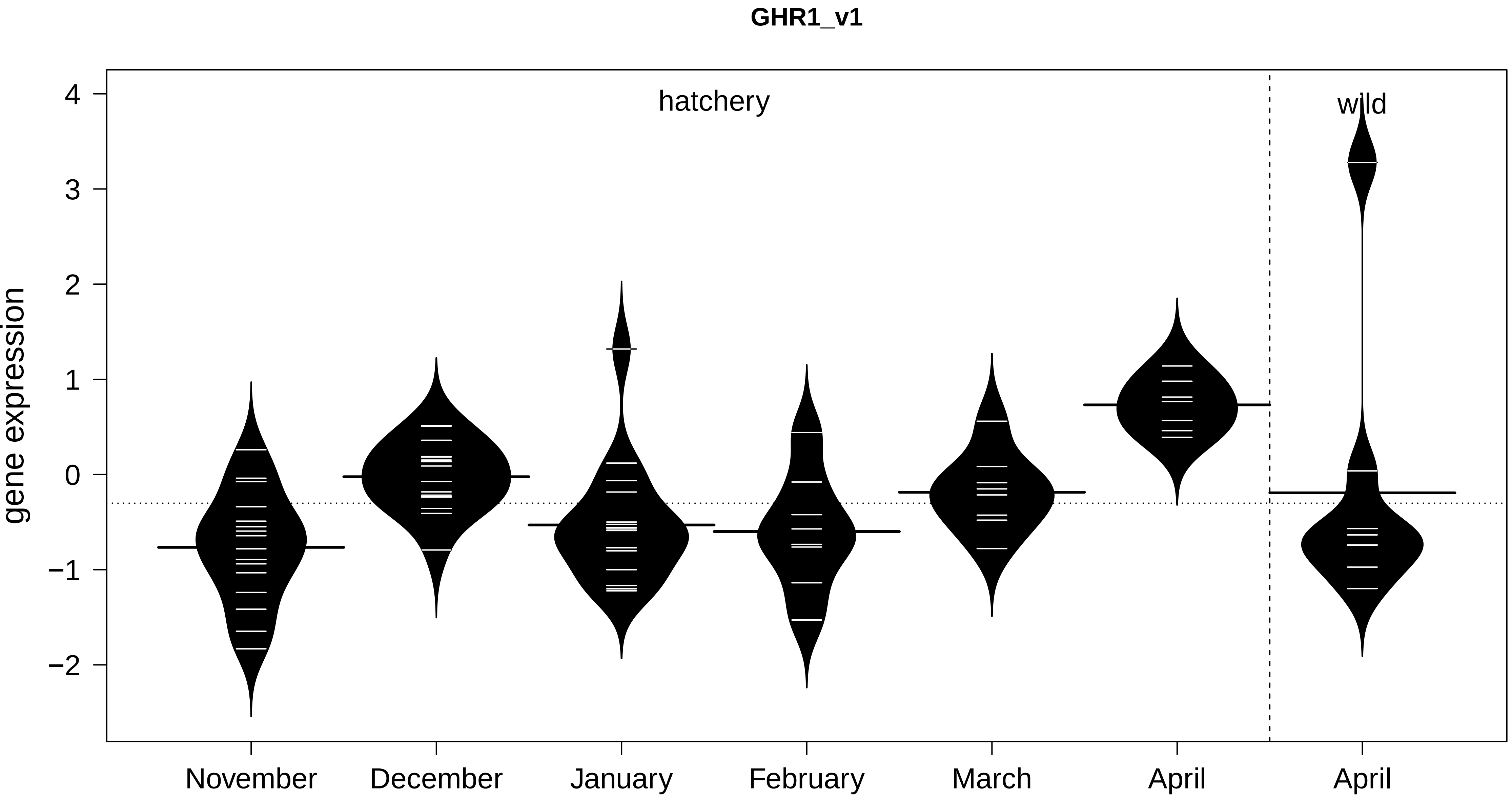
GHR1_v1
hatchery
wild
gene expression
4
3
2
1
0
−1
−2
November
December
January
February
March
April
April

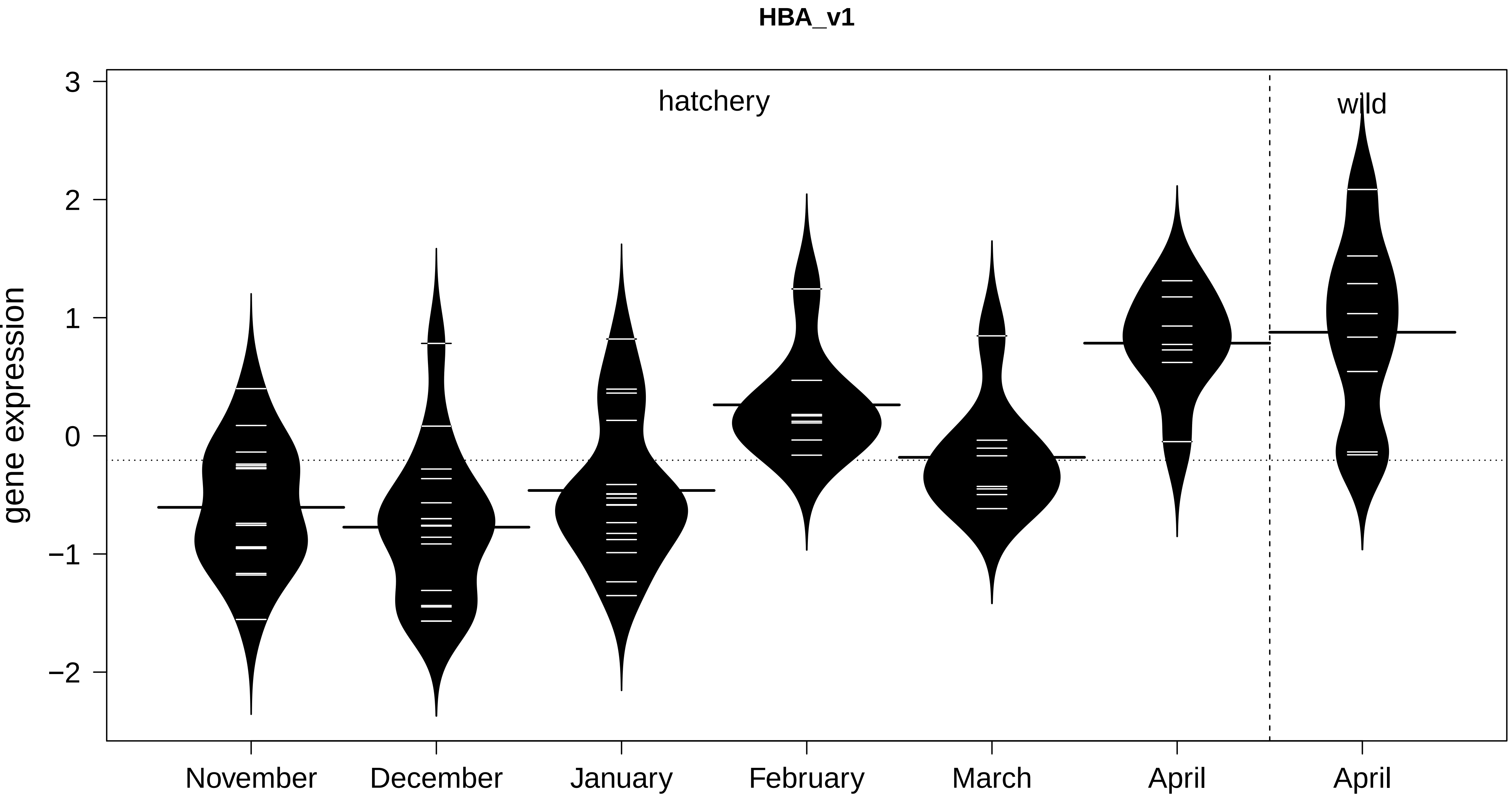

HBA_v1
hatchery
wild
gene expression
3
2
1
0
−1
−2
November
December
January
February
March
April
April

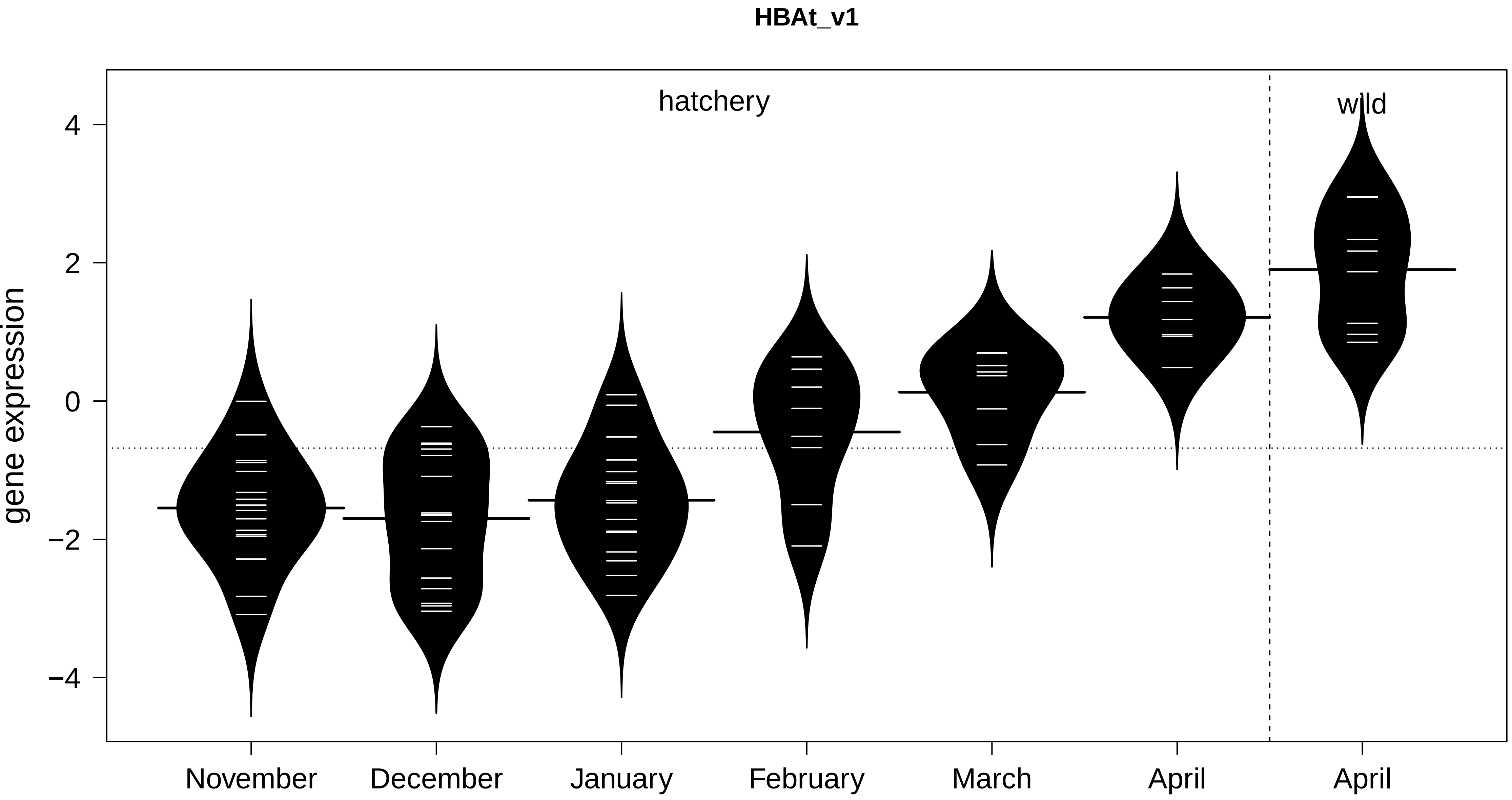
HBAt_v1
hatchery
wild
gene expression
4
2
0
−2
−4
November
December
January
February
March
April
April

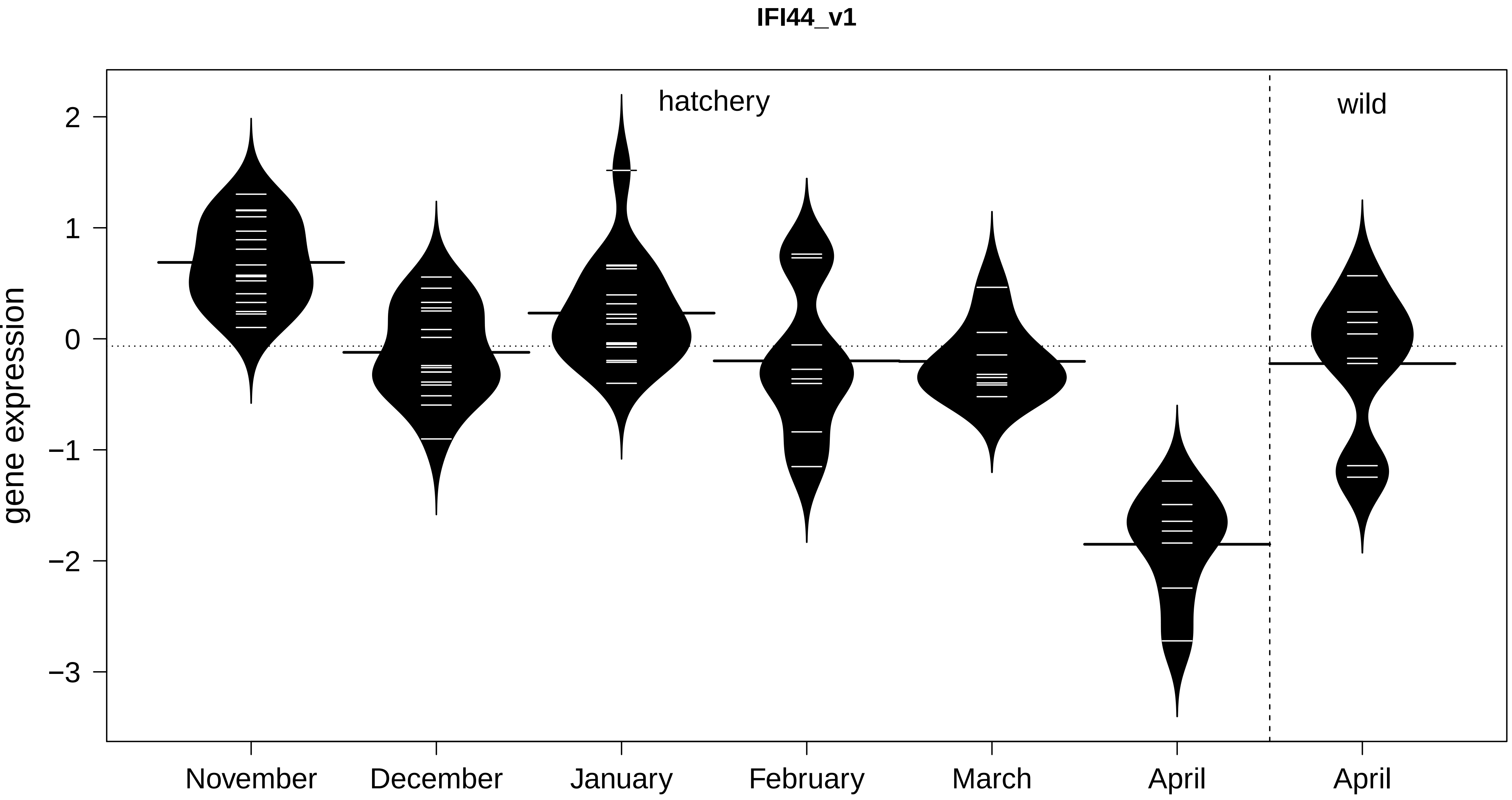

IFI44_v1
hatchery
wild
gene expression
2
1
0
−1
−2
−3
November
December
January
February
March
April
April

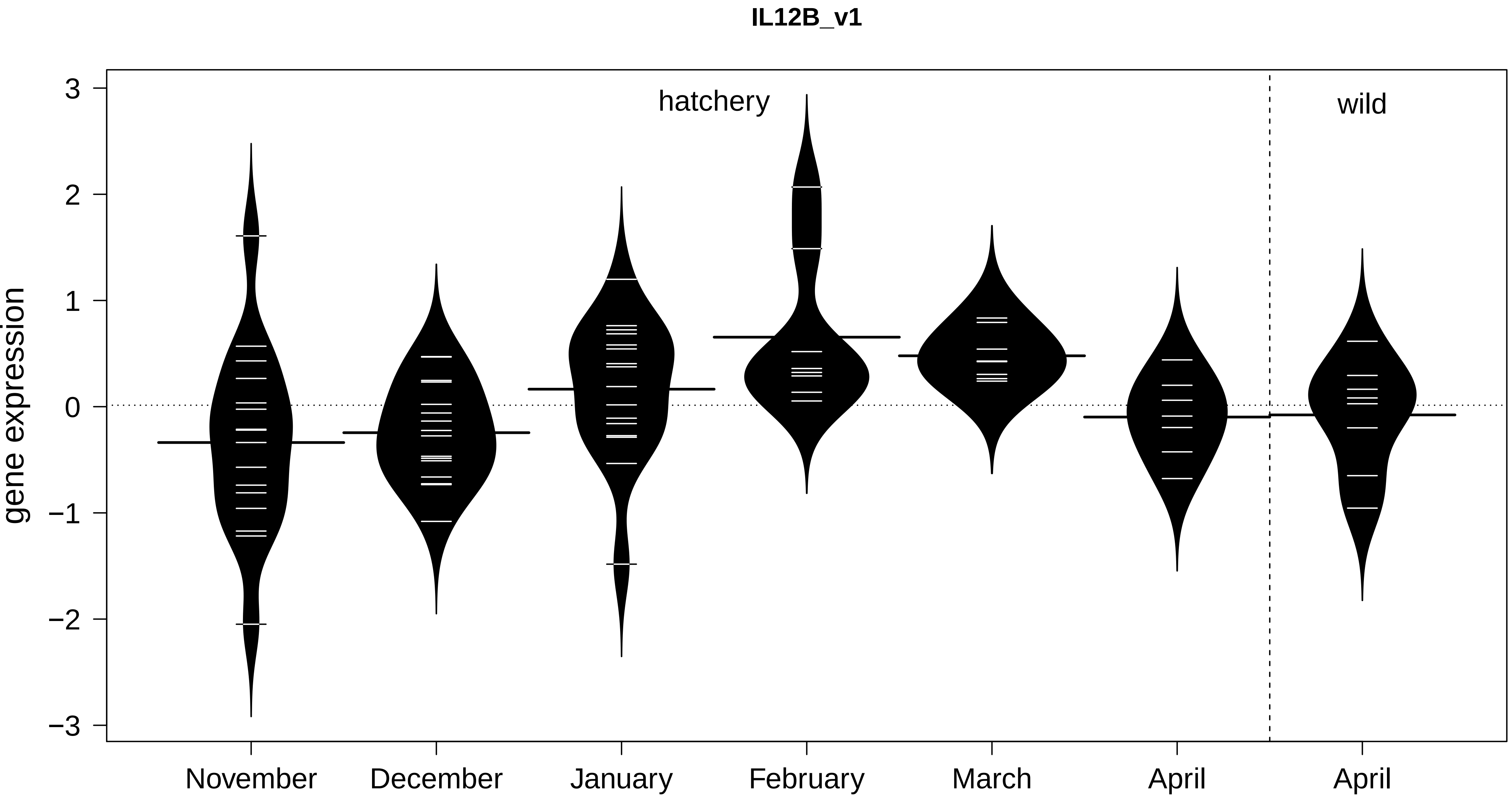
IL12B_v1
gene expression
3
2
1
0
−1
−2
−3
hatchery
wild
November
December
January
February
March
April
April

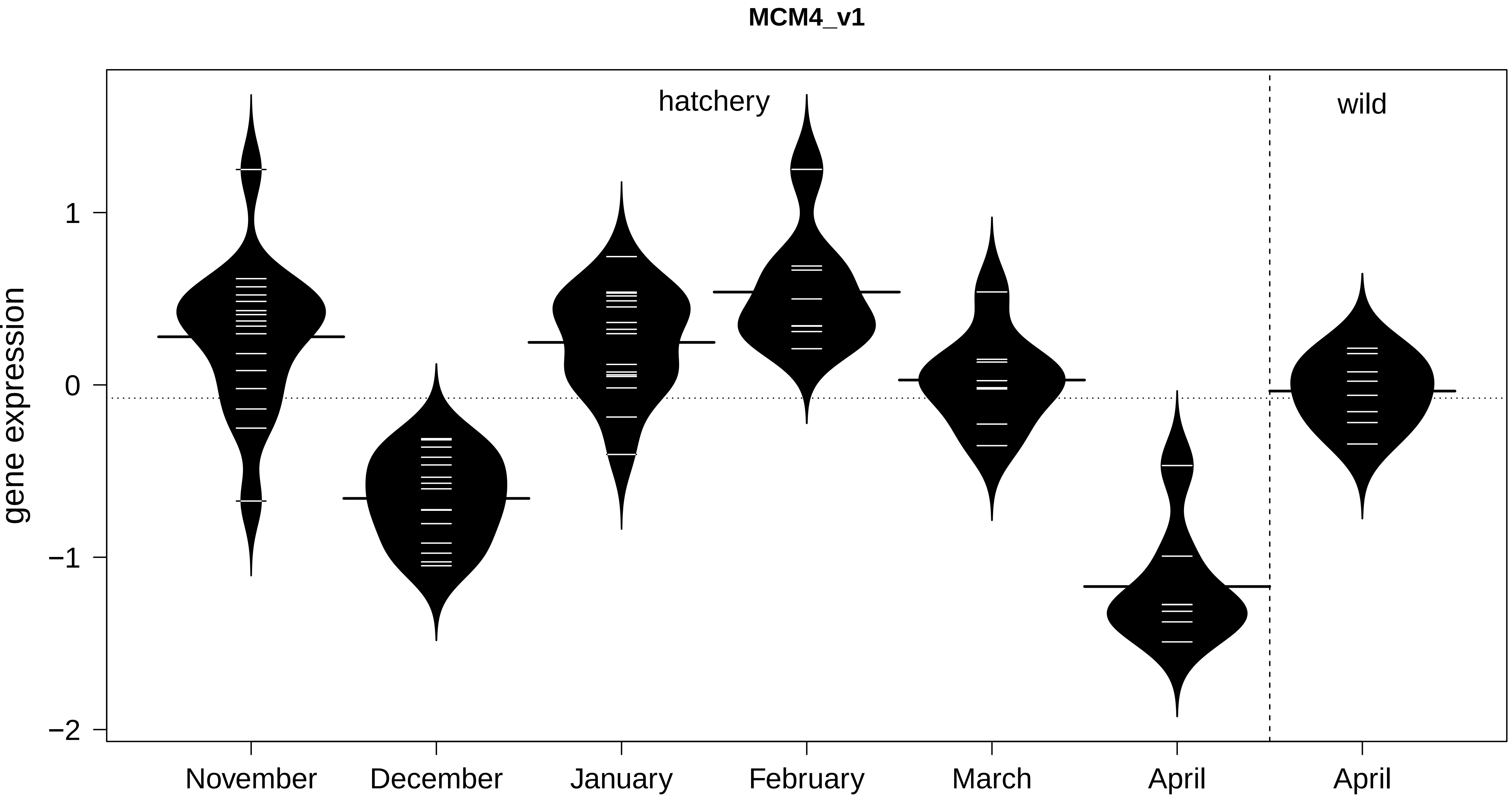

MCM4_v1
hatchery
wild
gene expression
1
0
−1
−2
November
December
January
February
March
April
April

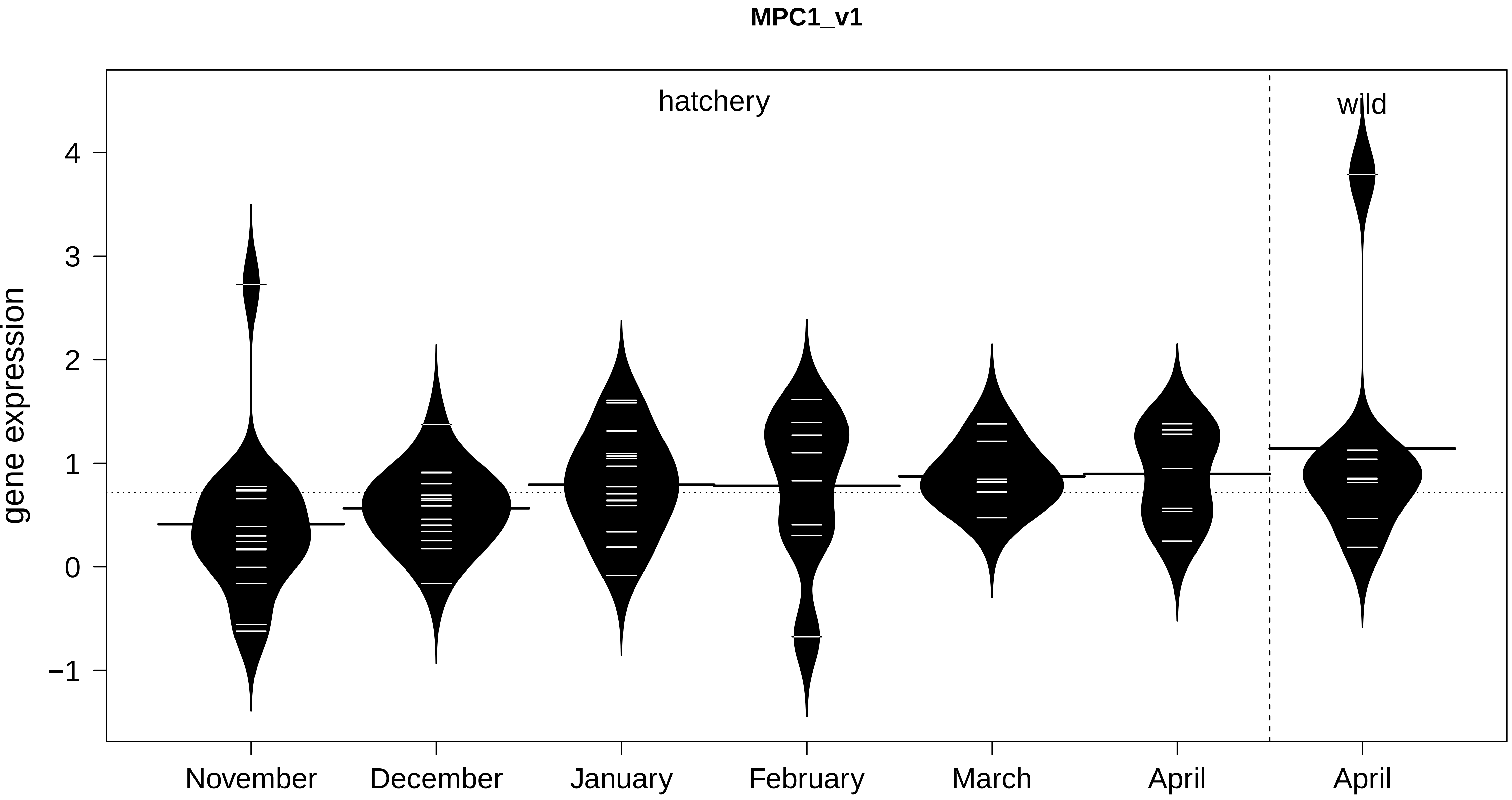

MPC1_v1
hatchery
wild
gene expression
4
3
2
1
0
−1
November
December
January
February
March
April
April

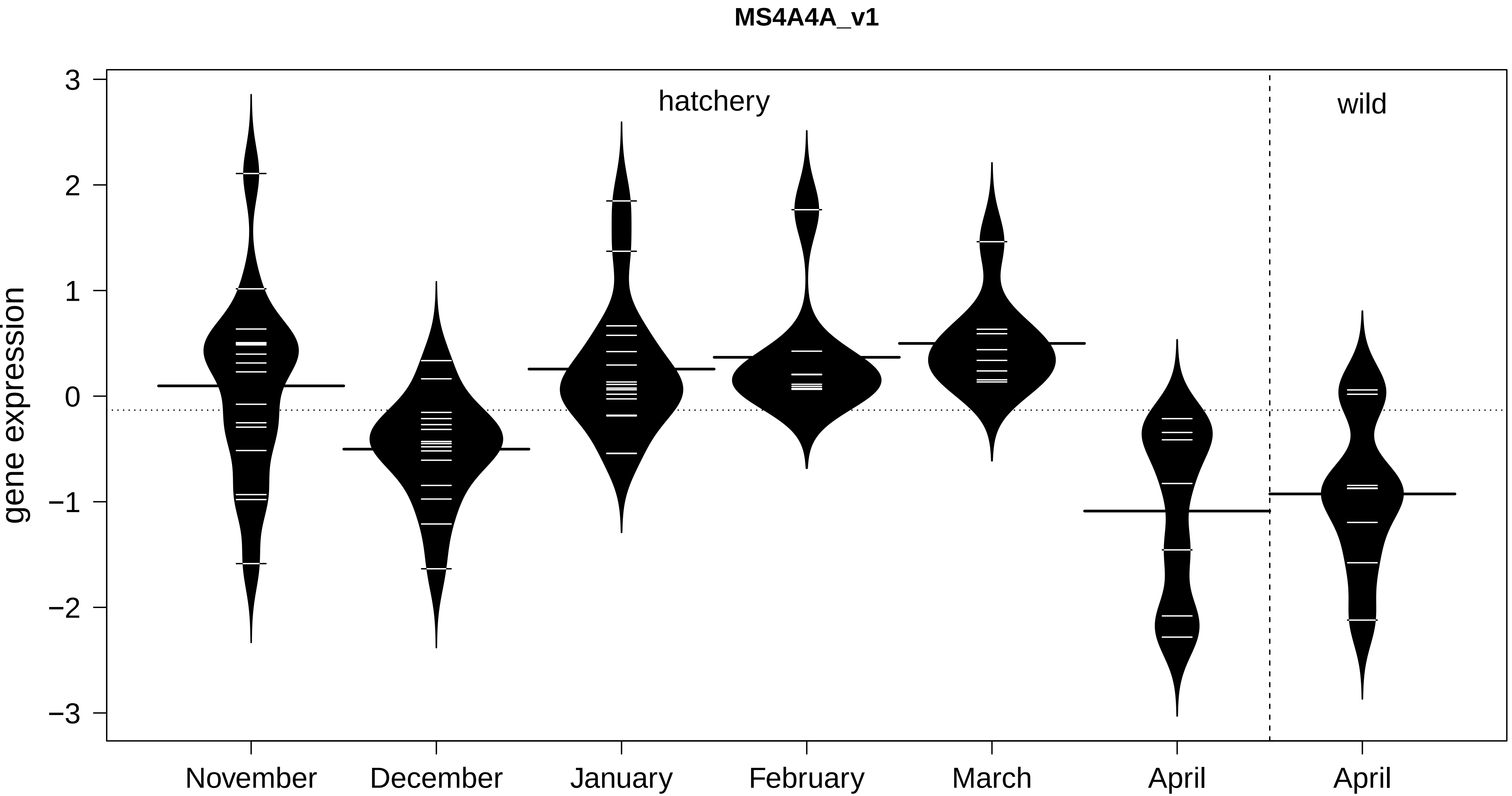
MS4A4A_v1
gene expression
3
2
1
0
−1
−2
−3
hatchery
wild
November
December
January
February
March
April
April

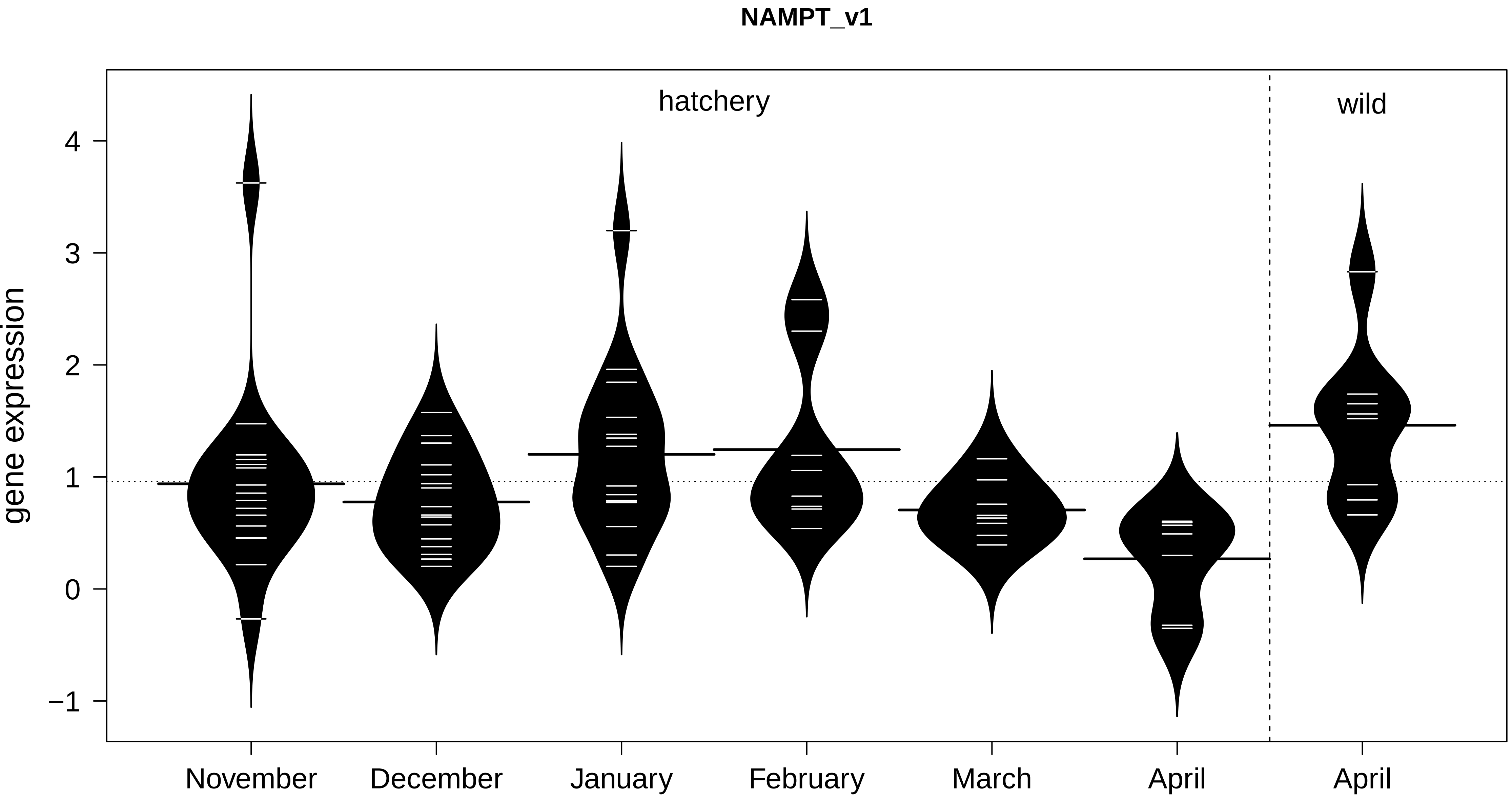
NAMPT_v1
hatchery
wild
gene expression
4
3
2
1
0
−1
November
December
January
February
March
April
April

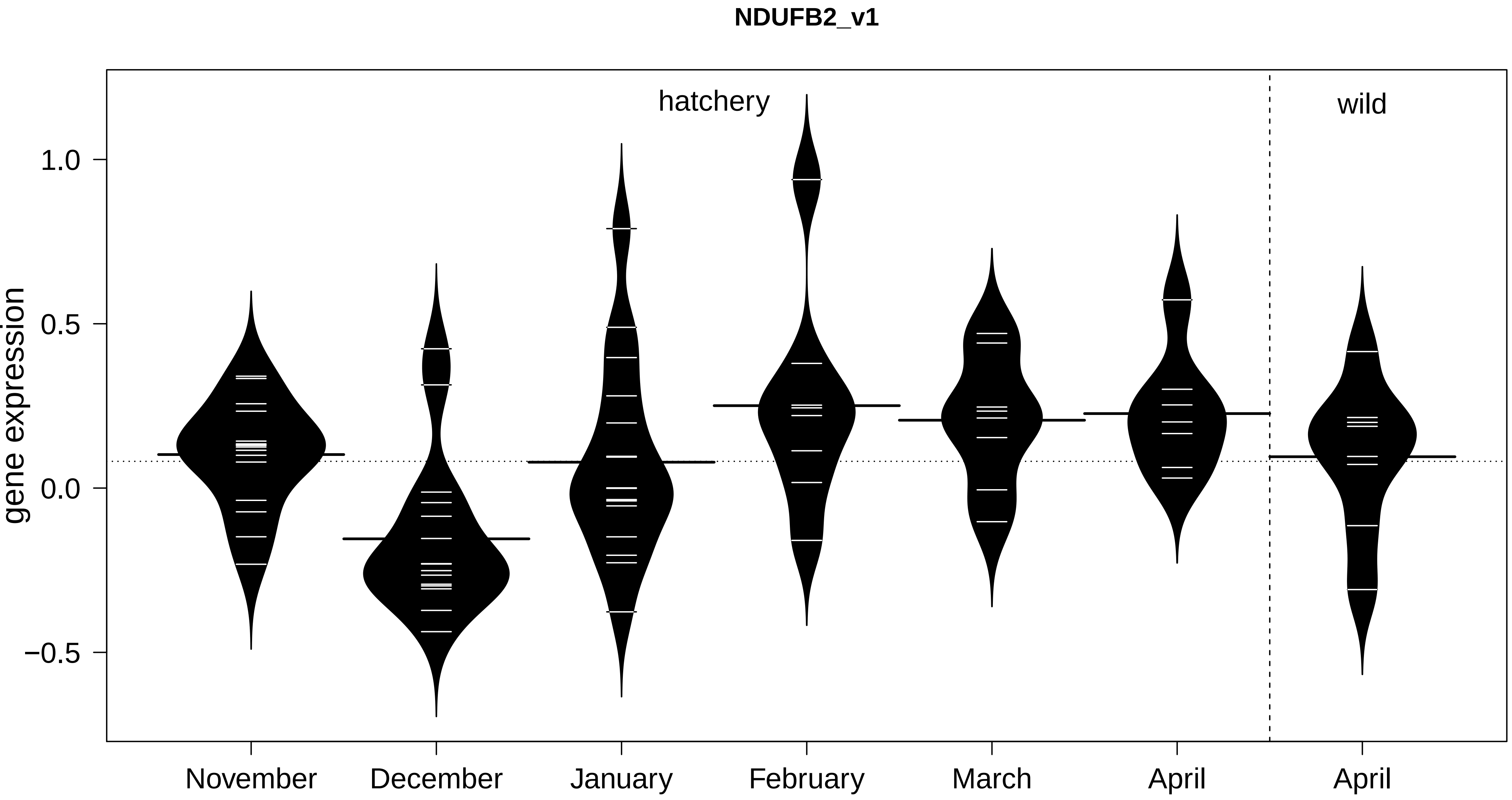
NDUFB2_v1
hatchery
wild
gene expression
1.0
0.5
0.0
−0.5
November
December
January
February
March
April
April

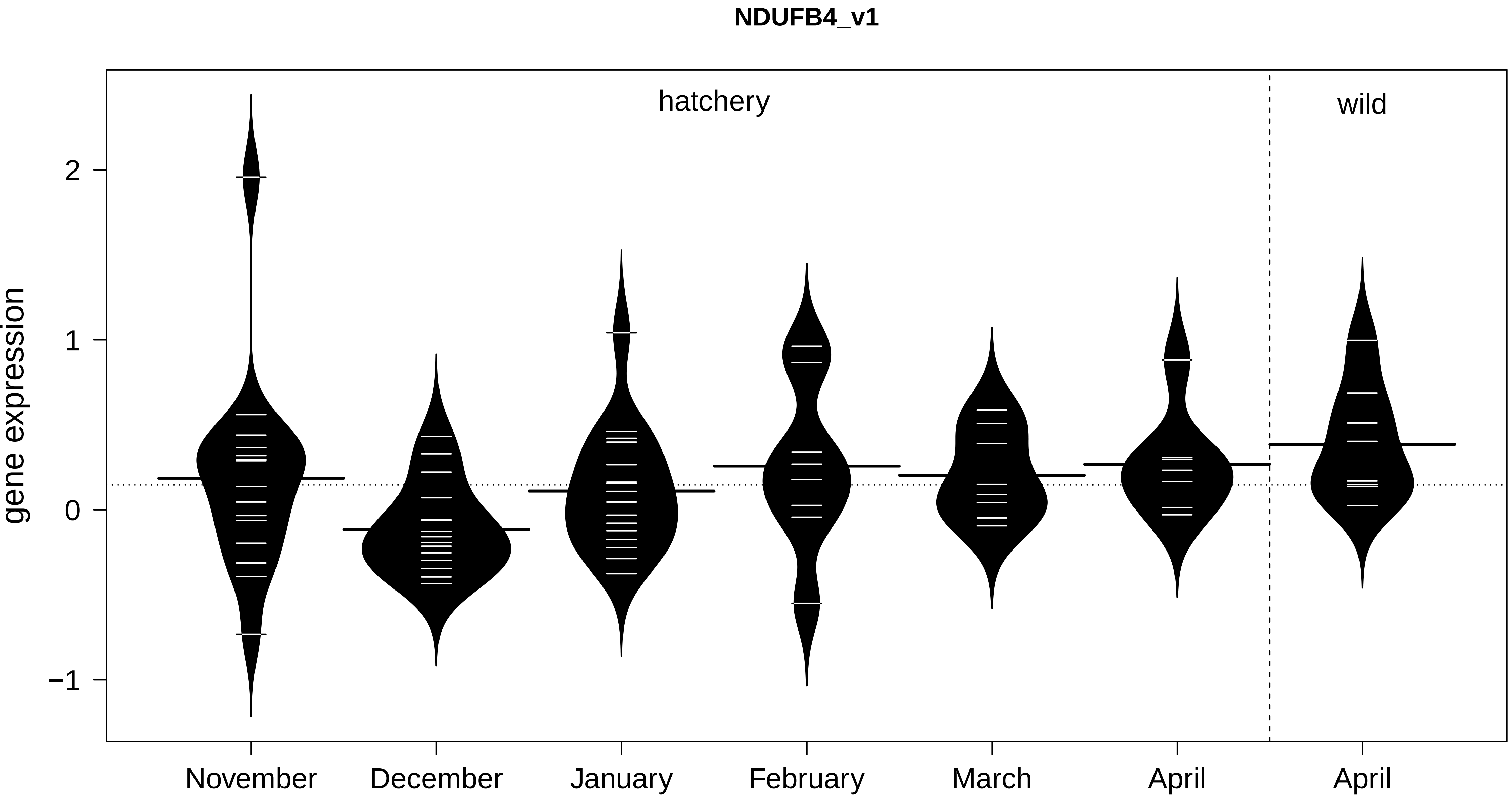

NDUFB4_v1
hatchery
wild
gene expression
2
1
0
−1
November
December
January
February
March
April
April

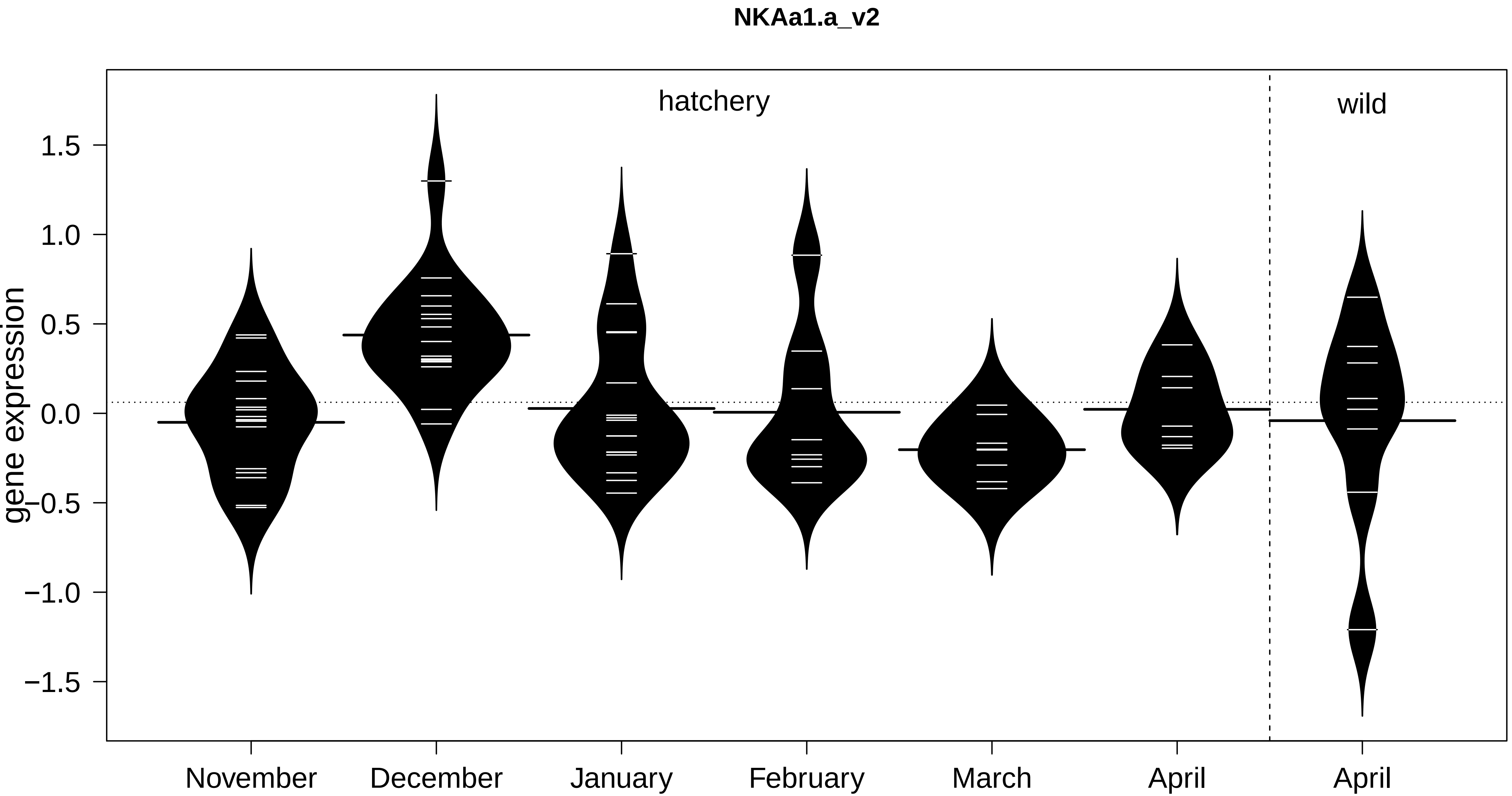

NKAa1.a_v2
hatchery
wild
gene expression
1.5
1.0
0.5
0.0
−0.5
−1.0
−1.5
November
December
January
February
March
April
April

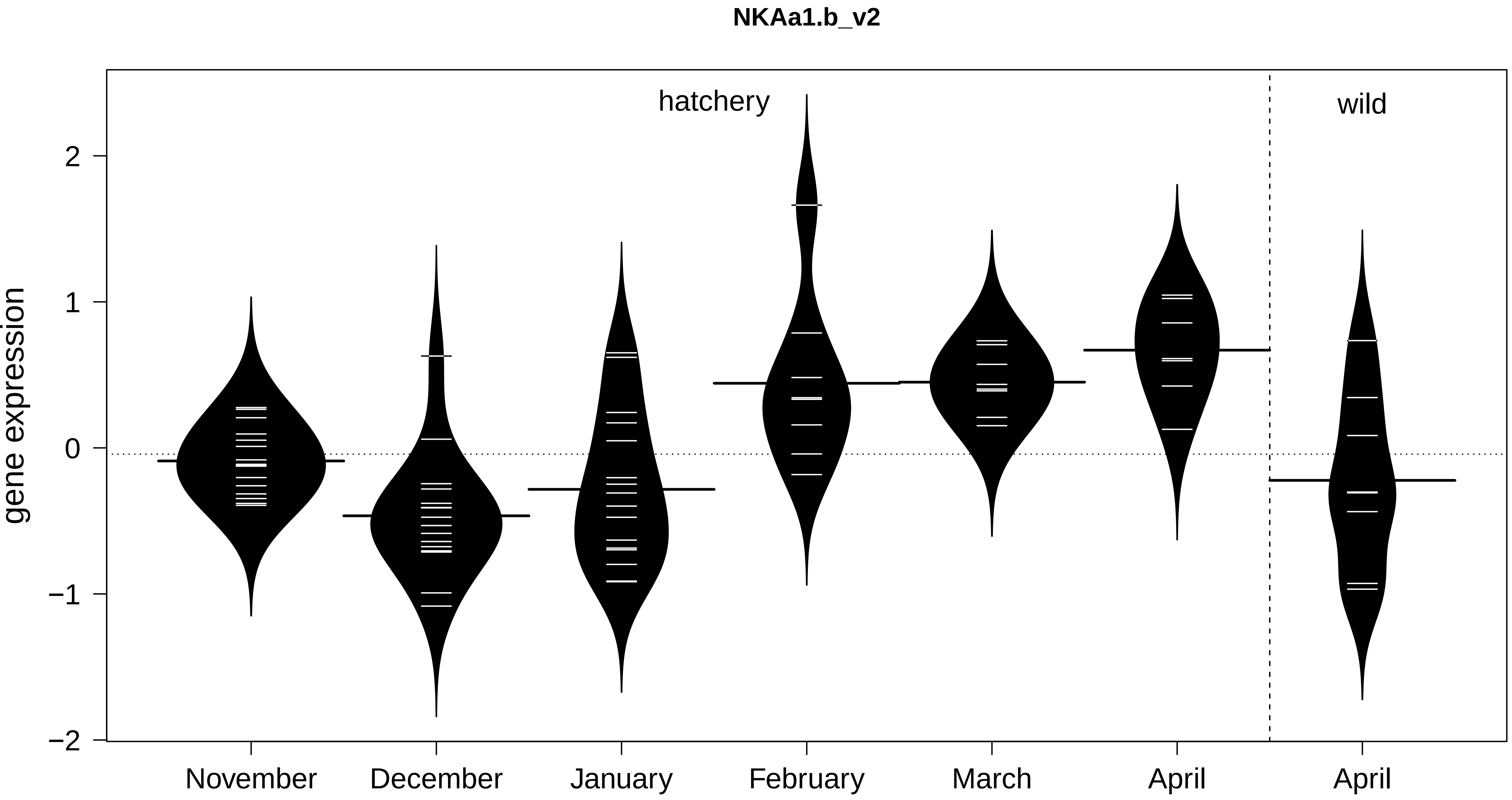

NKAa1.b_v2
hatchery
wild
gene expression
2
1
0
−1
−2
November
December
January
February
March
April
April

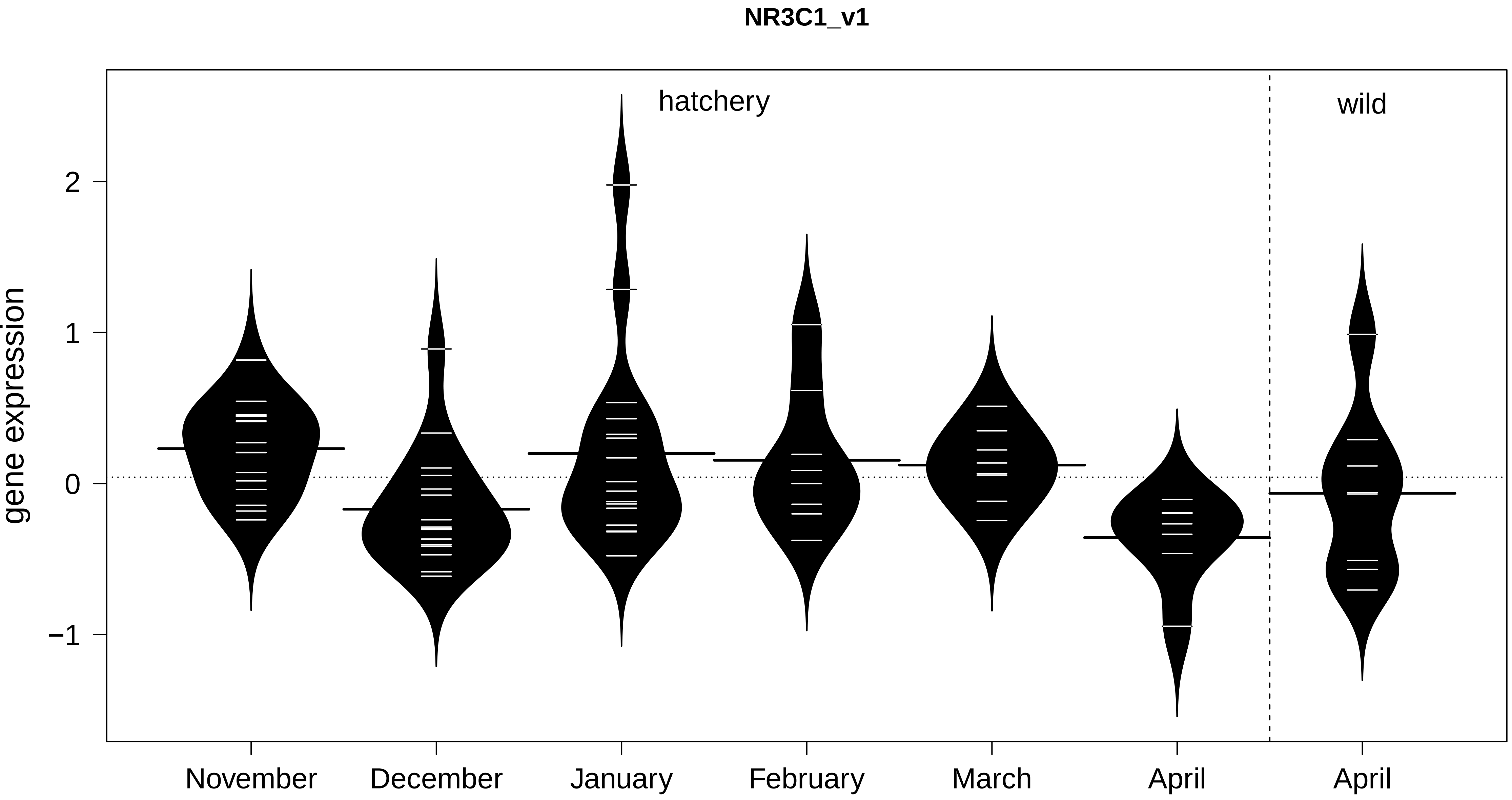

NR3C1_v1
hatchery
wild
2
1
0
−1
gene expression
November
December
January
February
March
April
April

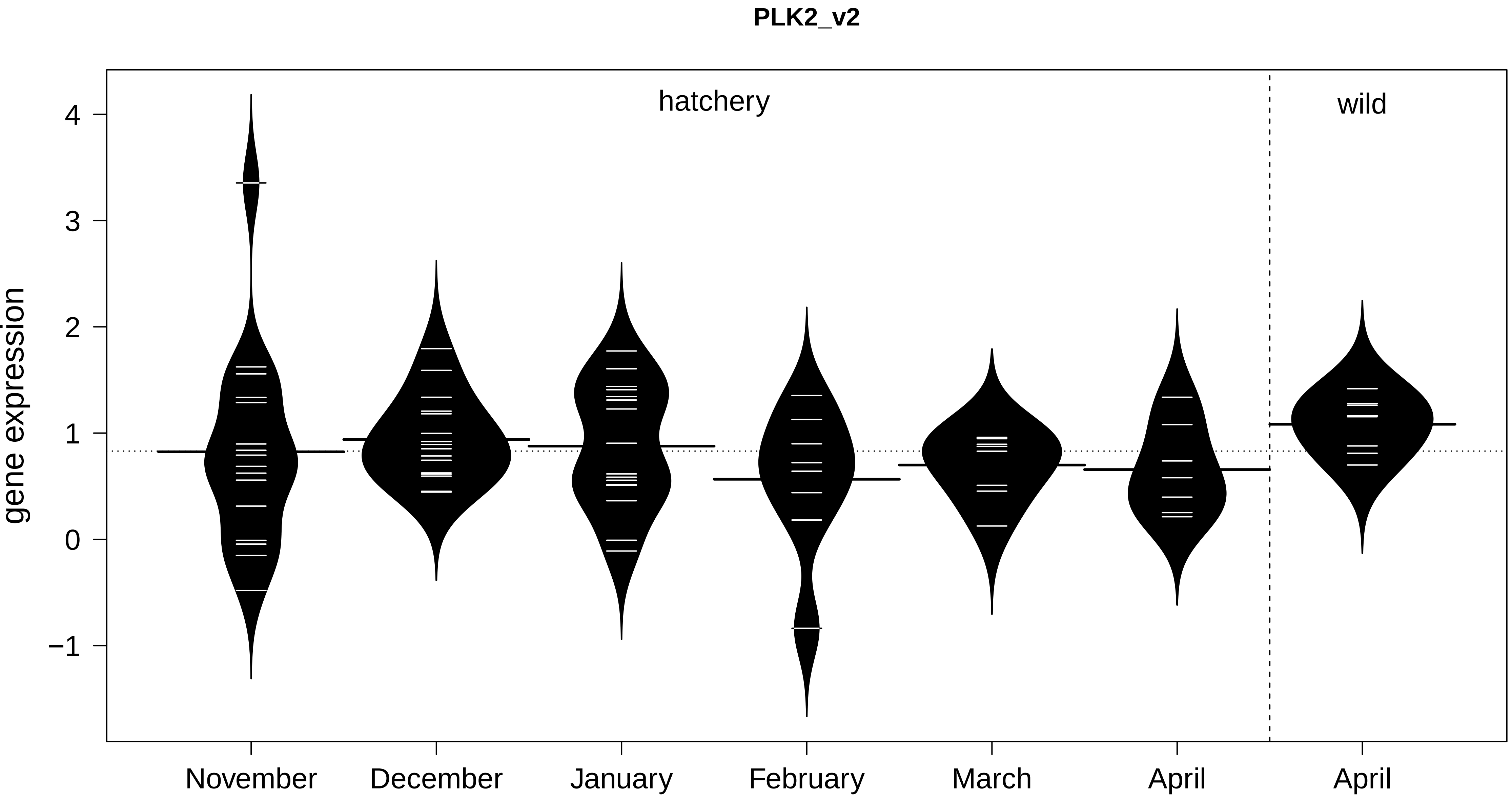
PLK2_v2
hatchery
wild
gene expression
4
3
2
1
0
−1
November
December
January
February
March
April
April

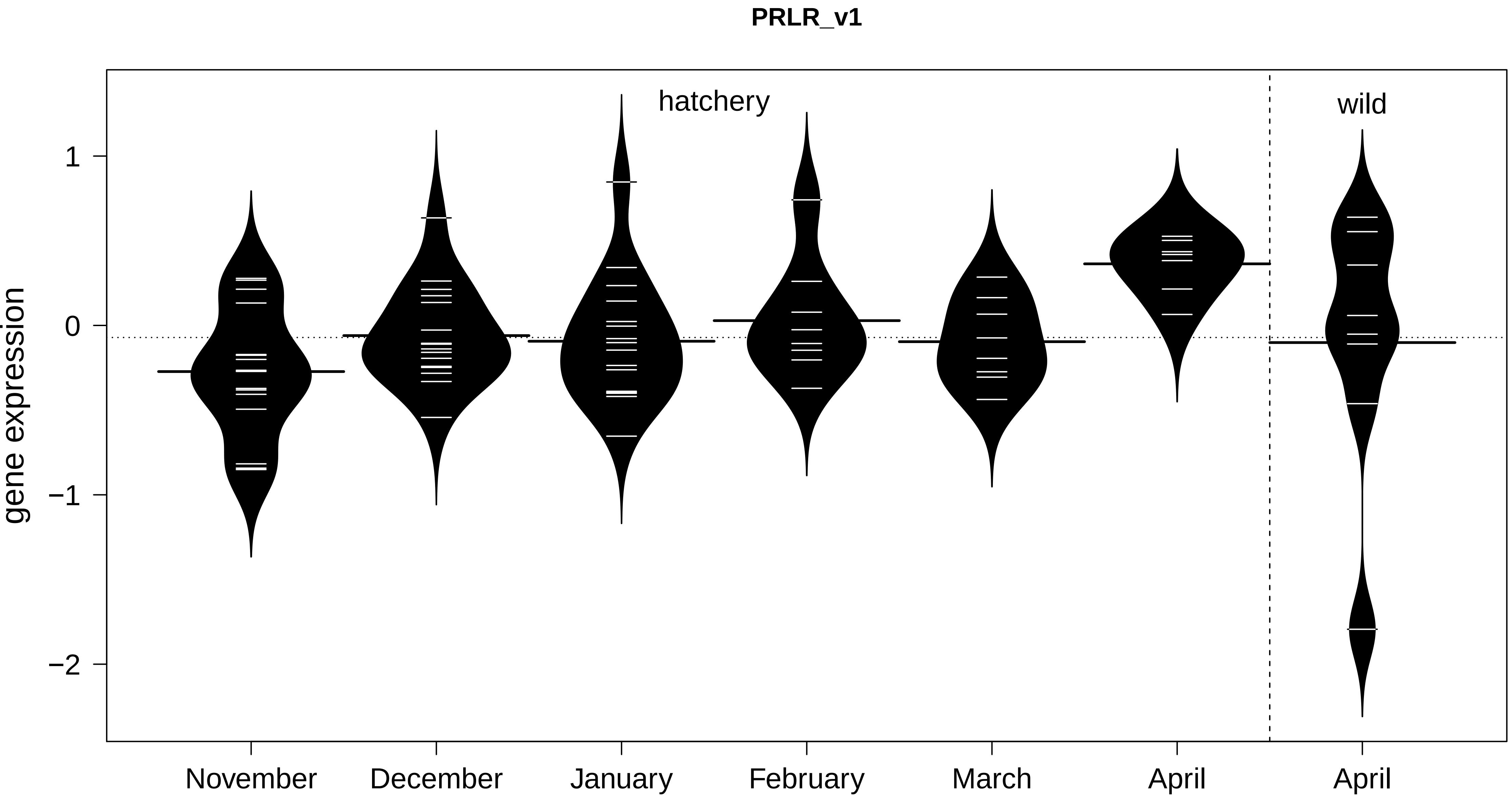
PRLR_v1
hatchery
wild
gene expression
1
0
−1
−2
November
December
January
February
March
April
April

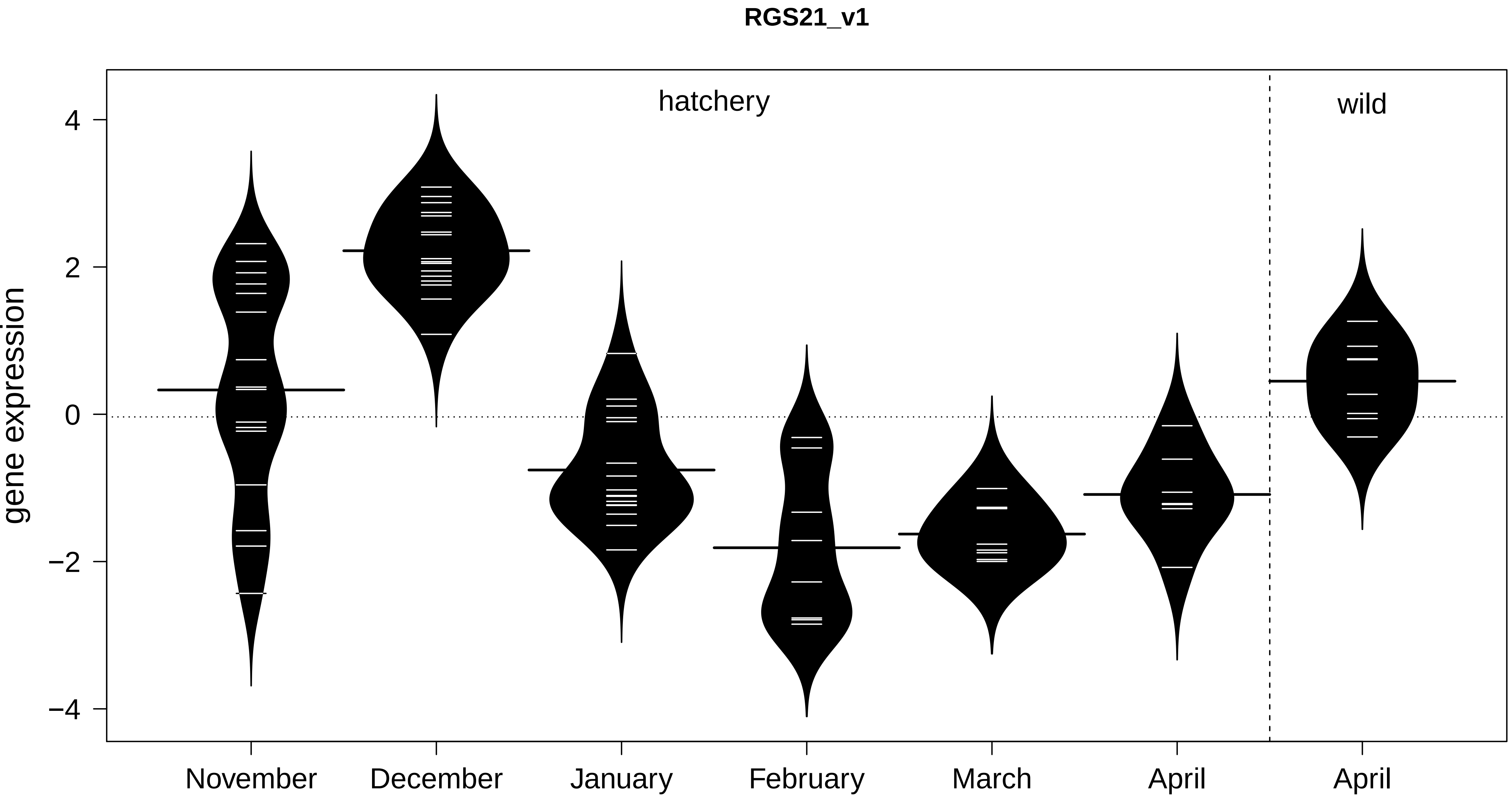
RGS21_v1
hatchery
wild
gene expression
4
2
0
−2
−4
November
December
January
February
March
April
April

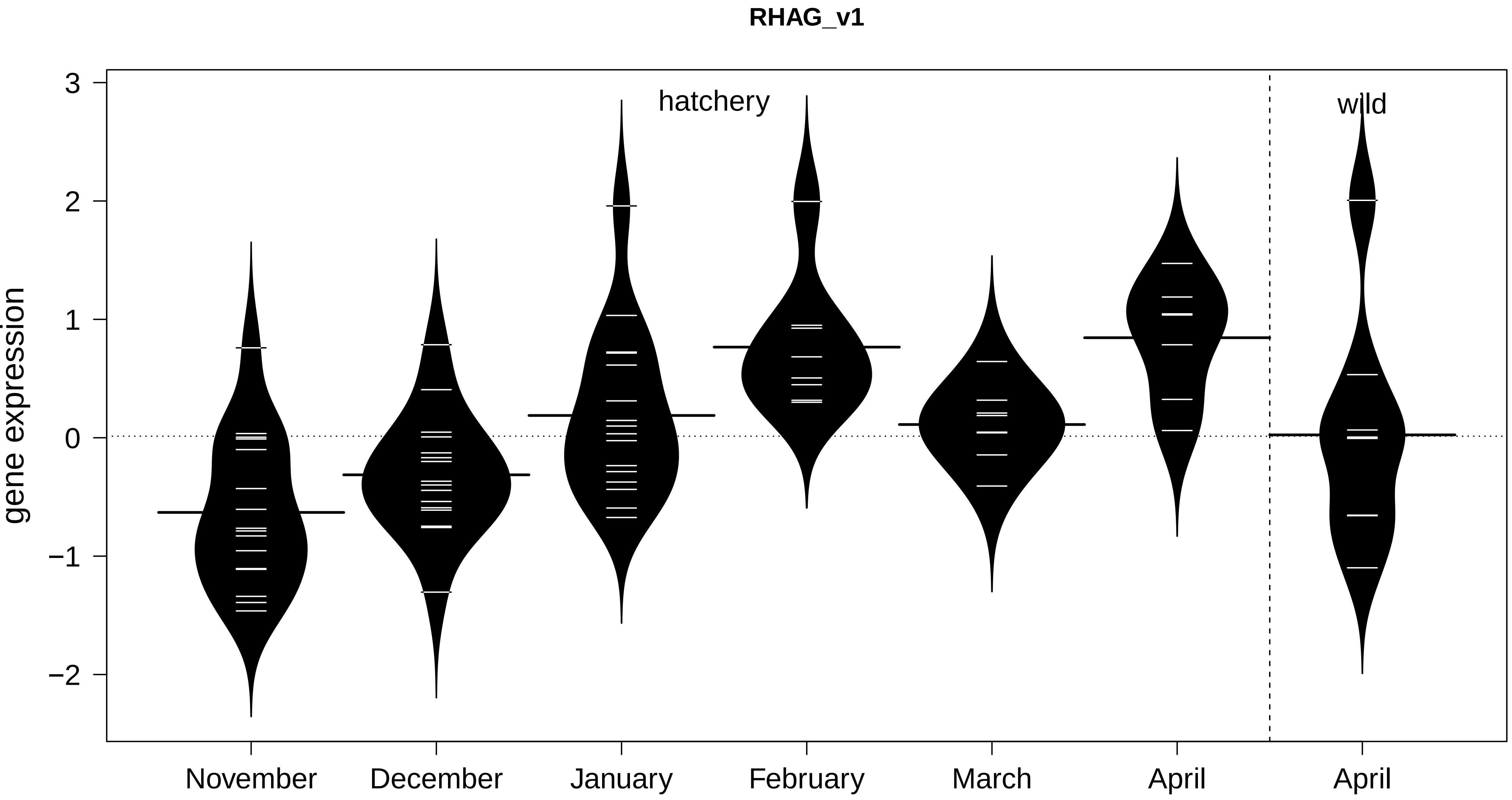

RHAG_v1
hatchery
wild
gene expression
3
2
1
0
−1
−2
November
December
January
February
March
April
April

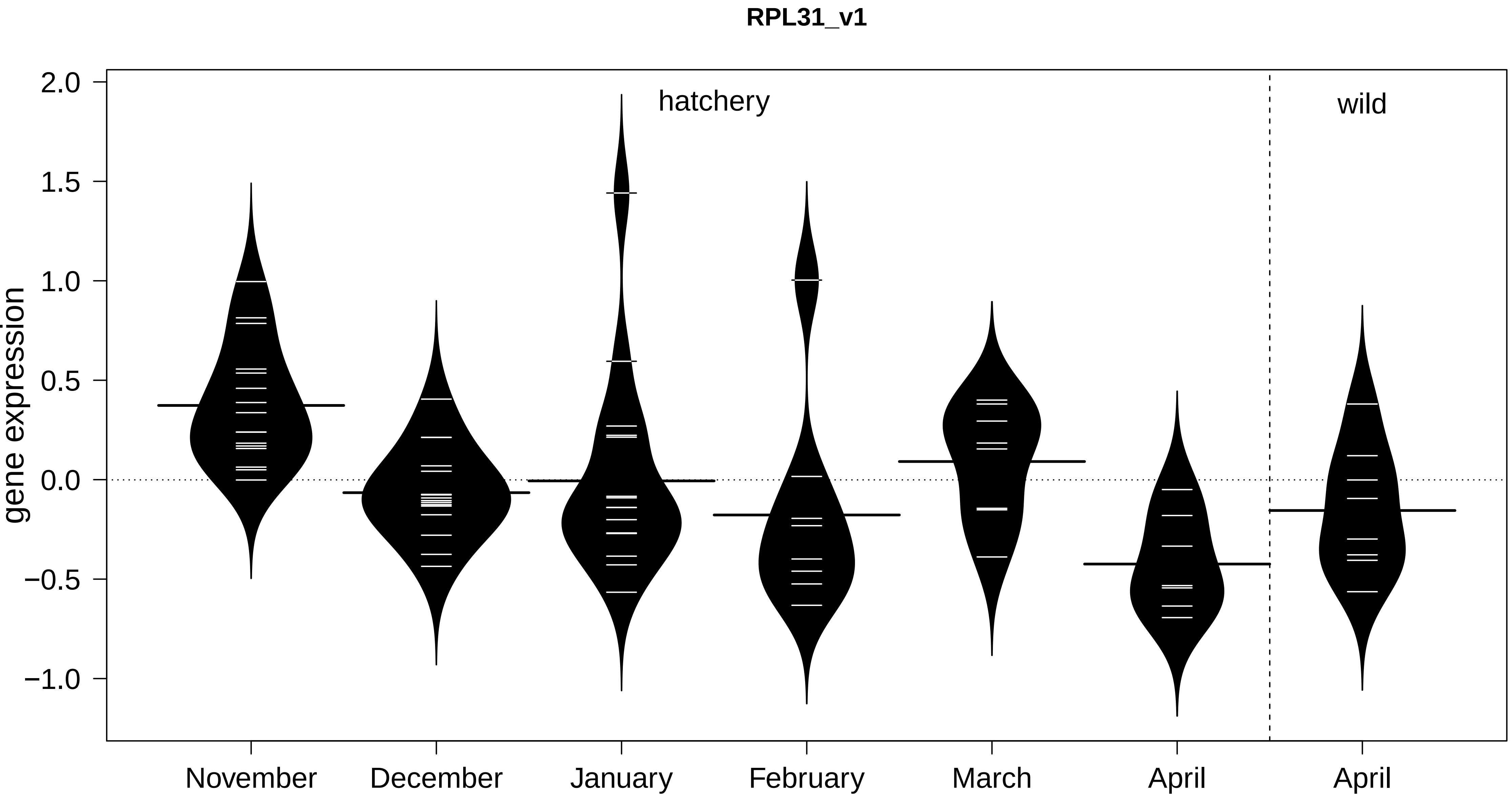
RPL31_v1
hatchery
wild
gene expression
2.0
1.5
1.0
0.5
0.0
−0.5
−1.0
November
December
January
February
March
April
April

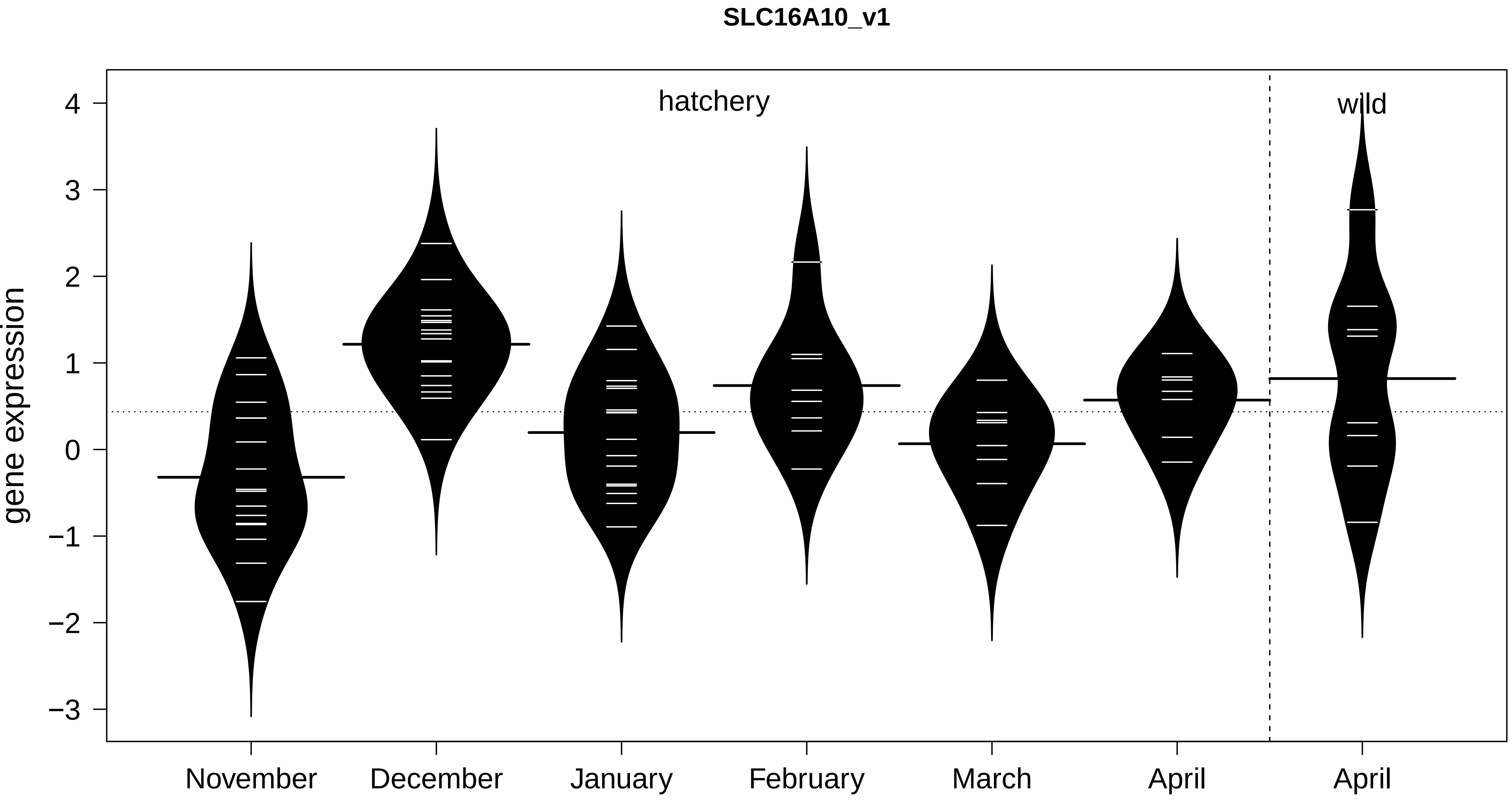
SLC16A10_v1
hatchery
wild
gene expression
4
3
2
1
0
−1
−2
−3
November
December
January
February
March
April
April

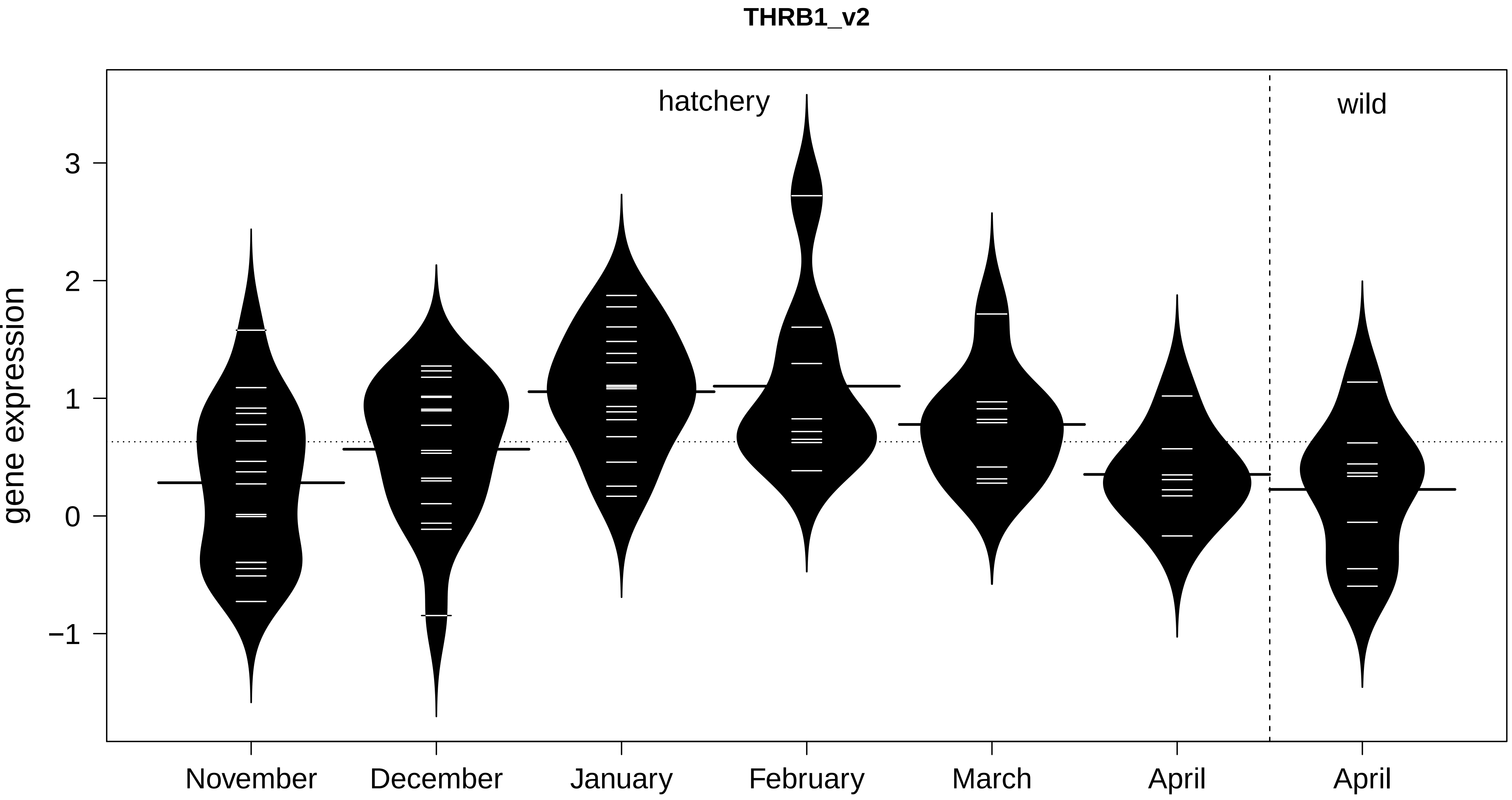
THRB1_v2
hatchery
wild
gene expression
3
2
1
0
−1
November
December
January
February
March
April
April

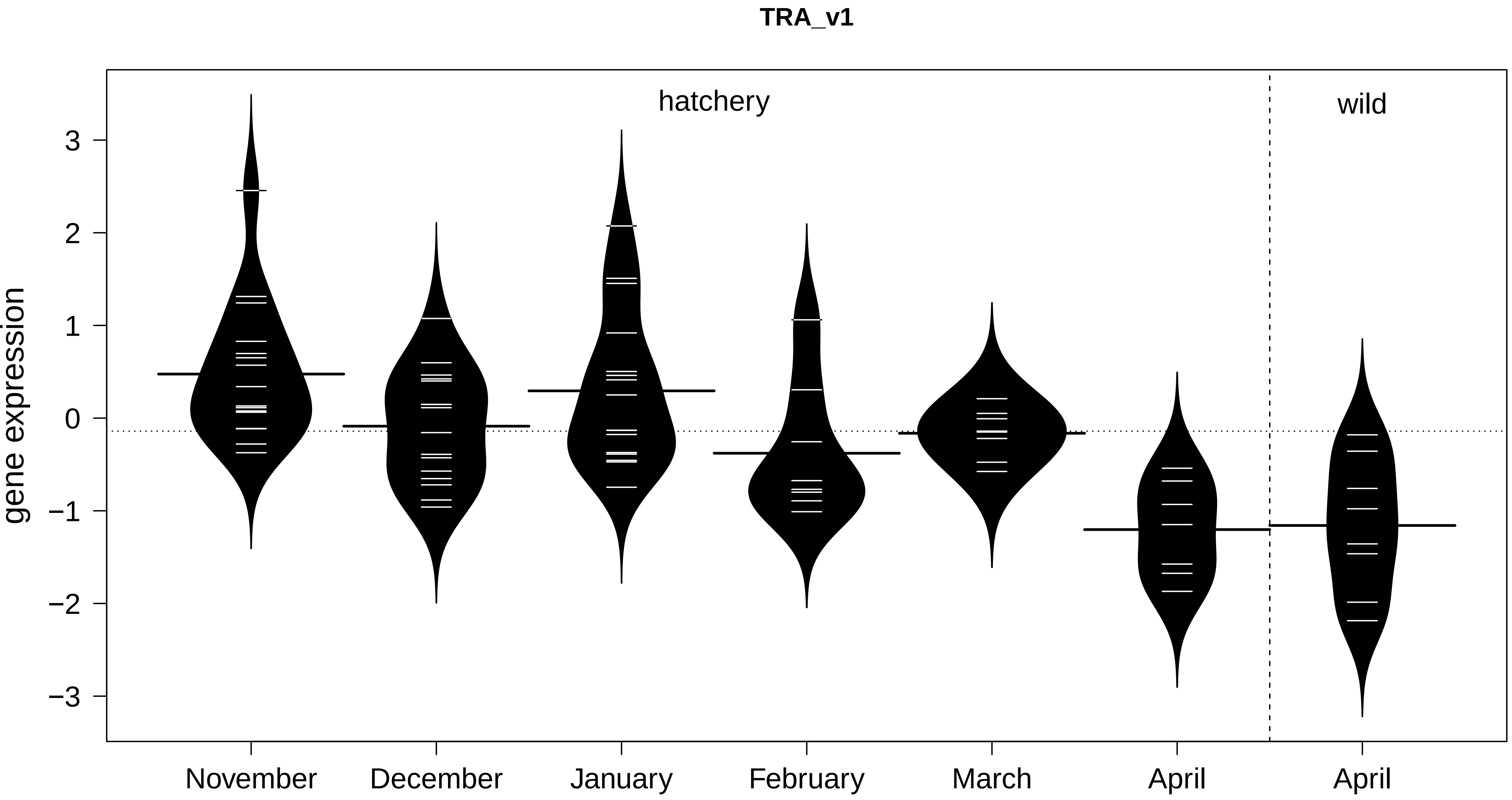
TRA_v1
hatchery
wild
3
2
1
0
−1
−2
−3
gene expression
November
December
January
February
March
April
April

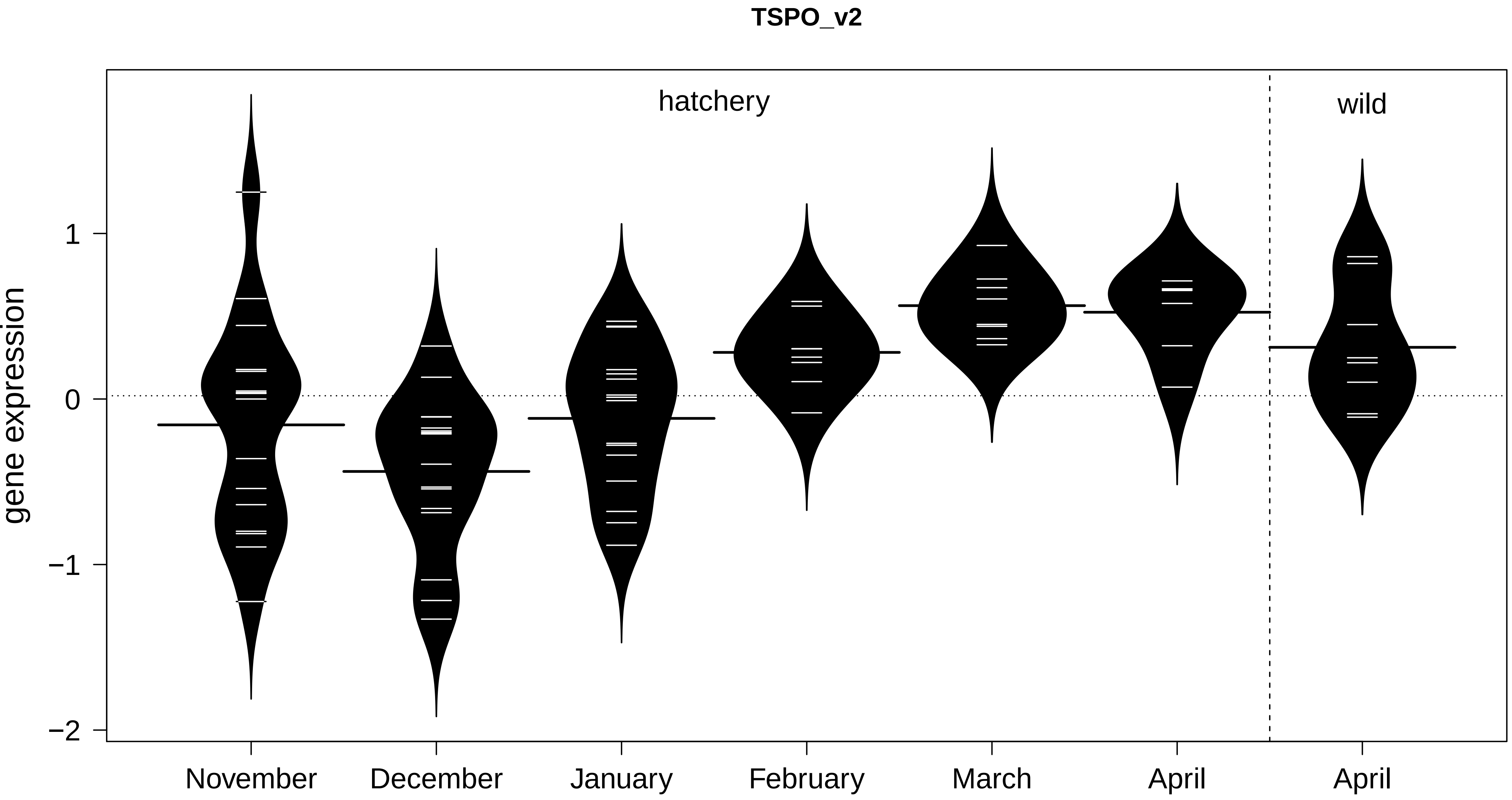

TSPO_v2
hatchery
wild
gene expression
1
0
−1
−2
November
December
January
February
March
April
April

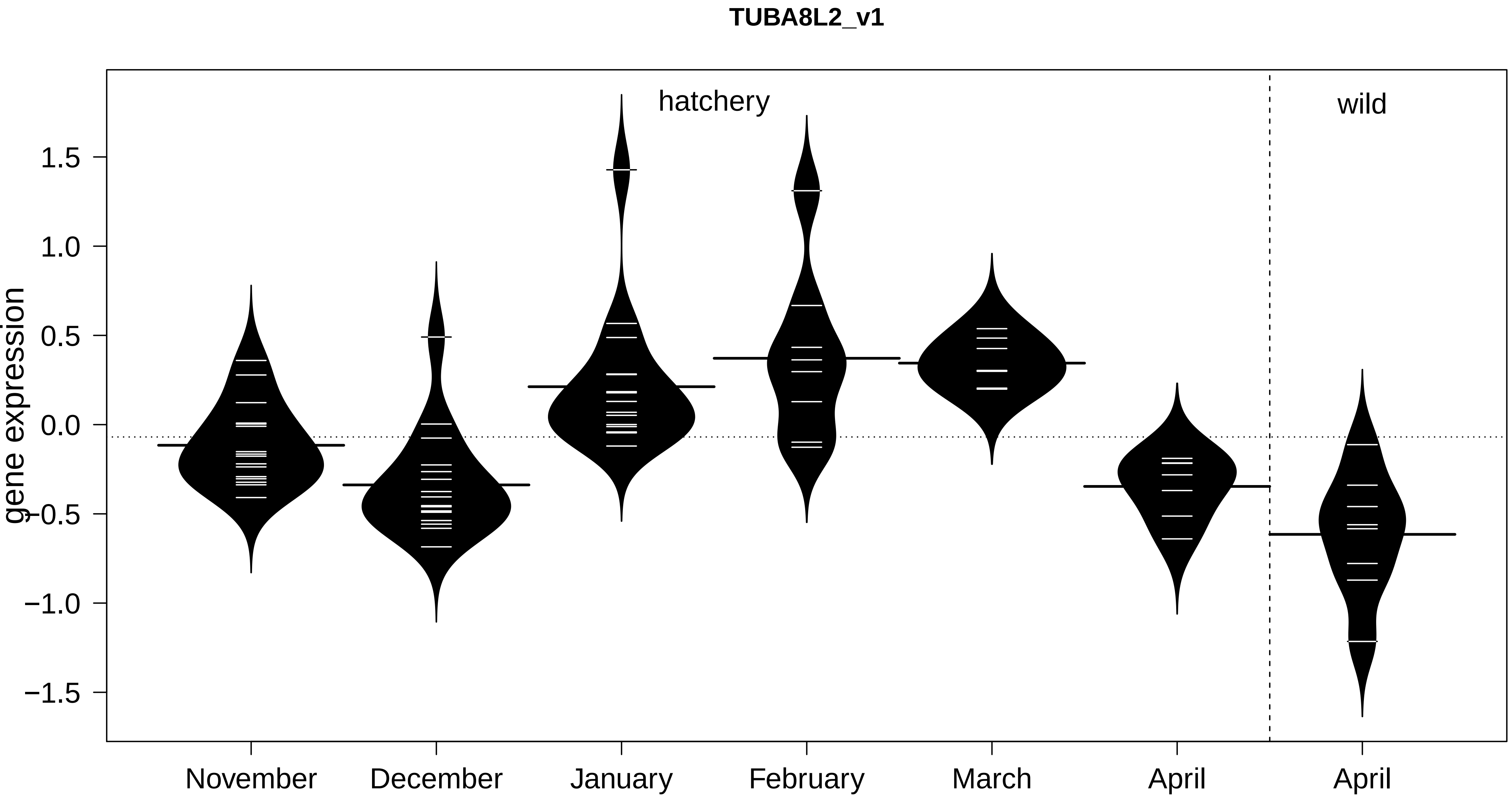

TUBA8L2_v1
hatchery
wild
gene expression
1.5
1.0
0.5
0.0
−0.5
−1.0
−1.5
November
December
January
February
March
April
April

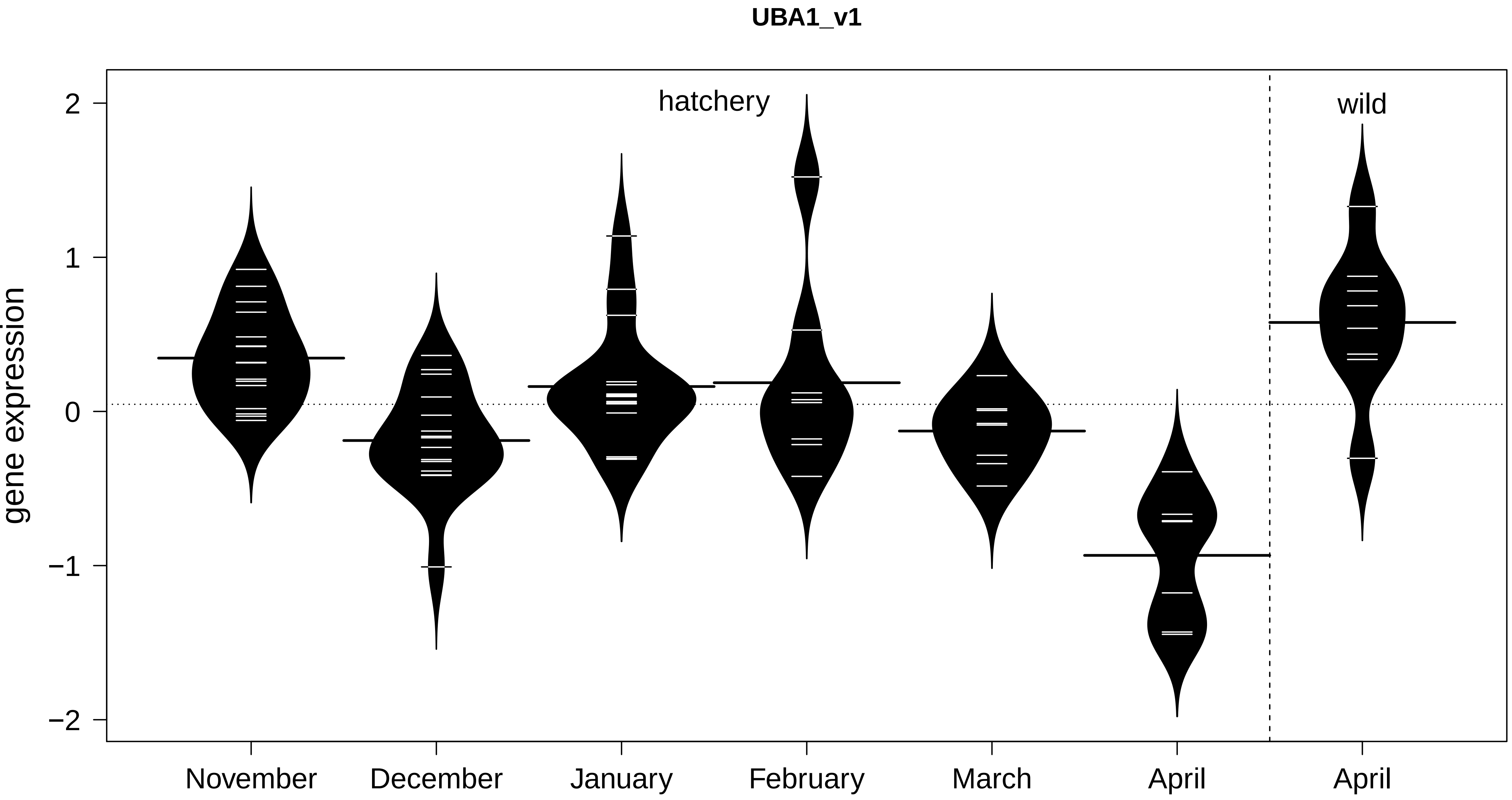

UBA1_v1
hatchery
wild
gene expression
2
1
0
−1
−2
November
December
January
February
March
April
April

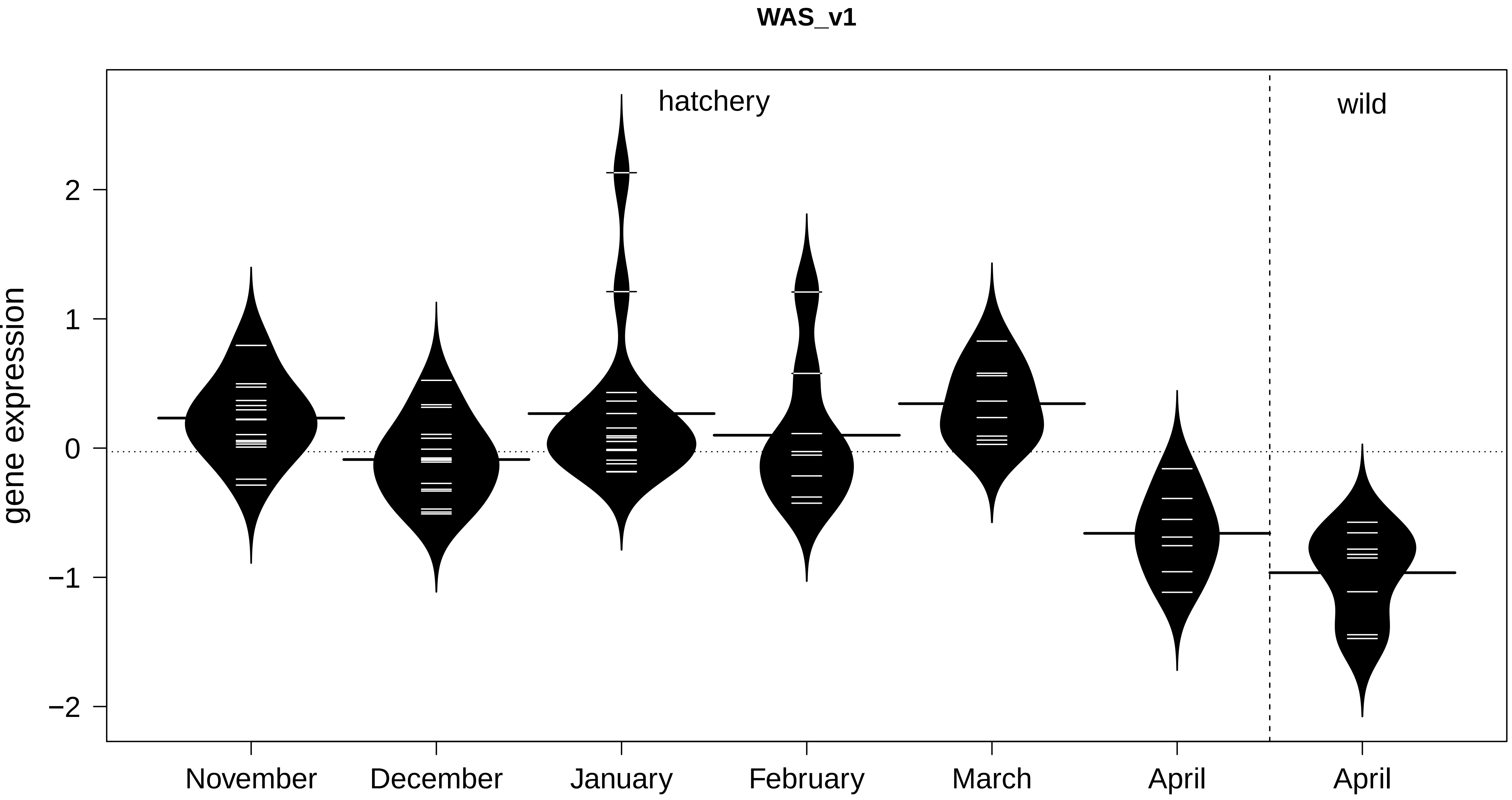

WAS_v1
hatchery
wild
gene expression
2
1
0
−1
−2
November
December
January
February
March
April
April
