## appendix 3 for "Discovery and validation of candidate smoltification gene expression biomarkers across multiple species and ecotypes of Pacific salmonids"

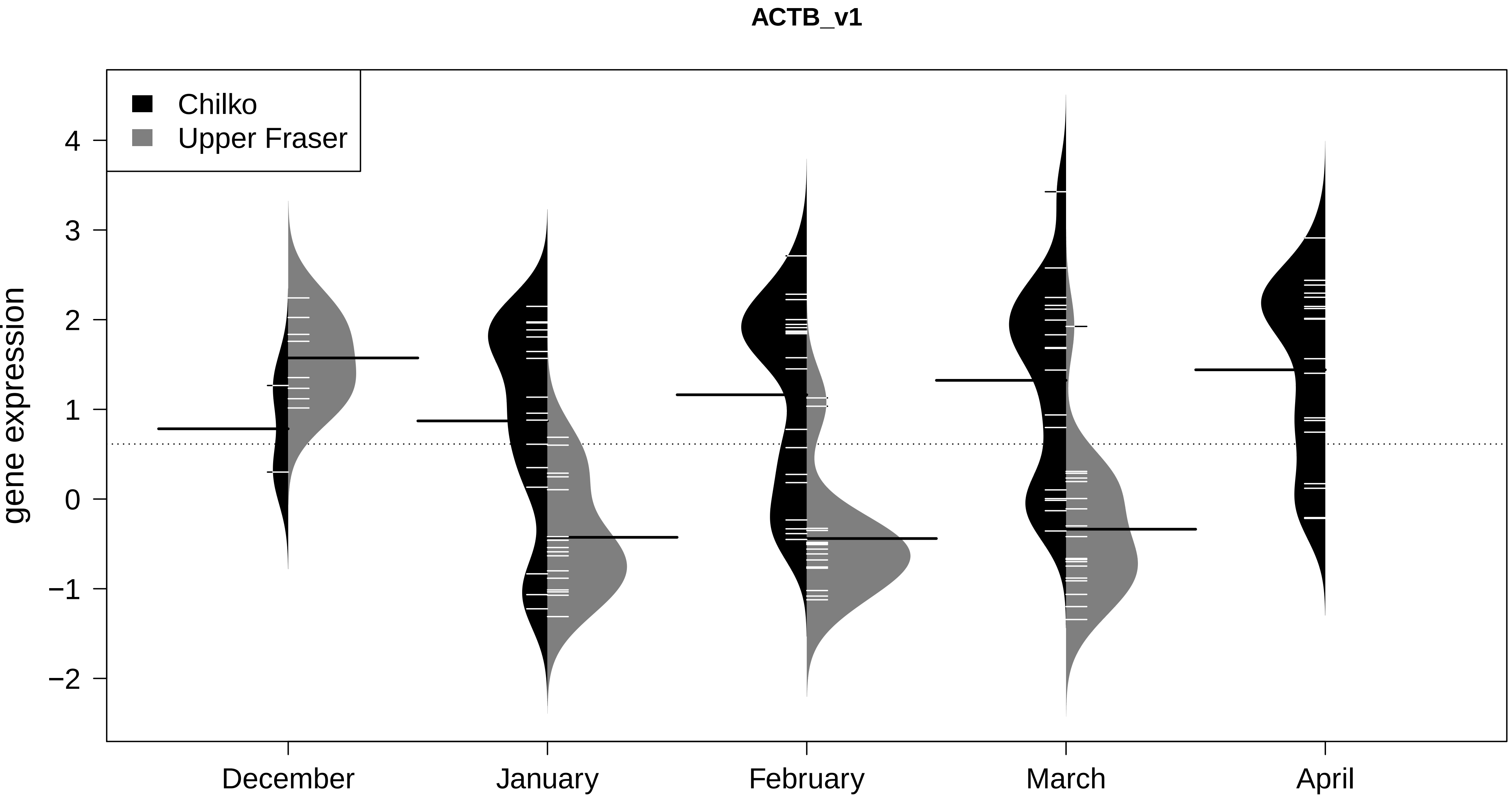
ACTB_v1
Chilko
Upper Fraser
gene expression
4
3
2
1
0
−1
−2
December
January
February
March
April

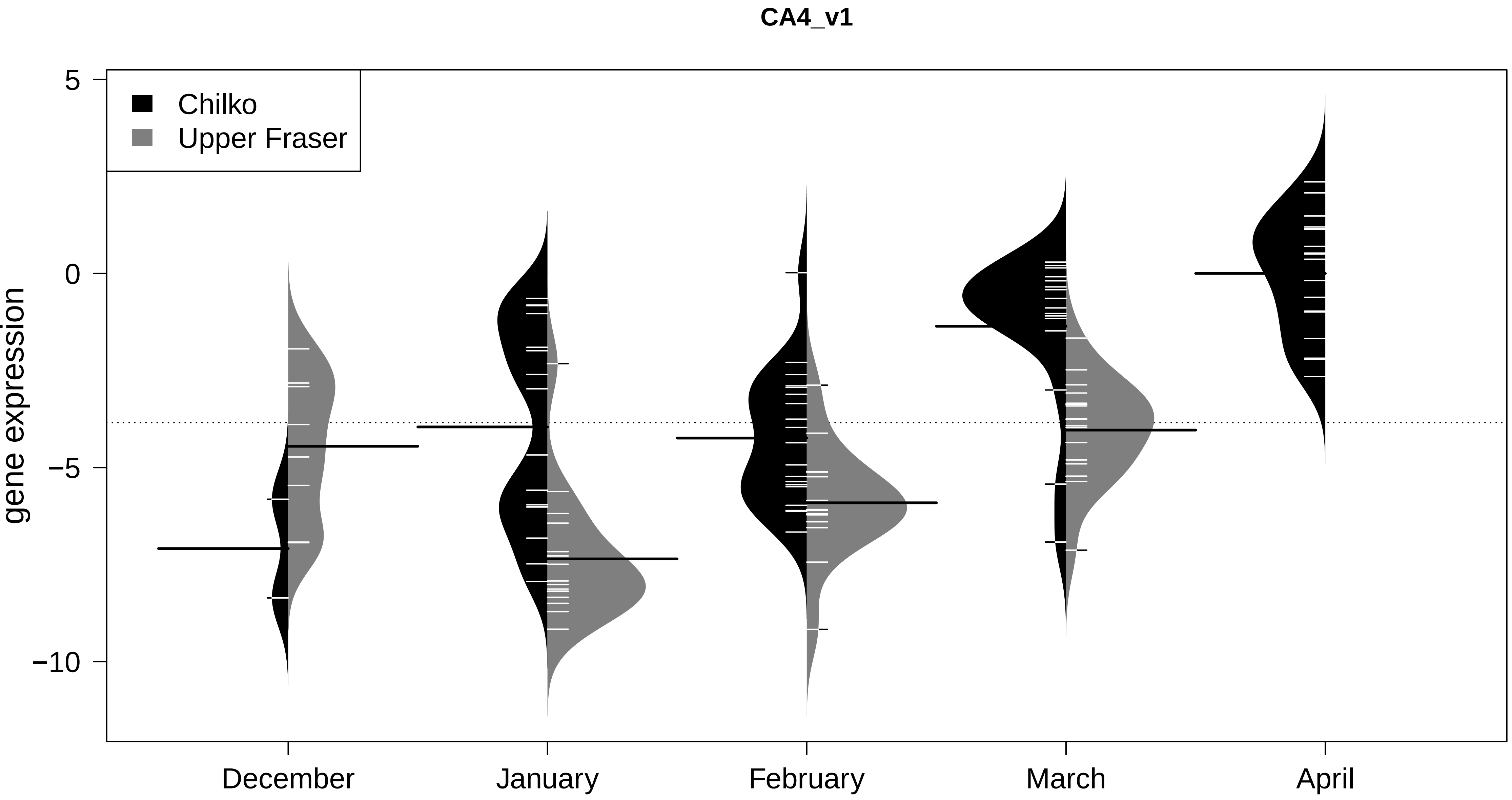
CA4_v1
Chilko
Upper Fraser
gene expression
5
0
−5
−10
December
January
February
March
April

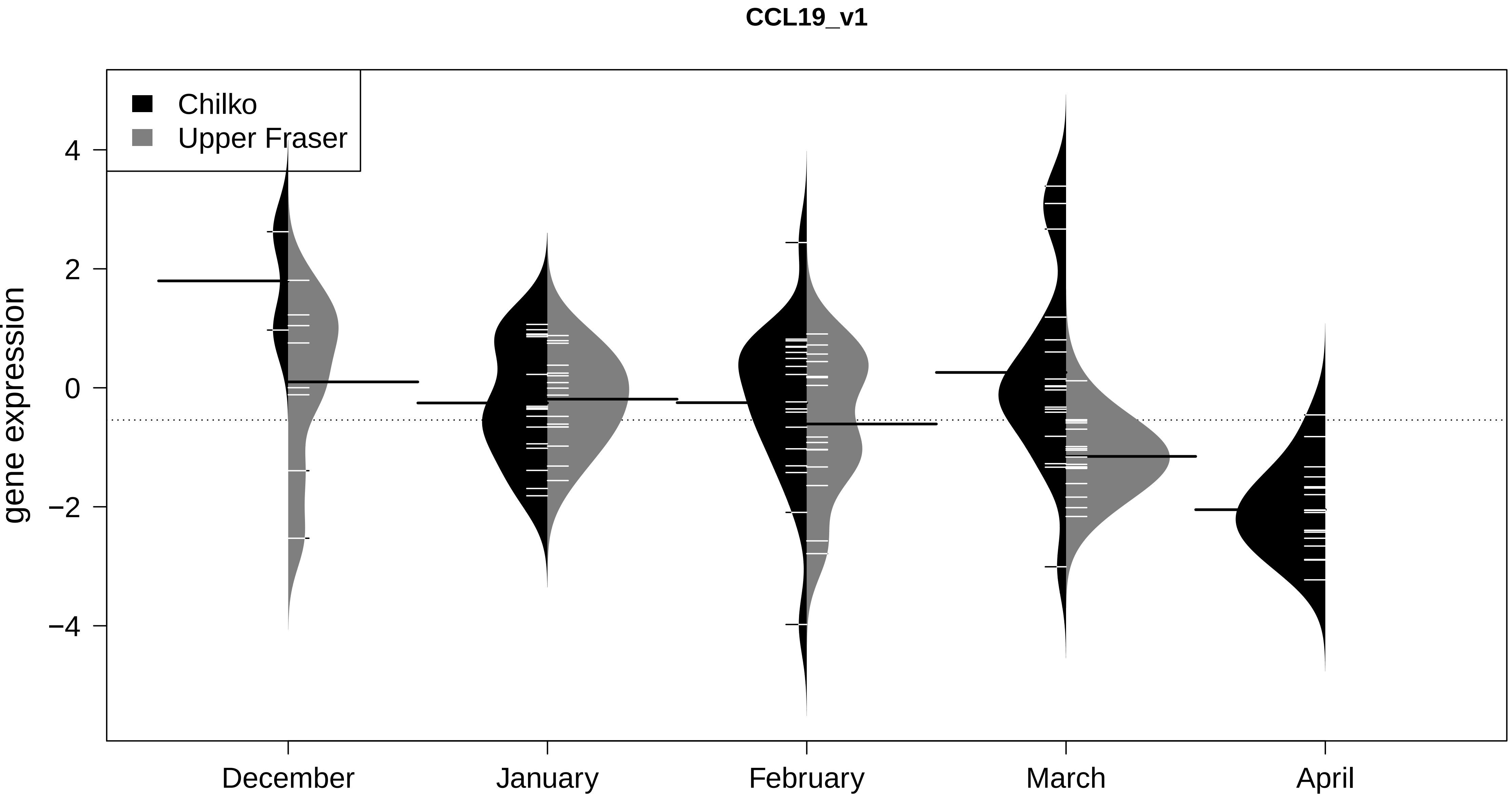

CCL19_v1
Chilko
Upper Fraser
gene expression
4
2
0
−2
−4
December
January
February
March
April

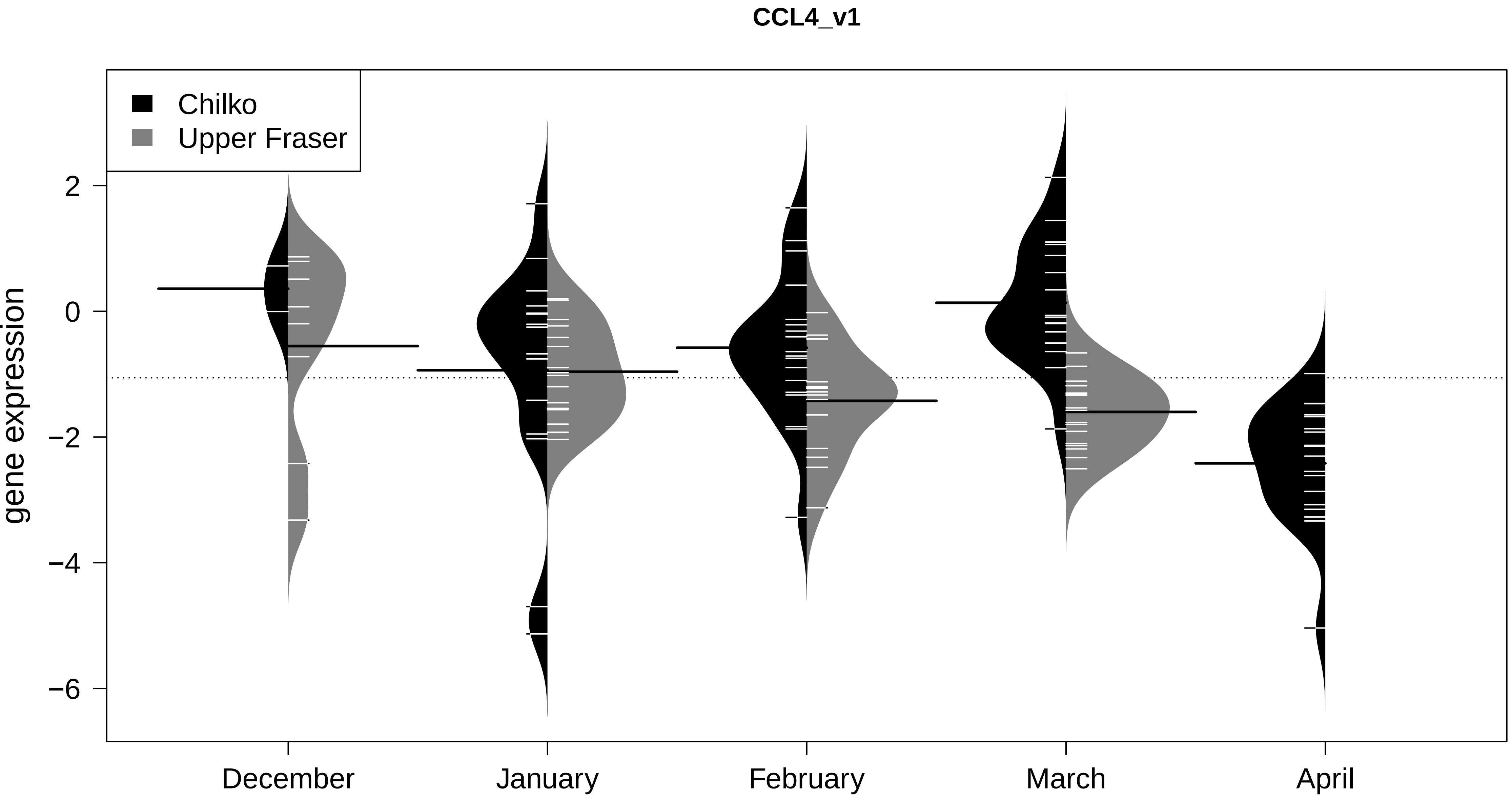
CCL4_v1
Chilko
Upper Fraser
gene expression
2
0
−2
−4
−6
December
January
February
March
April

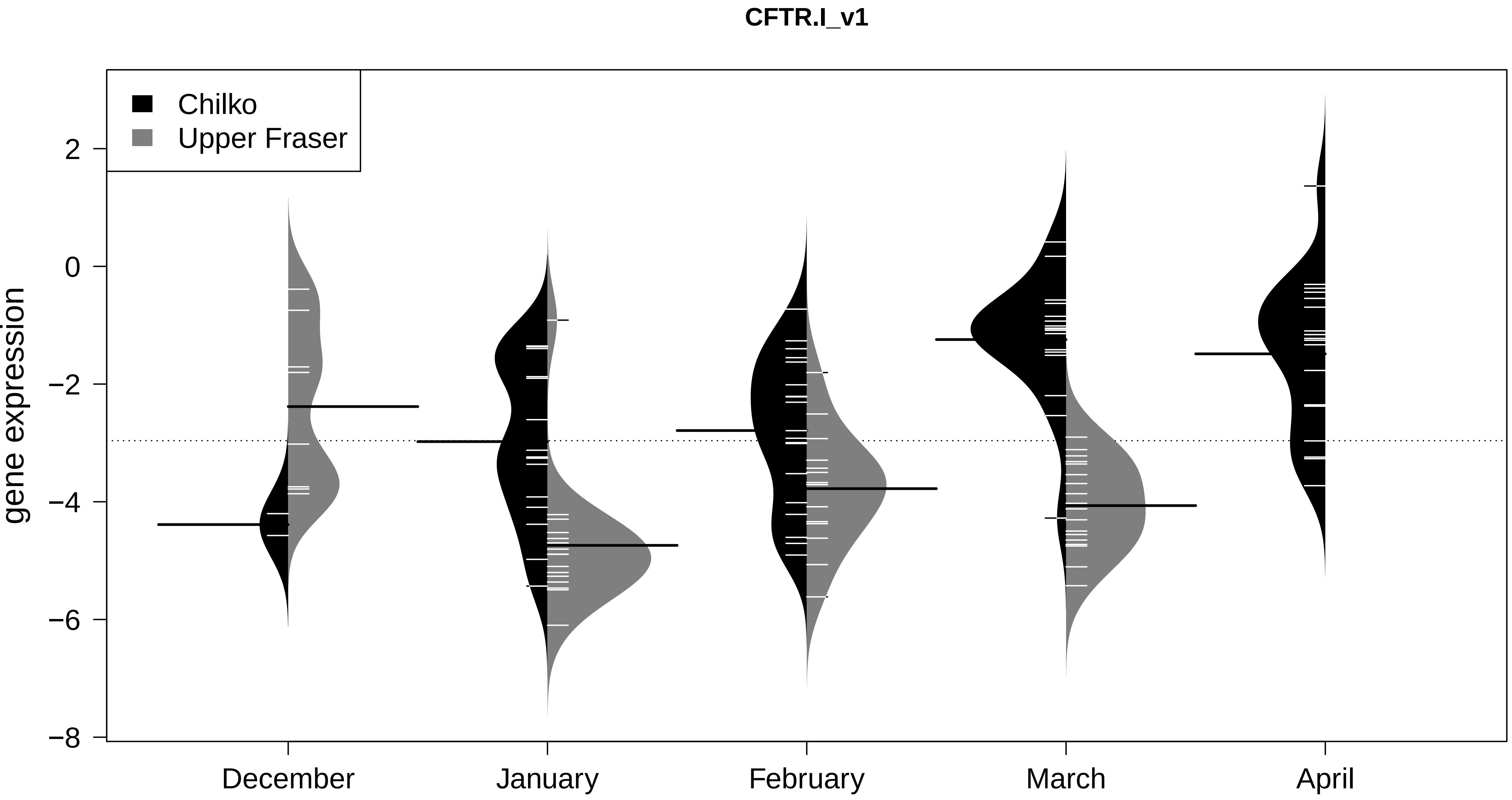
CFTR.I_v1
Chilko
Upper Fraser
gene expression
2
0
−2
−4
−6
−8
December
January
February
March
April

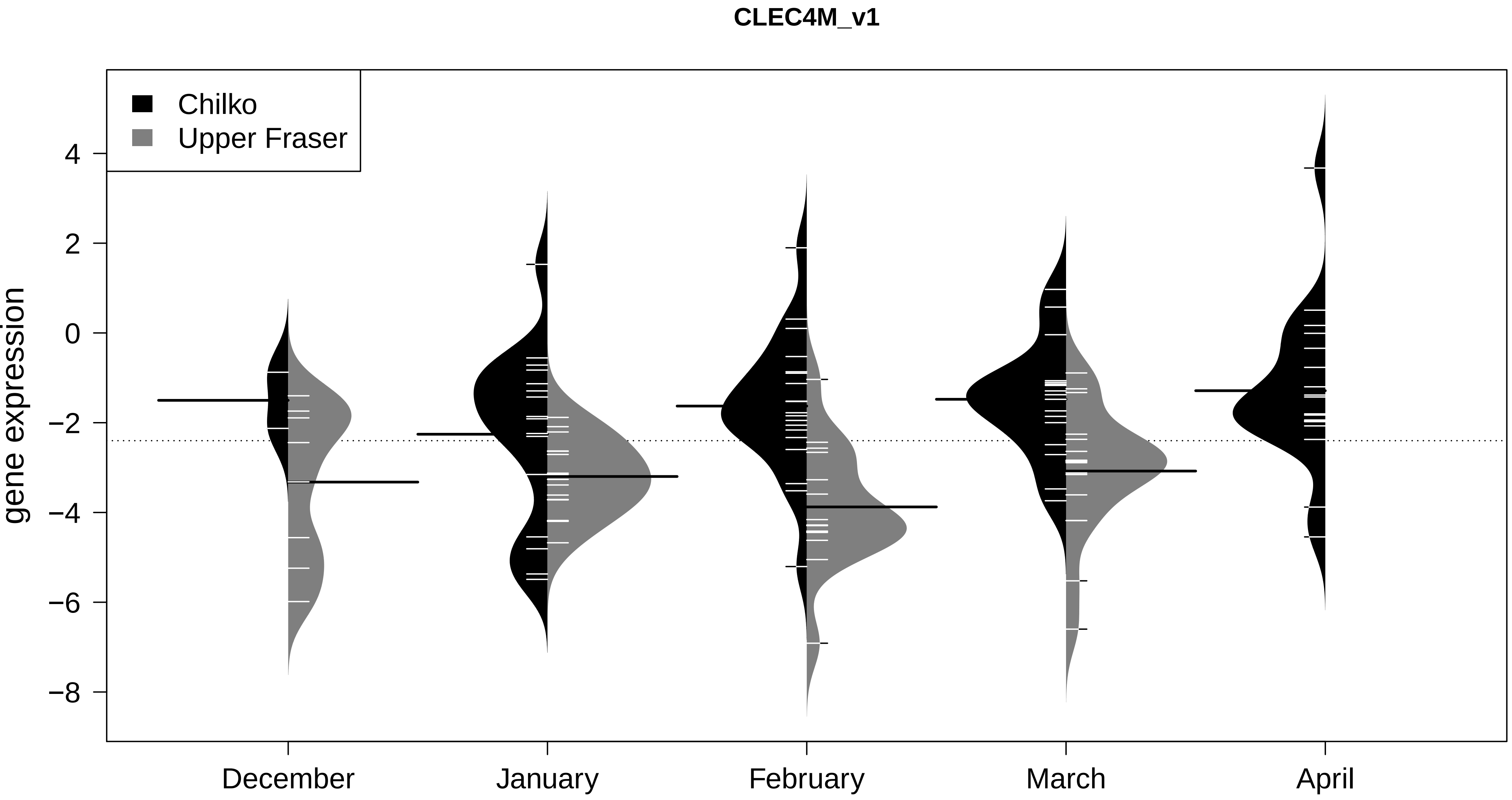

CLEC4M_v1
Chilko
Upper Fraser
gene expression
4
2
0
−2
−4
−6
−8
December
January
February
March
April

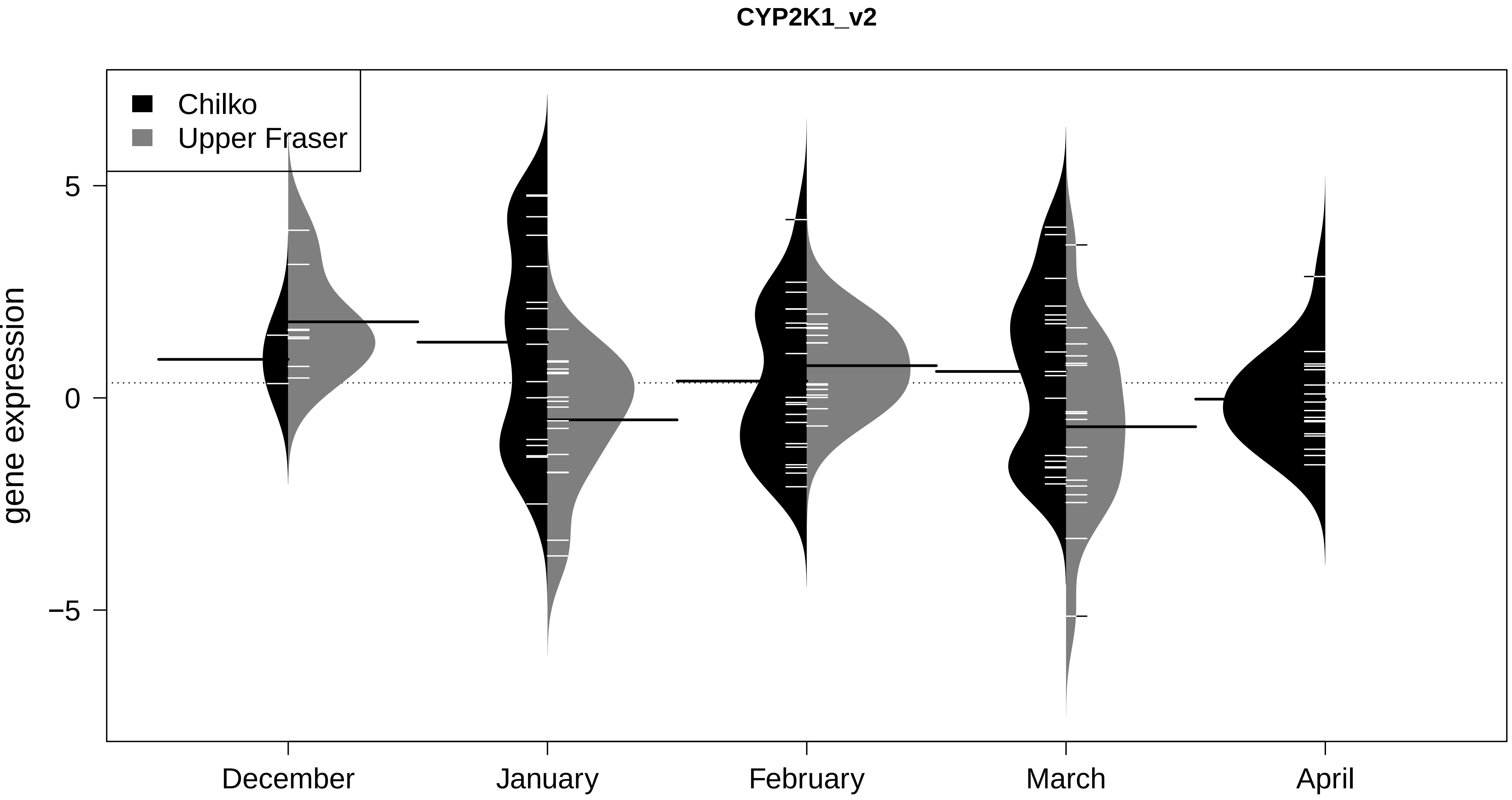
CYP2K1_v2
Chilko
Upper Fraser
gene expression
5
0
−5
December
January
February
March
April

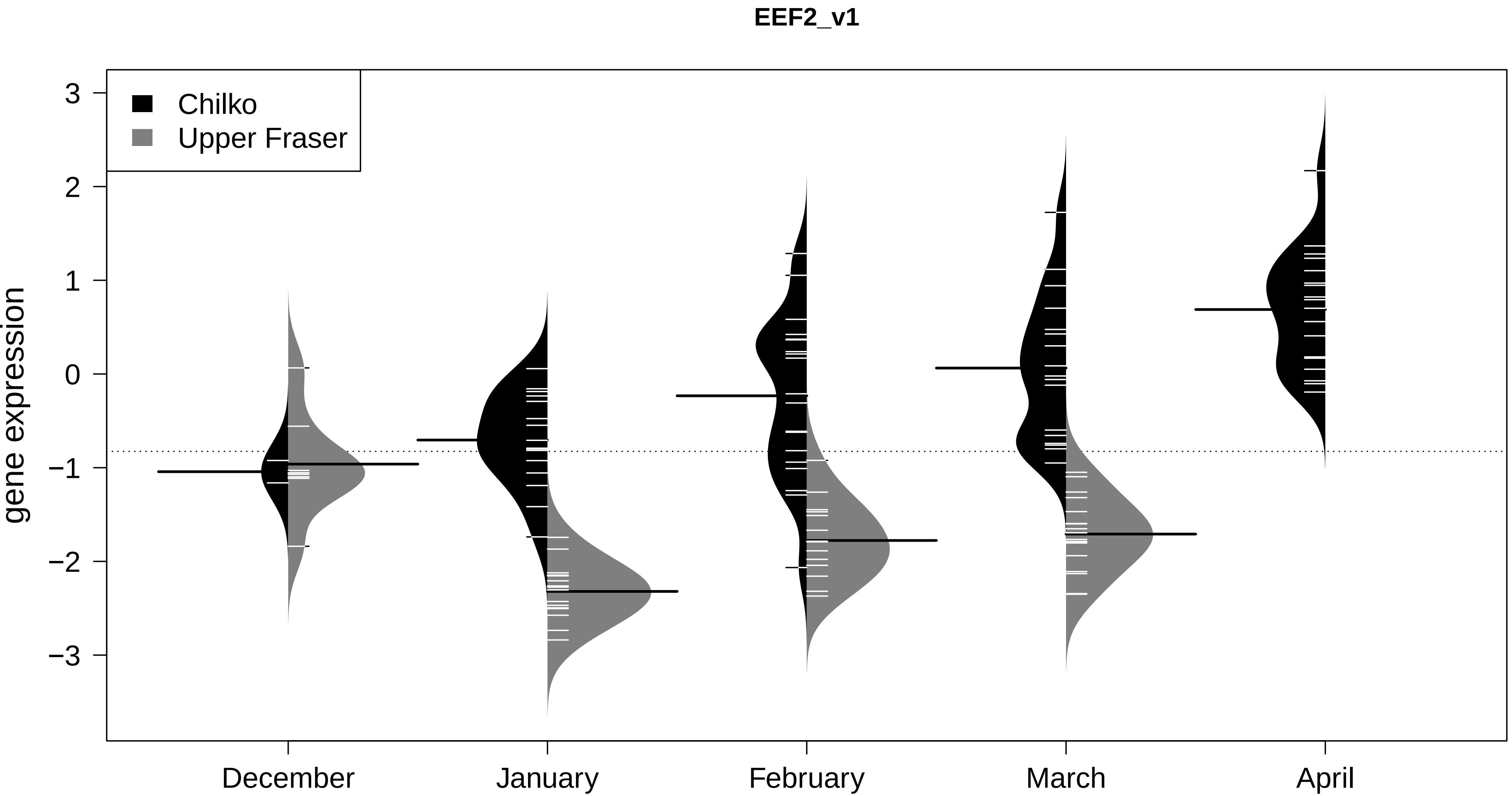
EEF2_v1
Chilko
Upper Fraser
gene expression
3
2
1
0
−1
−2
−3
December
January
February
March
April

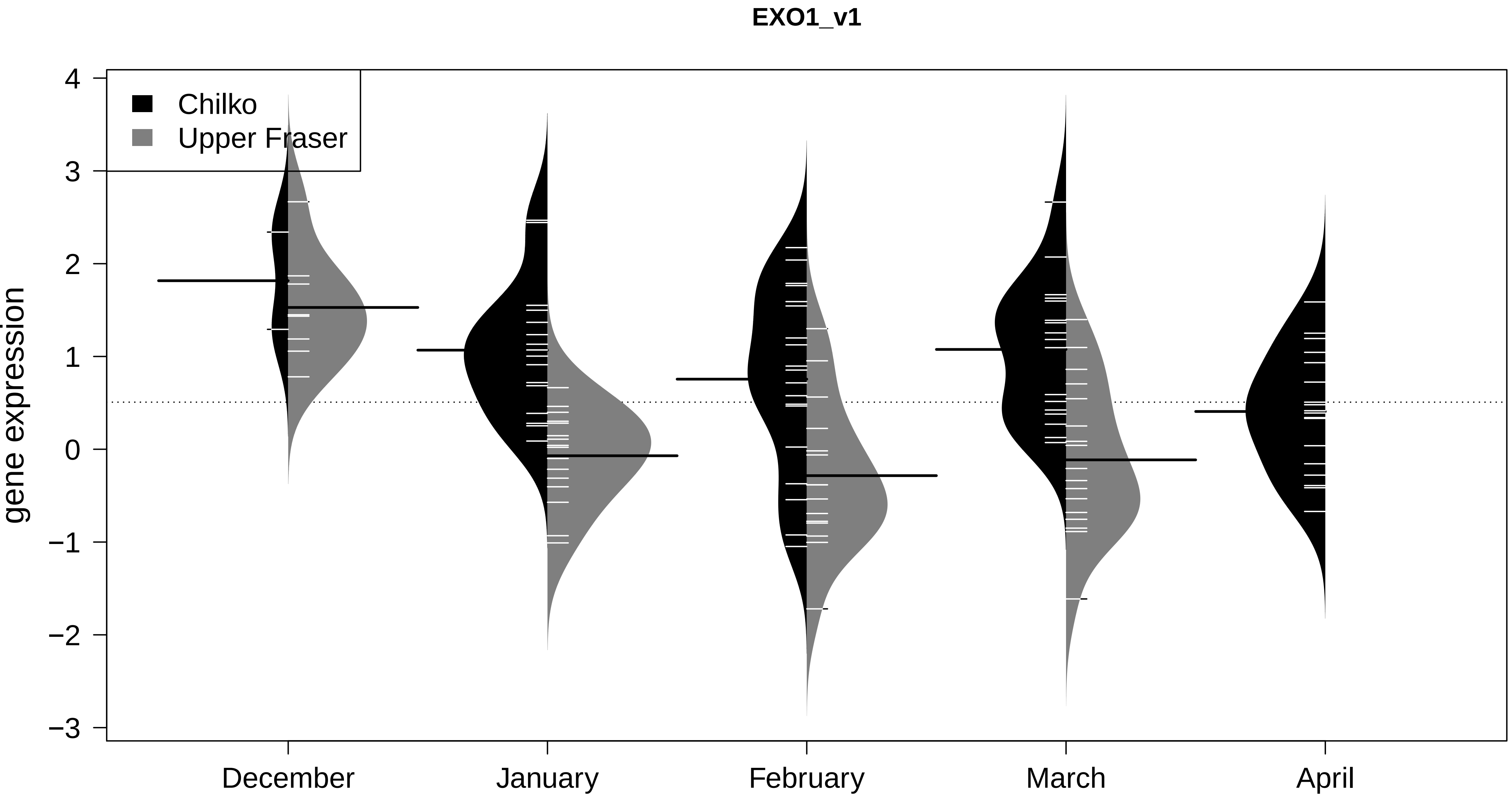

EXO1_v1
Chilko
Upper Fraser
gene expression
4
3
2
1
0
−1
−2
−3
December
January
February
March
April

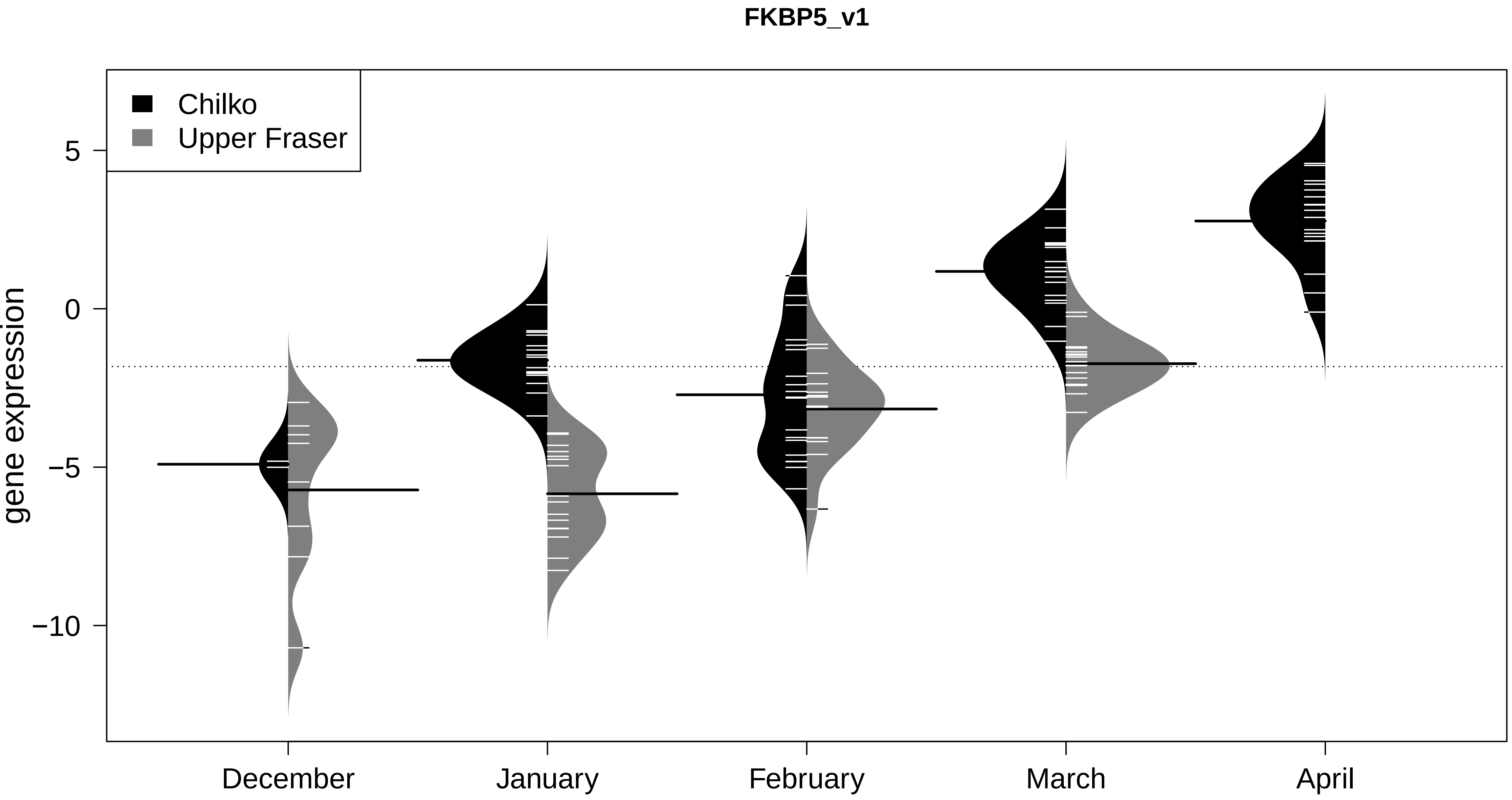

FKBP5_v1
Chilko
Upper Fraser
gene expression
5
0
−5
−10
December
January
February
March
April

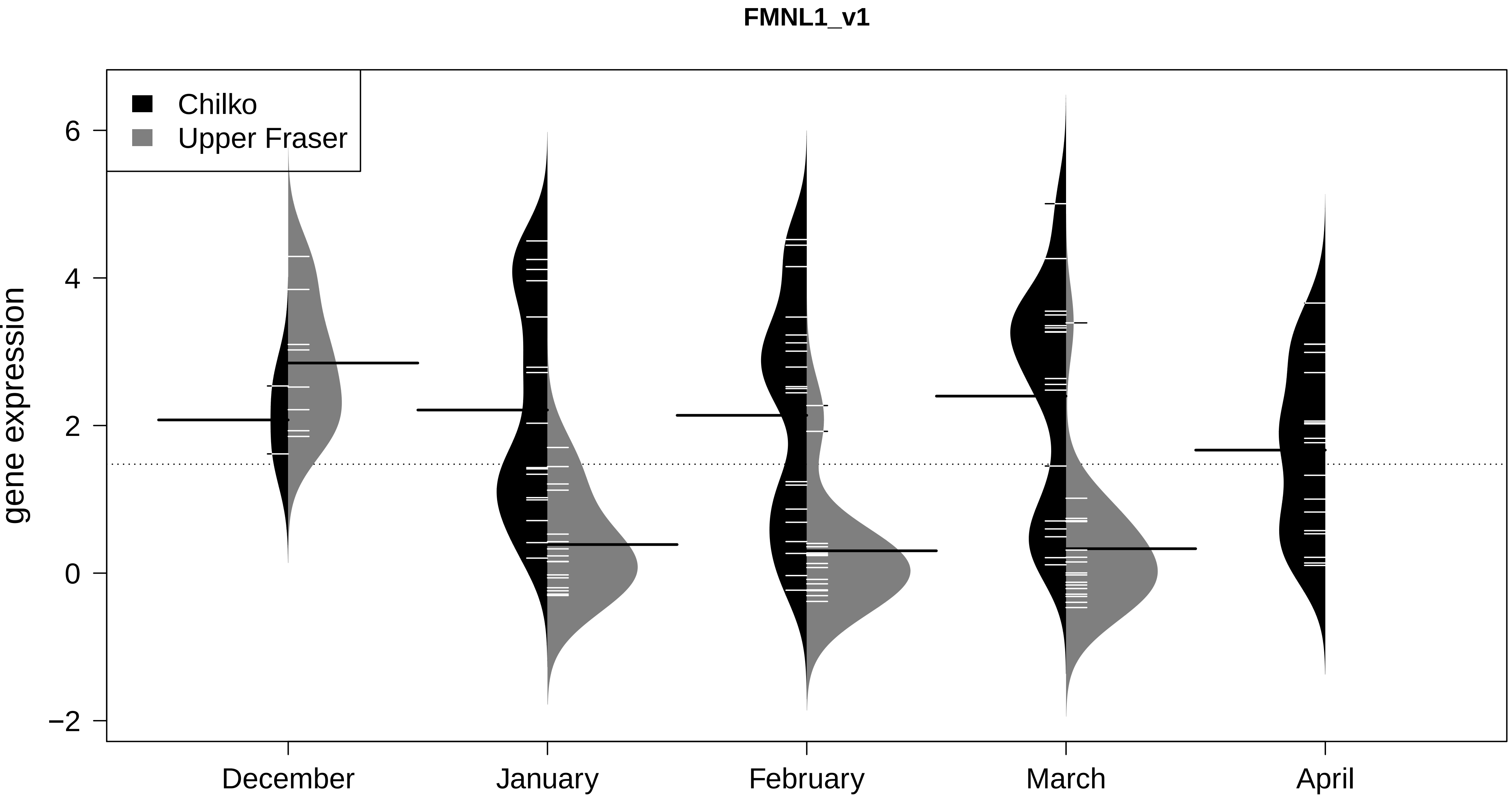
FMNL1_v1
Chilko
Upper Fraser
gene expression
6
4
2
0
−2
December
January
February
March
April

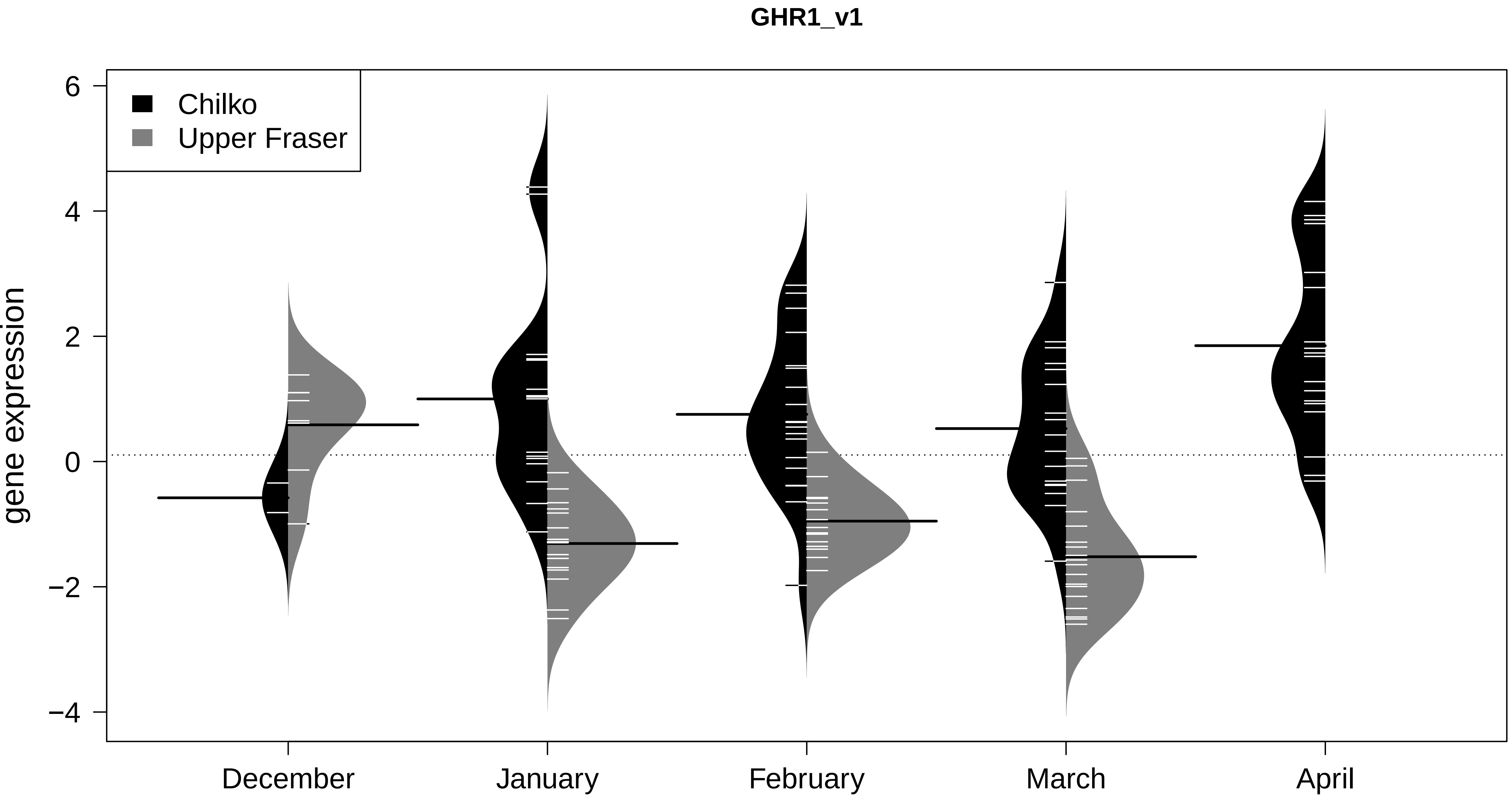

GHR1_v1
Chilko
Upper Fraser
gene expression
6
4
2
0
−2
−4
December
January
February
March
April

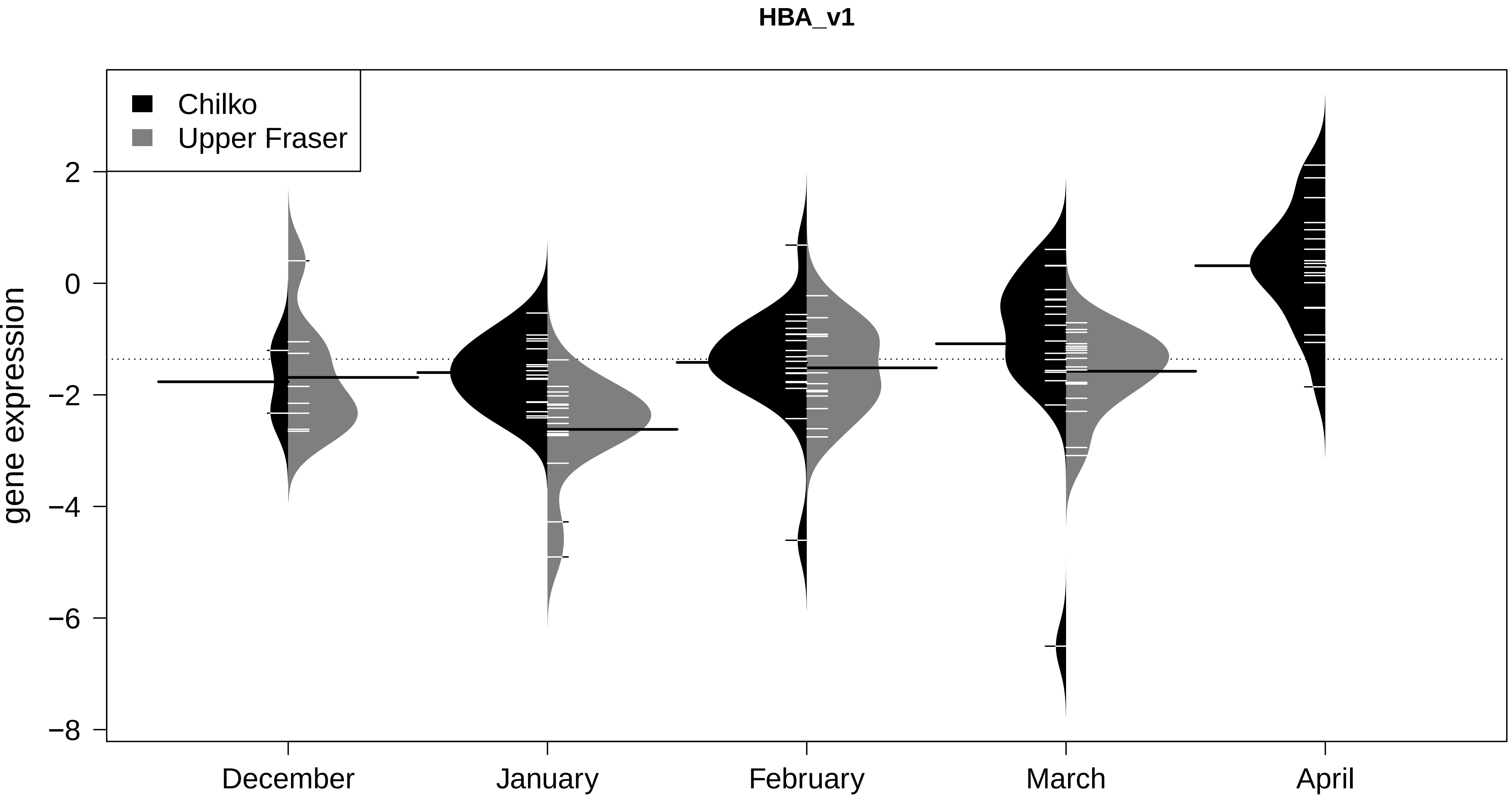
HBA_v1
Chilko
Upper Fraser
gene expression
2
0
−2
−4
−6
−8
December
January
February
March
April

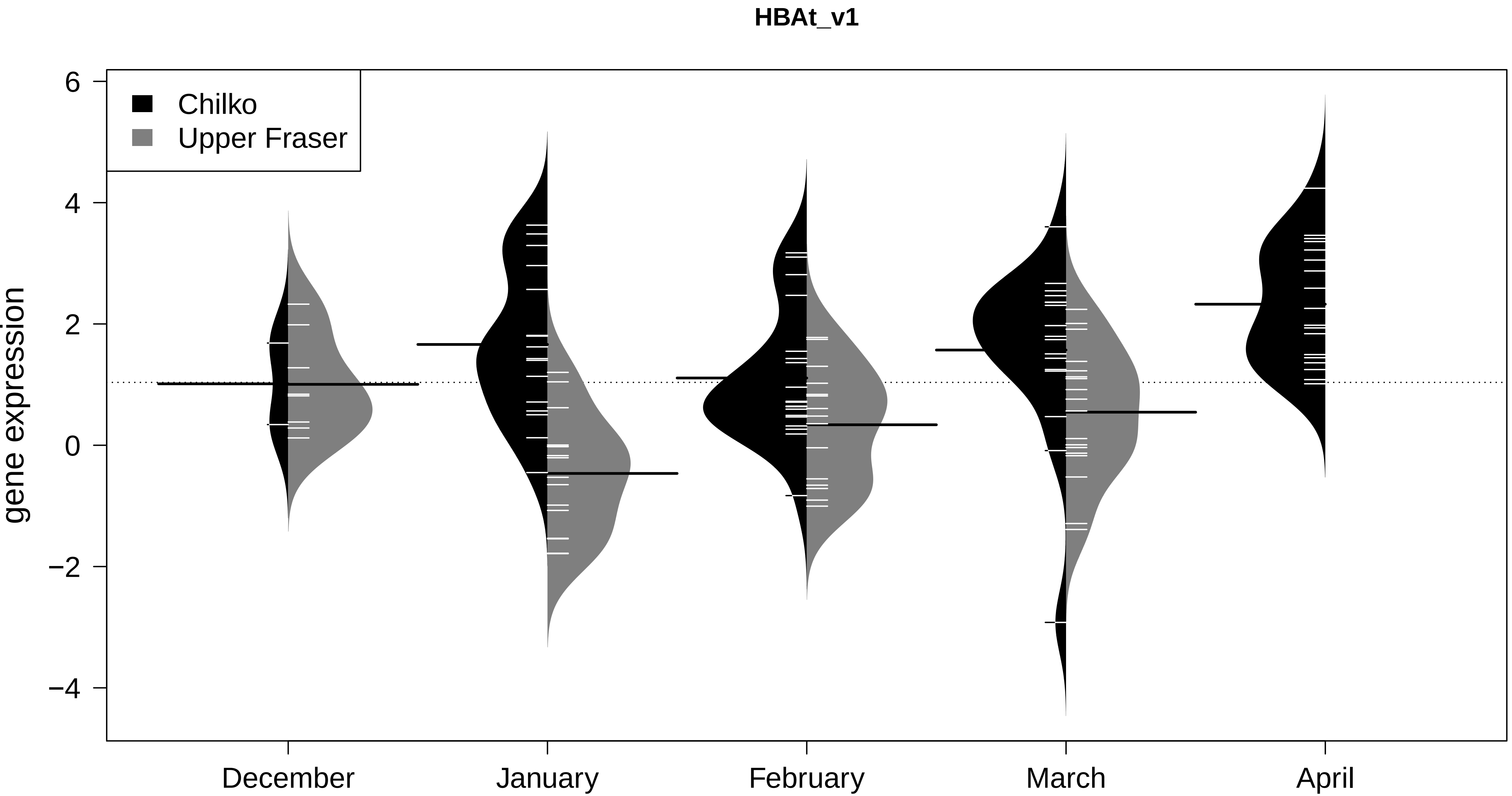
HBAt_v1
Chilko
Upper Fraser
gene expression
6
4
2
0
−2
−4
December
January
February
March
April

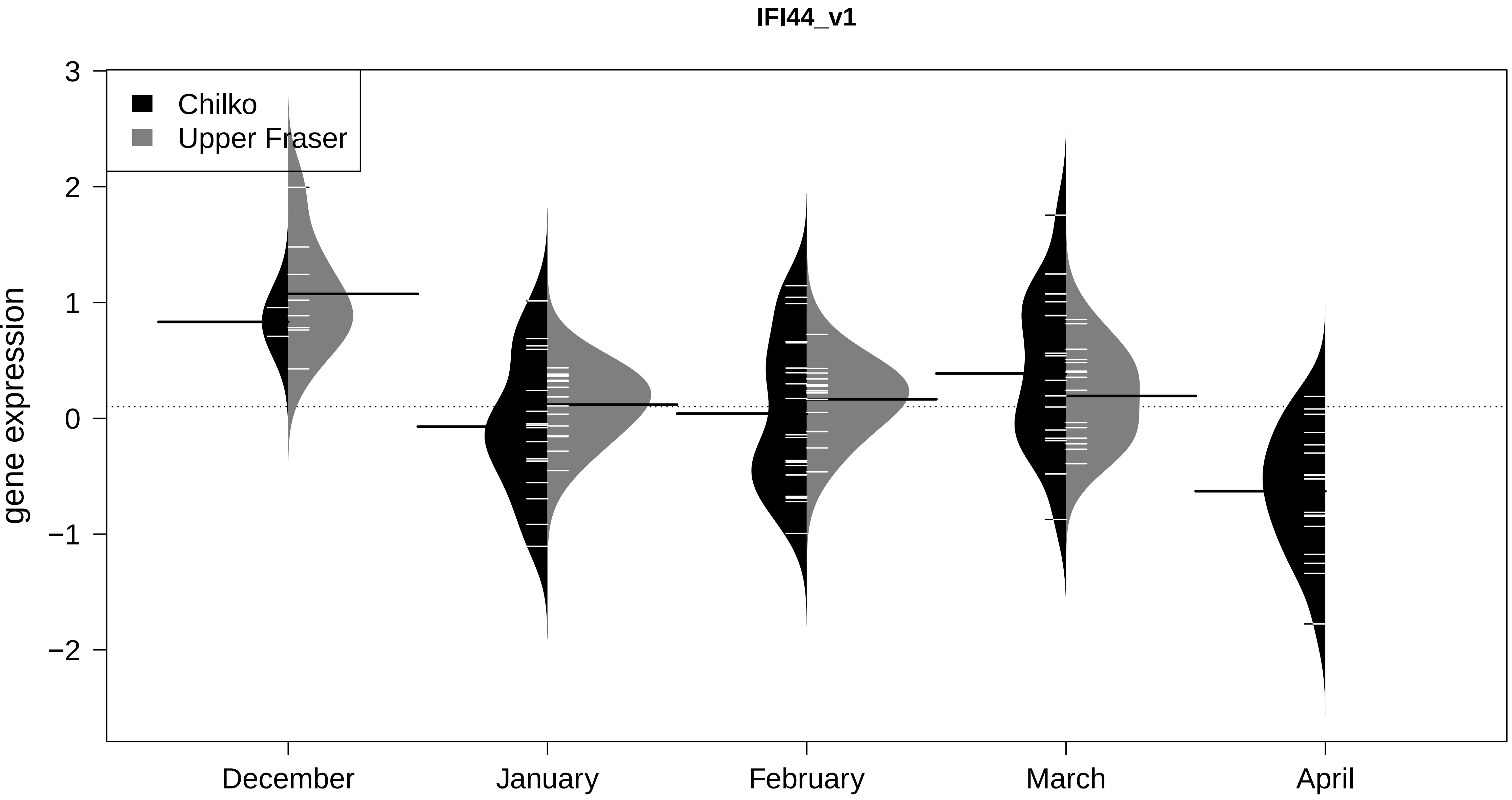
IFI44_v1
Chilko
Upper Fraser
gene expression
3
2
1
0
−1
−2
December
January
February
March
April

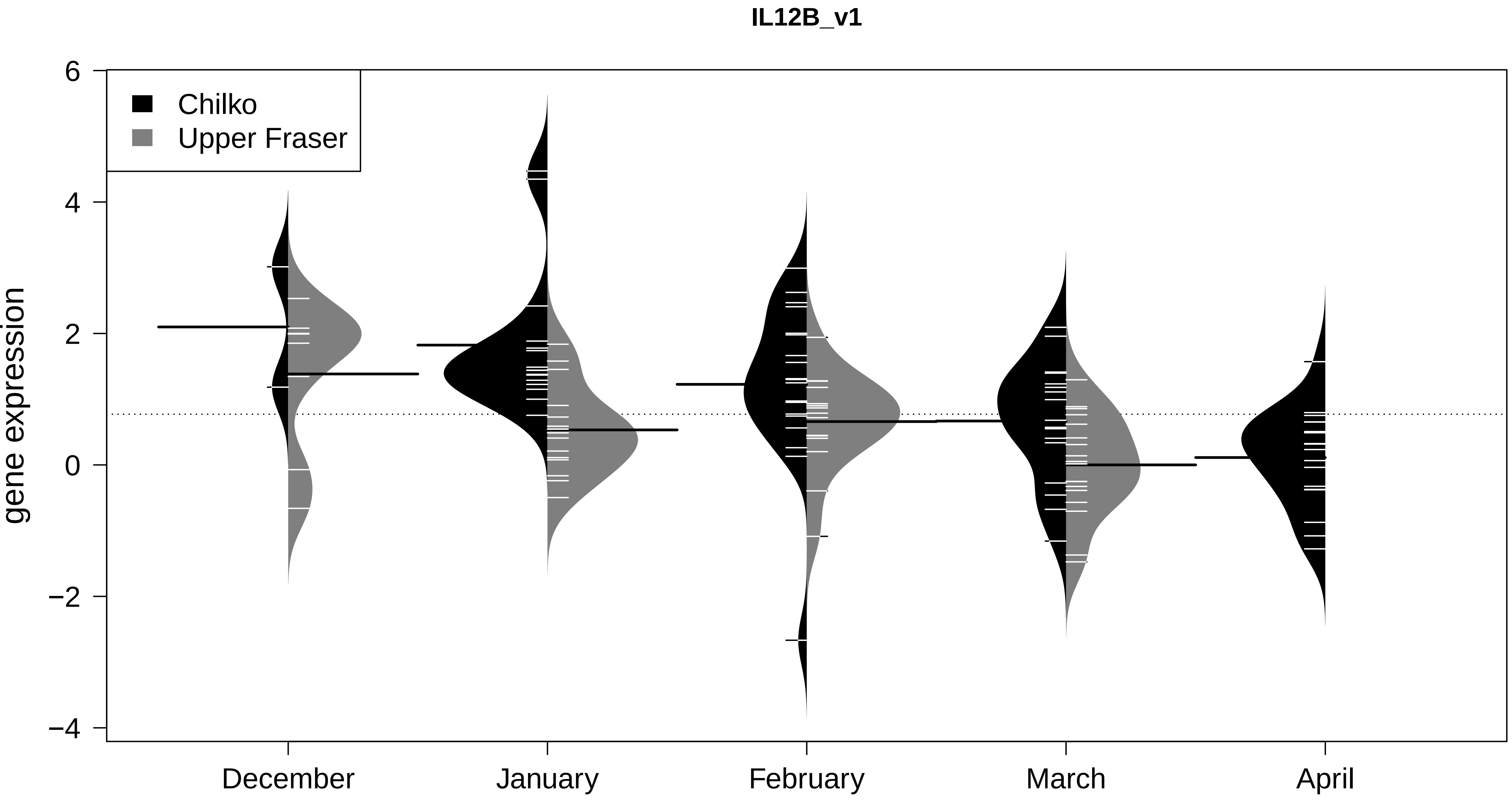
IL12B_v1
Chilko
Upper Fraser
gene expression
6
4
2
0
−2
−4
December
January
February
March
April

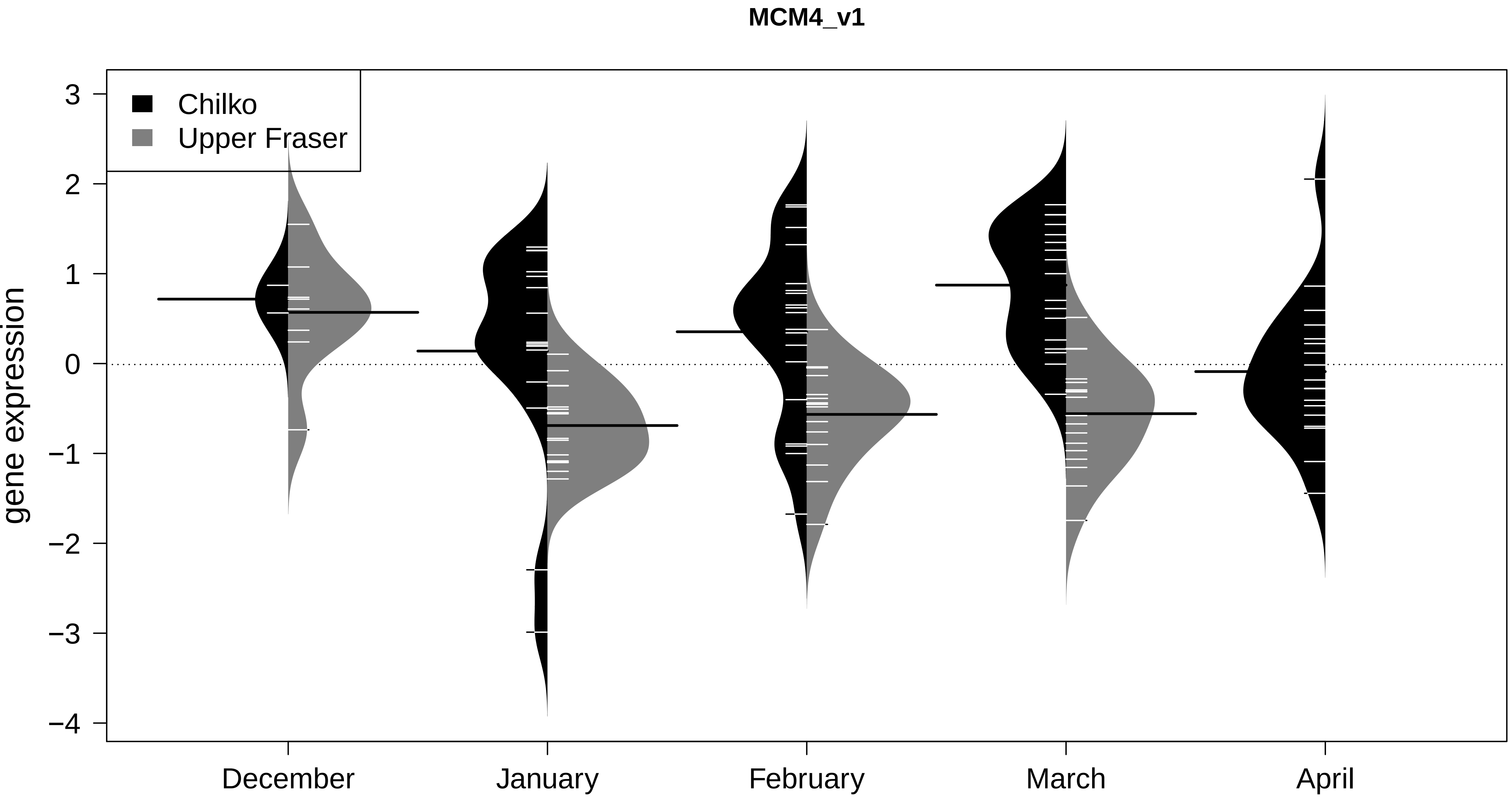

MCM4_v1
Chilko
Upper Fraser
gene expression
3
2
1
0
−1
−2
−3
−4
December
January
February
March
April

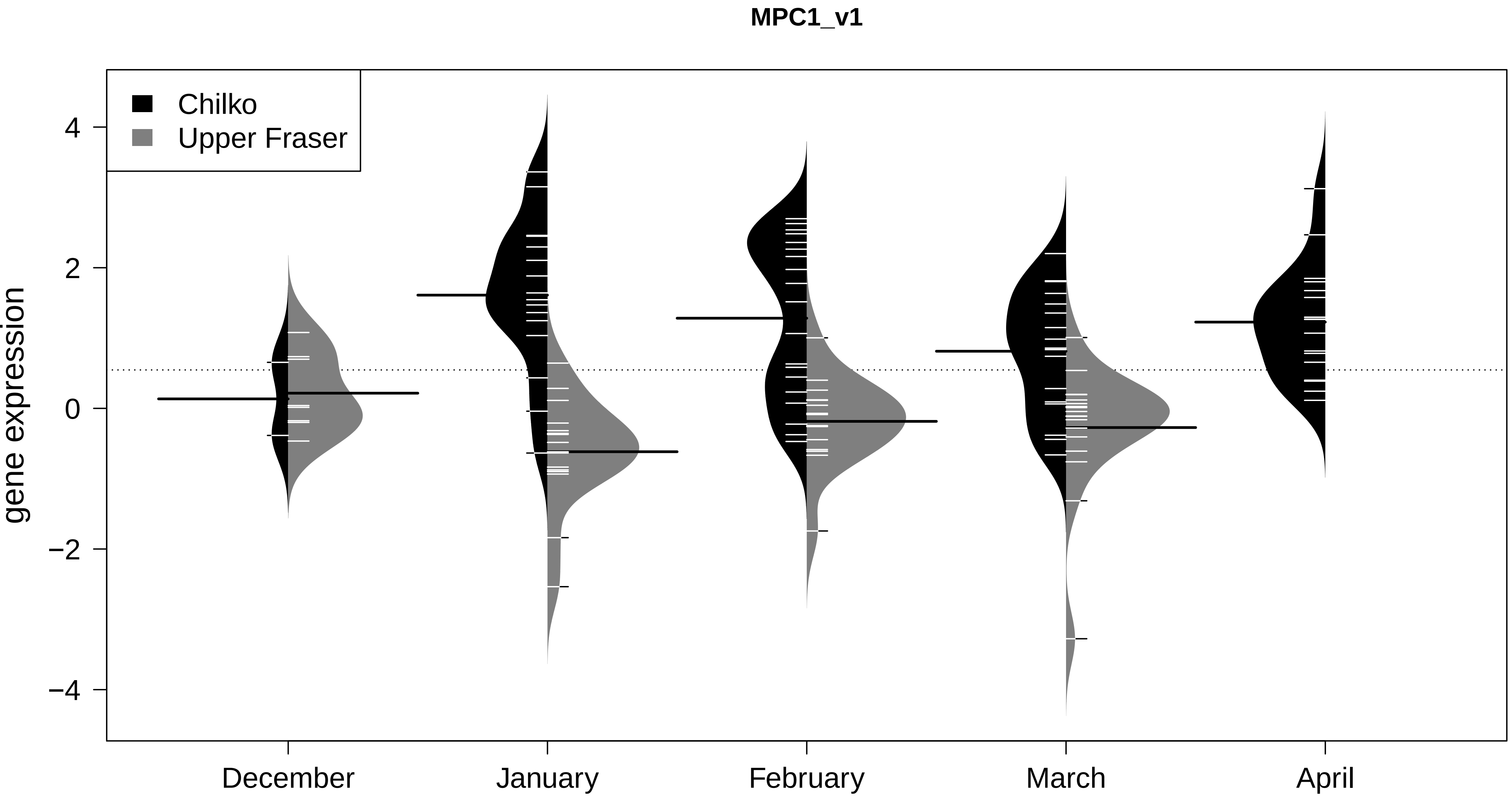
MPC1_v1
Chilko
Upper Fraser
gene expression
4
2
0
−2
−4
December
January
February
March
April

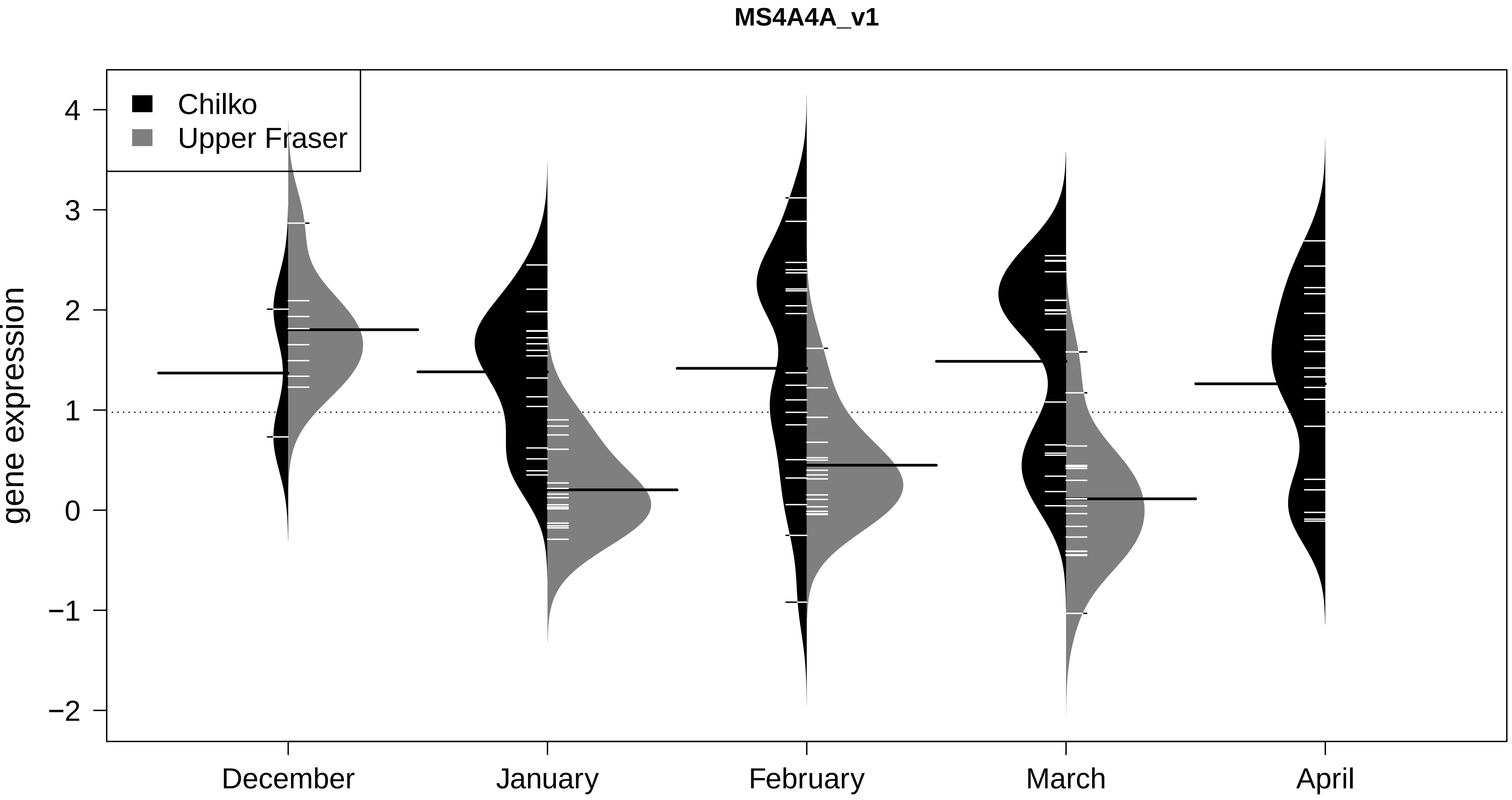

MS4A4A_v1
Chilko
Upper Fraser
gene expression
4
3
2
1
0
−1
−2
December
January
February
March
April

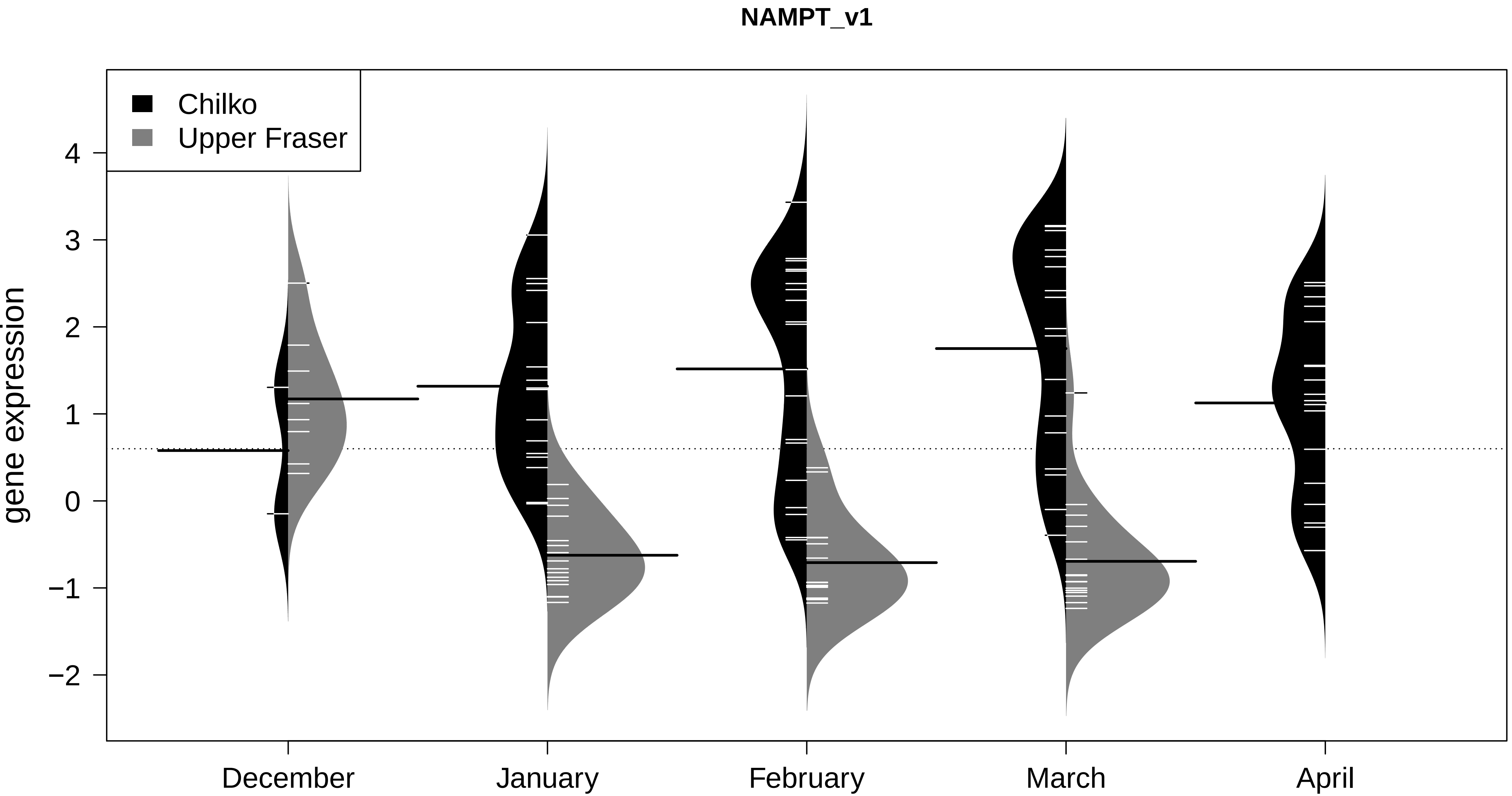

NAMPT_v1
Chilko
Upper Fraser
gene expression
4
3
2
1
0
−1
−2
December
January
February
March
April

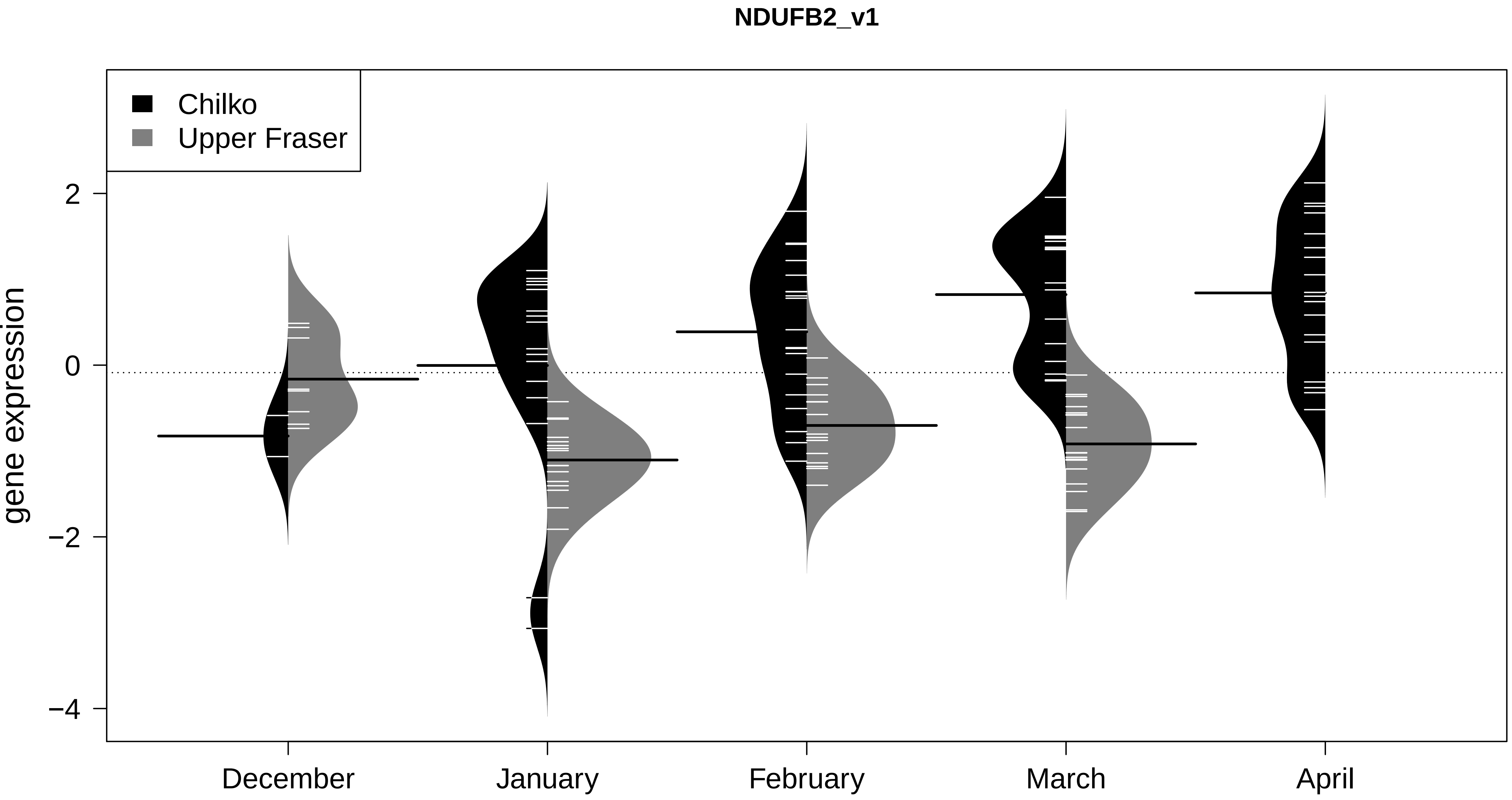

NDUFB2_v1
Chilko
Upper Fraser
gene expression
2
0
−2
−4
December
January
February
March
April

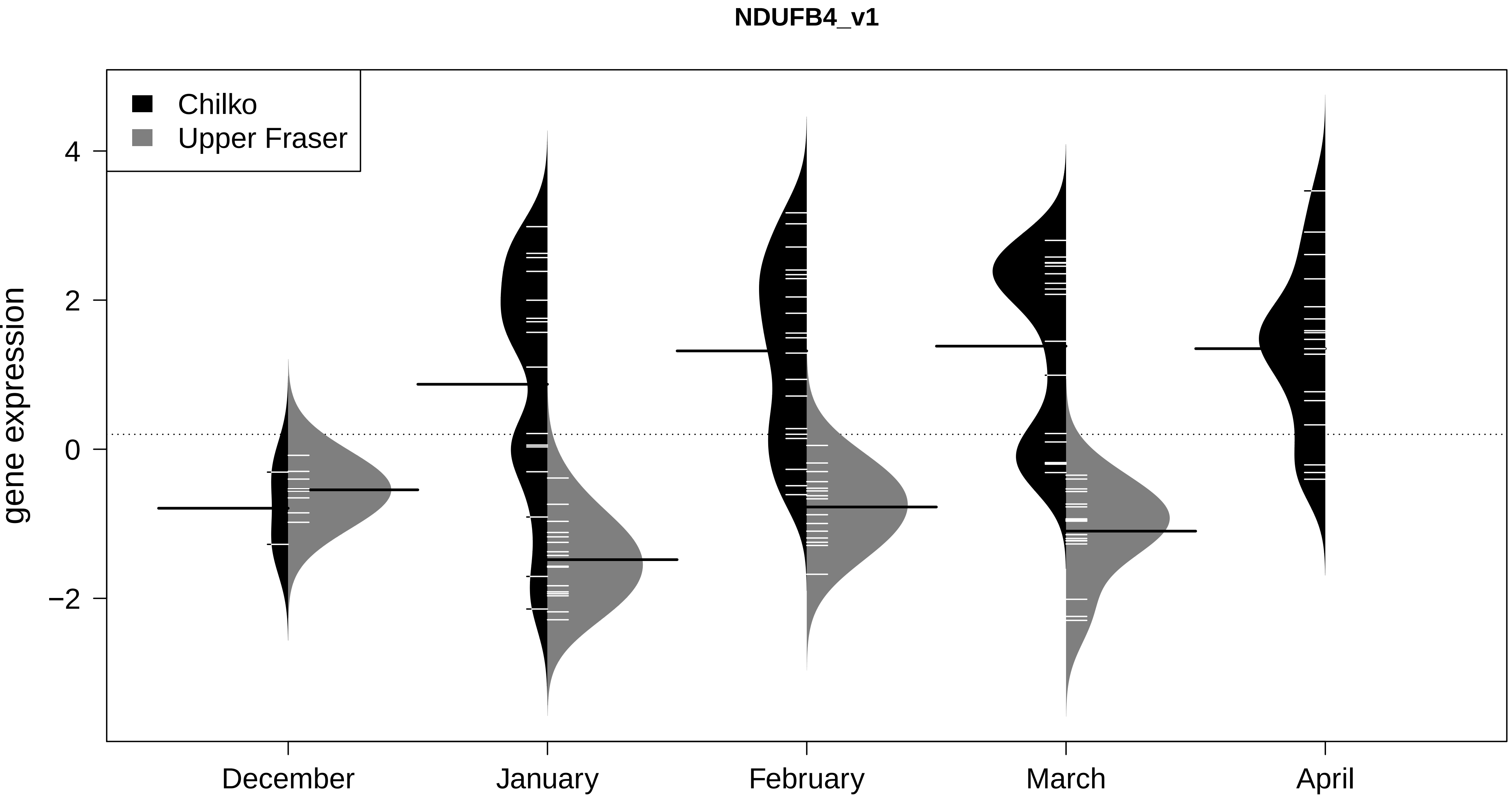
NDUFB4_v1
Chilko
Upper Fraser
gene expression
4
2
0
−2
December
January
February
March
April

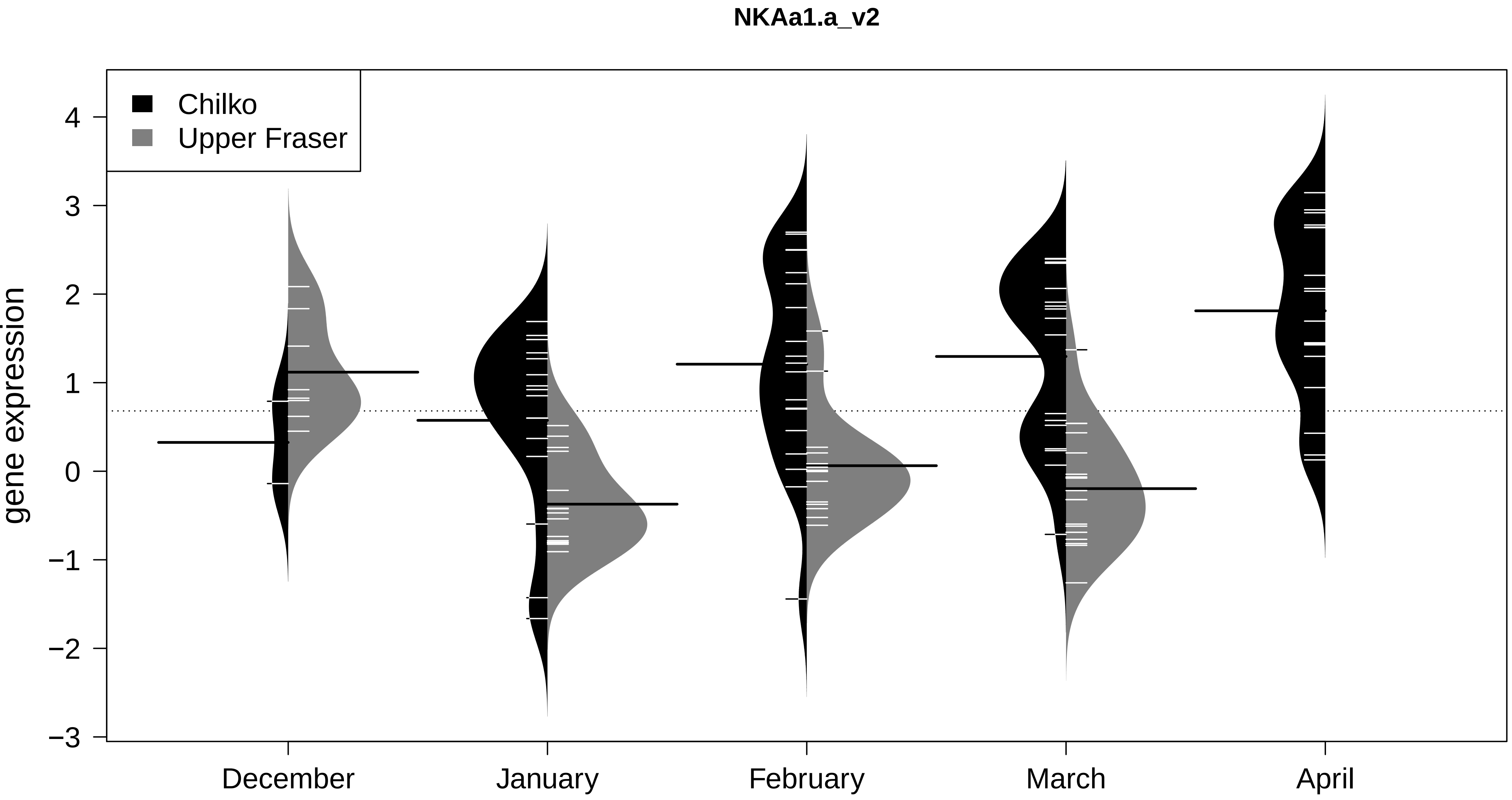
NKAa1.a_v2
Chilko
Upper Fraser
gene expression
4
3
2
1
0
−1
−2
−3
December
January
February
March
April

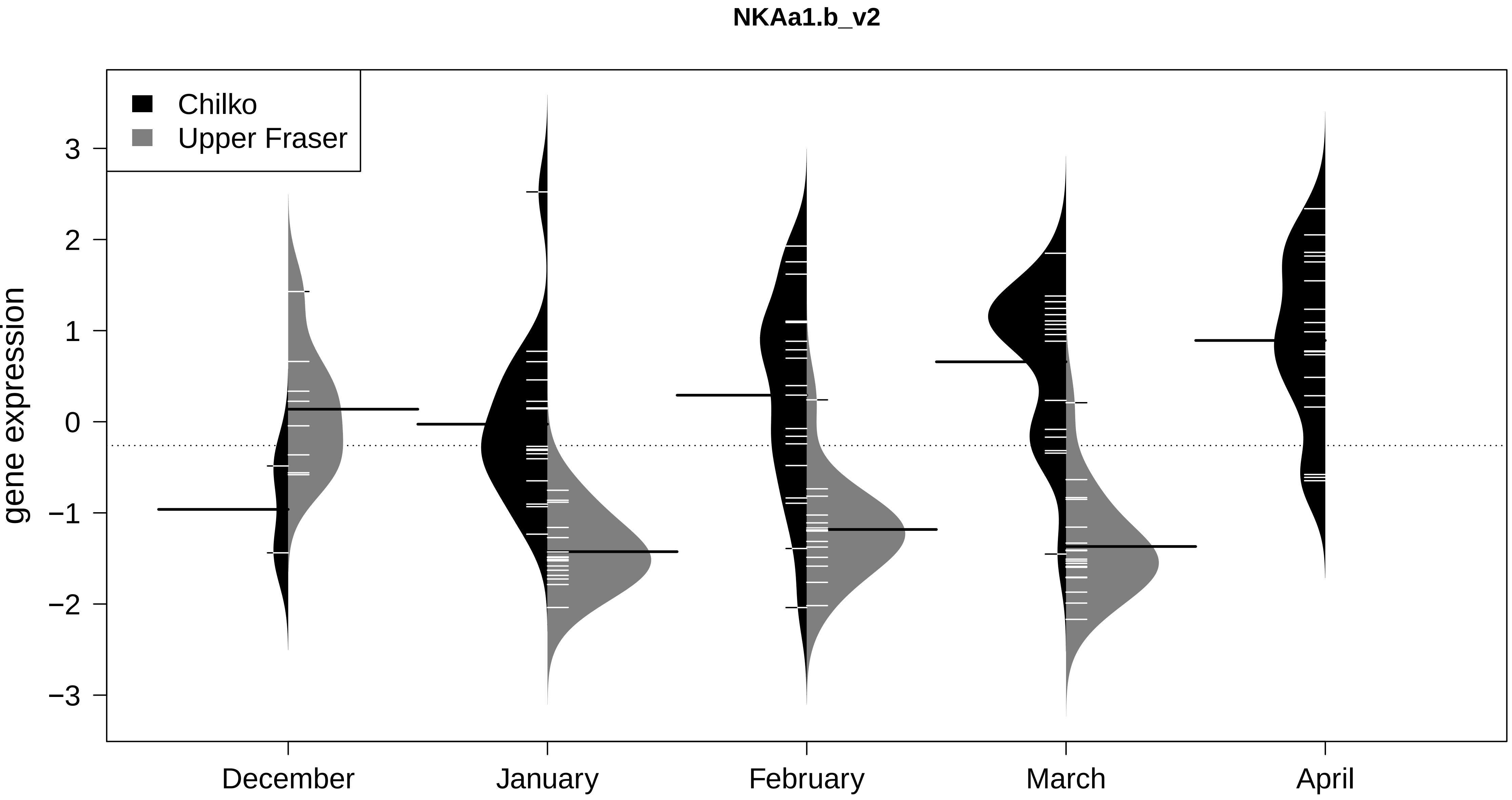

NKAa1.b_v2
Chilko
Upper Fraser
gene expression
3
2
1
0
−1
−2
−3
December
January
February
March
April

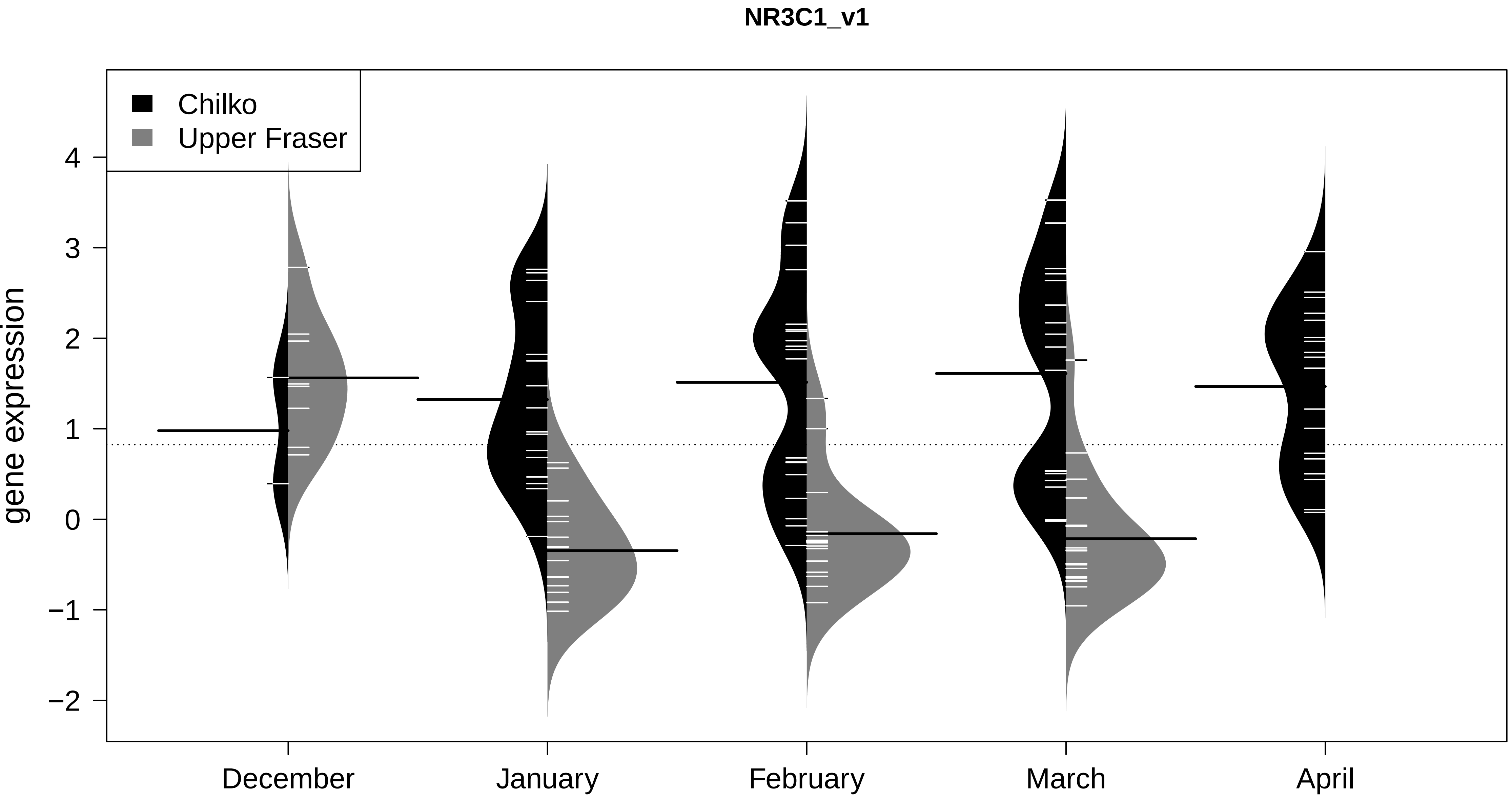

NR3C1_v1
Chilko
Upper Fraser
gene expression
4
3
2
1
0
−1
−2
December
January
February
March
April

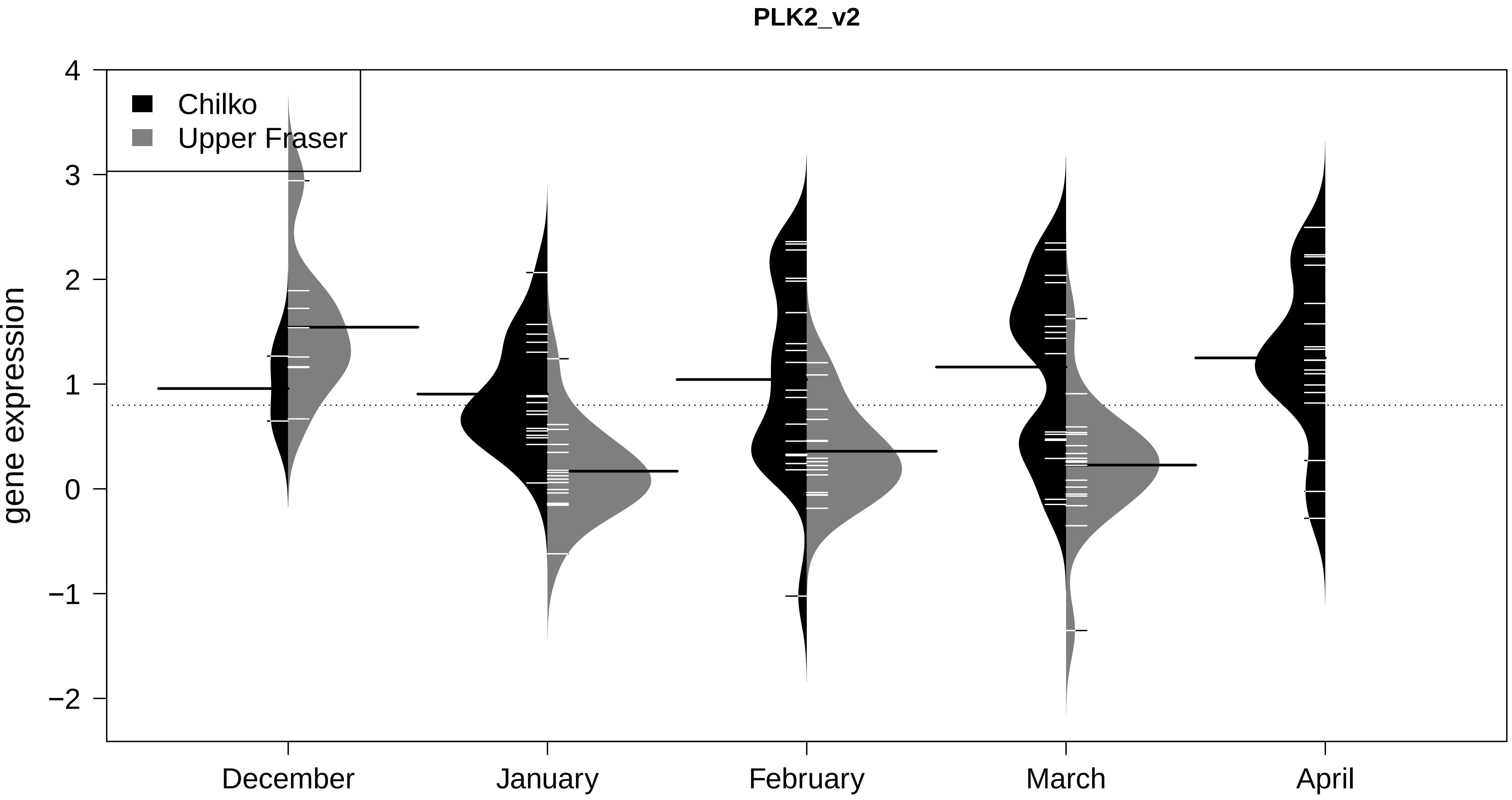

PLK2_v2
Chilko
Upper Fraser
gene expression
4
3
2
1
0
−1
−2
December
January
February
March
April

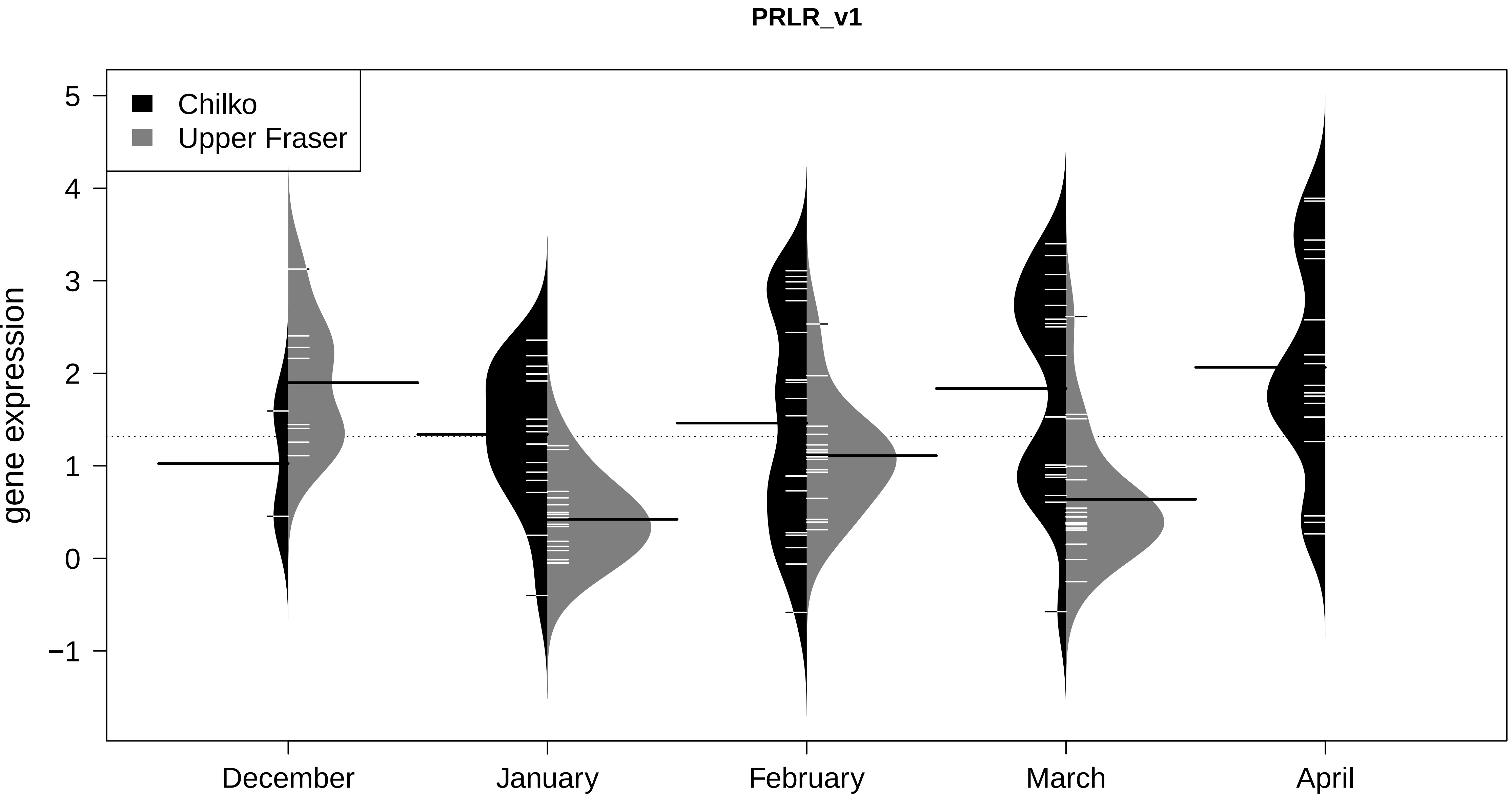

PRLR_v1
Chilko
Upper Fraser
gene expression
5
4
3
2
1
0
−1
December
January
February
March
April

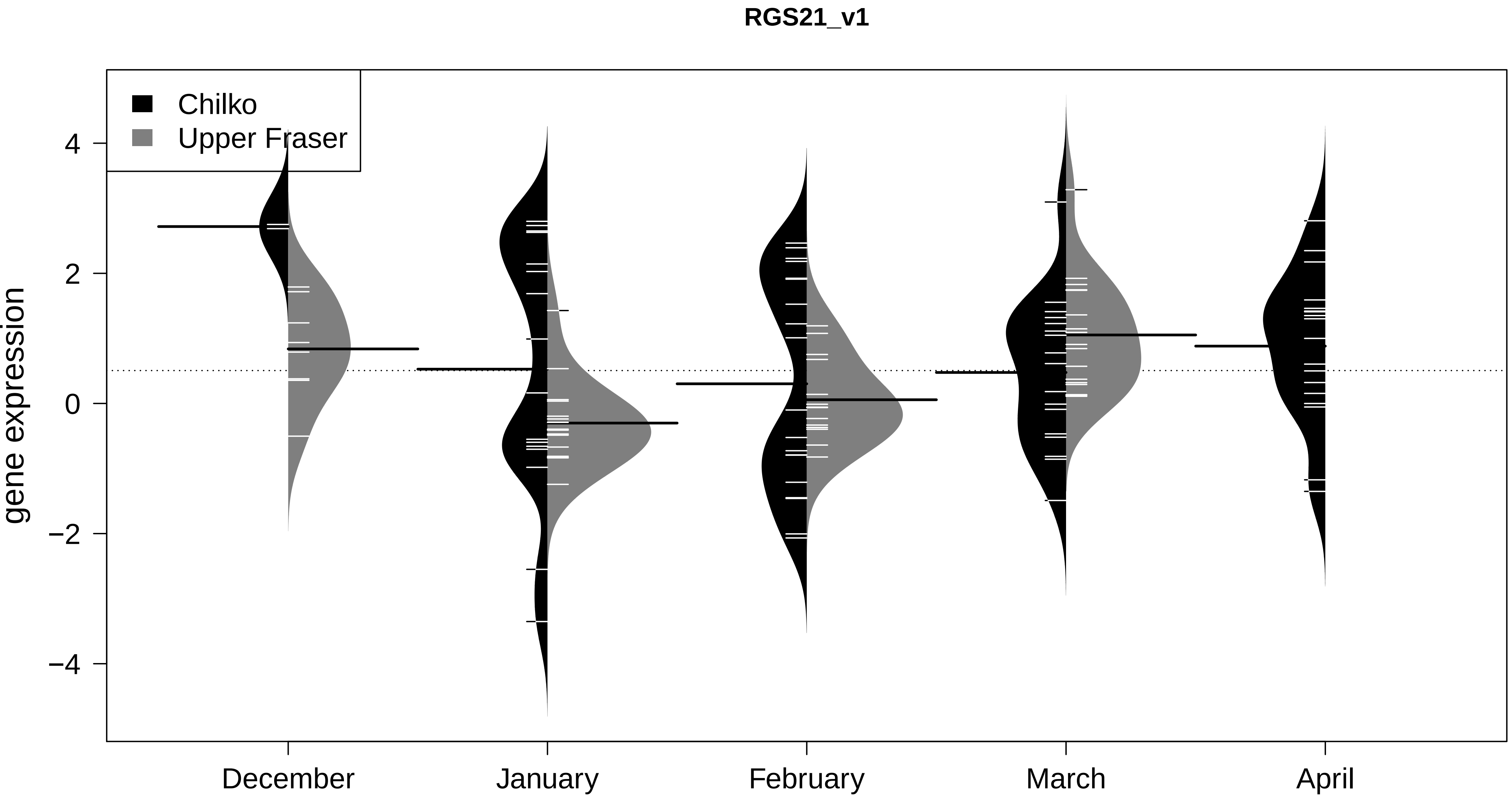

RGS21_v1
Chilko
Upper Fraser
gene expression
4
2
0
−2
−4
December
January
February
March
April

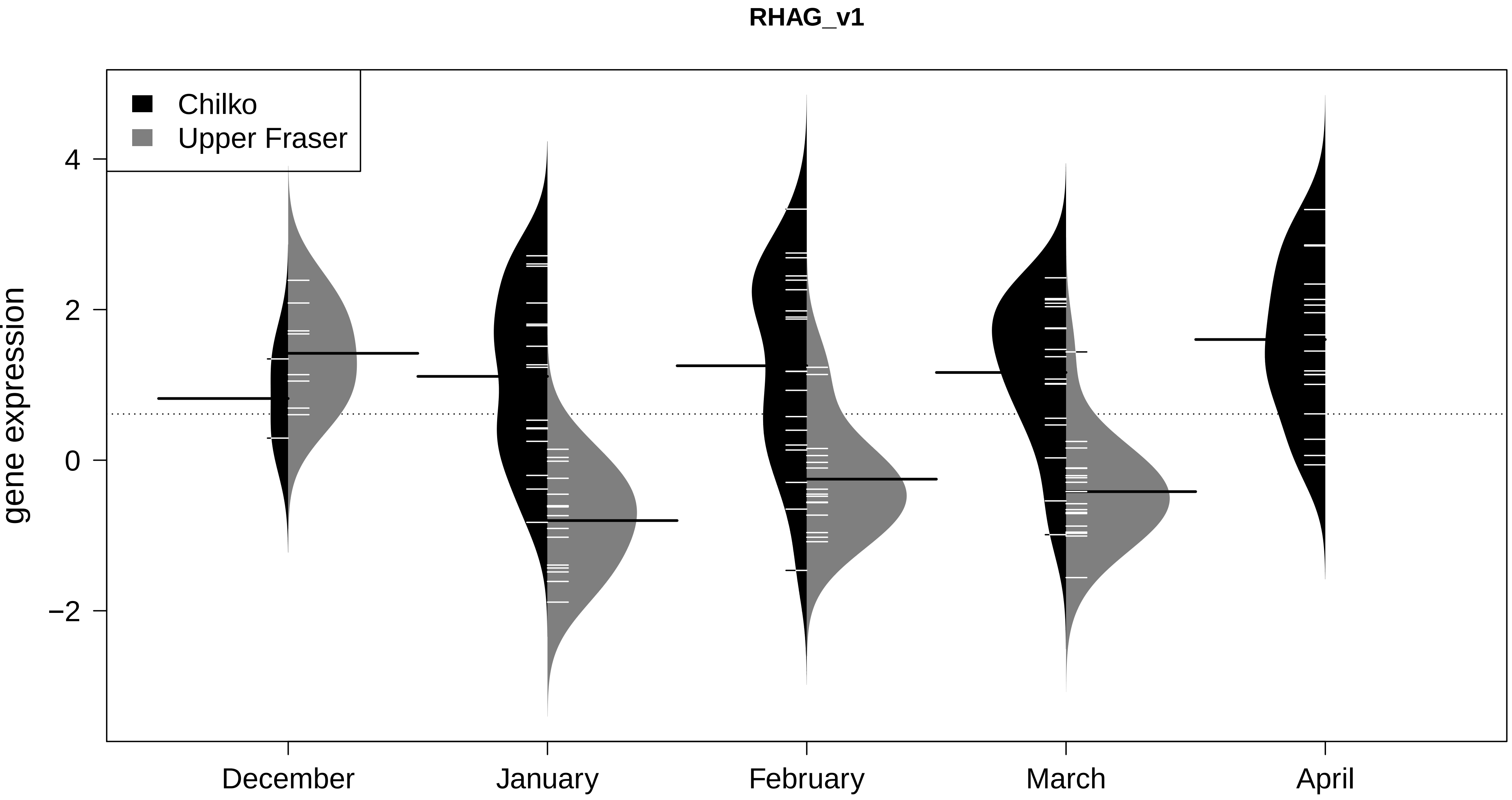
RHAG_v1
Chilko
Upper Fraser
gene expression
4
2
0
−2
December
January
February
March
April

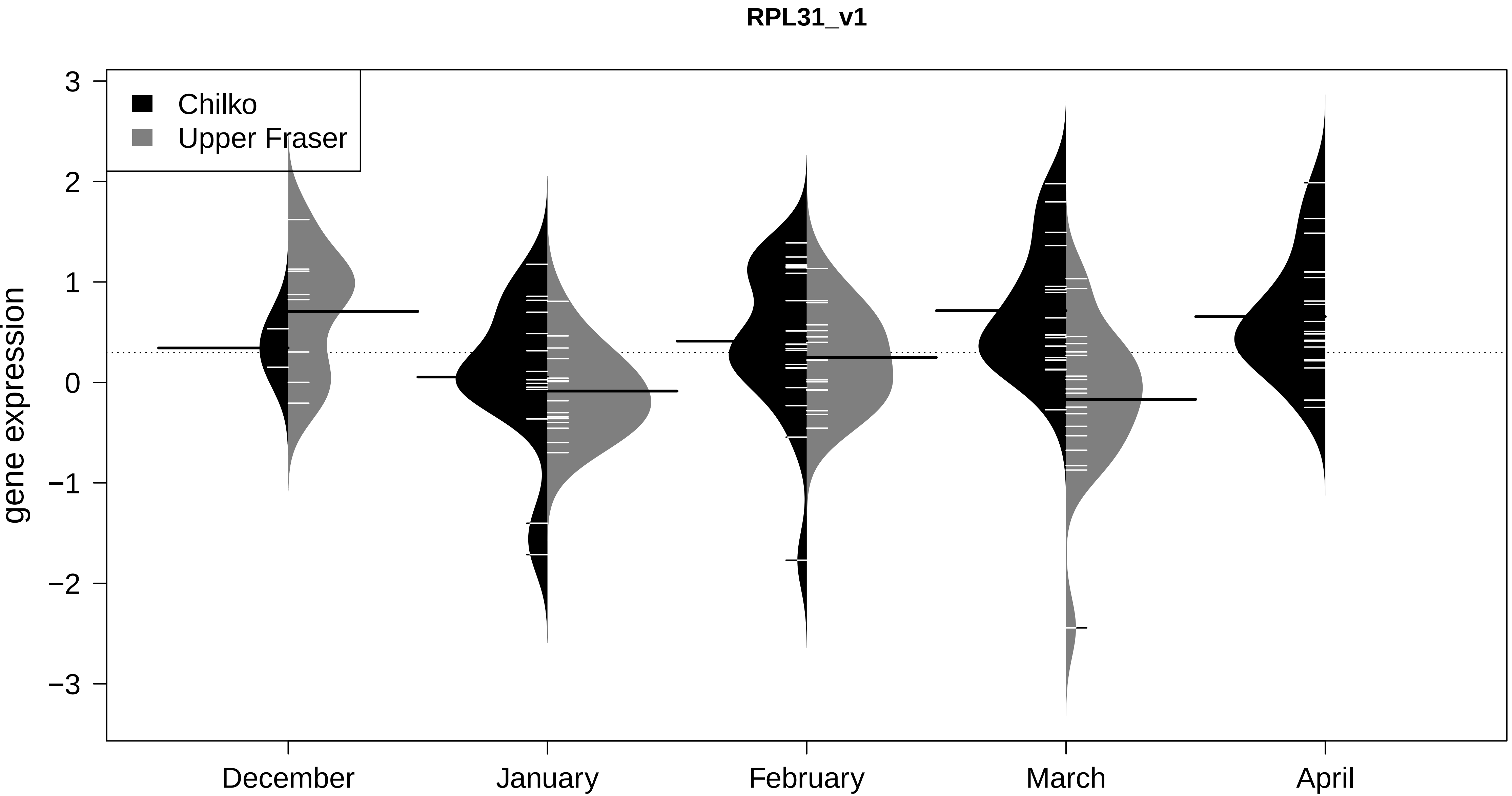

RPL31_v1
Chilko
Upper Fraser
gene expression
3
2
1
0
−1
−2
−3
December
January
February
March
April

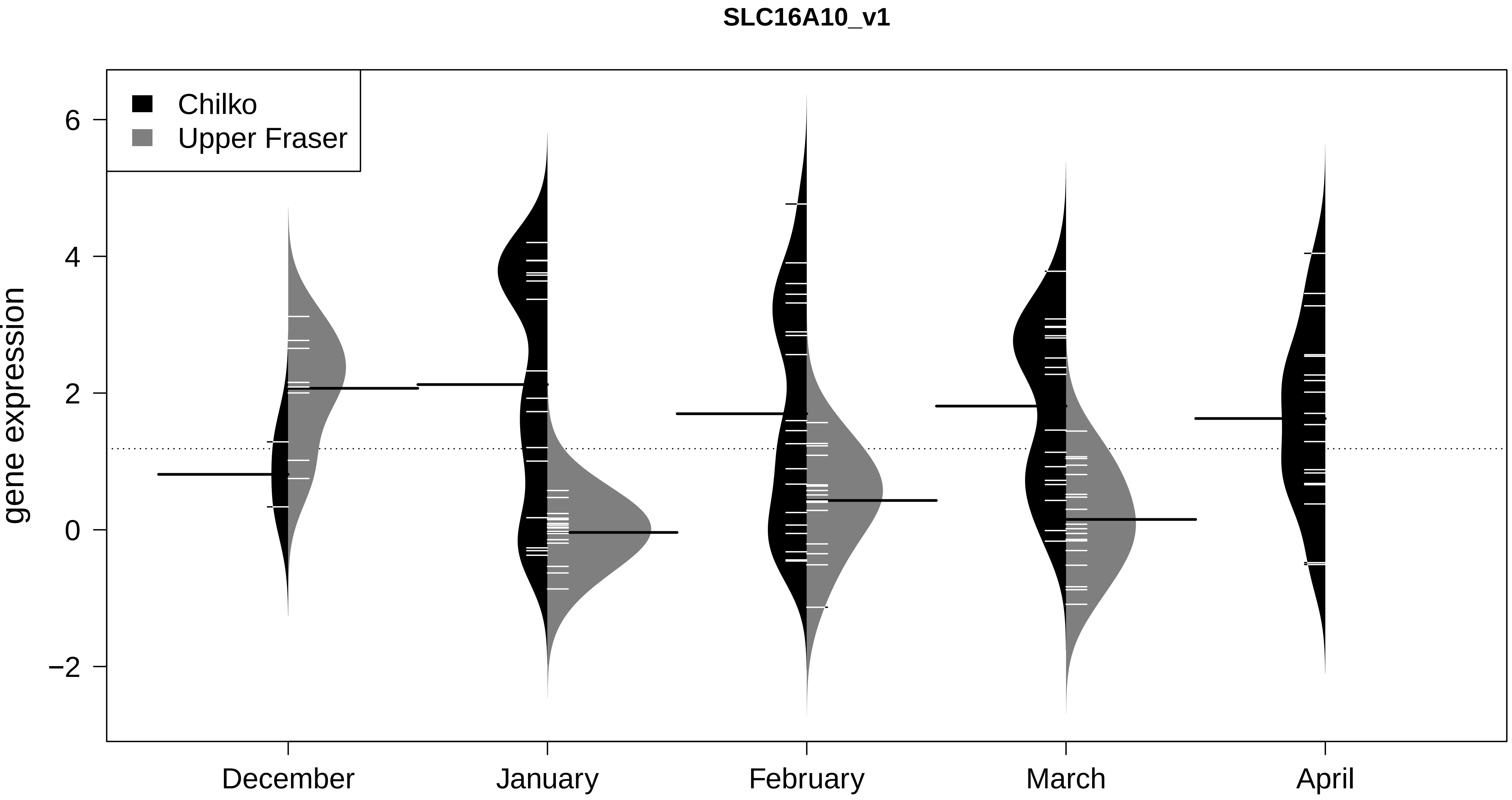
SLC16A10_v1
Chilko
Upper Fraser
gene expression
6
4
2
0
−2
December
January
February
March
April

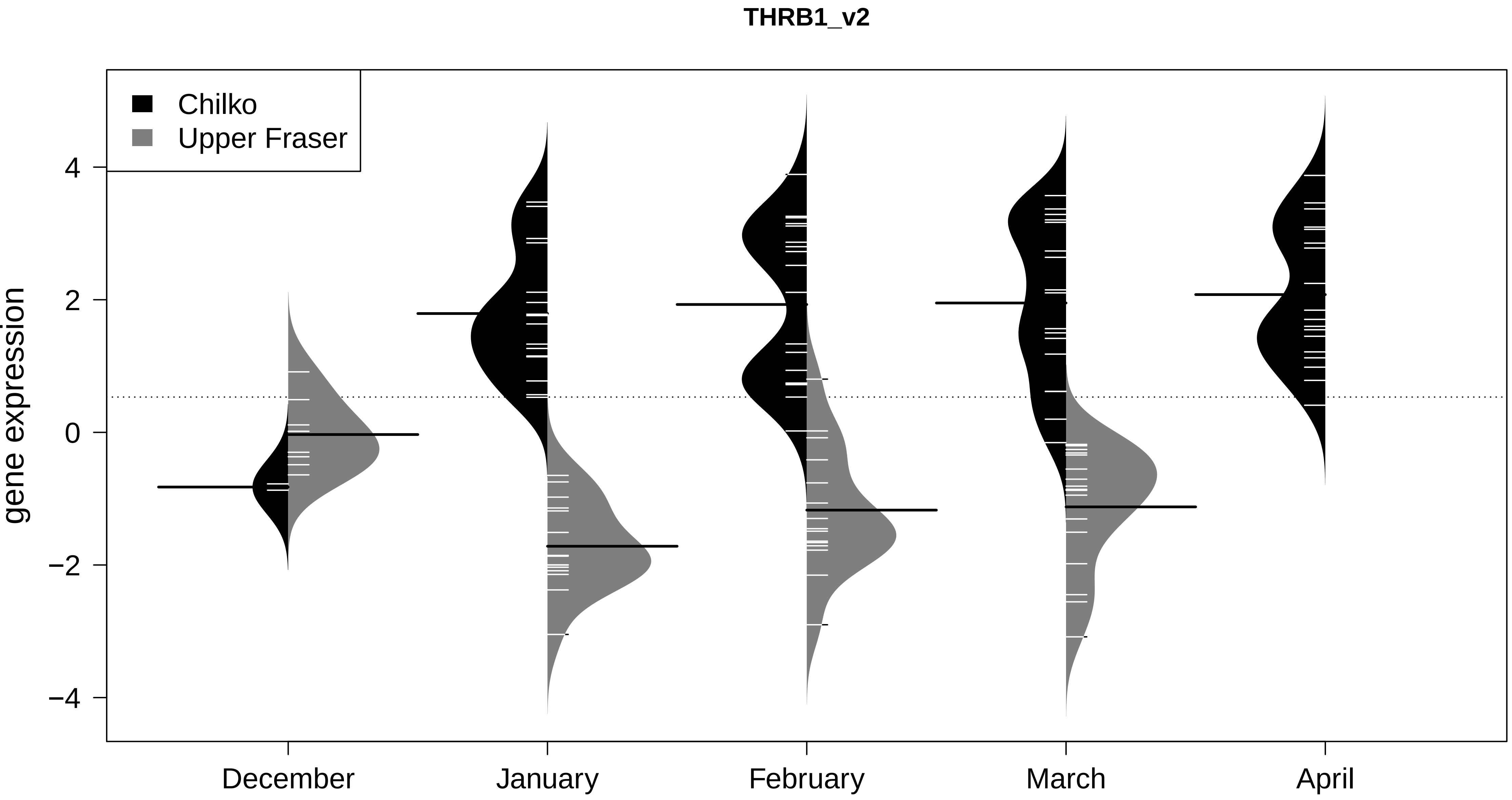
THRB1_v2
Chilko
Upper Fraser
gene expression
4
2
0
−2
−4
December
January
February
March
April

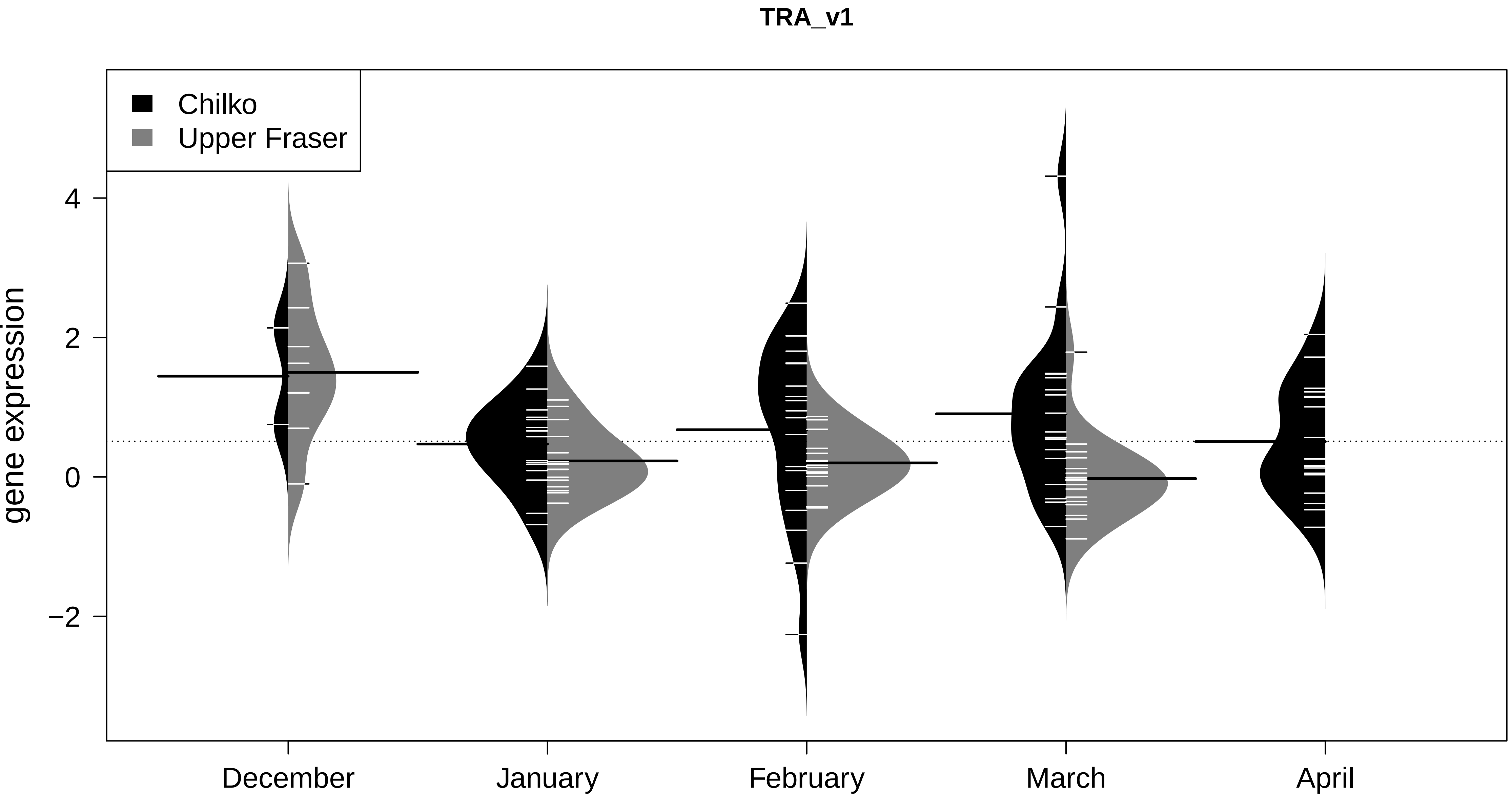
TRA_v1
Chilko
Upper Fraser
gene expression
4
2
0
−2
December
January
February
March
April

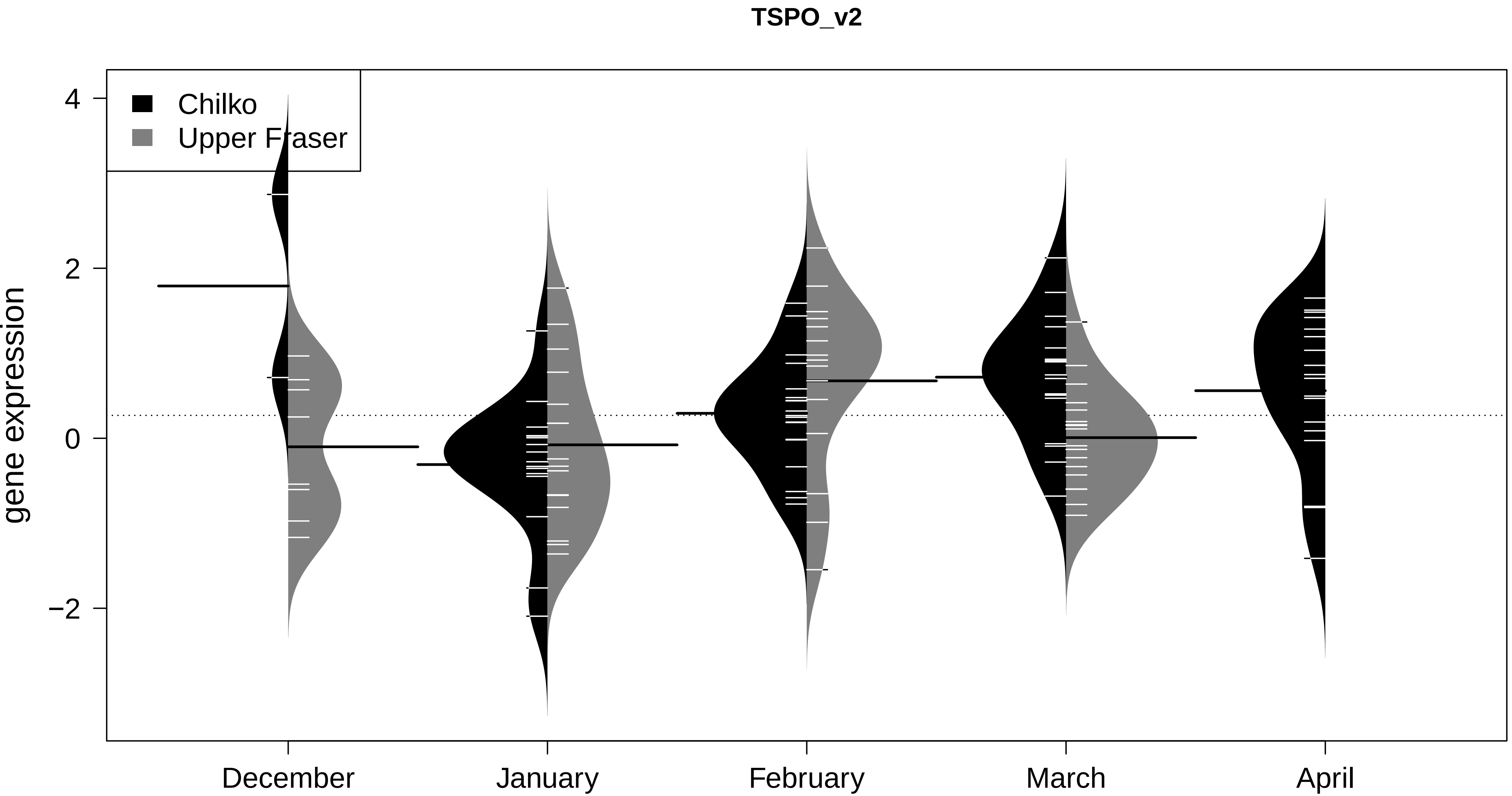
TSPO_v2
Chilko
Upper Fraser
gene expression
4
2
0
−2
December
January
February
March
April

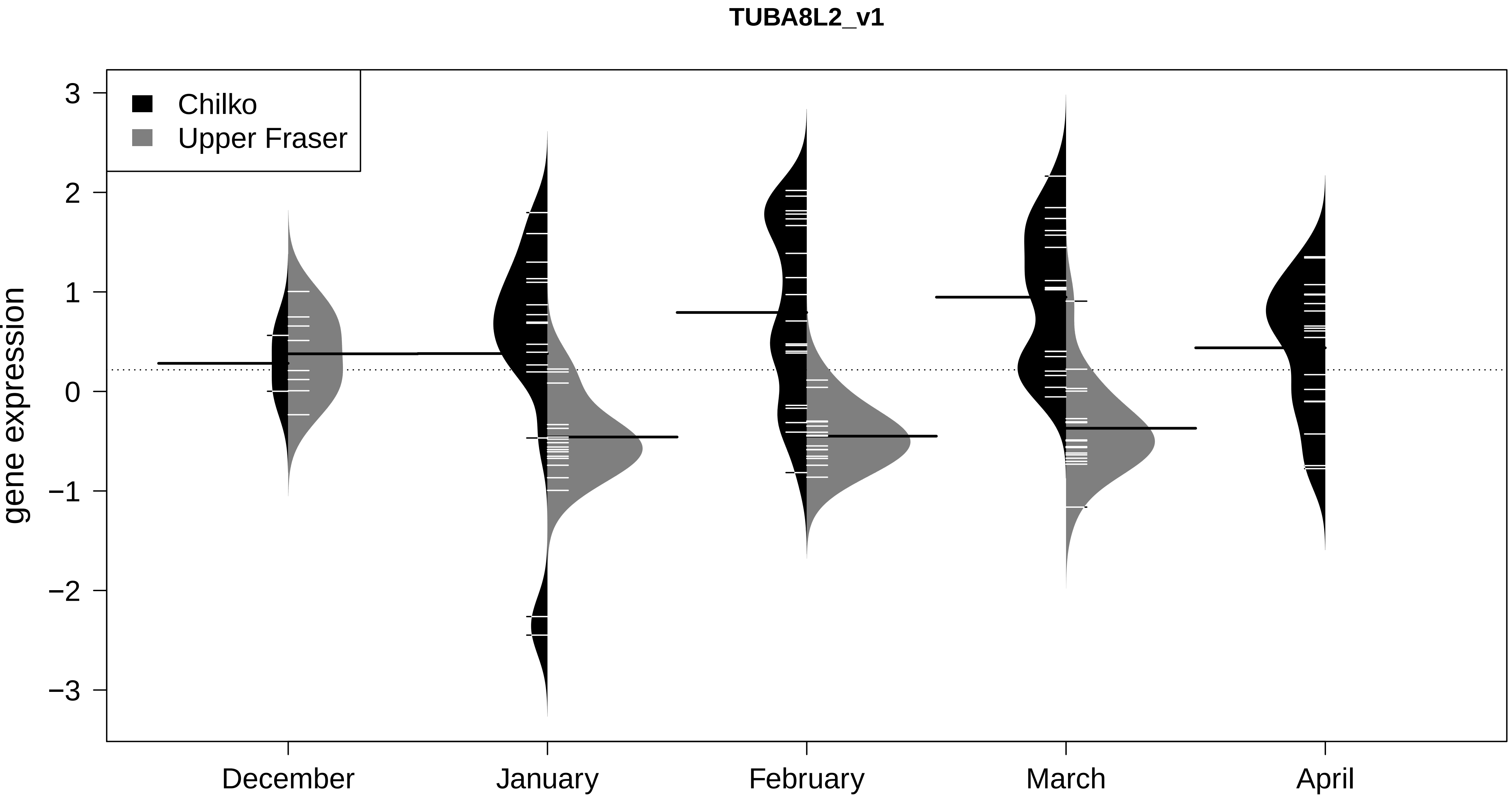

TUBA8L2_v1
Chilko
Upper Fraser
gene expression
3
2
1
0
−1
−2
−3
December
January
February
March
April

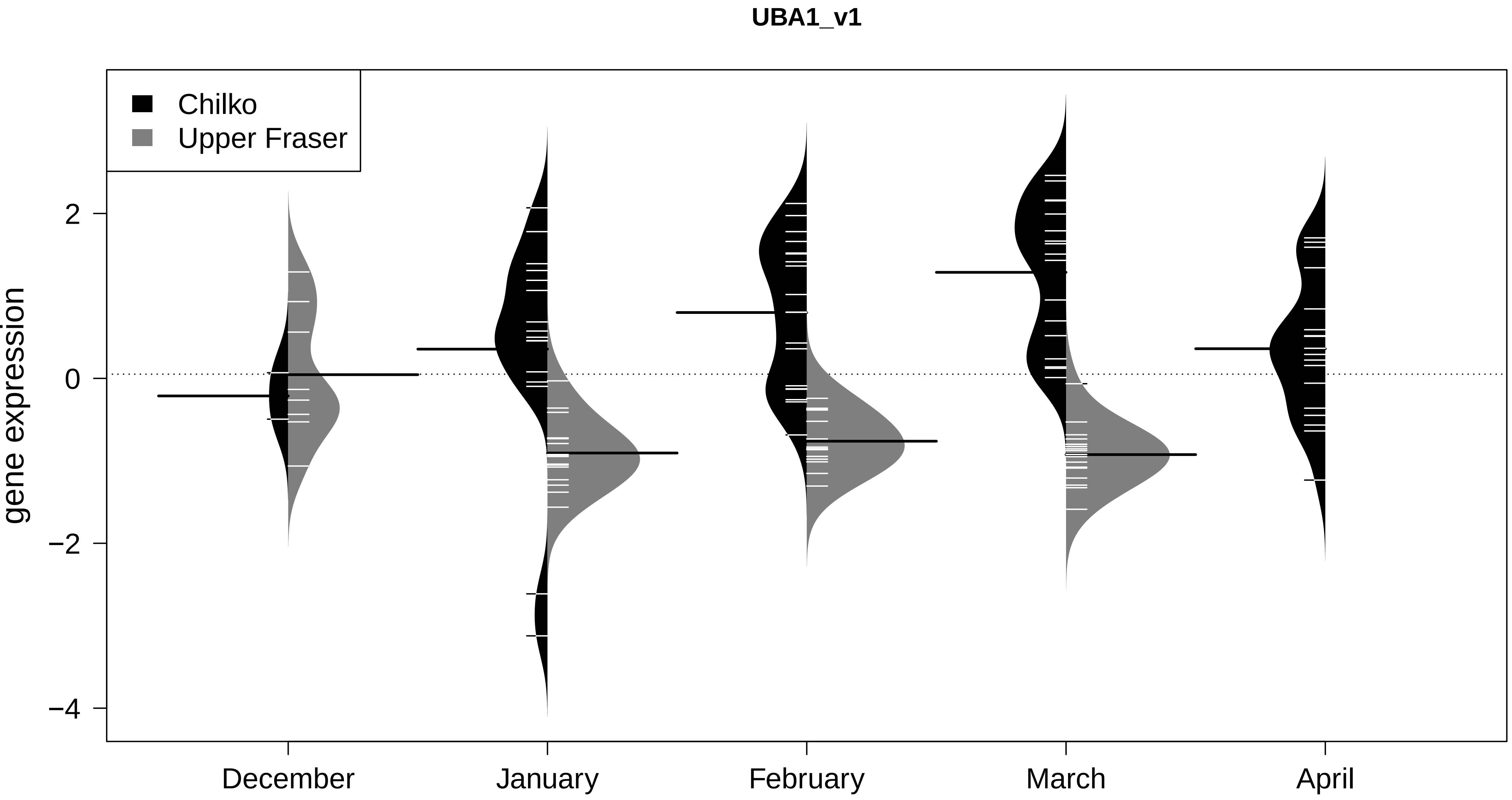

UBA1_v1
Chilko
Upper Fraser
gene expression
2
0
−2
−4
December
January
February
March
April

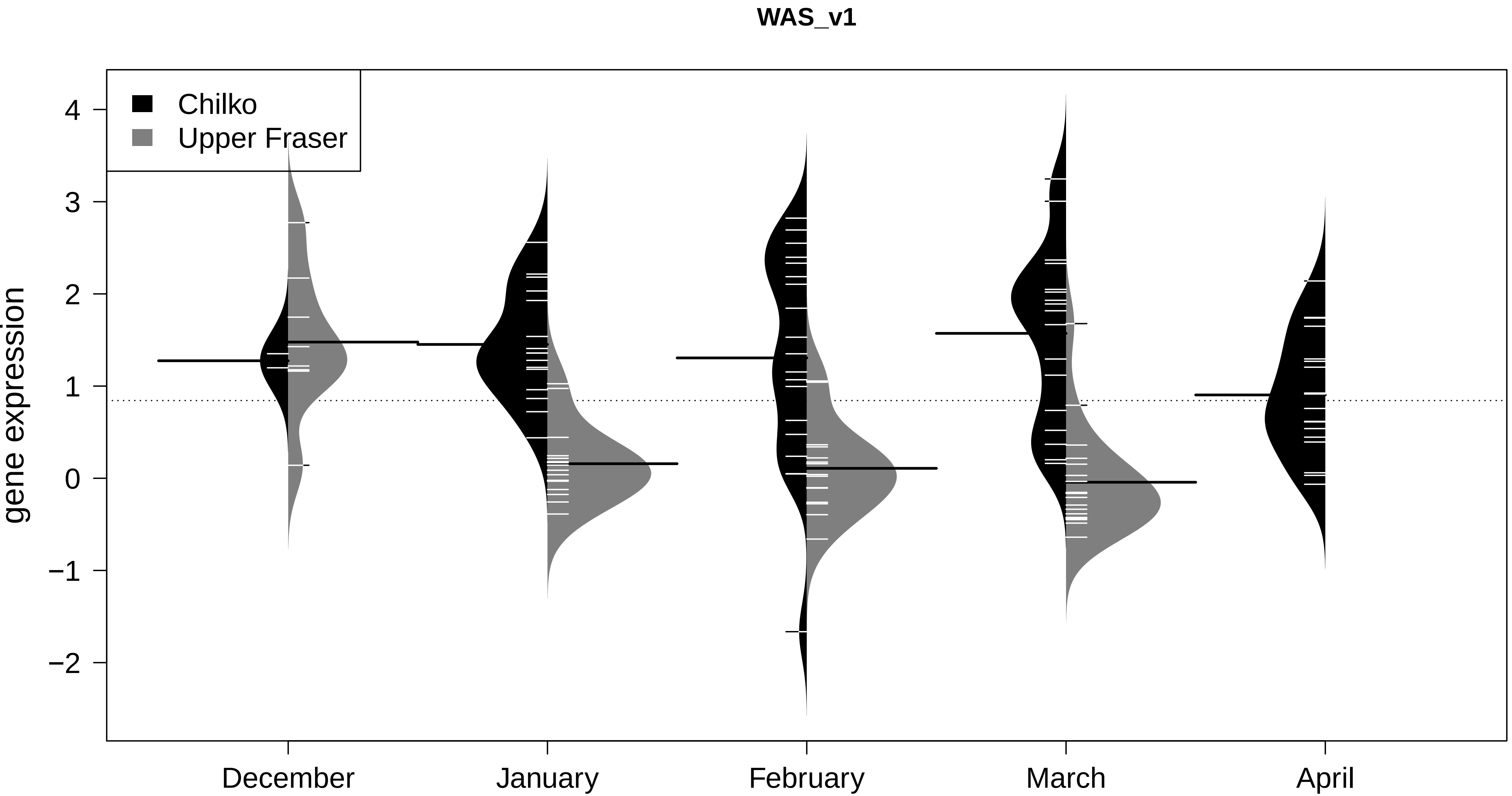

WAS_v1
Chilko
Upper Fraser
gene expression
4
3
2
1
0
−1
−2
December
January
February
March
April
