## appendix 4 for "Discovery and validation of candidate smoltification gene expression biomarkers across multiple species and ecotypes of Pacific salmonids"

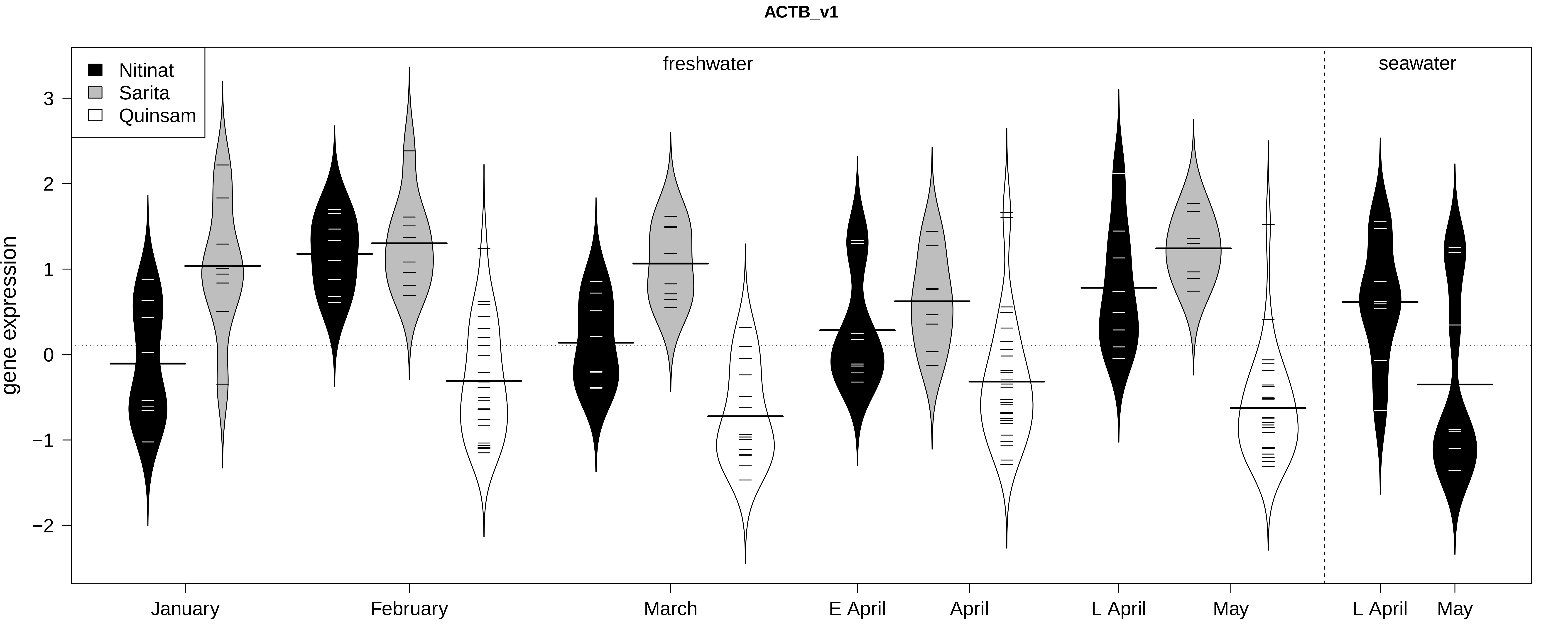
ACTB_v1
Nitinat
Sarita
Quinsam
freshwater
seawater
gene expression
3
2
1
0
−1
−2
January
February
March
E April
April
L April
May
L April
May

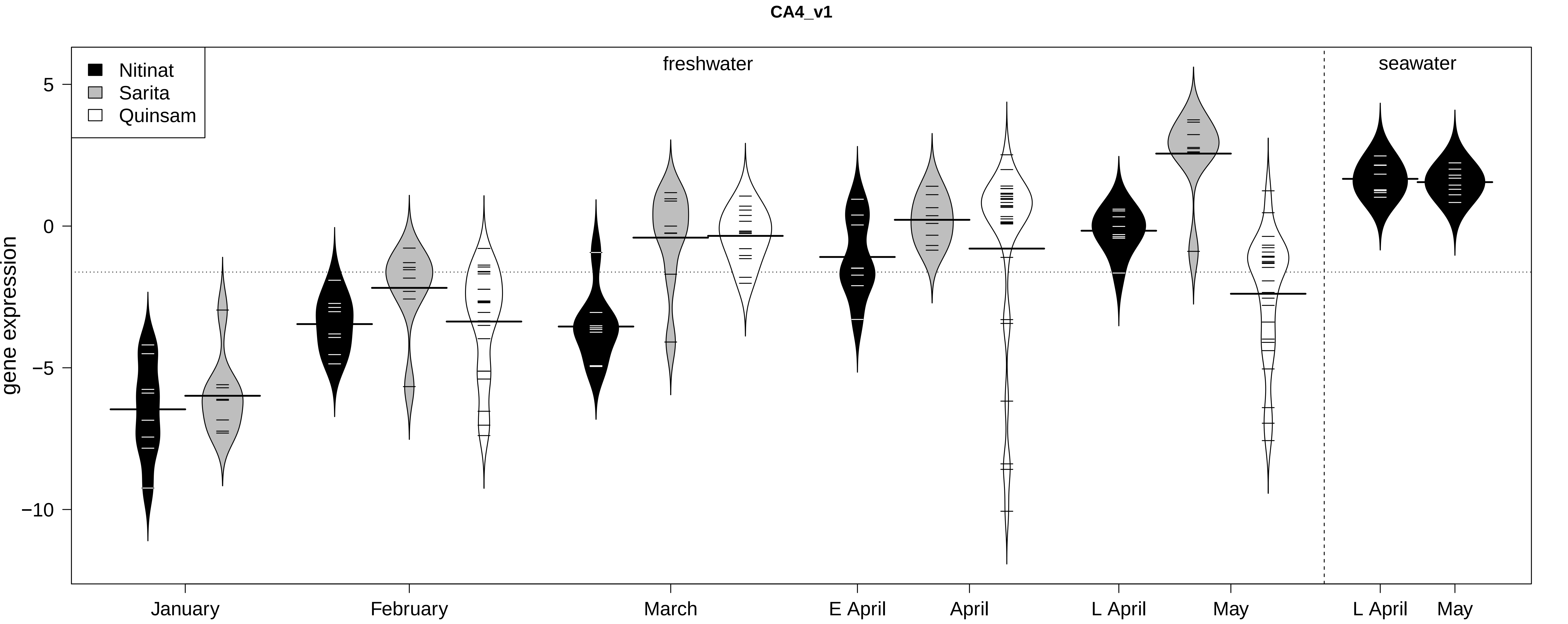
CA4_v1
Nitinat
Sarita
Quinsam
freshwater
seawater
gene expression
5
0
−5
−10
January
February
March
E April
April
L April
May
L April
May

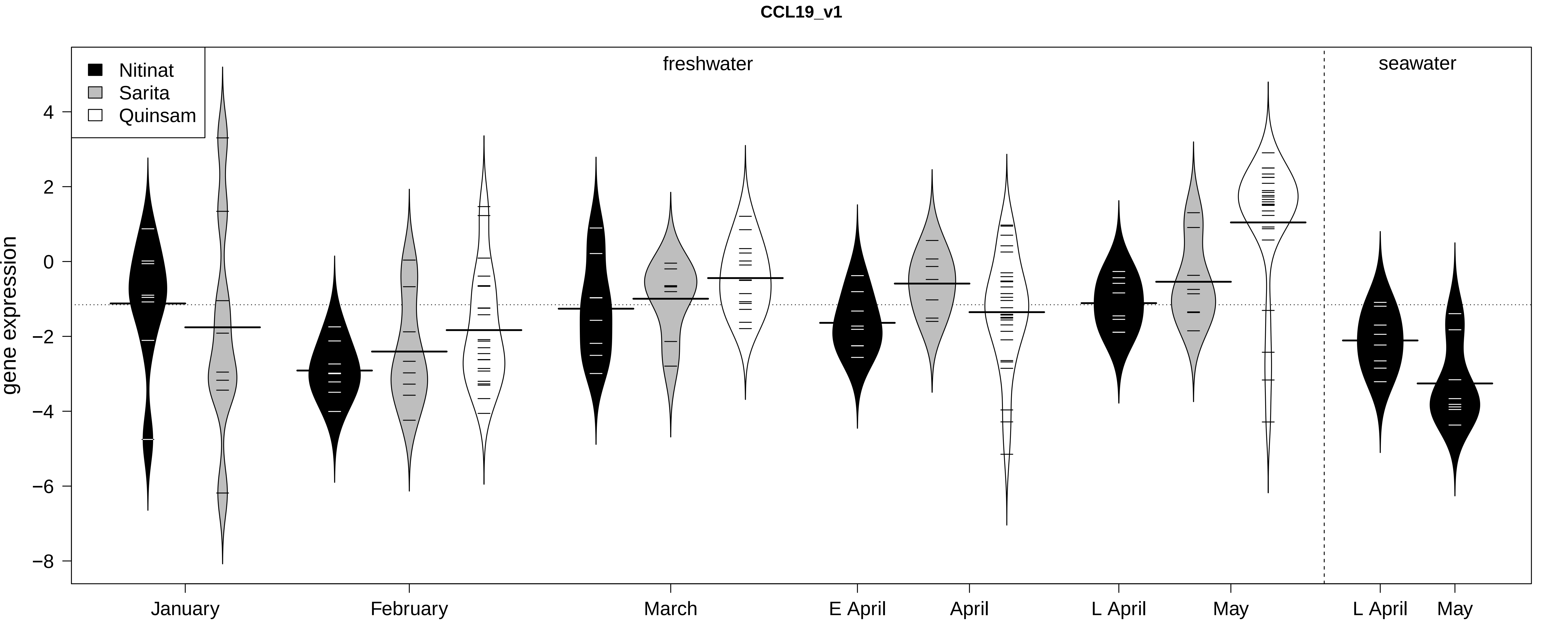

CCL19_v1
Nitinat
Sarita
Quinsam
freshwater
seawater
gene expression
4
2
0
−2
−4
−6
−8
January
February
March
E April
April
L April
May
L April
May

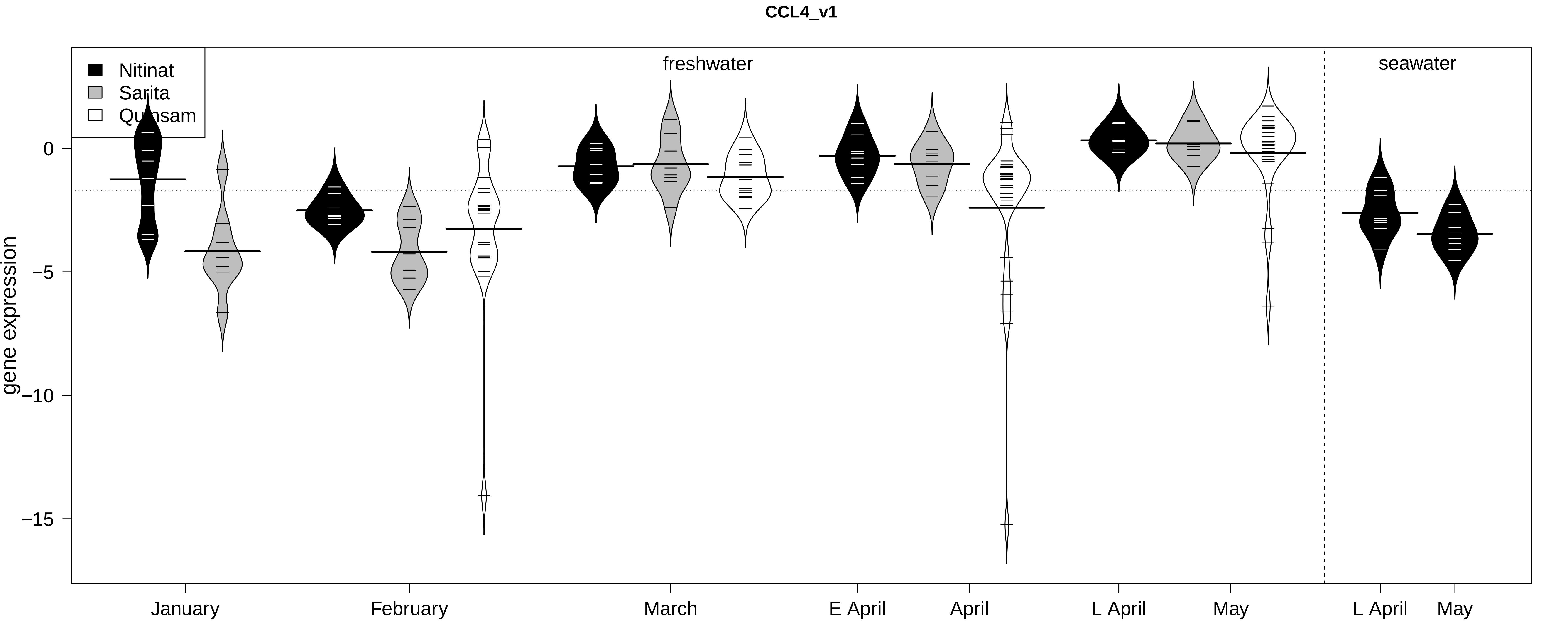
CCL4_v1
Nitinat
Sarita
Qualicum
freshwater
seawater
gene expression
0
−5
−10
−15
January
February
March
E April
April
L April
May
L April
May

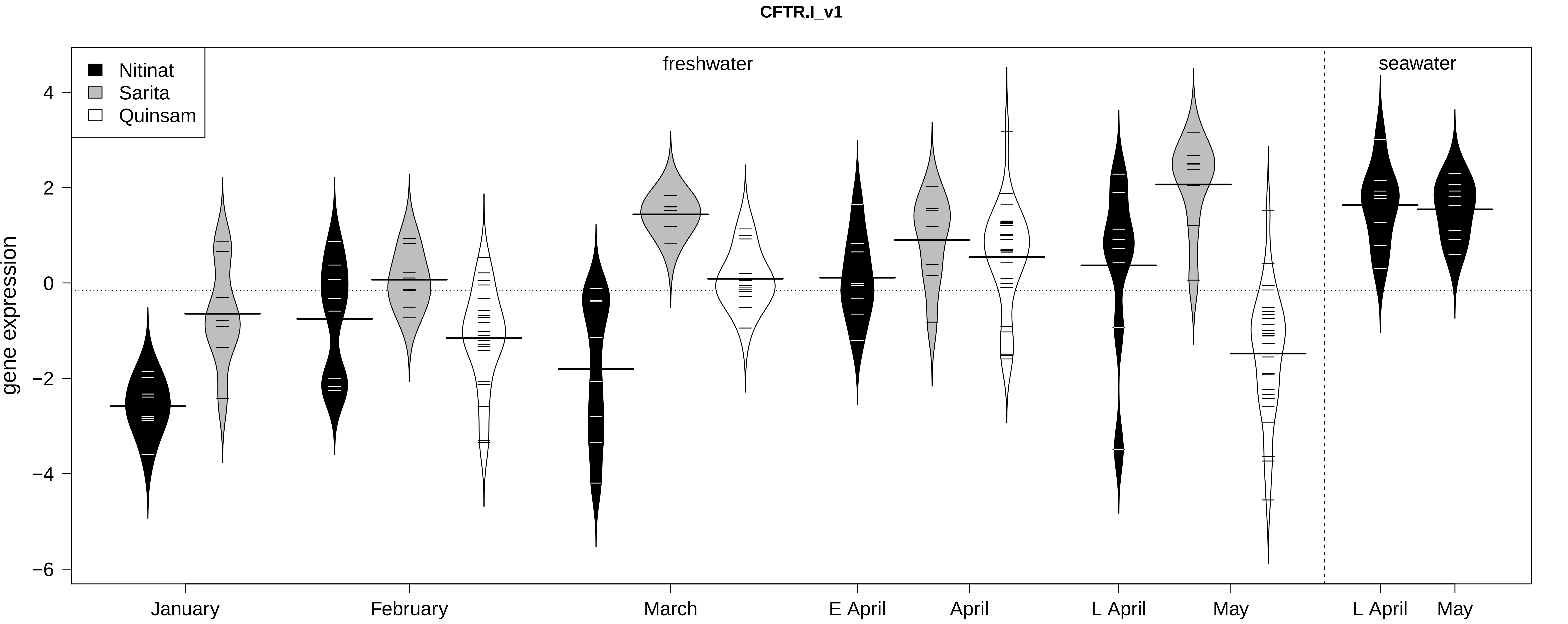

CFTR.I_v1
Nitinat
Sarita
Quinsam
freshwater
seawater
gene expression
4
2
0
−2
−4
−6
January
February
March
E April
April
L April
May
L April
May

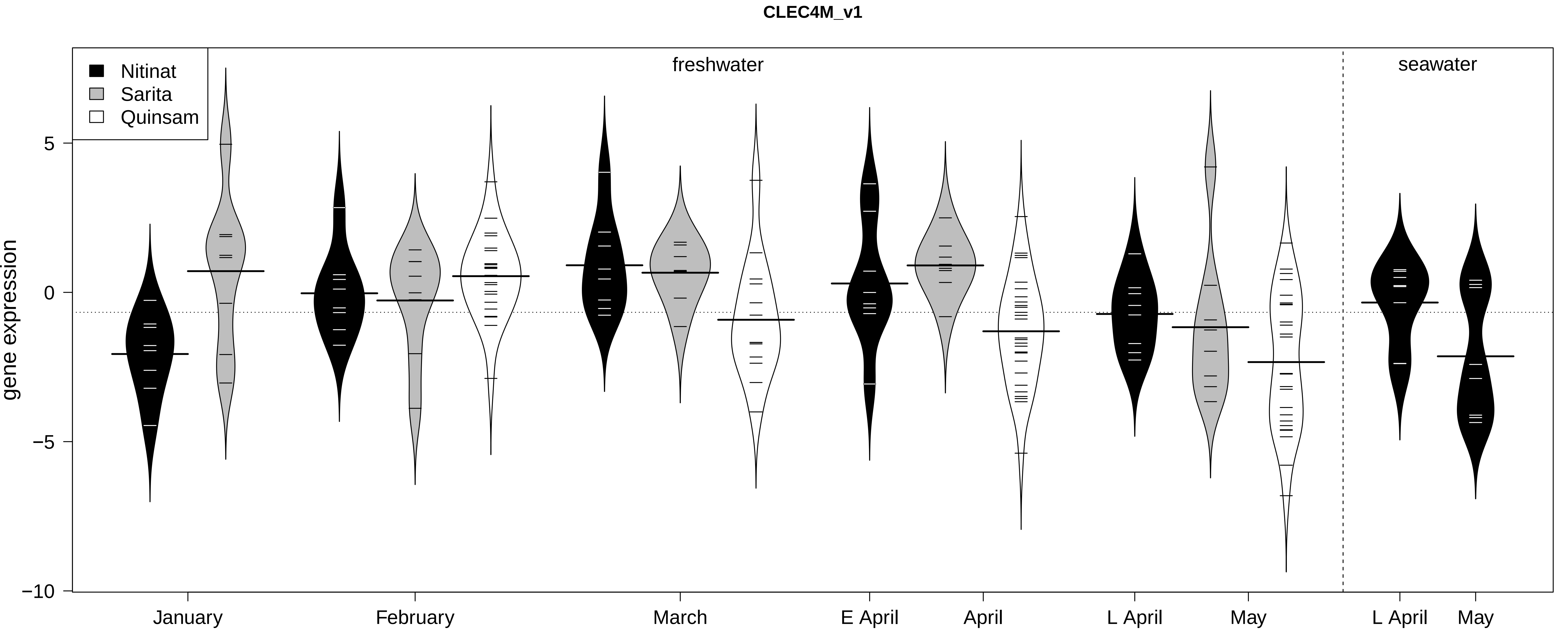

CLEC4M_v1
Nitinat
Sarita
Quinsam
freshwater
seawater
gene expression
5
0
−5
−10
January
February
March
E April
April
L April
May
L April
May

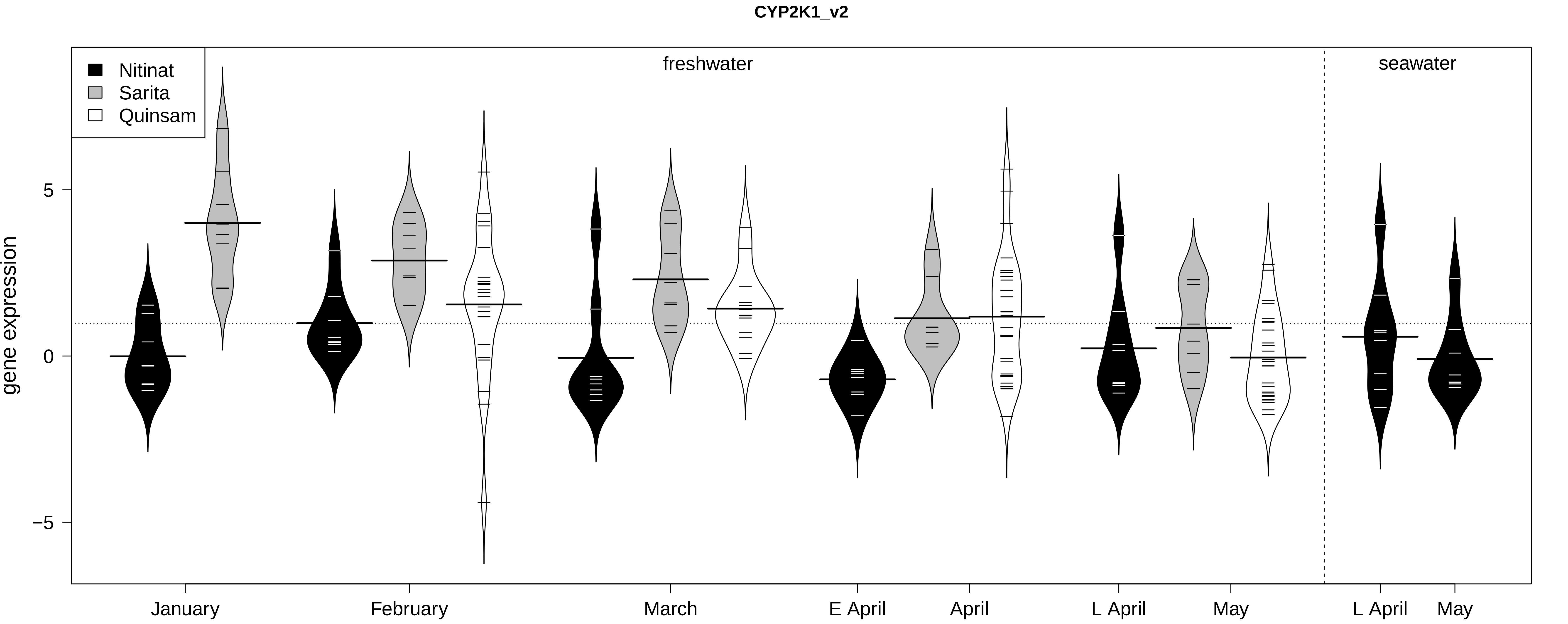
CYP2K1_v2
Nitinat
Sarita
Quinsam
freshwater
seawater
gene expression
5
0
−5
January
February
March
E April
April
L April
May
L April
May

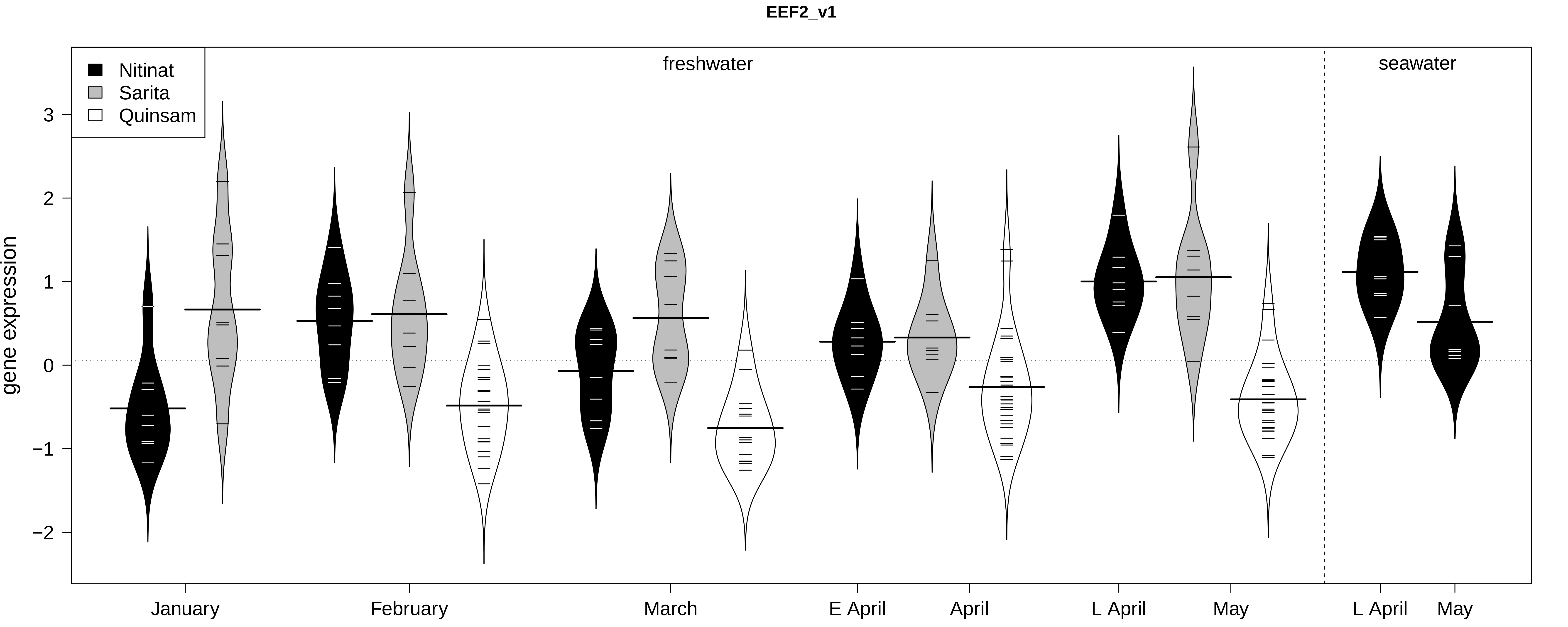

EEF2_v1
Nitinat
Sarita
Quinsam
freshwater
seawater
gene expression
3
2
1
0
−1
−2
January
February
March
E April
April
L April
May
L April
May

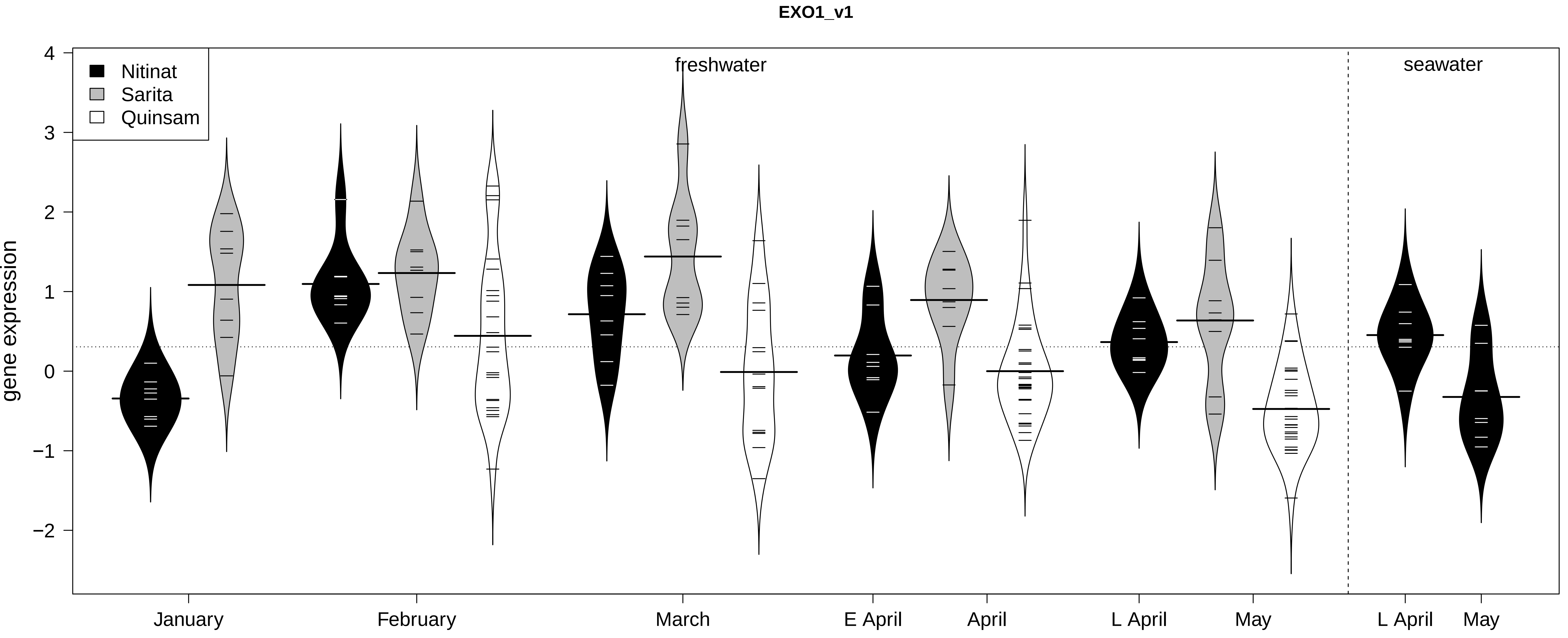

EXO1_v1
Nitinat
Sarita
Quinsam
freshwater
seawater
gene expression
4
3
2
1
0
−1
−2
January
February
March
E April
April
L April
May
L April
May

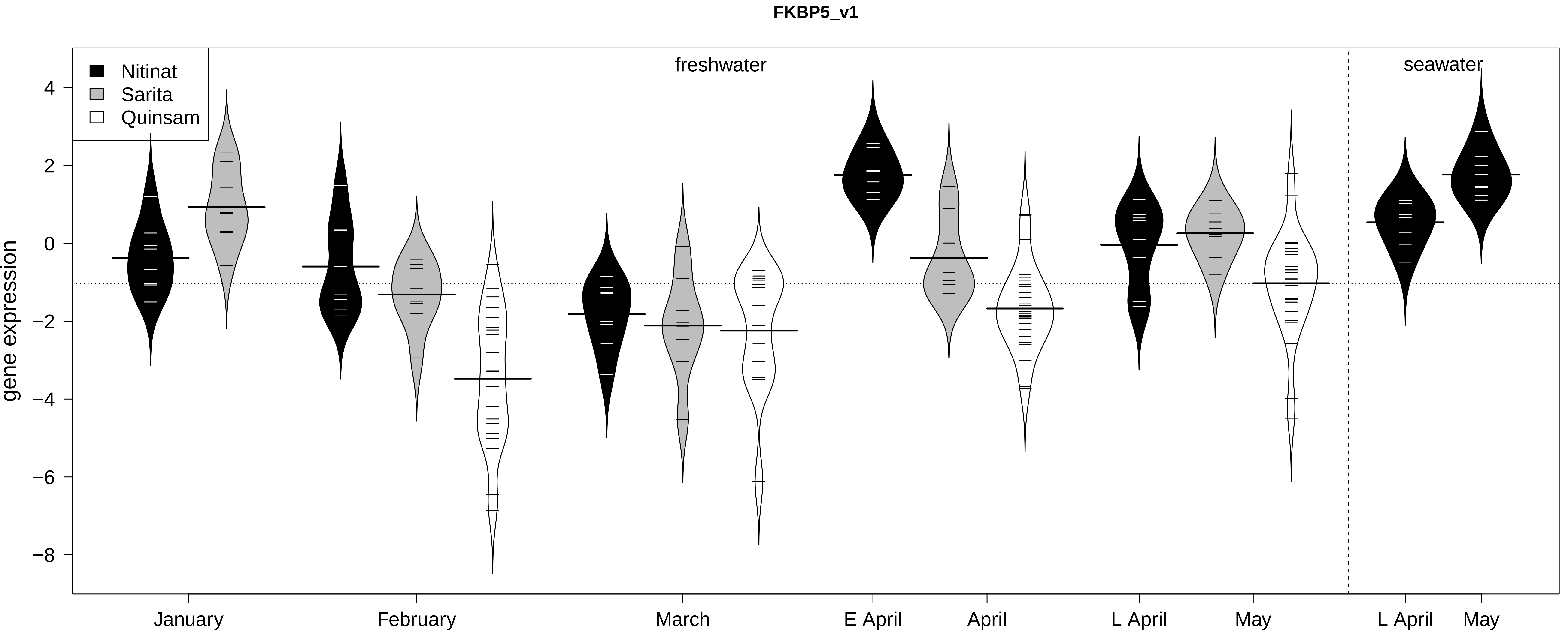
FKBP5_v1
Nitinat
Sarita
Quinsam
freshwater
seawater
gene expression
4
2
0
−2
−4
−6
−8
January
February
March
E April
April
L April
May
L April
May

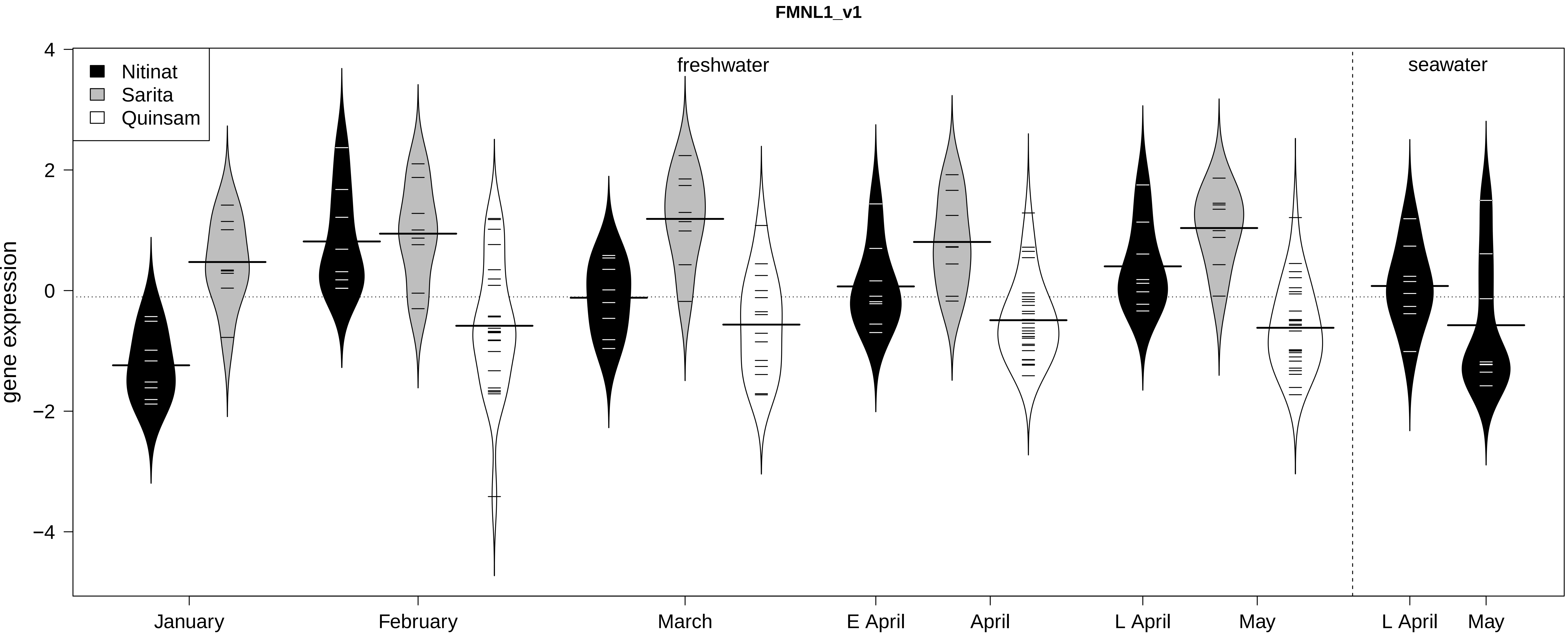
FMNL1_v1
Nitinat
Sarita
Quinsam
freshwater
seawater
gene expression
4
2
0
−2
−4
January
February
March
E April
April
L April
May
L April
May

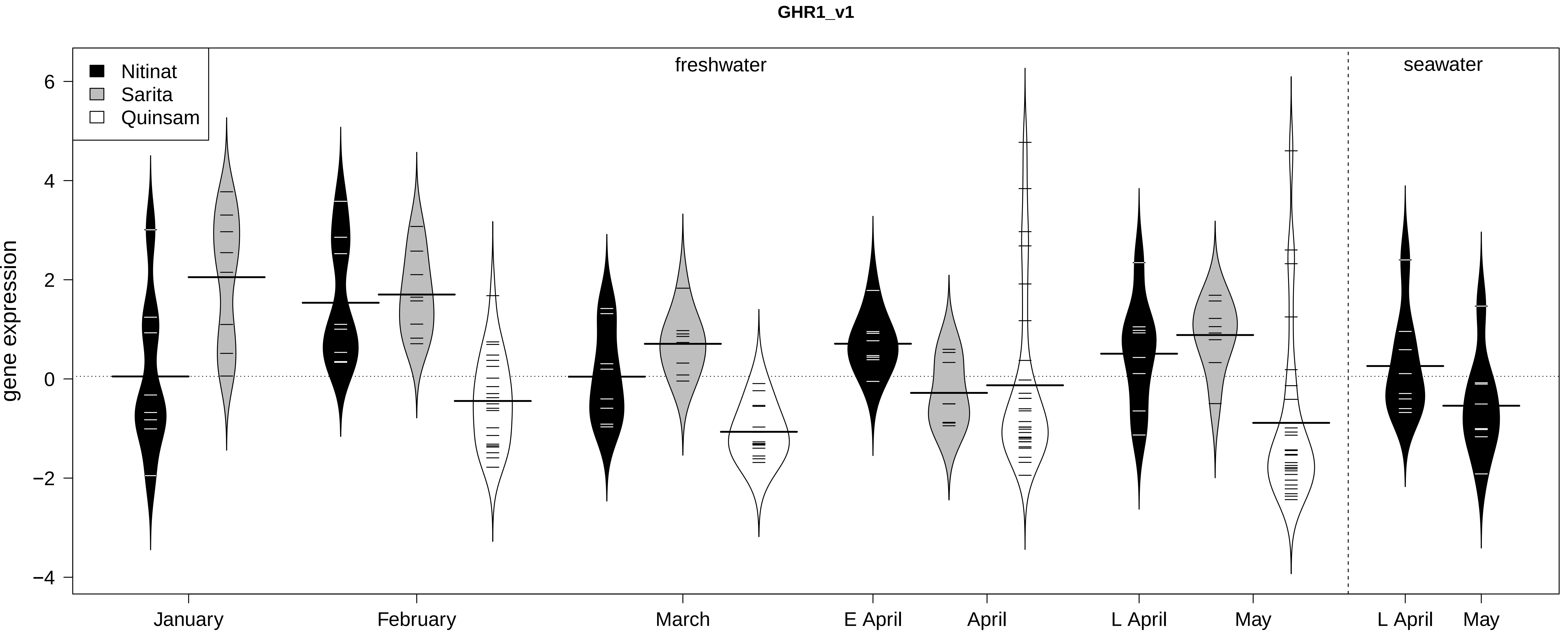

GHR1_v1
Nitinat
Sarita
Quinsam
freshwater
seawater
gene expression
6
4
2
0
−2
−4
January
February
March
E April
April
L April
May
L April
May

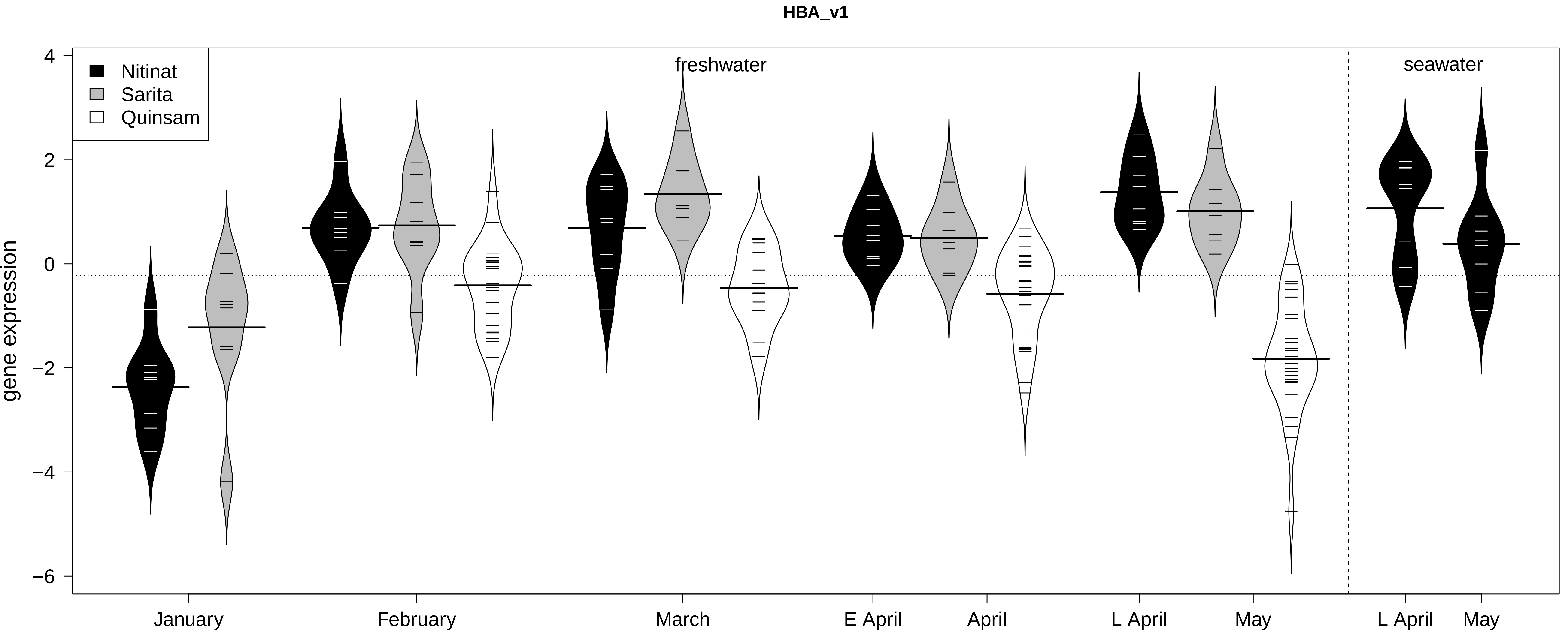

HBA_v1
Nitinat
Sarita
Quinsam
freshwater
seawater
gene expression
4
2
0
−2
−4
−6
January
February
March
E April
April
L April
May
L April
May

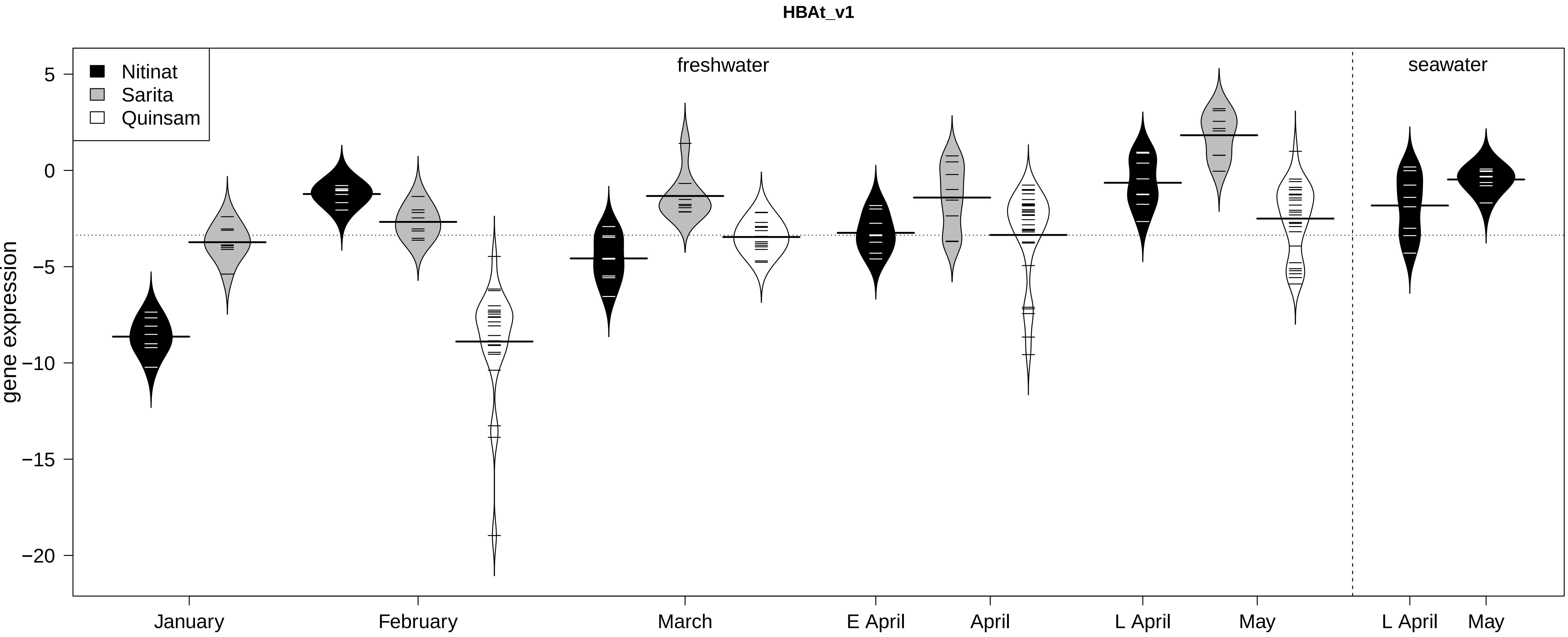

HBAt_v1
freshwater
seawater
Nitinat
Sarita
Quinsam
gene expression
5
0
−5
−10
−15
−20
January
February
March
E April
April
L April
May
L April
May

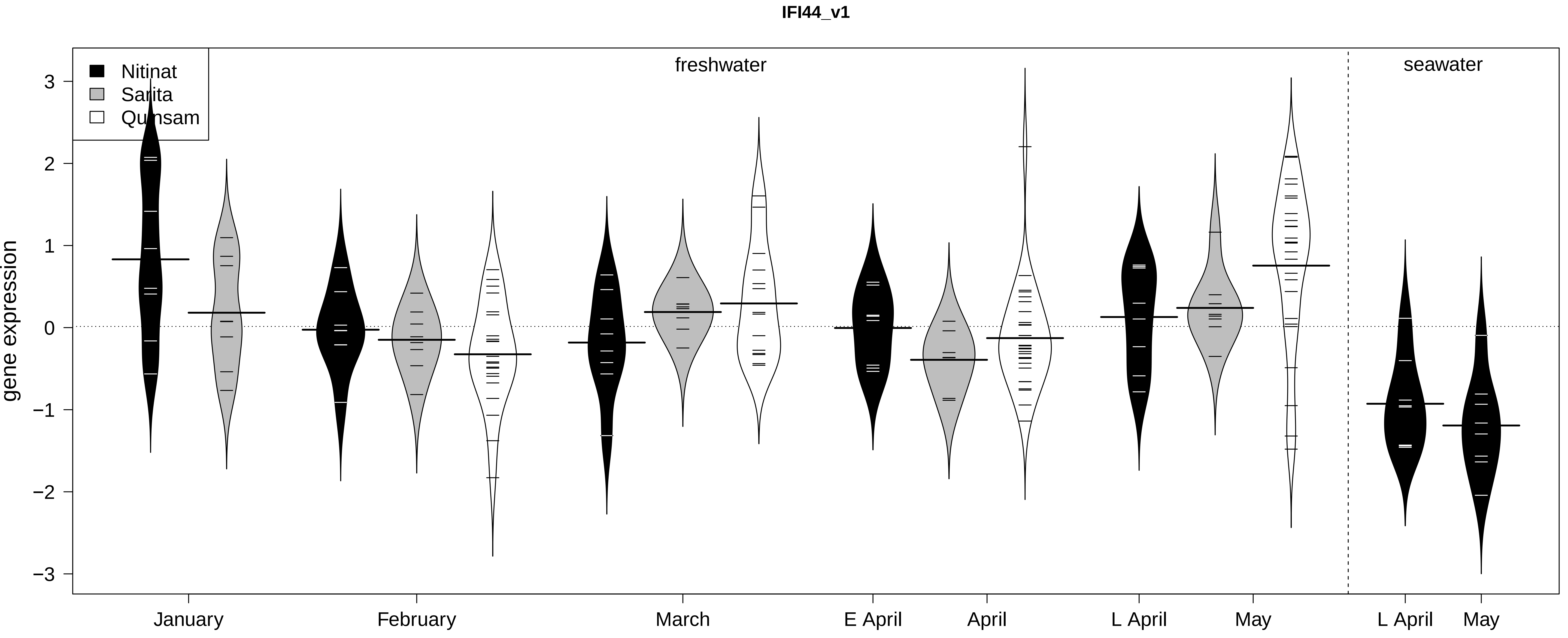
IFI44_v1
Nitinat
Sarita
Qumsam
freshwater
seawater
gene expression
3
2
1
0
−1
−2
−3
January
February
March
E April
April
L April
May
L April
May

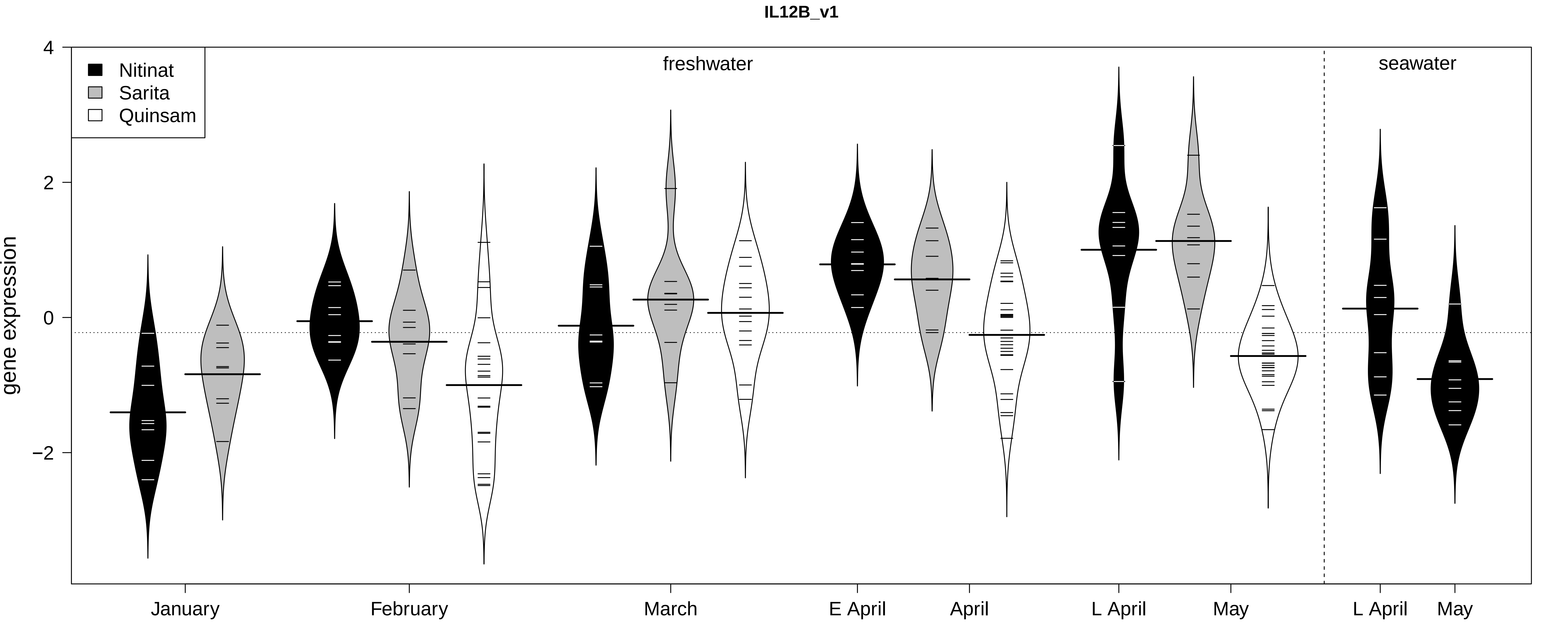
IL12B_v1
Nitinat
Sarita
Quinsam
freshwater
seawater
gene expression
4
2
0
−2
January
February
March
E April
April
L April
May
L April
May

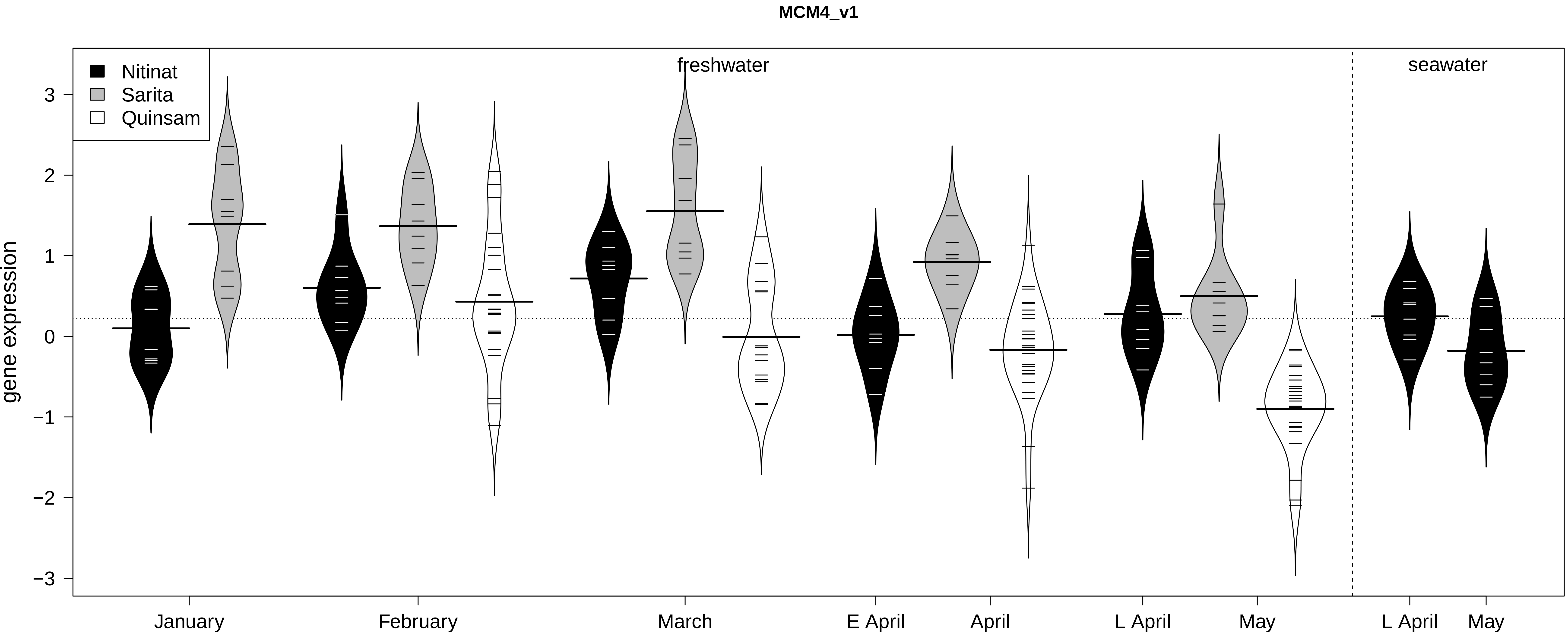

MCM4_v1
Nitinat
Sarita
Quinsam
freshwater
seawater
gene expression
3
2
1
0
−1
−2
−3
January
February
March
E April
April
L April
May
L April
May

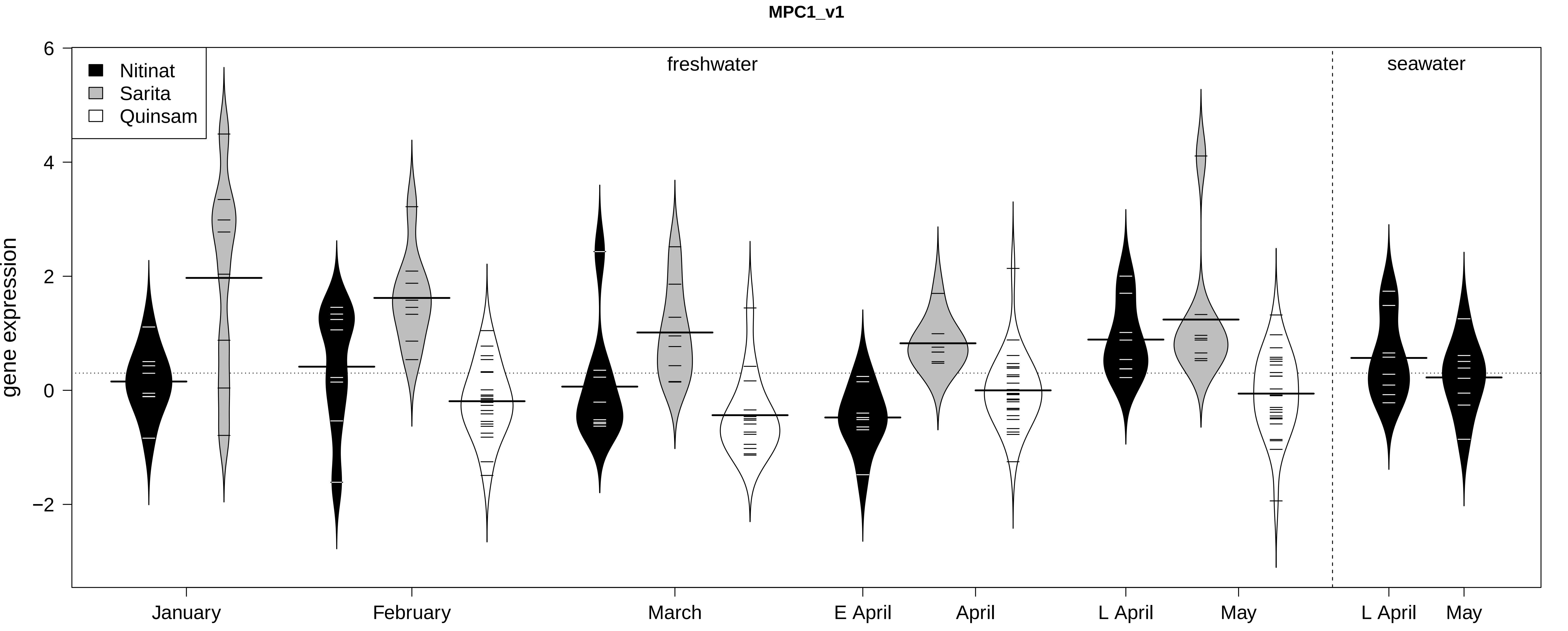

MPC1_v1
Nitinat
Sarita
Quinsam
freshwater
seawater
gene expression
6
4
2
0
−2
January
February
March
E April
April
L April
May
L April
May

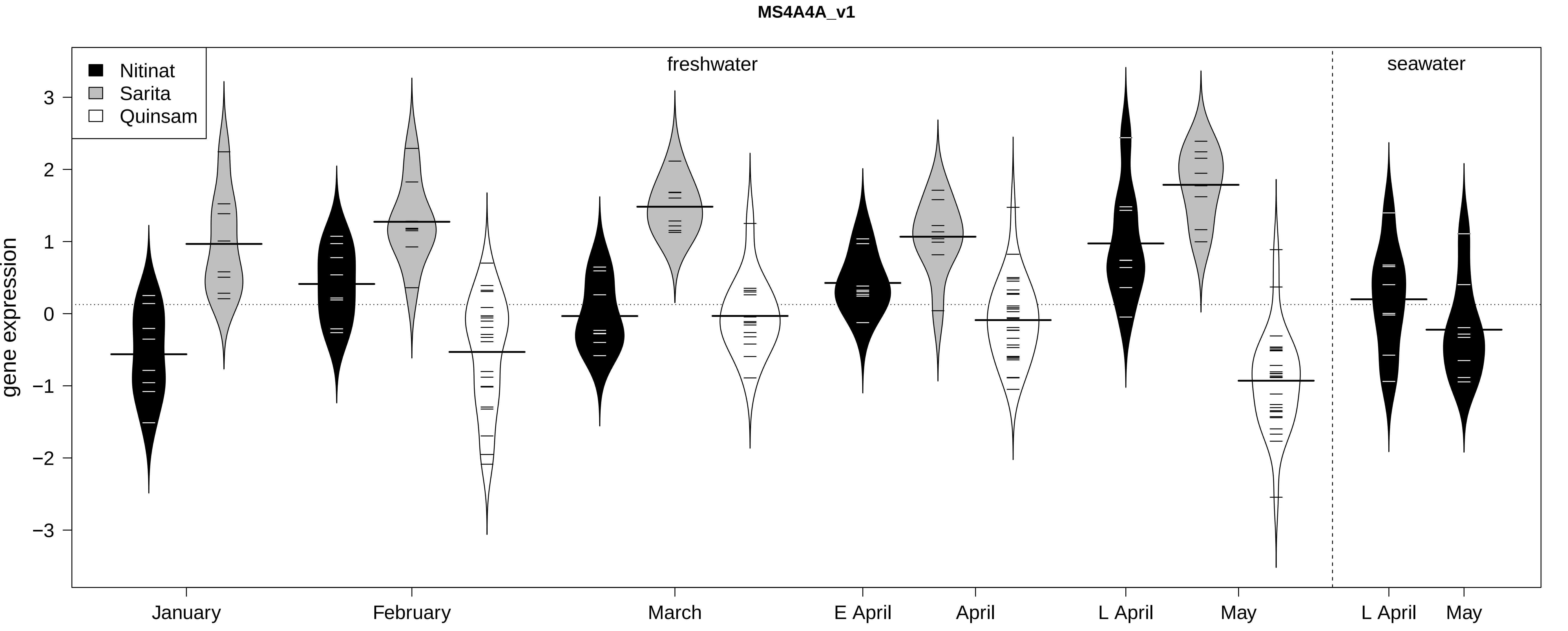

MS4A4A_v1
Nitinat
Sarita
Quinsam
freshwater
seawater
gene expression
3
2
1
0
−1
−2
−3
January
February
March
E April
April
L April
May
L April
May

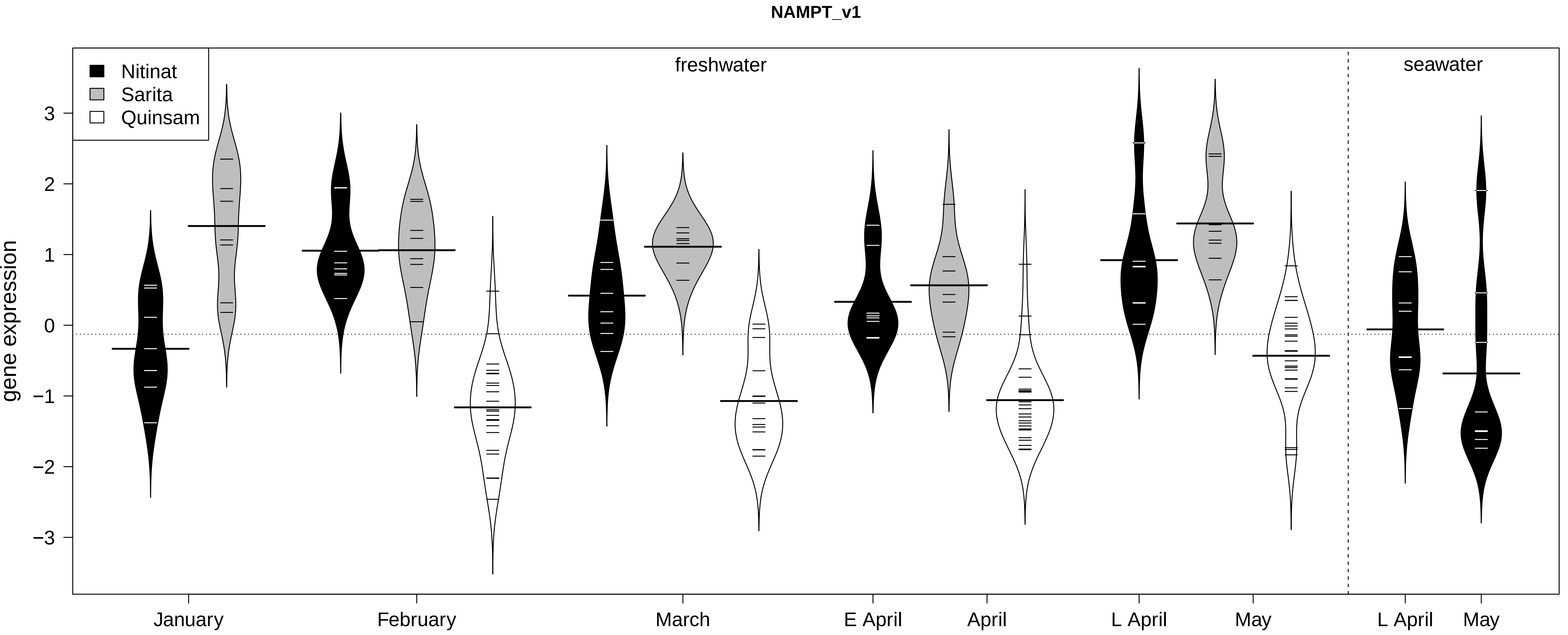

NAMPT_v1
Nitinat
Sarita
Quinsam
freshwater
seawater
gene expression
3
2
1
0
−1
−2
−3
January
February
March
E April
April
L April
May
L April
May

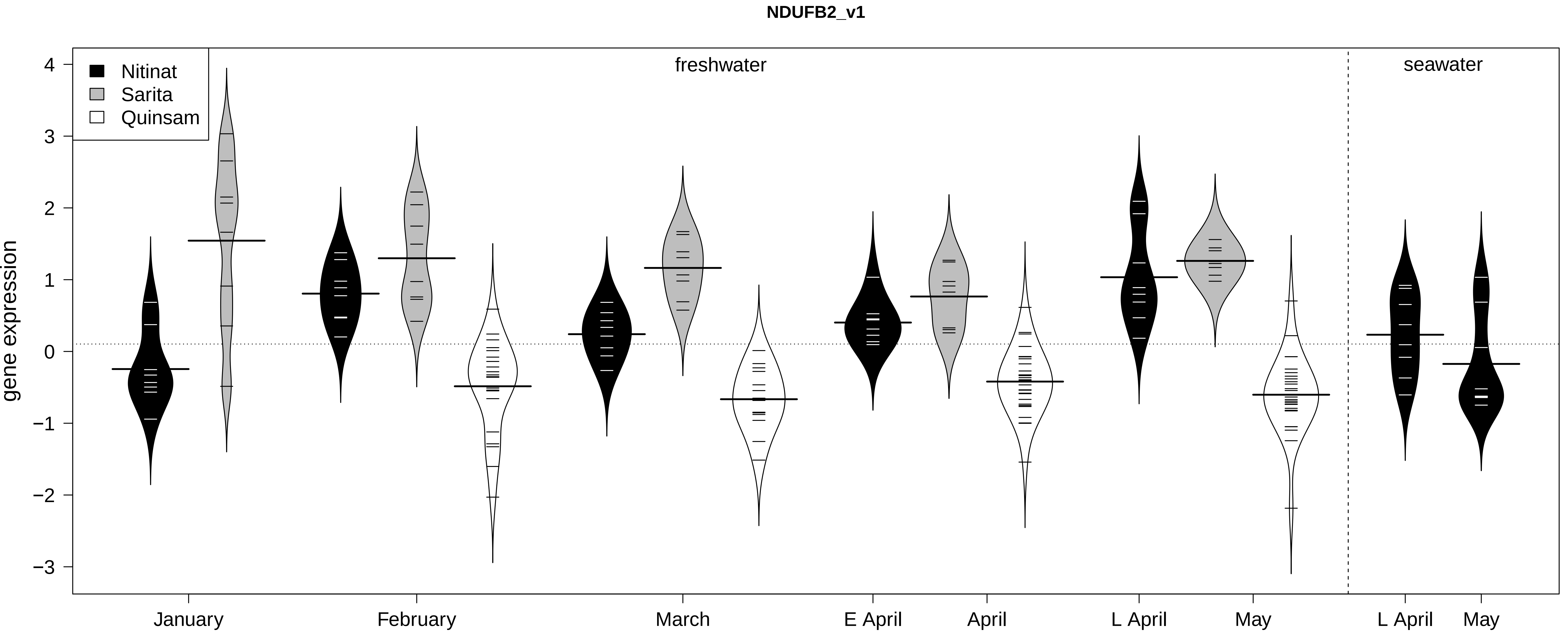

NDUFB2_v1
Nitinat
Sarita
Quinsam
freshwater
seawater
gene expression
4
3
2
1
0
−1
−2
−3
January
February
March
E April
April
L April
May
L April
May

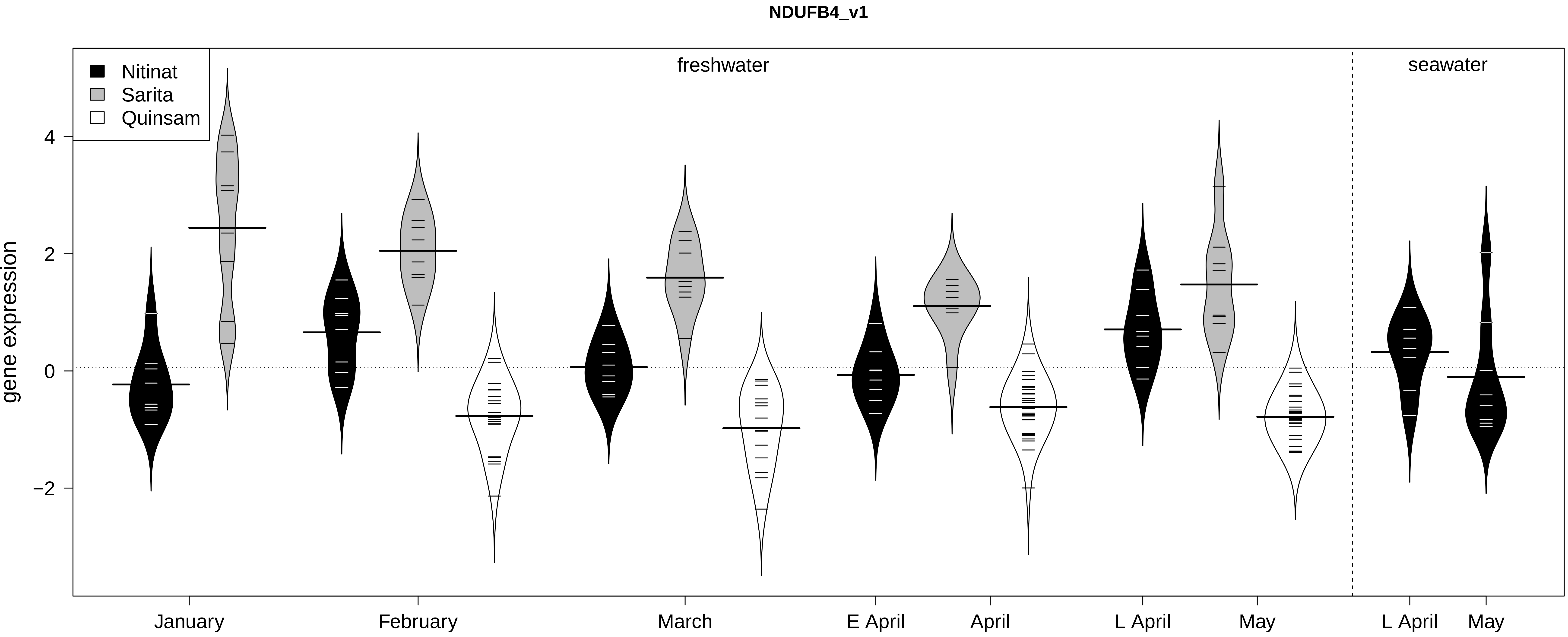

NDUFB4_v1
Nitinat
Sarita
Quinsam
freshwater
seawater
gene expression
4
2
0
−2
January
February
March
E April
April
L April
May
L April
May

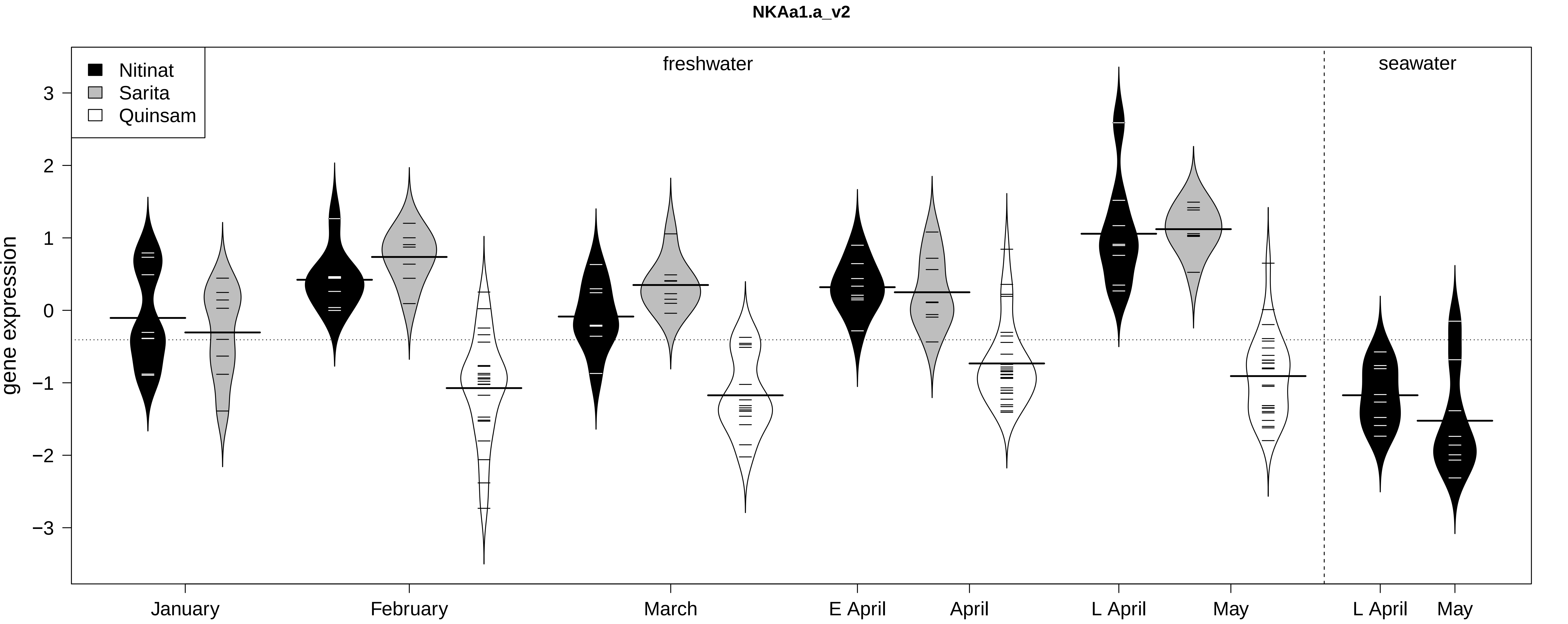

NKAa1.a_v2
Nitinat
Sarita
Quinsam
freshwater
seawater
gene expression
3
2
1
0
−1
−2
−3
January
February
March
E April
April
L April
May
L April
May

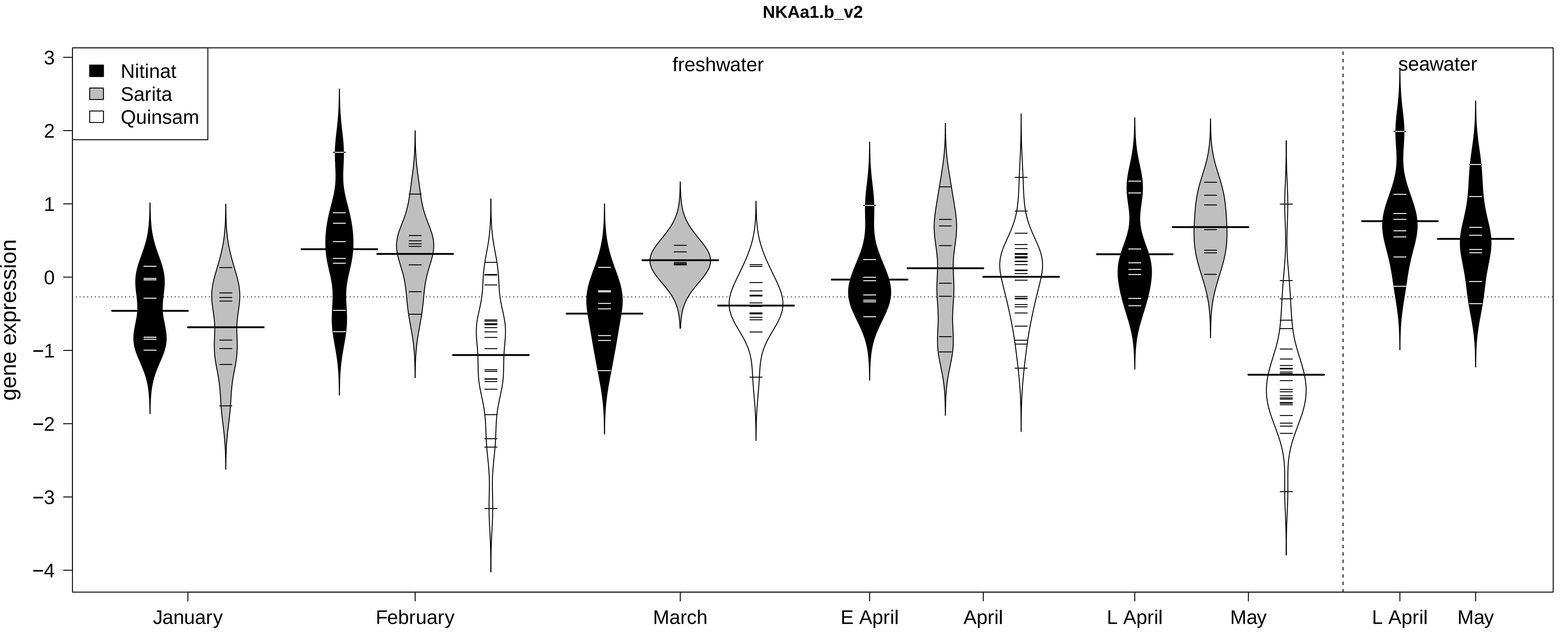
NKAa1.b_v2
Nitinat
Sarita
Quinsam
freshwater
seawater
gene expression
3
2
1
0
−1
−2
−3
−4
January
February
March
E April
April
L April
May
L April
May

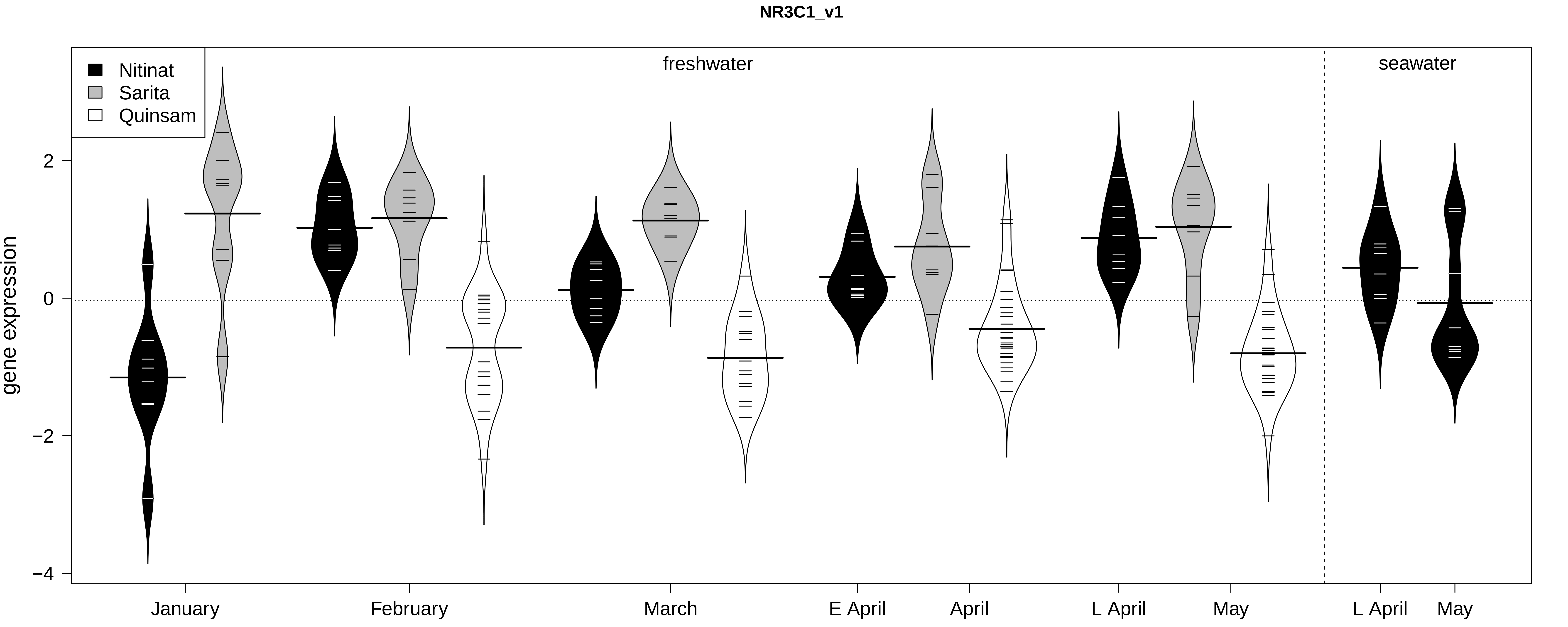
NR3C1_v1
Nitinat
Sarita
Quinsam
freshwater
seawater
gene expression
2
0
−2
−4
January
February
March
E April
April
L April
May
L April
May

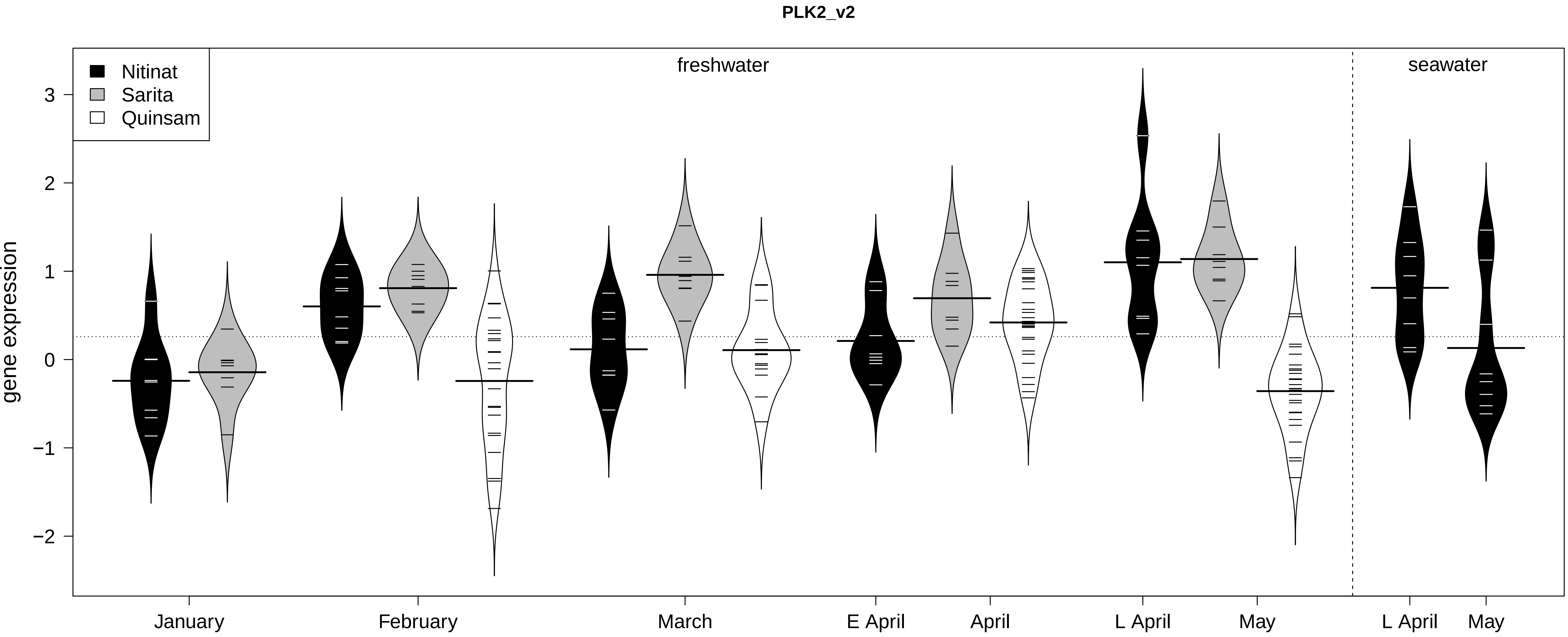
PLK2_v2
Nitinat
Sarita
Quinsam
freshwater
seawater
gene expression
3
2
1
0
−1
−2
January
February
March
E April
April
L April
May
L April
May

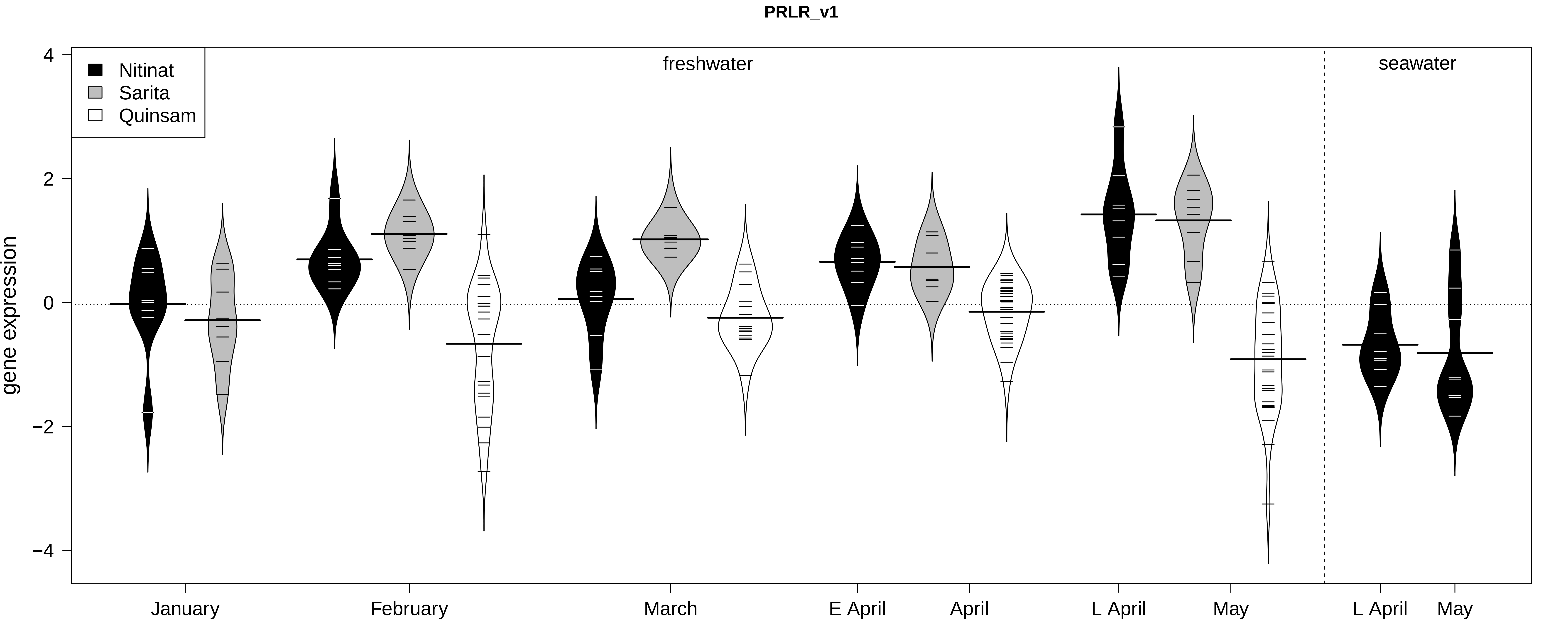

PRLR_v1
Nitinat
Sarita
Quinsam
freshwater
seawater
gene expression
4
2
0
−2
−4
January
February
March
E April
April
L April
May
L April
May

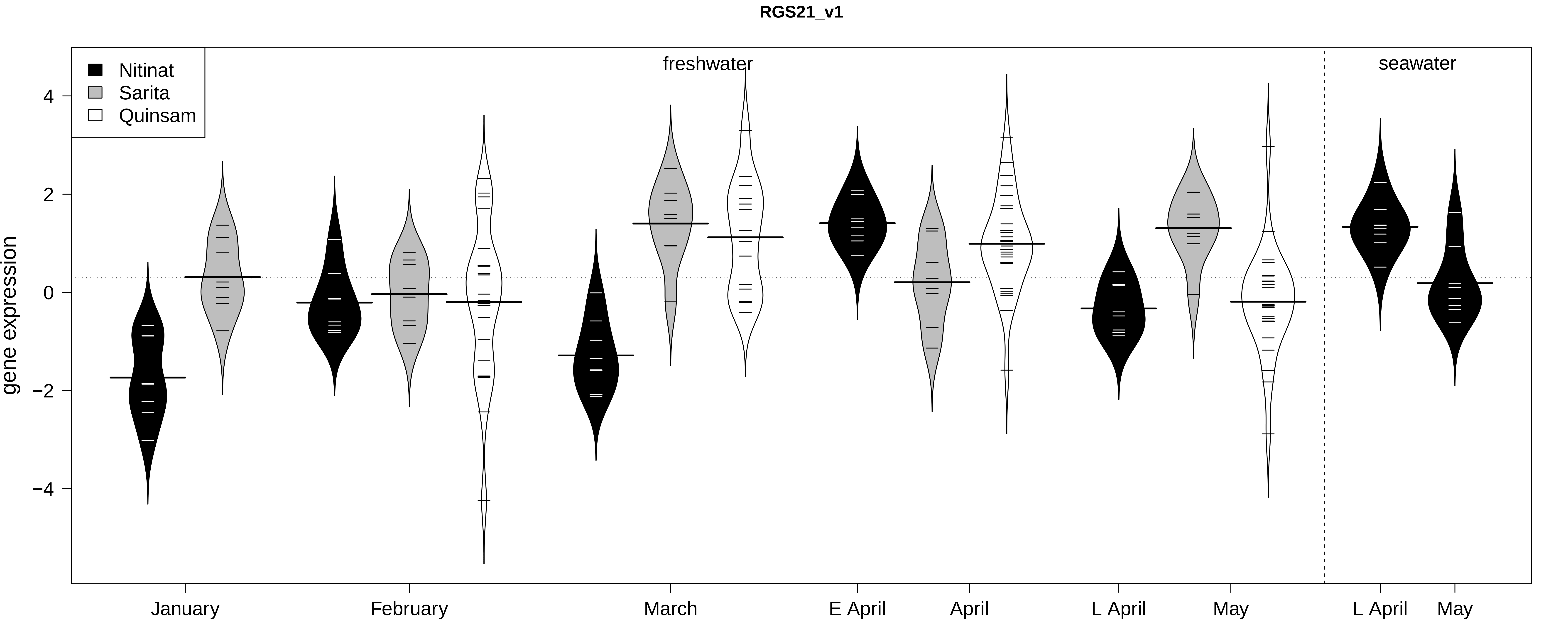

RGS21_v1
Nitinat
Sarita
Quinsam
freshwater
seawater
gene expression
4
2
0
−2
−4
January
February
March
E April
April
L April
May
L April
May

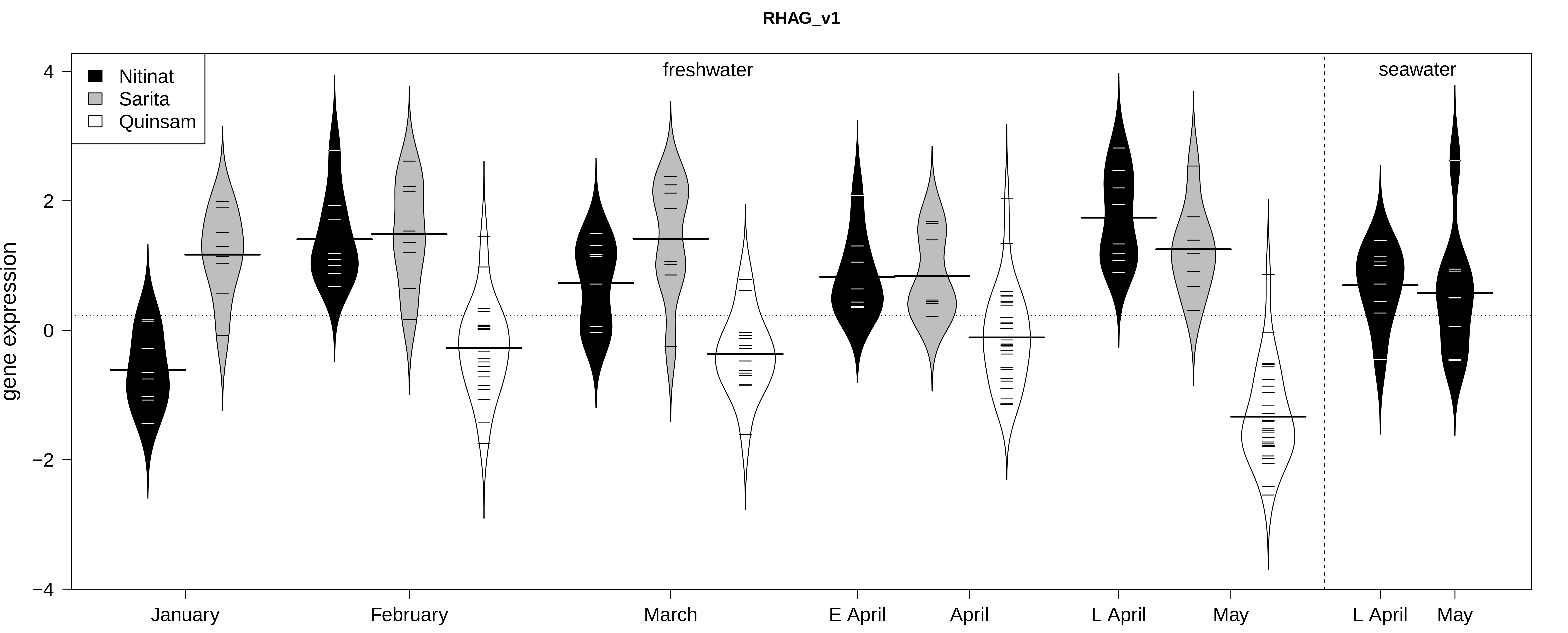

RHAG_v1
Nitinat
Sarita
Quinsam
freshwater
seawater
gene expression
4
2
0
−2
−4
January
February
March
E April
April
L April
May
L April
May

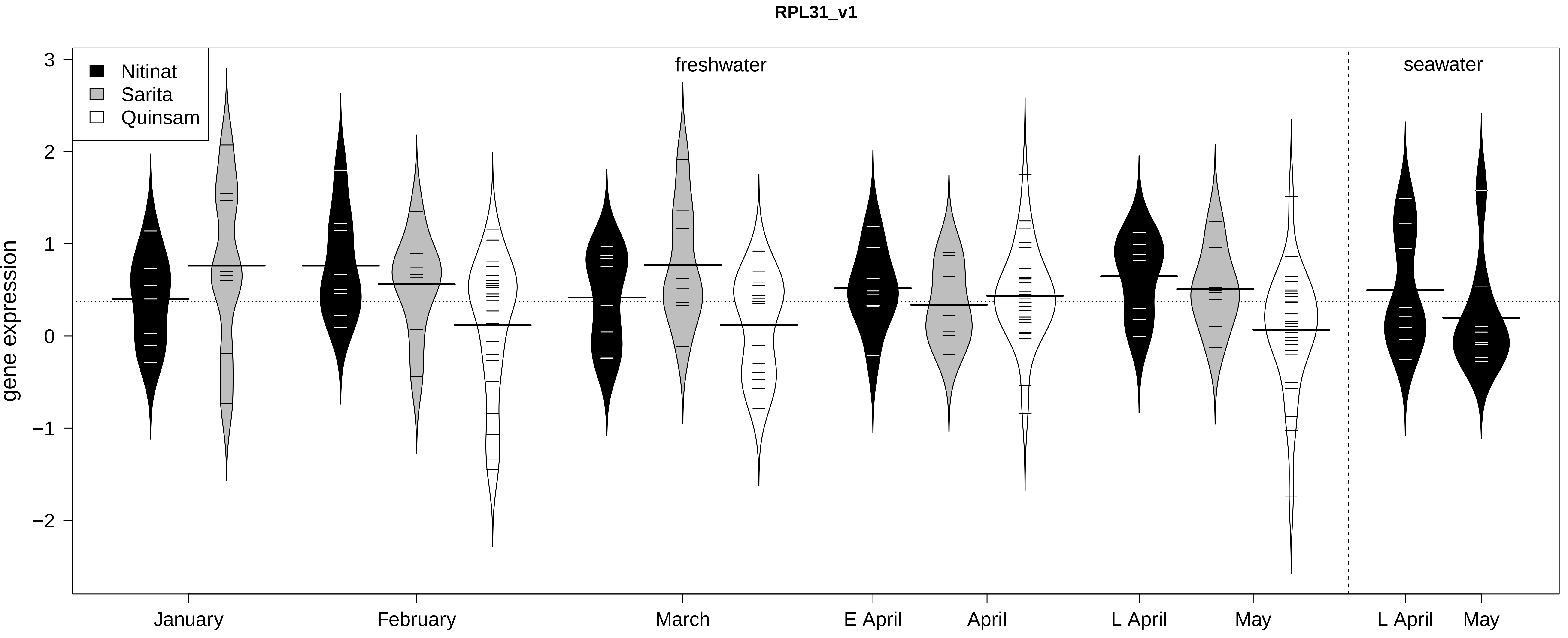

RPL31_v1
Nitinat
Sarita
Quinsam
freshwater
seawater
gene expression
3
2
1
0
−1
−2
January
February
March
E April
April
L April
May
L April
May

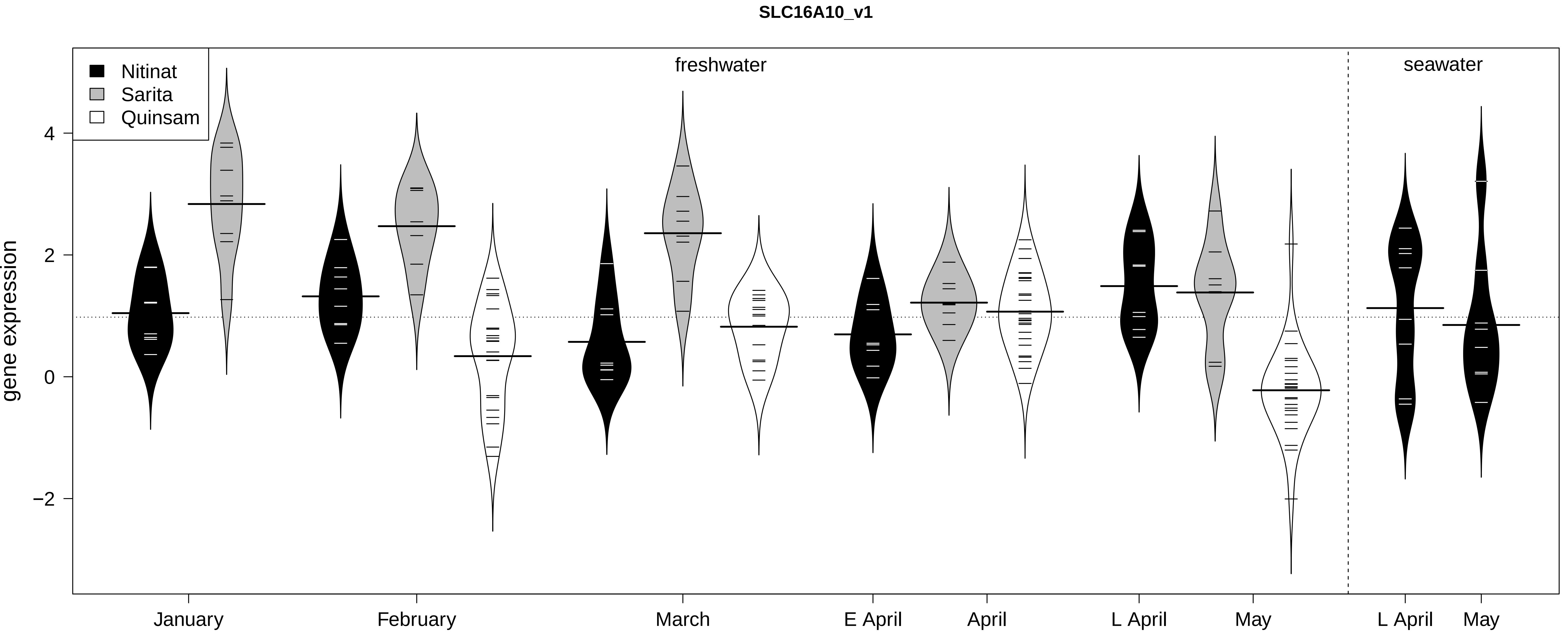
SLC16A10_v1
Nitinat
Sarita
Quinsam
freshwater
seawater
gene expression
4
2
0
−2
January
February
March
E April
April
L April
May
L April
May

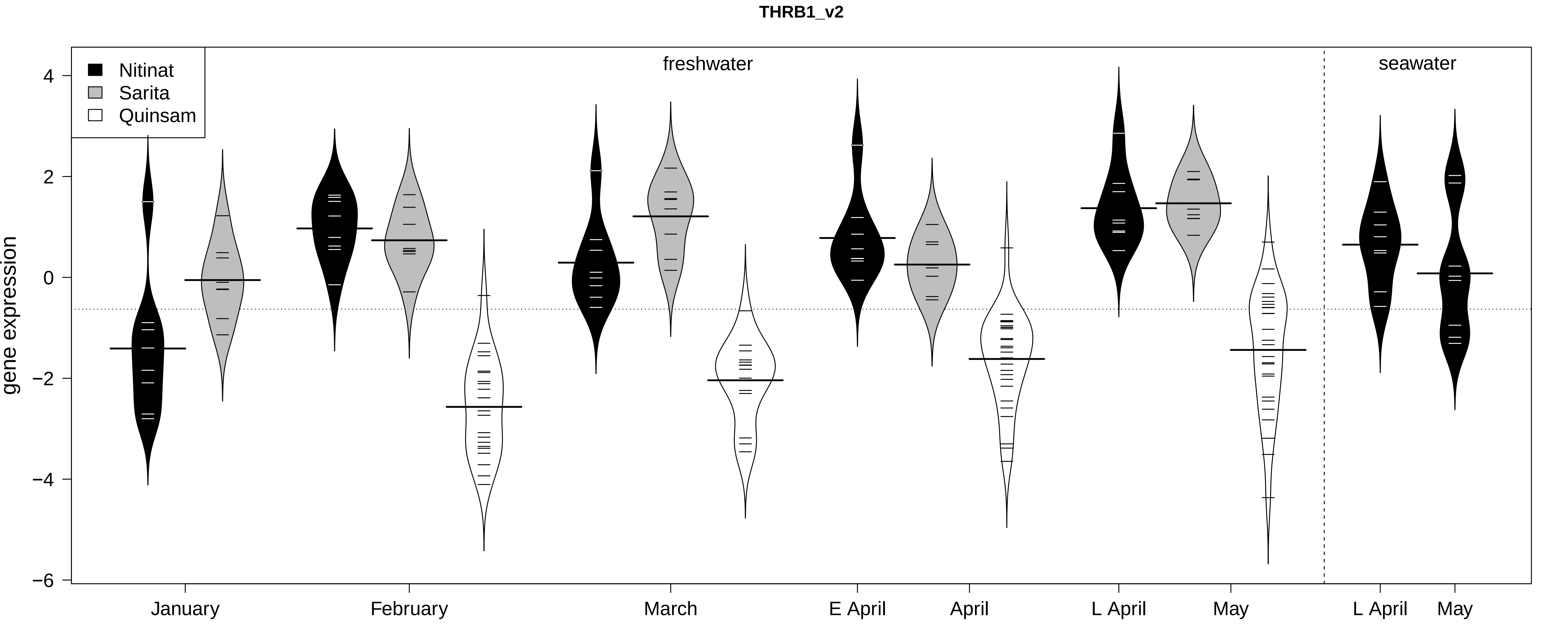
THRB1_v2
Nitinat
Sarita
Quinsam
freshwater
seawater
gene expression
4
2
0
−2
−4
−6
January
February
March
E April
April
L April
May
L April
May

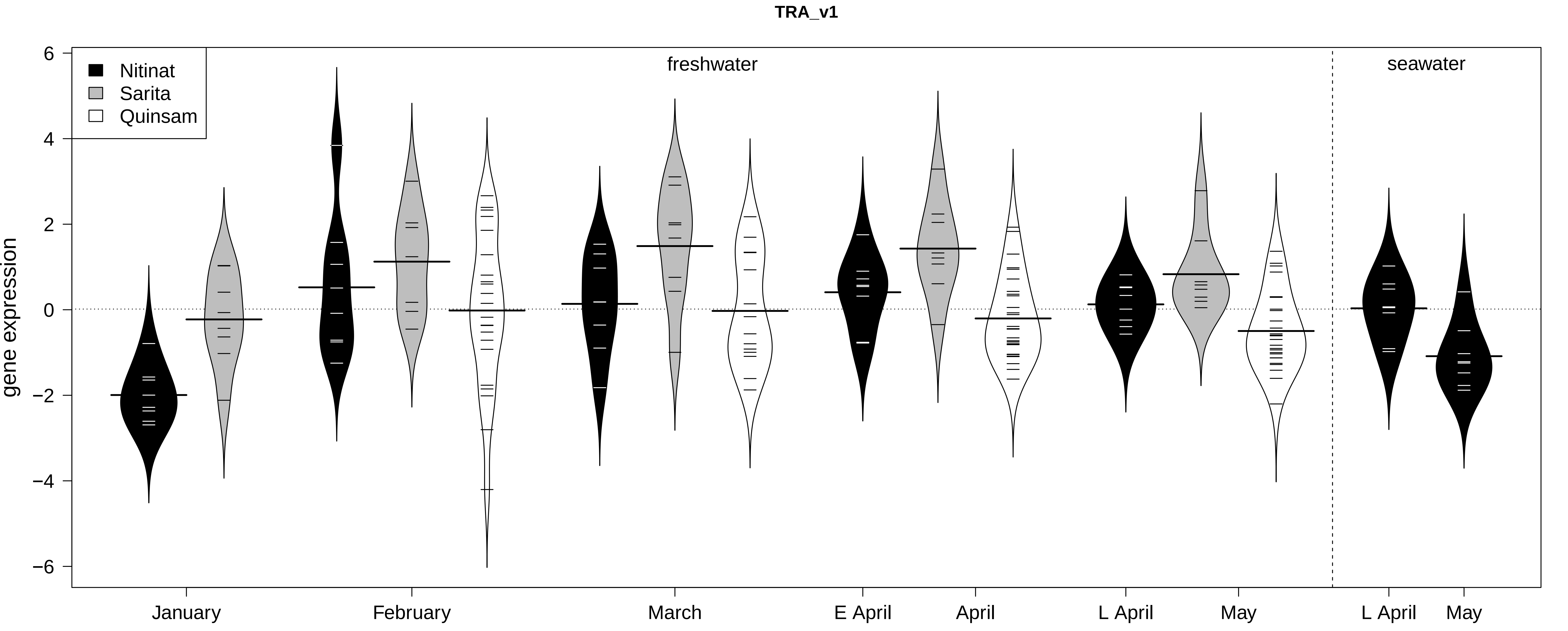

TRA_v1
Nitinat
Sarita
Quinsam
freshwater
seawater
gene expression
6
4
2
0
−2
−4
−6
January
February
March
E April
April
L April
May
L April
May

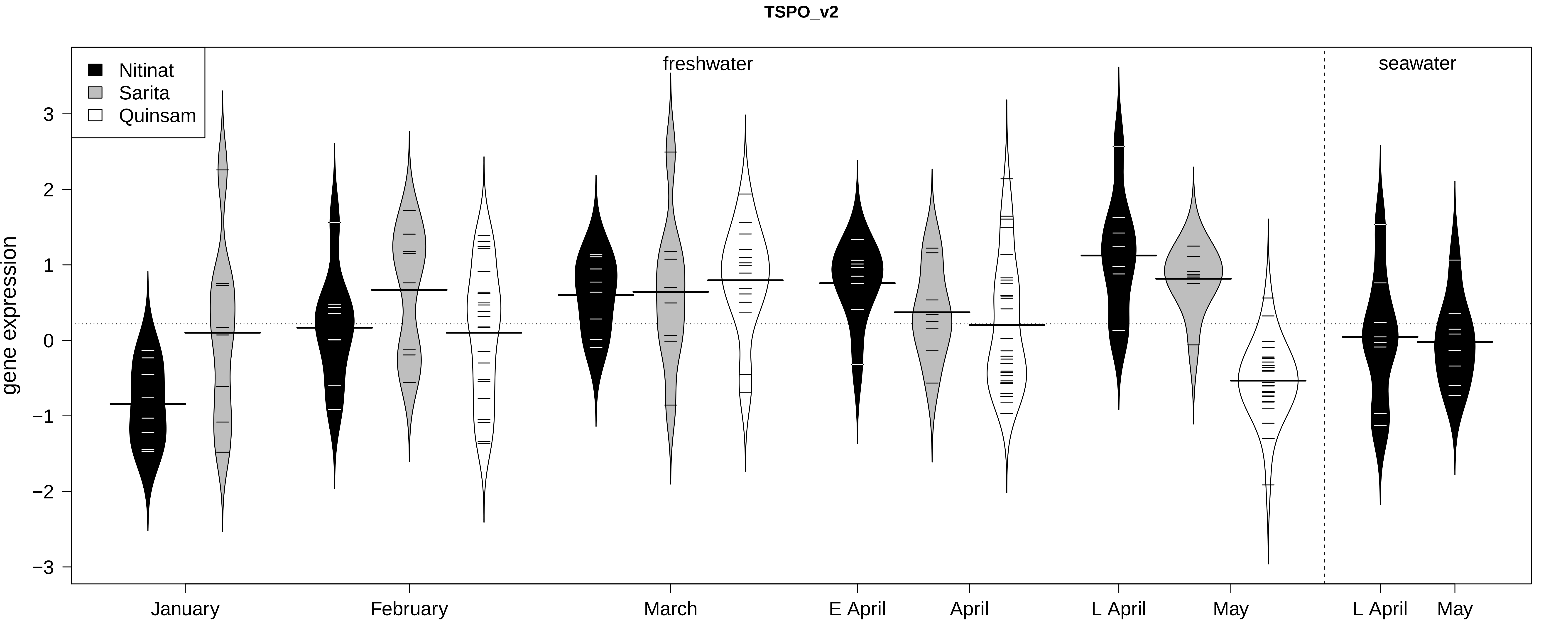

TSPO_v2
Nitinat
Sarita
Quinsam
freshwater
seawater
gene expression
3
2
1
0
−1
−2
−3
January
February
March
E April
April
L April
May
L April
May

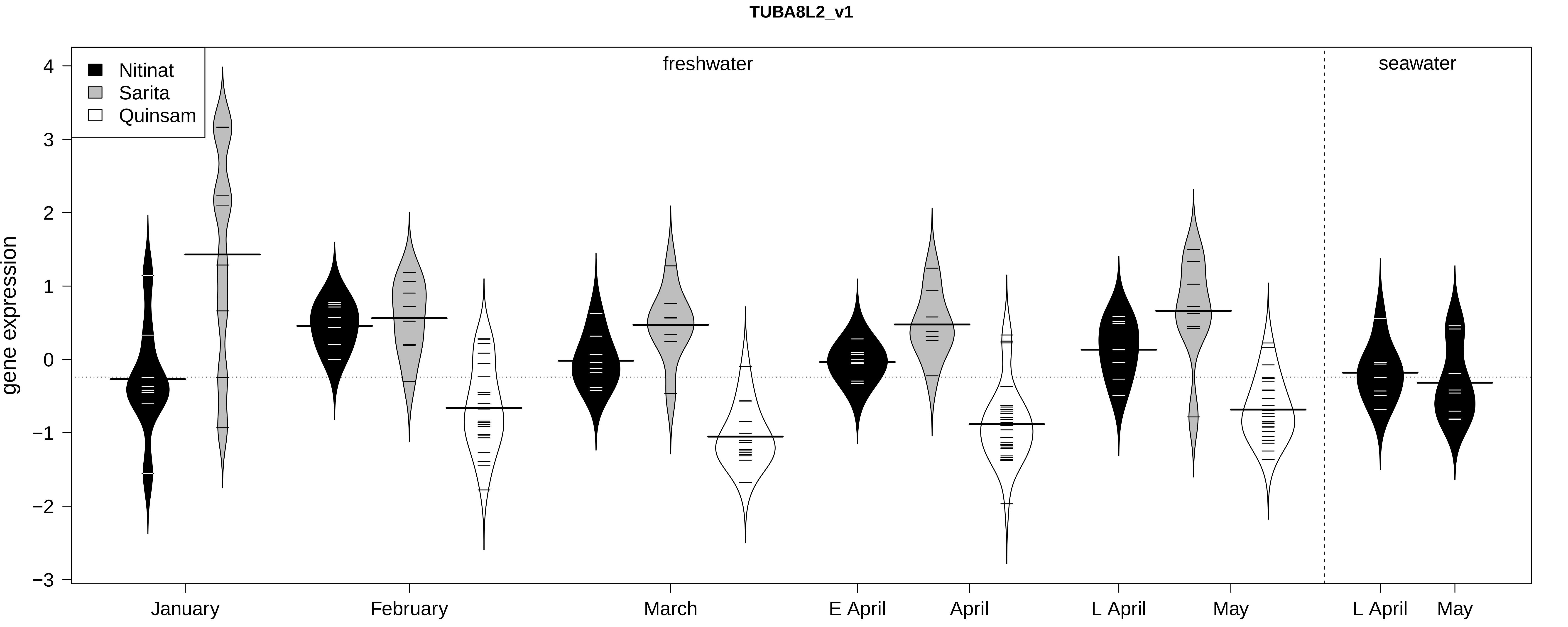

TUBA8L2_v1
Nitinat
Sarita
Quinsam
freshwater
seawater
gene expression
4
3
2
1
0
−1
−2
−3
January
February
March
E April
April
L April
May
L April
May

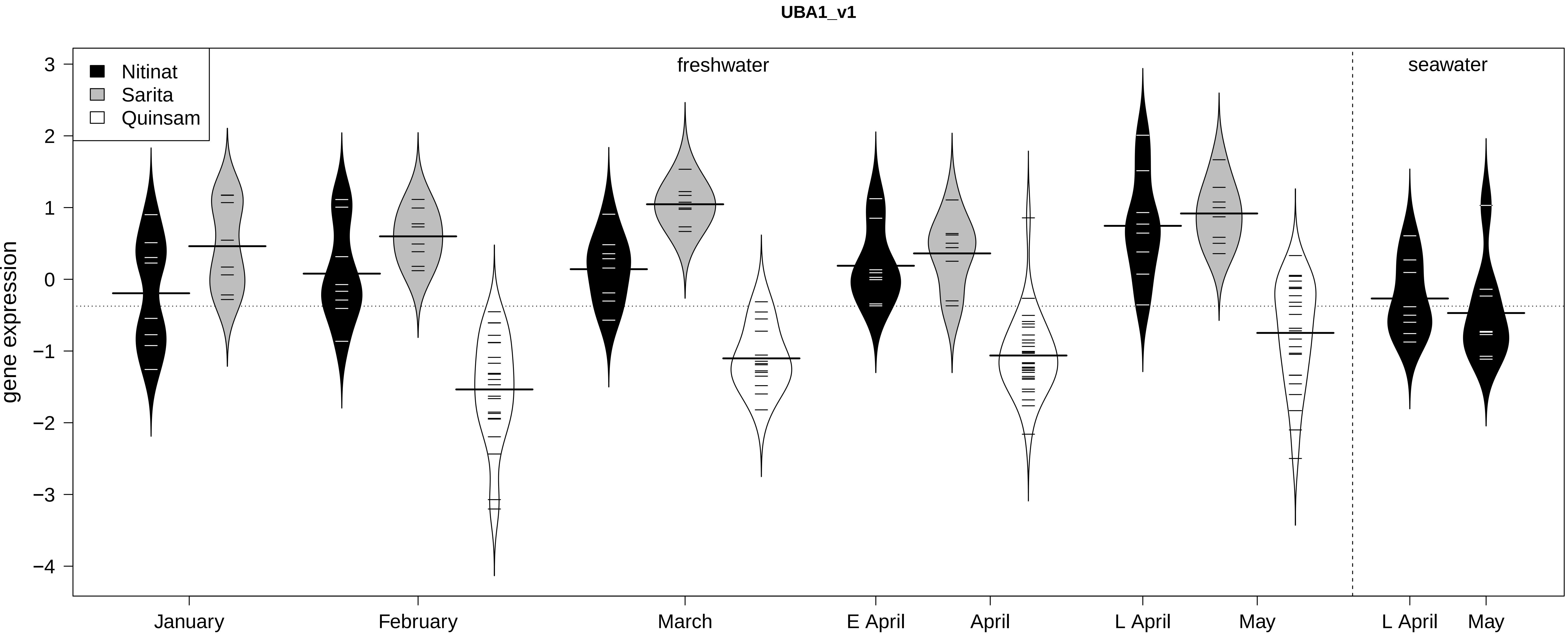

UBA1_v1
Nitinat
Sarita
Quinsam
freshwater
seawater
gene expression
3
2
1
0
−1
−2
−3
−4
January
February
March
E April
April
L April
May
L April
May

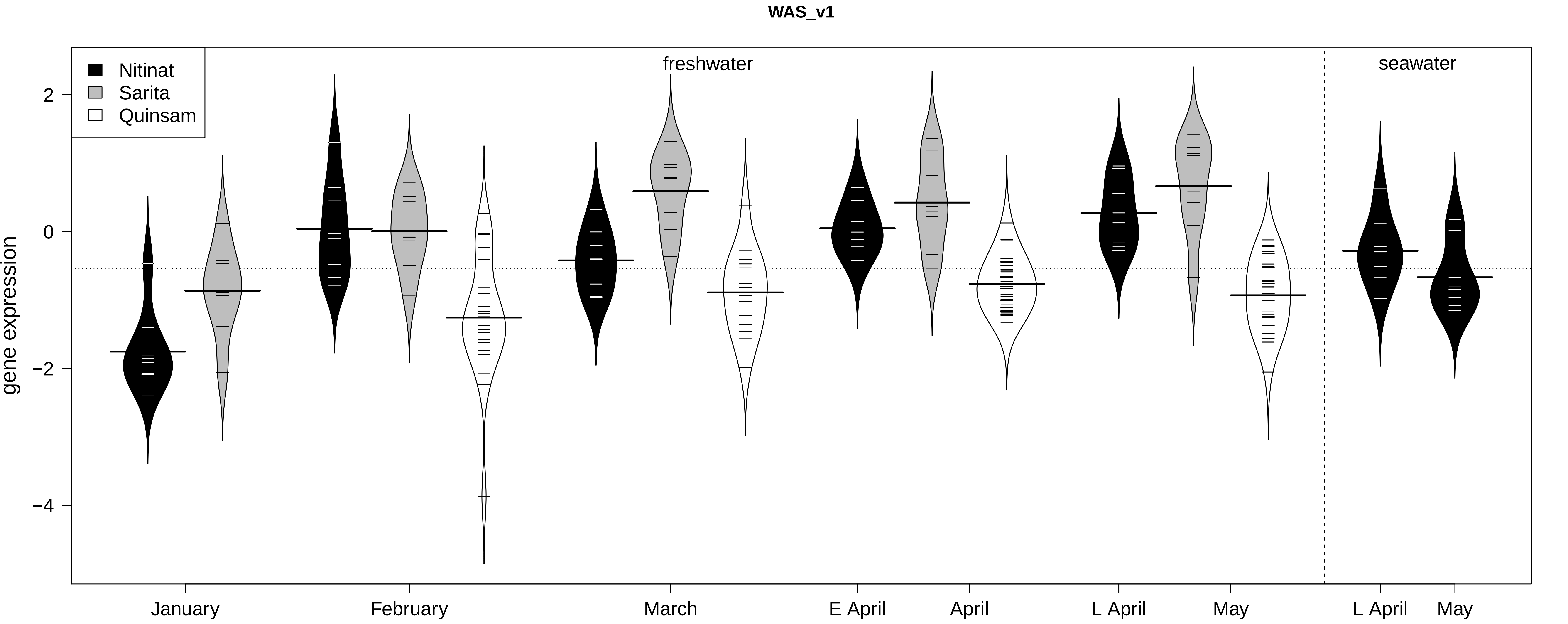

WAS_v1
Nitinat
Sarita
Quinsam
freshwater
seawater
gene expression
2
0
−2
−4
January
February
March
E April
April
L April
May
L April
May
